# Identification of hub genes and key pathways in pediatric Crohn’s disease using next generation sequencing and bioinformatics analysis

**DOI:** 10.1101/2022.07.27.501664

**Authors:** Basavaraj Vastrad, Chanabasayya Vastrad

## Abstract

Pediatric Crohn Disease (CD) also known as inflammatory bowel diseases, affects millions of people all over the world. The aim of this investigation is to identify the key genes in CD and uncover their potential functions. We downloaded the next generation sequencing (NGS) dataset GSE101794 from the Gene Expression Omnibus (GEO) database. The NGS dataset GSE101794 was used to screen differentially expressed genes (DEGs) between samples from patients with CD and healthy controls. Gene ontology (GO) and REACTOME pathway enrichment analyses were applied for the DEGs. Subsequently, a protein - protein interaction (PPI) network, modules, miRNA- hub gene regulatory network and TF - hub gene regulatory network were constructed to identify hub genes, miRNAs and TFs. Receiver operating characteristic curve (ROC) analysis was applied to validate the hub genes. A total of 957 DEGs were identified, including 478 up regulated genes and 479 down regulated genes. GO and REACTOME results suggested that several Go terms and pathways are involved in response to stimulus, extracellular region, signaling receptor binding, small molecule metabolic process, membrane, transporter activity, immune system and biological oxidations. The top centrality hub genes MDFI, MNDA, FBXO6, TFRC, STAT1, DPP4, MME, SLC39A4, APOA1 and TMEM25 were screened out as the critical genes among the DEGs from the PPI network, modules, miRNA-hub gene regulatory network and TF-hub gene regulatory network. This investigation identified key genes and signal pathways, which might help us improve our understanding of the molecular mechanisms of CD and identify some novel therapeutic targets for CD.

## Introduction

Pediatric Crohn Disease (CD) is an group of inflammatory bowel diseases and characterized by transmural inflammation in gastrointestinal tract [1]. CD predominately affects before the age of 18, however 25% of cases are diagnosed [2]. The prevalence of CD has been growing steadily in both developed and developing nations [3]. The pathogenesis of CD remains incompletely understood, but genetics factors, epigenetic factors, microbial exposure, immune response and environment factors are believed to contribute [4]. Moreover, CD is commonly associated with other complications such as anemia [5], autoimmune liver disease [6], type 1 diabetes mellitus [7], coagulation and fibrinolysis [8] and colorectal cancer [9]. The etiology of CD has been investigated extensively, but the exact pathogenic factors or triggering agents for CD are still unknown, and the underlying molecular mechanisms for induction and advancement of CD remain largely unidentified. Therefore, the discovery of effective biomarkers for the treatment of CD is very essential.

Accumulating evidence had shows that genes [10] and signaling pathways [11] mainly contribute to the occurrence and advancement of CD. Genes include MDR1 [12], ACE2 [13], NUDT15 [14], HNF4A [15] and IL23R [16] are responsible for progression of CD. Signaling pathways include JAK/STAT signaling pathway [17], STING signaling pathway [18], TLRs and dectin-1 signaling pathways [19], NF□κB and MAPK signaling pathways [20] and P2X7R-Pannexin-1 signaling pathway [21] are involved in advancement of CD. However, the prevalence and factors responsible for CD etiology are still not fully known.

Next generation sequencing (NGS) is the latest technology, which can detect multiple genes at the same time and minimize system errors and has extremely high sensitivity [22]. At present situation, this technology has been widely used in the investigation of various diseases, which has opened up a milestone in the field of genomics research on CD [23]. In the process of this investigation on CD, based on this technology, we can find and analyze the gene expression of CD at the molecular level.

In this investigation, we downloaded NGS dataset GSE101794 [24] based on the Gene Expression Omnibus database (GEO, https://www.ncbi.nlm.nih.gov/geo/) [25]. First, differentially expressed genes (DEGs) were analyzed from the samples by limma. These underwent gene ontology (GO) and REACTOME pathway enrichment analyses, followed by the construction of a protein-protein interaction (PPI) network, modules, miRNA-hub gene regulatory network and TF-hub gene regulatory network to identify hub genes. Receiver operating characteristic curve (ROC) analysis was conducted to validate the hub genes, which could be used as molecular biomarkers or diagnostic targets for CD therapy. Collectively, our investigation will help the advancement of a genetic diagnosis for CD and more effective measures of prevention and treatment.

## Materials and Methods

### Next generation sequencing data source

NGS data was downloaded from the GEO database. GSE101794 [24] includes 254 CD samples and 50 healthy control samples. This dataset was obtained from the NGS platform of GPL11154 Illumina HiSeq 2000 (Homo sapiens).

### Identification of DEGs

As a fully functional package, the limma package [26] in R software includes the original data input of NGS, as well as a liner model for analyzing differentially expressed genes. We screened DEGs between CD samples and healthy control samples by utilizing limma package with a adjust p < 0.05, and a log (Fold Change) > 0.822 for up regulated genes and log (Fold Change) < −0.825 for down regulated genes. And the volcano plot and heat map were drawn by ggplot2 package and gplot package in R software.

### GO and pathway enrichment analyses of DEGs

One online tool, g:Profiler (http://biit.cs.ut.ee/gprofiler/) [27], was applied to carried out the functional annotation for DEGs. GO (http://www.geneontology.org) [28] generally perform enrichment analysis of genomes. GO include biological processes (BP), cellular components (CC) and molecular functions (MF) in the GO enrichment analysis. REACTOME (https://reactome.org/) [29] is a comprehensive database of genomic, chemical, and systemic functional information. Therefore, g:Profiler was used to make analysis of GO and REACTOME. P□<□0.05 value was set as the cutoff criterion for significant GO and pathway enrichment.

### Construction of the PPI network and module analysis

Human Integrated Protein-Protein Interaction rEference (HiPPIE) interactome database [30] was used to construct a PPI network for the DEGs, which was visualized in Cytoscape (version 3.9.1) [31]. The Network Analyzer plug-in can be used to screen hub genes with the node degree [32], betweenness [33], stress [34] and closeness [35]. PEWCC1 [36] Plug-in was used to filter key modules in the PPI network with a degree□cutoff ≥ 2, node□score□cutoff = 0.2, K − core ≥ 2, and max.depth = 100 as the cutoff criteria.

### miRNA-hub gene regulatory network construction

The hub genes and miRNA network was generated by miRNet database (https://www.mirnet.ca/) [37]. miRNA-hub gene interaction database: TarBase, miRTarBase, miRecords, miRanda (S mansoni only), miR2Disease, HMDD, PhenomiR, SM2miR, PharmacomiR, EpimiR, starBase, TransmiR, ADmiRE, and TAM 2.0 databases. The analysis results of miRNet was then imported into Cytoscape software (version 3.9.1) [31] for further visualization.

### TF-hub gene regulatory network construction

The hub genes and TF network was generated by NetworkAnalyst database (https://www.networkanalyst.ca/) [38]. TF-hub gene interaction database: JASPAR database. The analysis results of NetworkAnalyst was then imported into Cytoscape software (version 3.9.1) [31] for further visualization.

### Receiver operating characteristic curve (ROC) analysis

ROC curve analysis was performed to evaluate the sensitivity (true positive rate) and specificity (true negative rate) of the hub genes for CD diagnosis and we investigated how large the area under the curve (AUC) was by using the statistical R software package pROC package [39].

## Results

### Identification of DEGs

NGS dataset was obtained from the NCBI GEO database. Using the limma R software tool, DEGs were extracted from the GSE101794 NGS dataset (∣logFC | > 0.822 for up regulated genes, ∣logFC | < −0.825 for down regulated genes and adj. p value < 0.05). As the volcano plots illustrated, NGS data from GSE101794 identified 957 differentially expressed genes with 478 genes up regulated genes and 479 genes down regulated genes in CD samples compared with the expression in healthy control samples (Fig. 1) and are listed in Table 1. The heatmap exhibited the expression difference genes between CD samples and healthy control samples (Fig. 2).

**Fig. 1.**
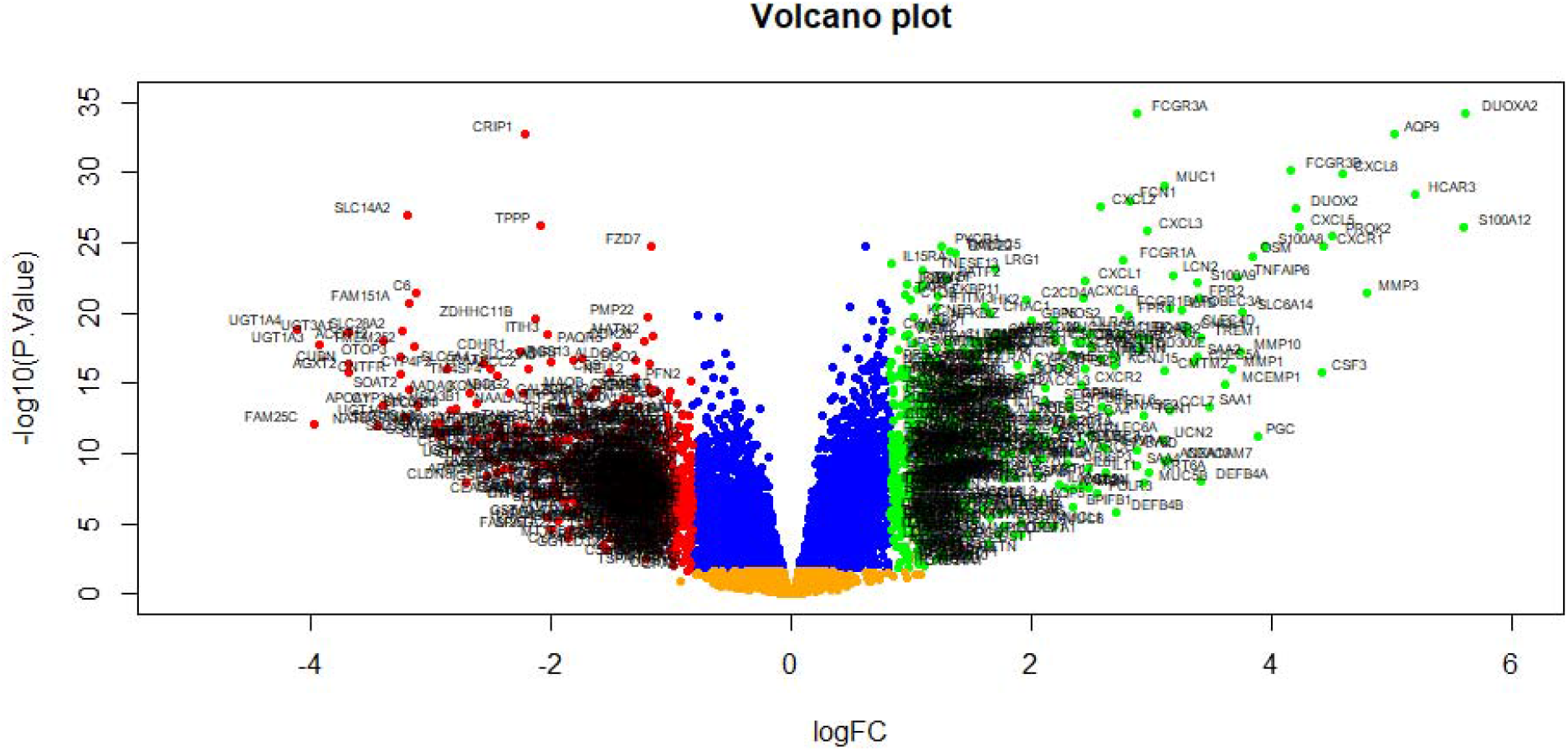
Volcano plot of differentially expressed genes. Genes with a significant change of more than two-fold were selected. Green dot represented up regulated significant genes and red dot represented down regulated significant genes.

**Table 1.**
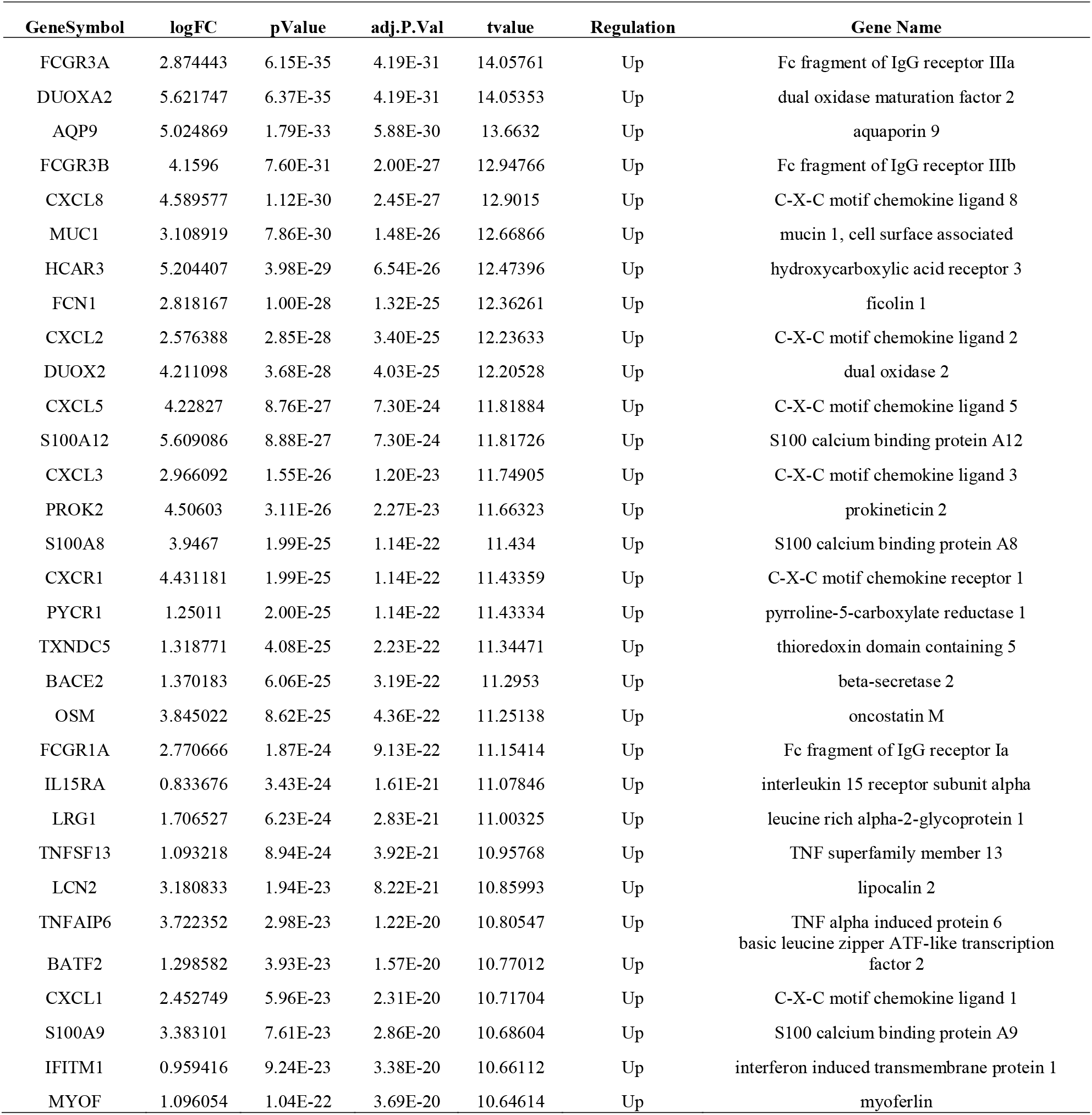
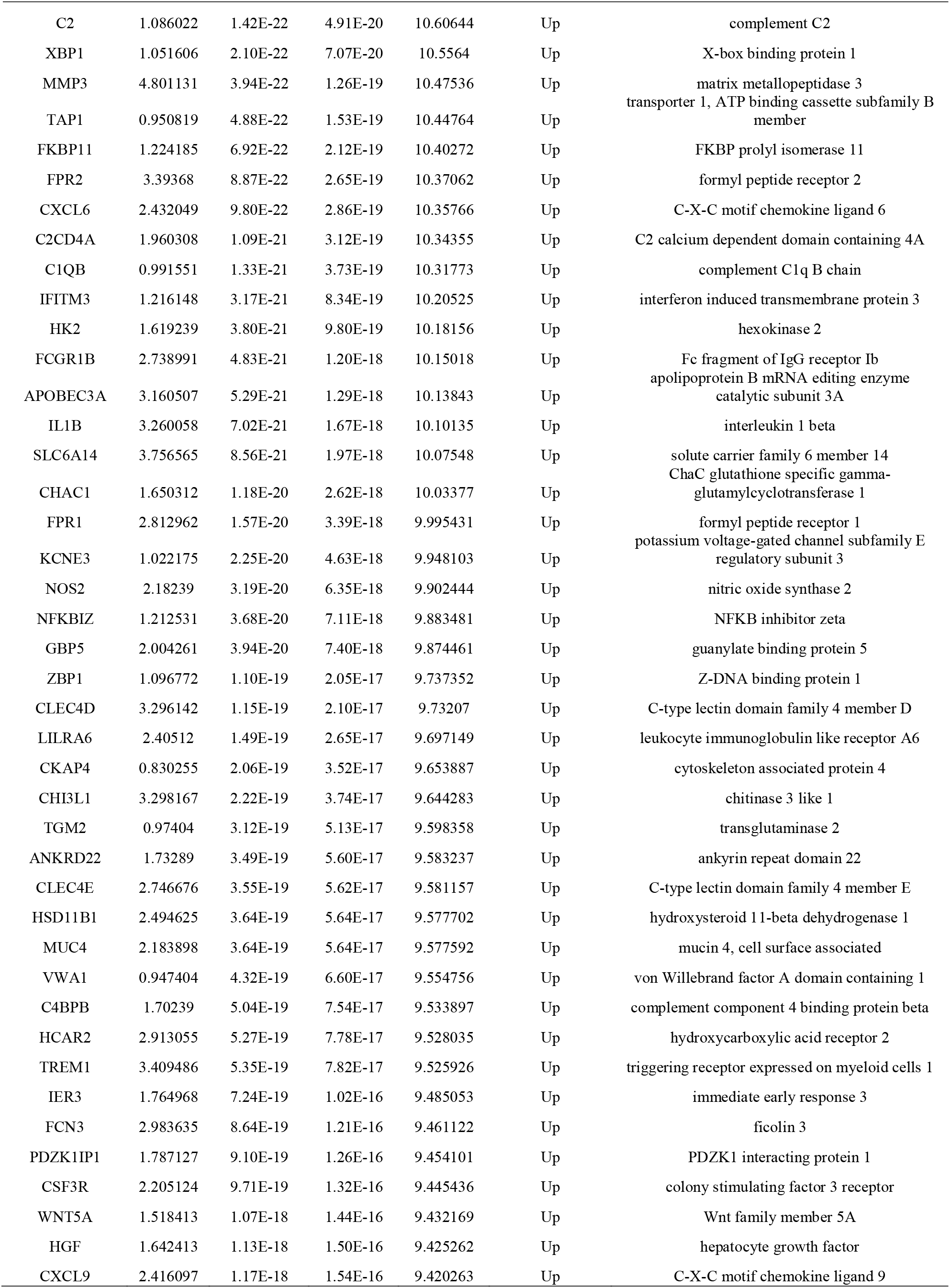
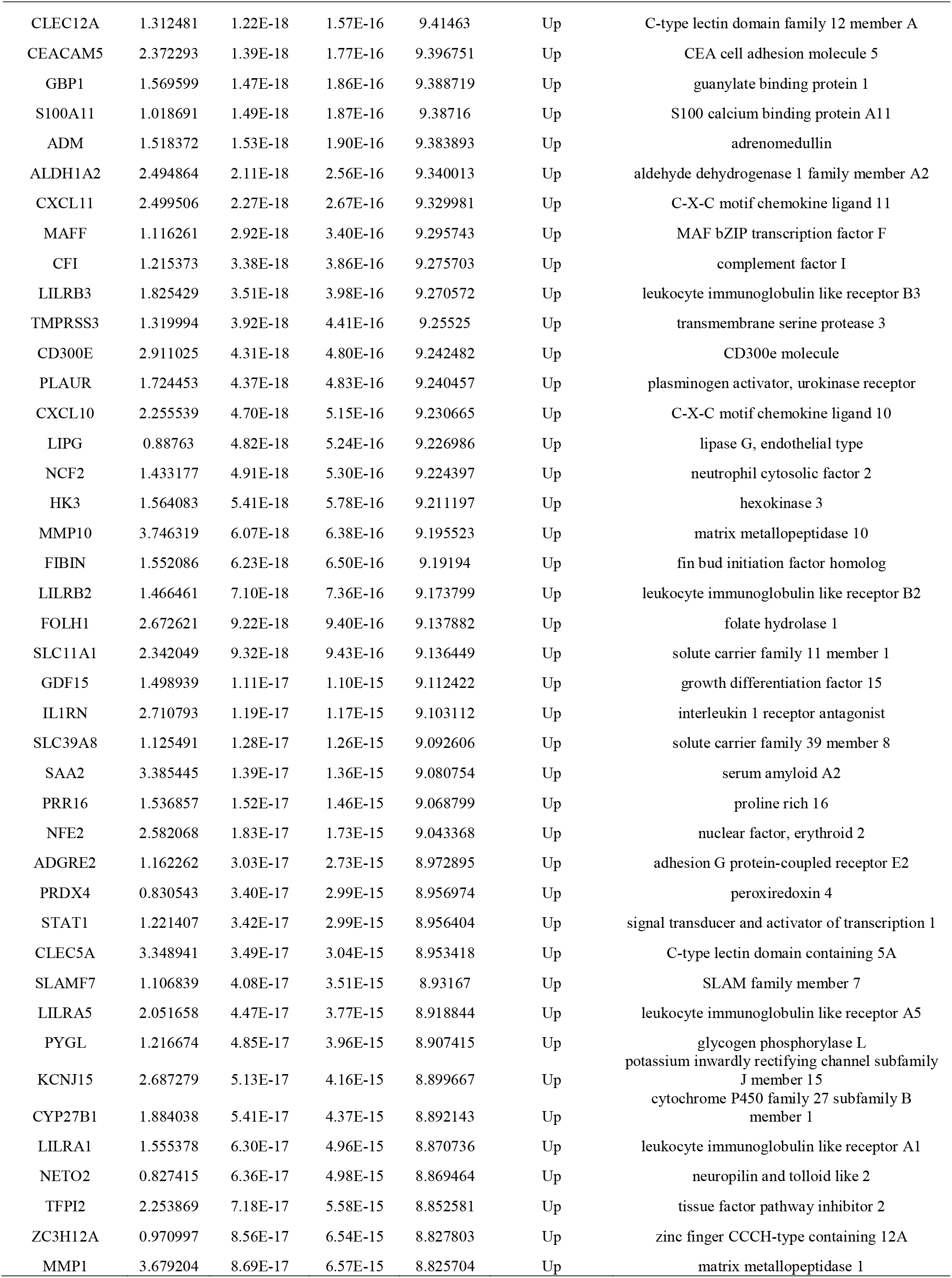
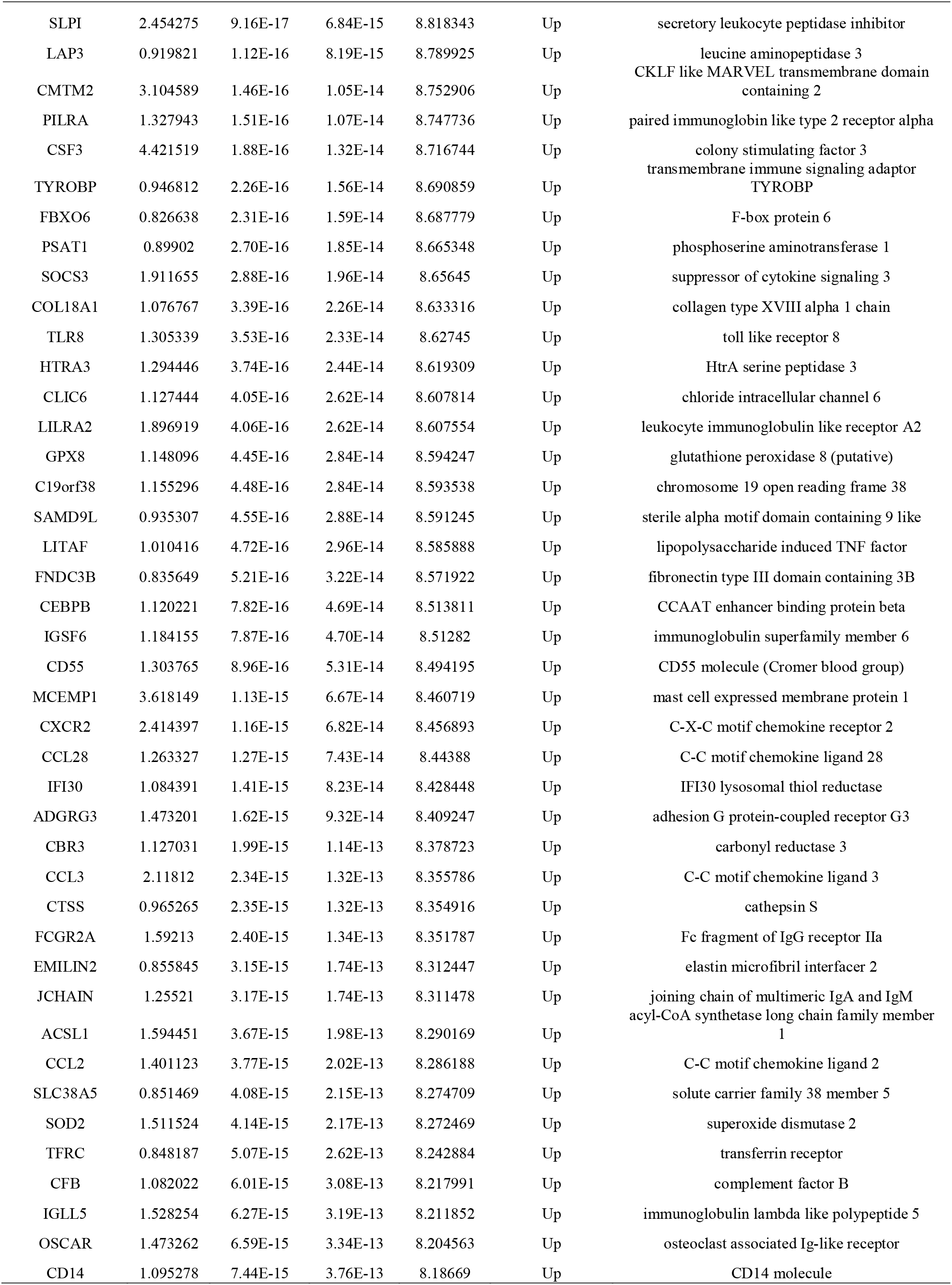
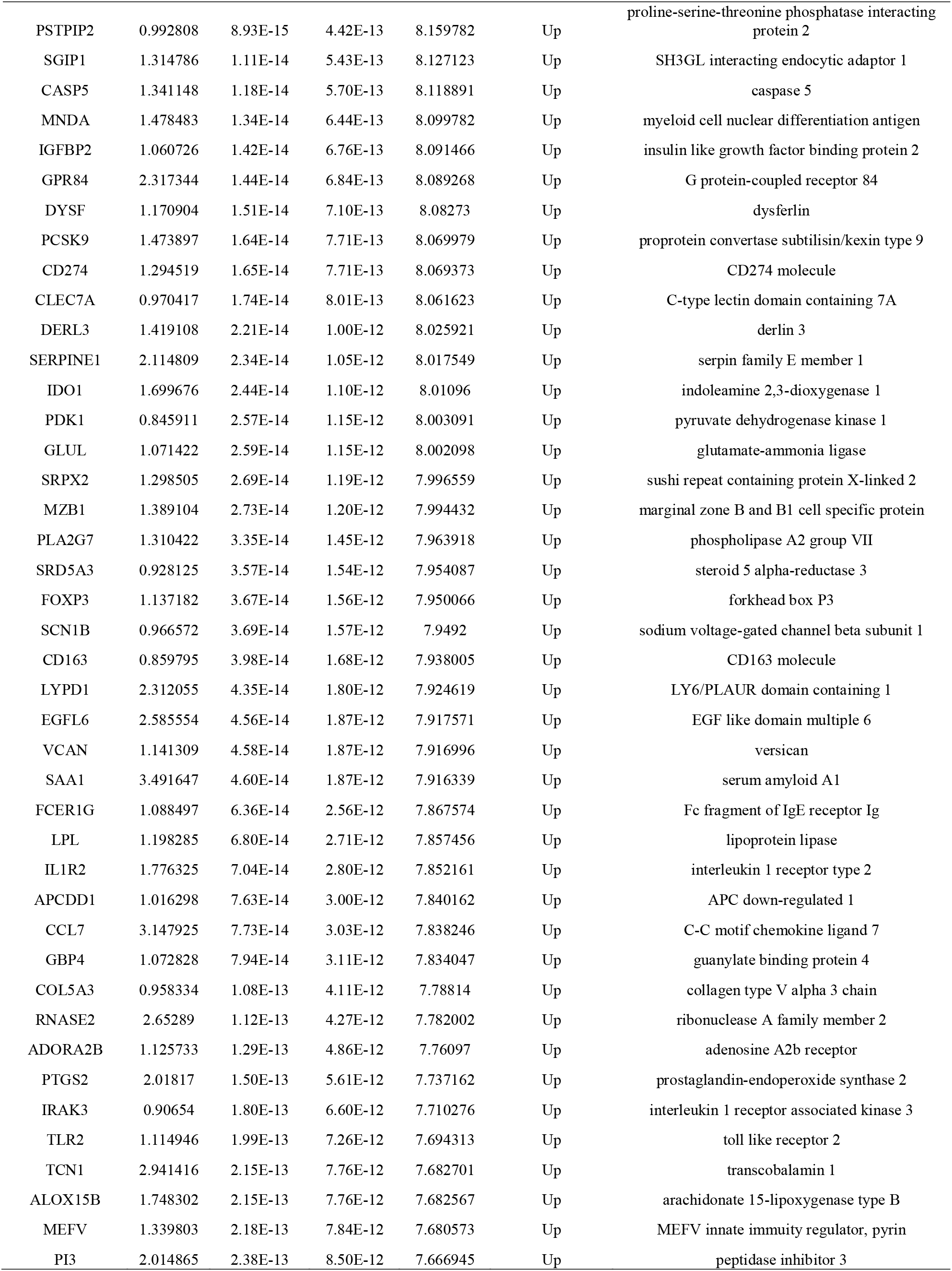
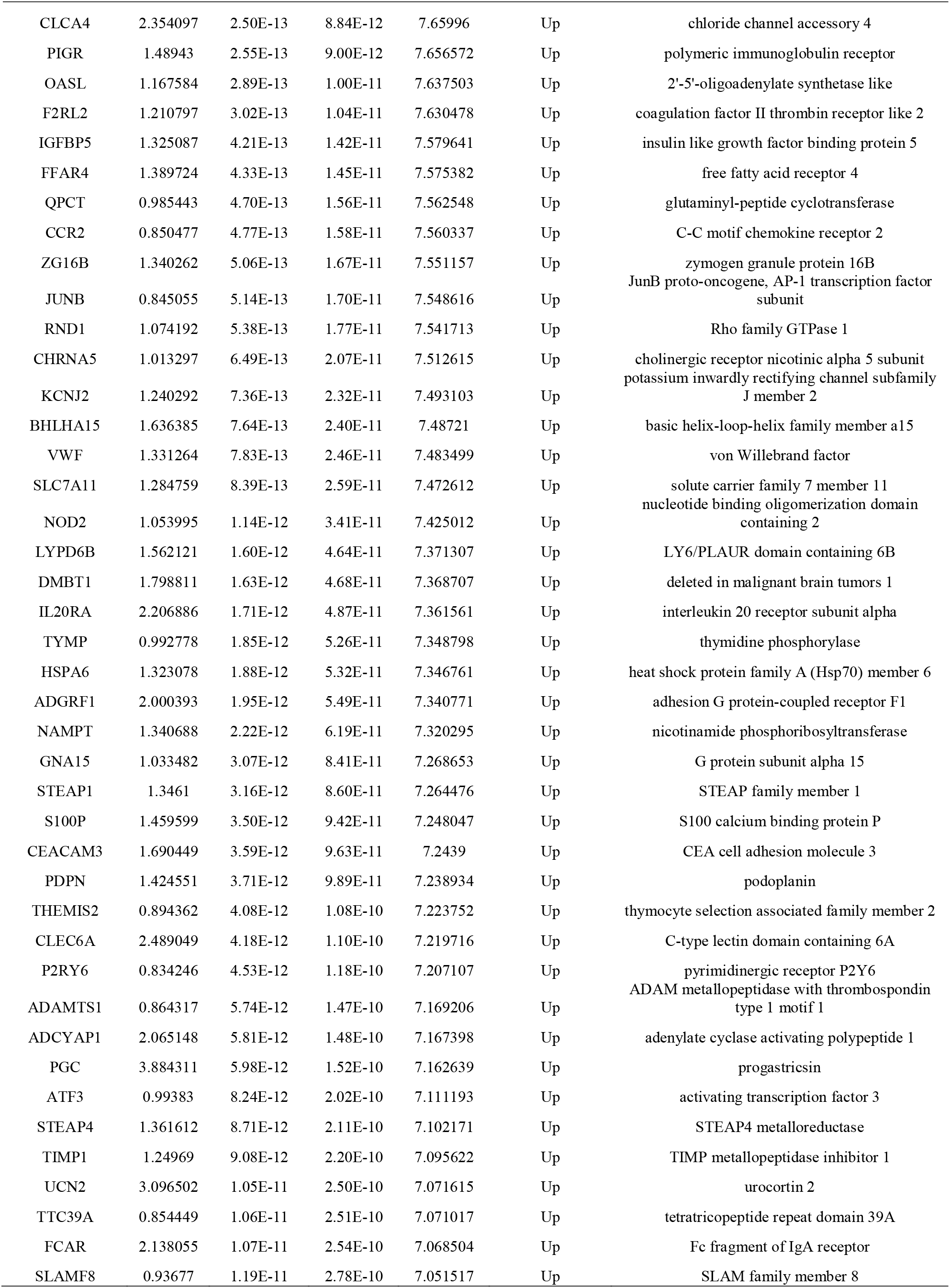
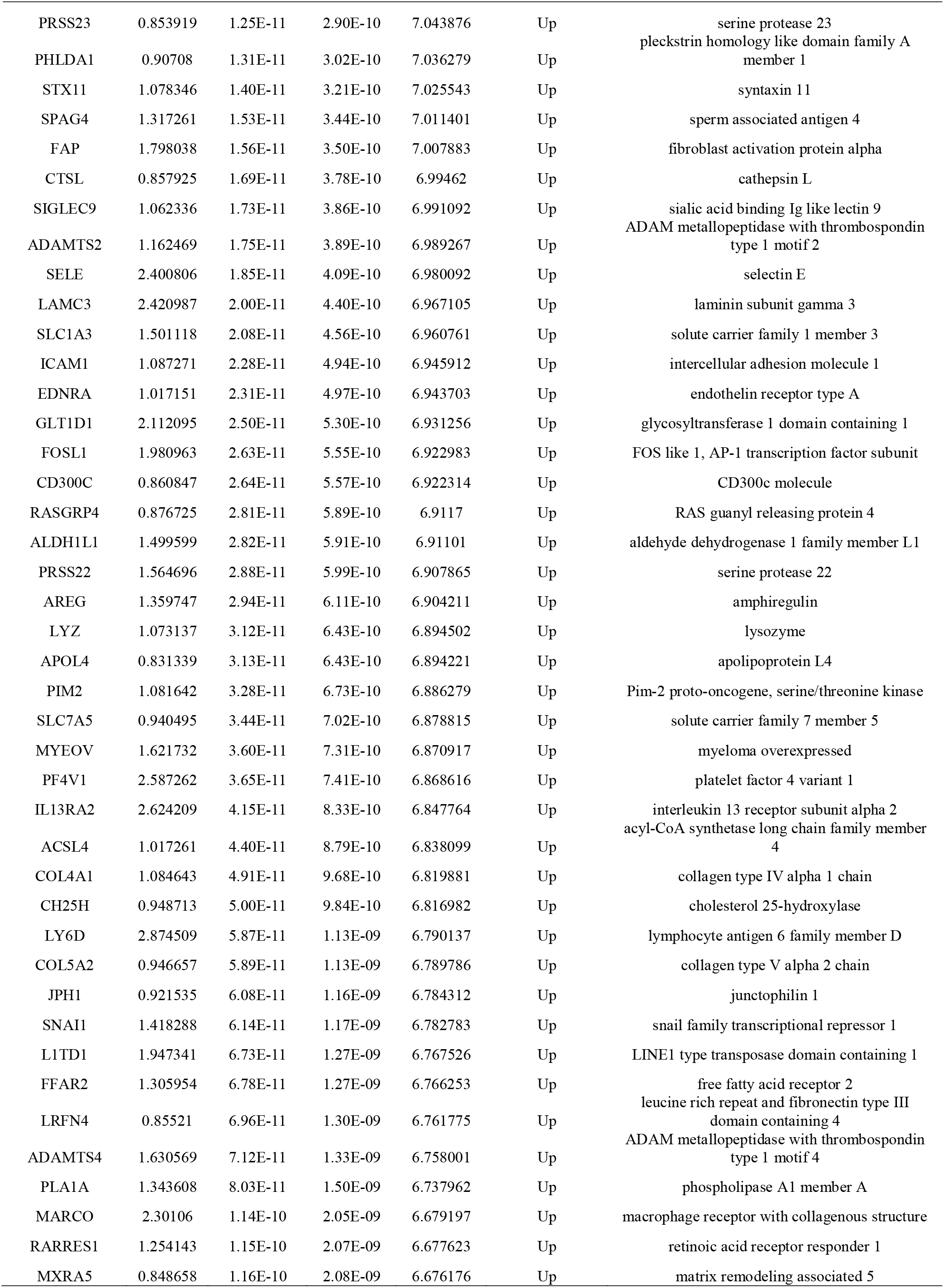
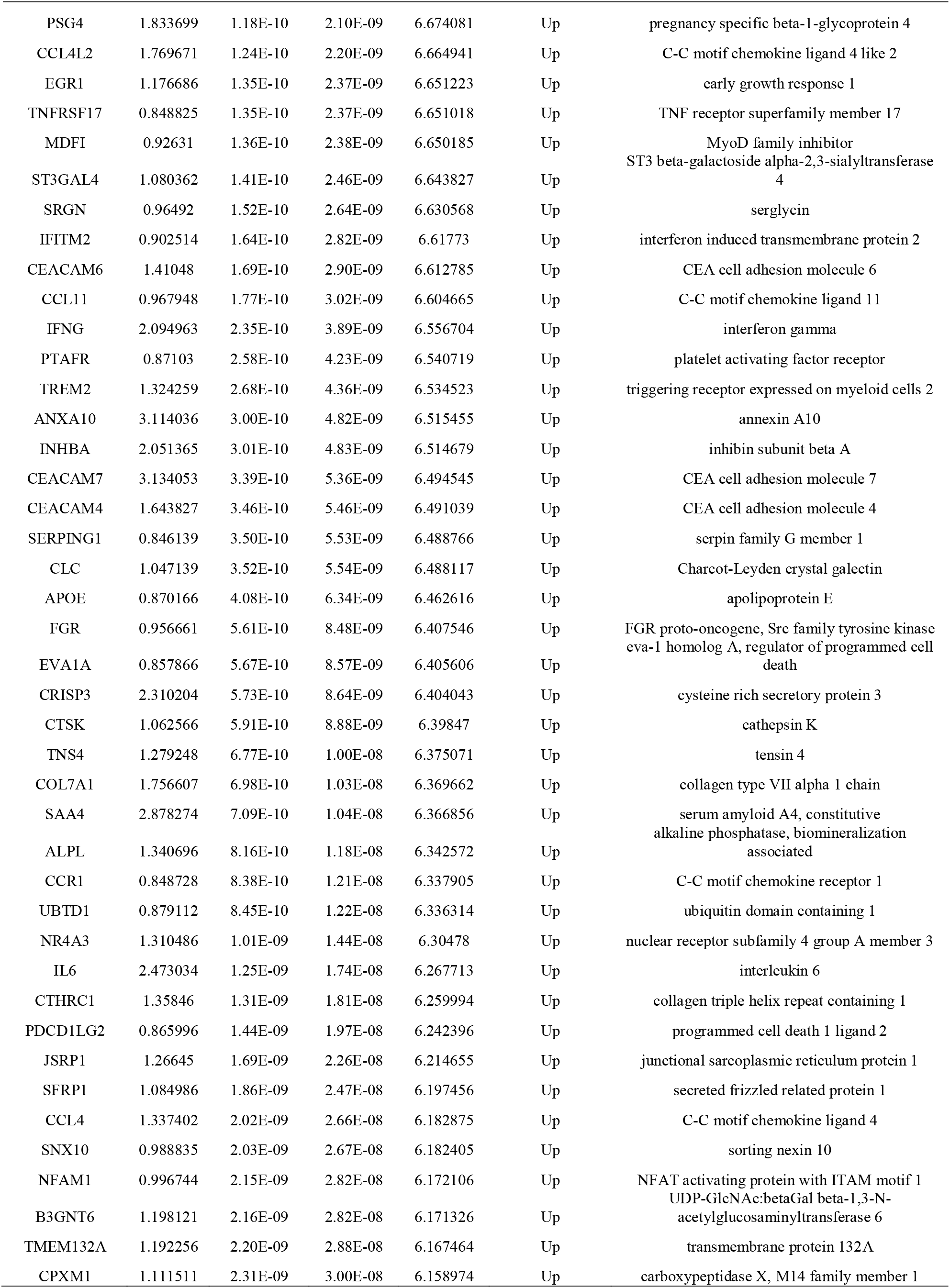
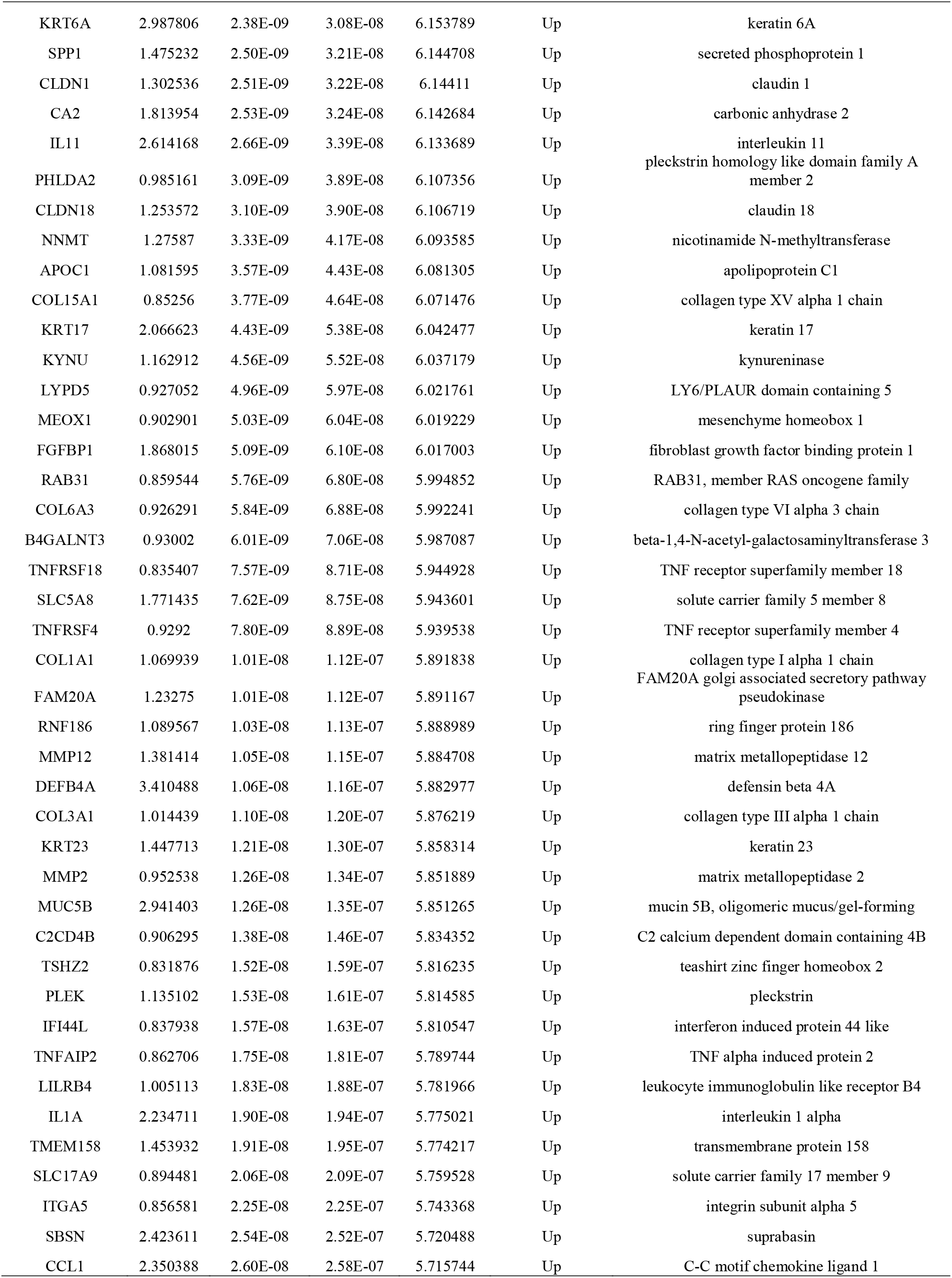
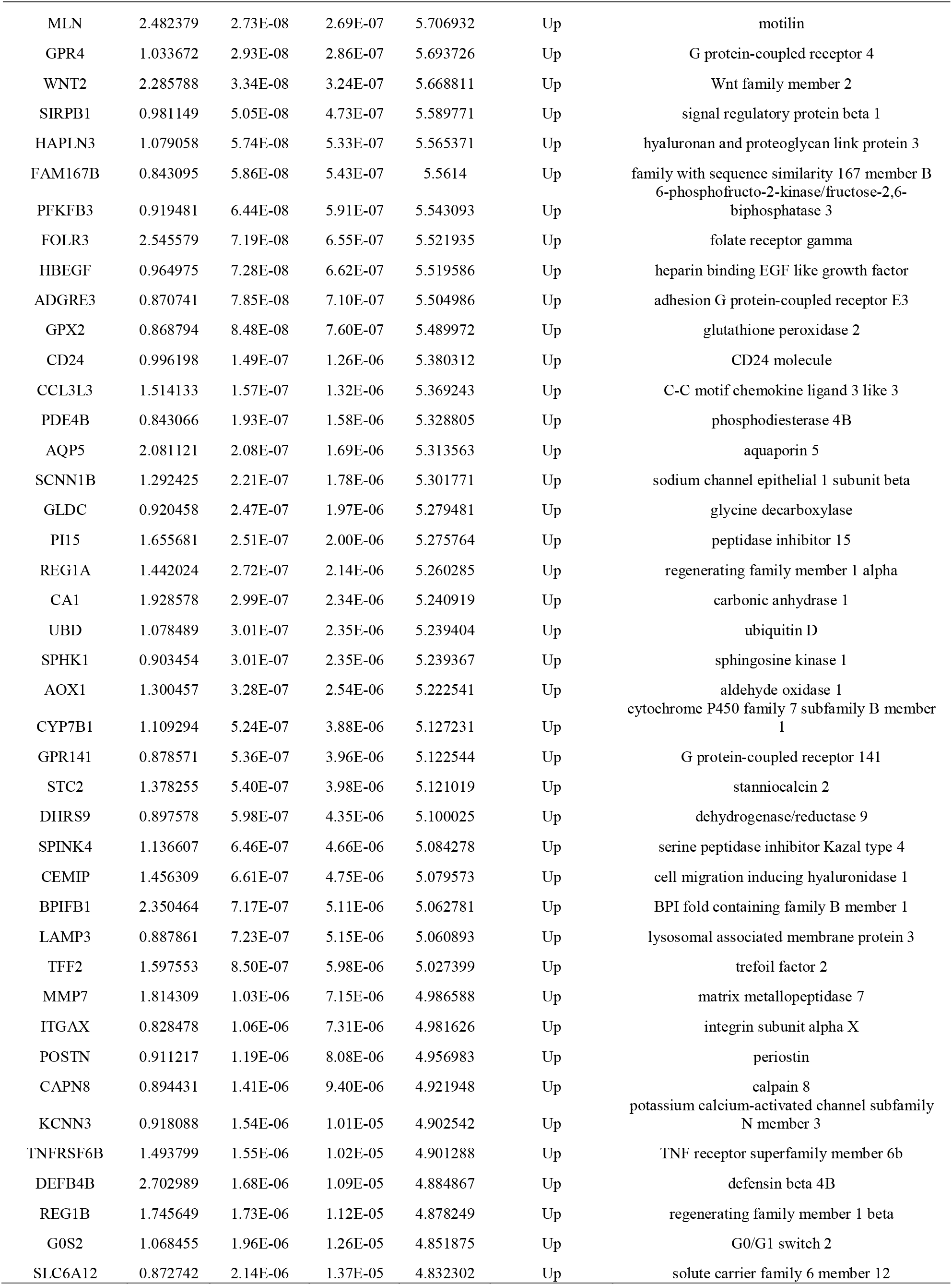
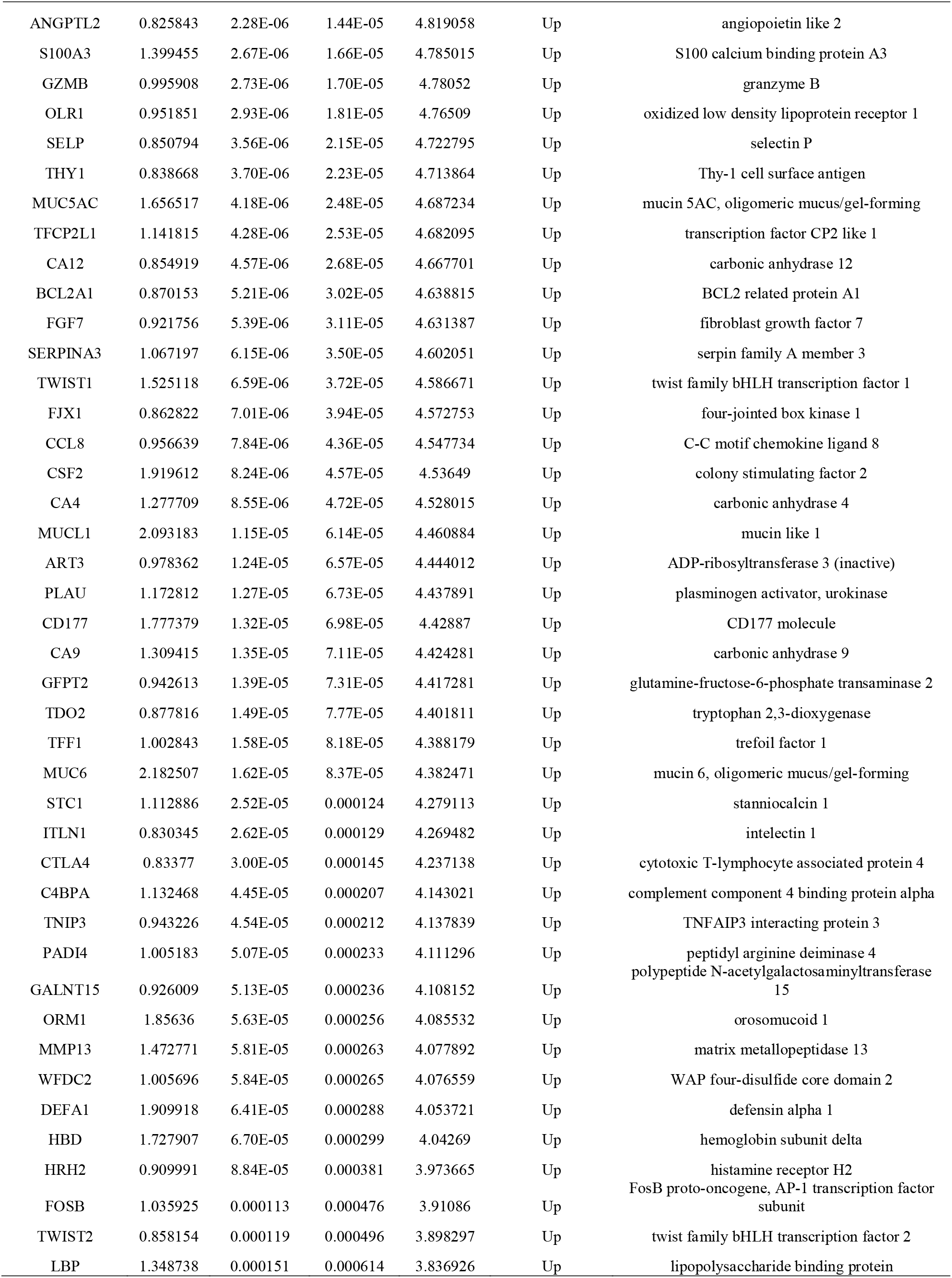
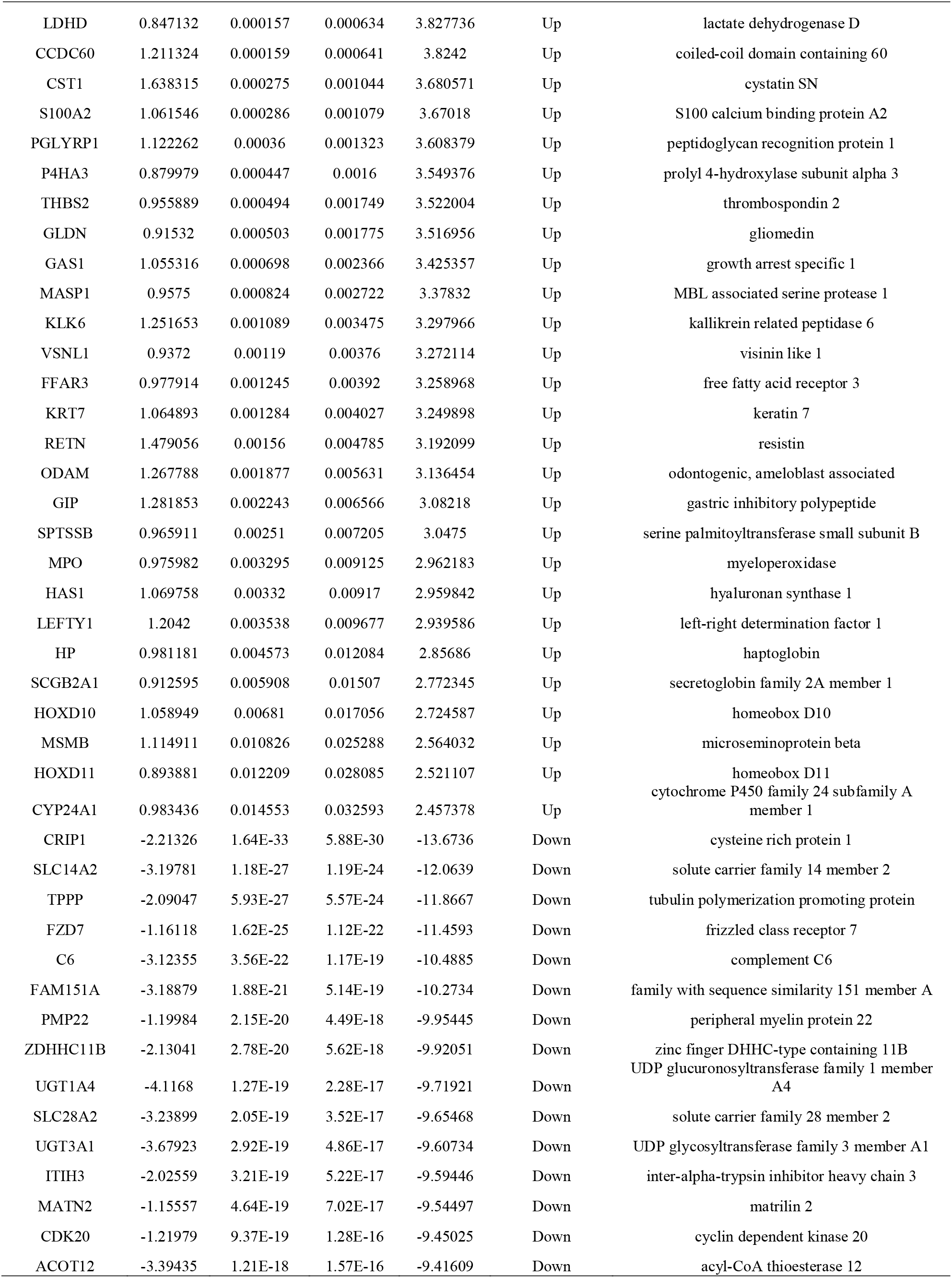
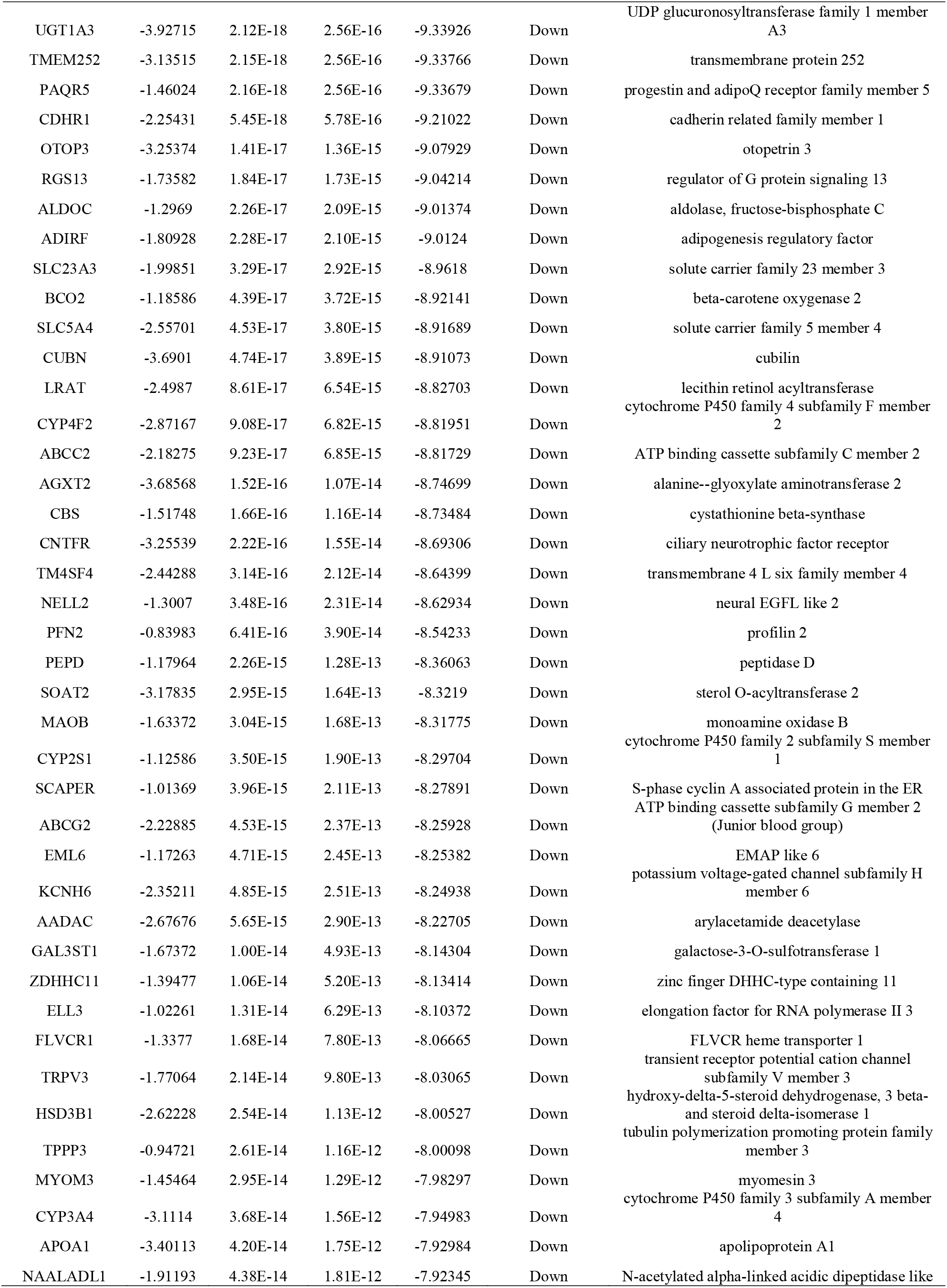
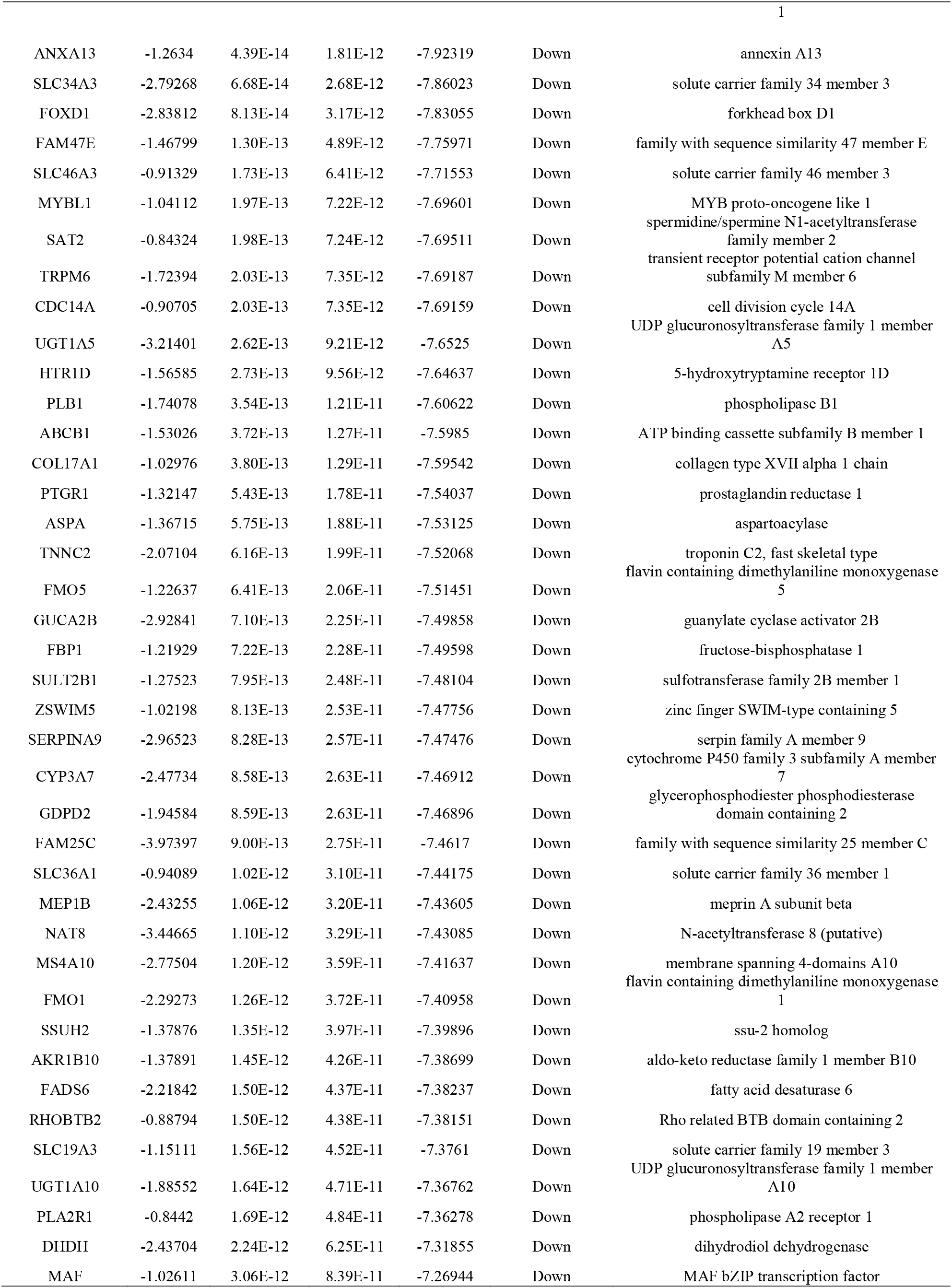
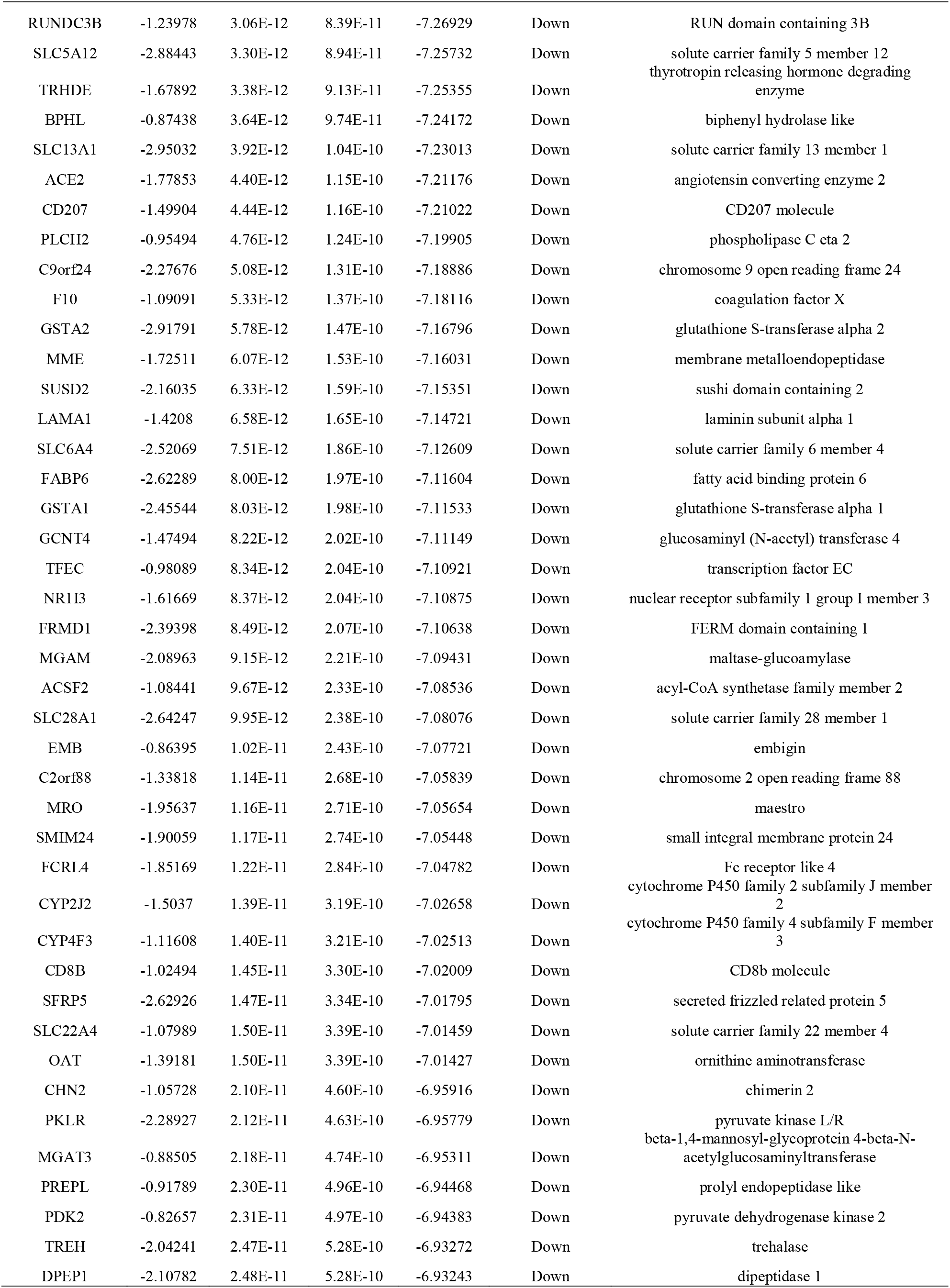
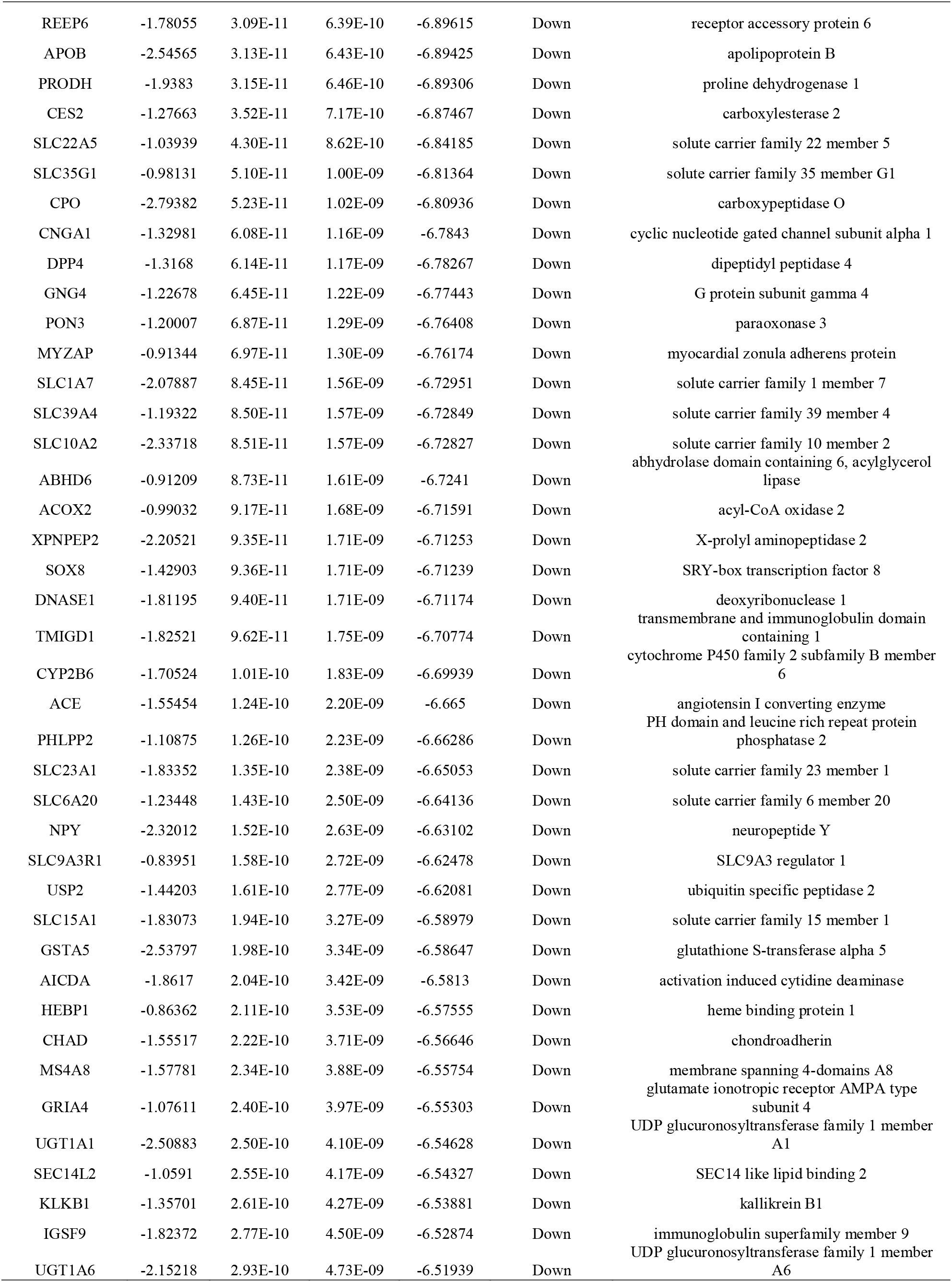
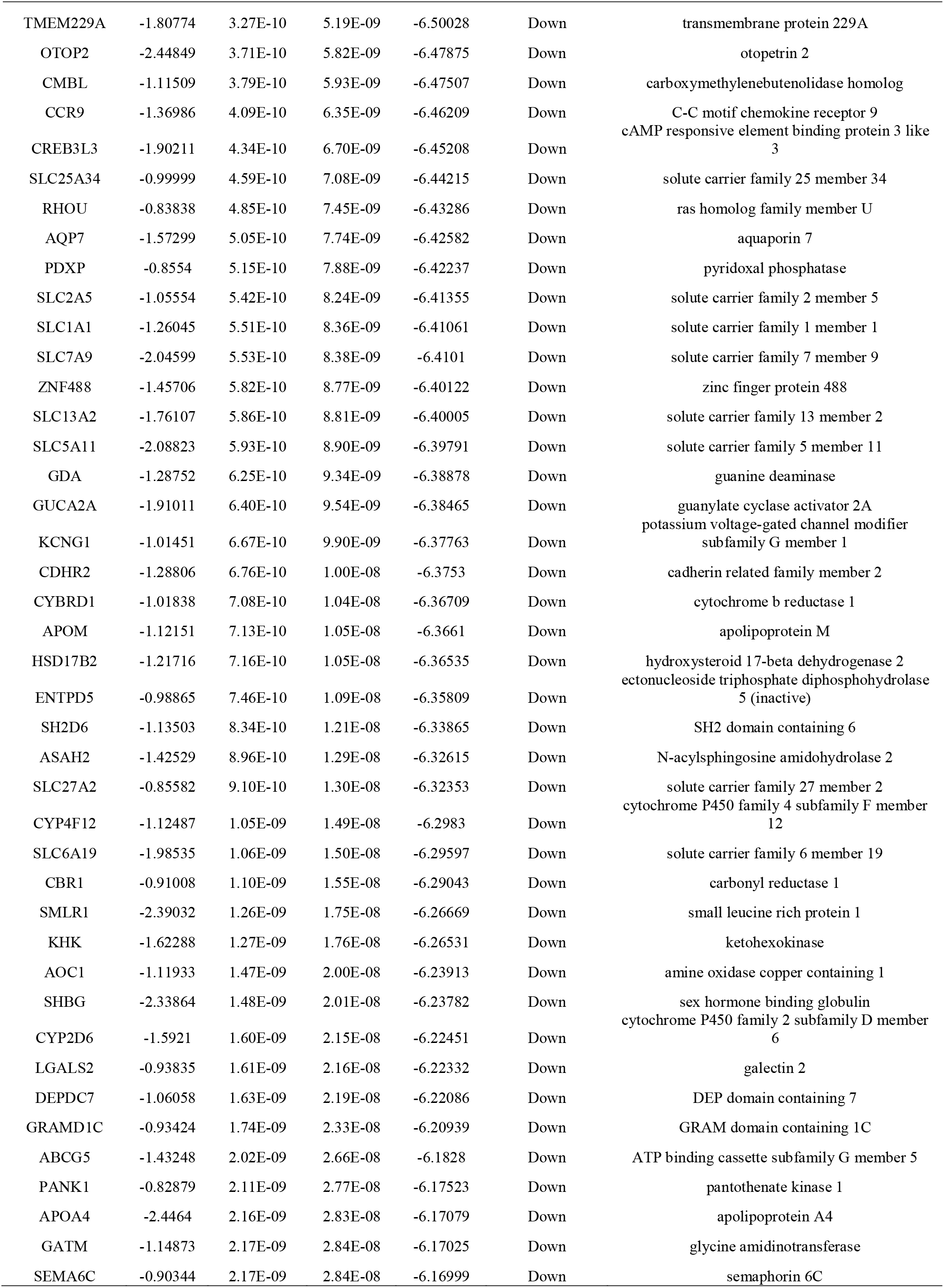
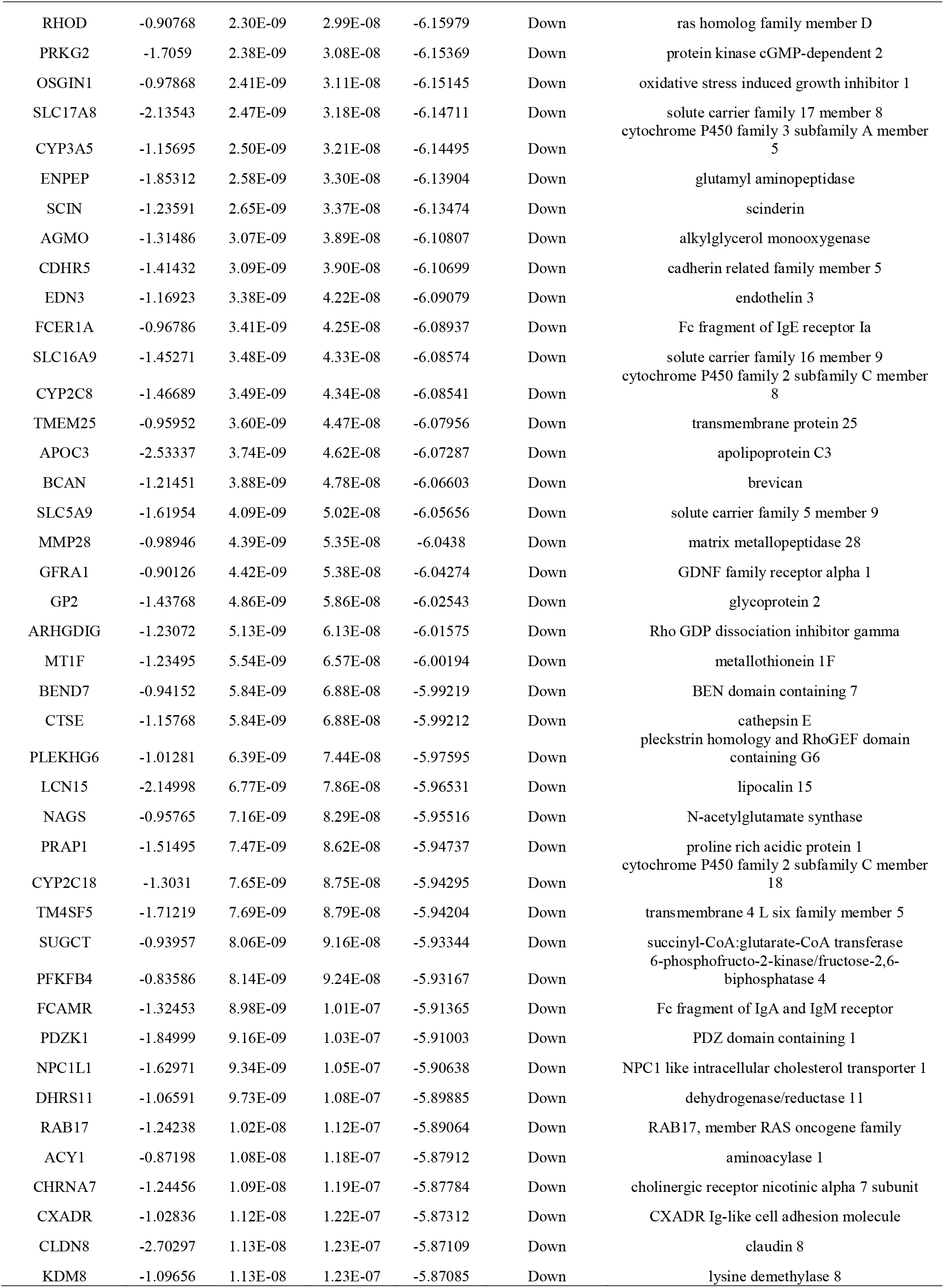
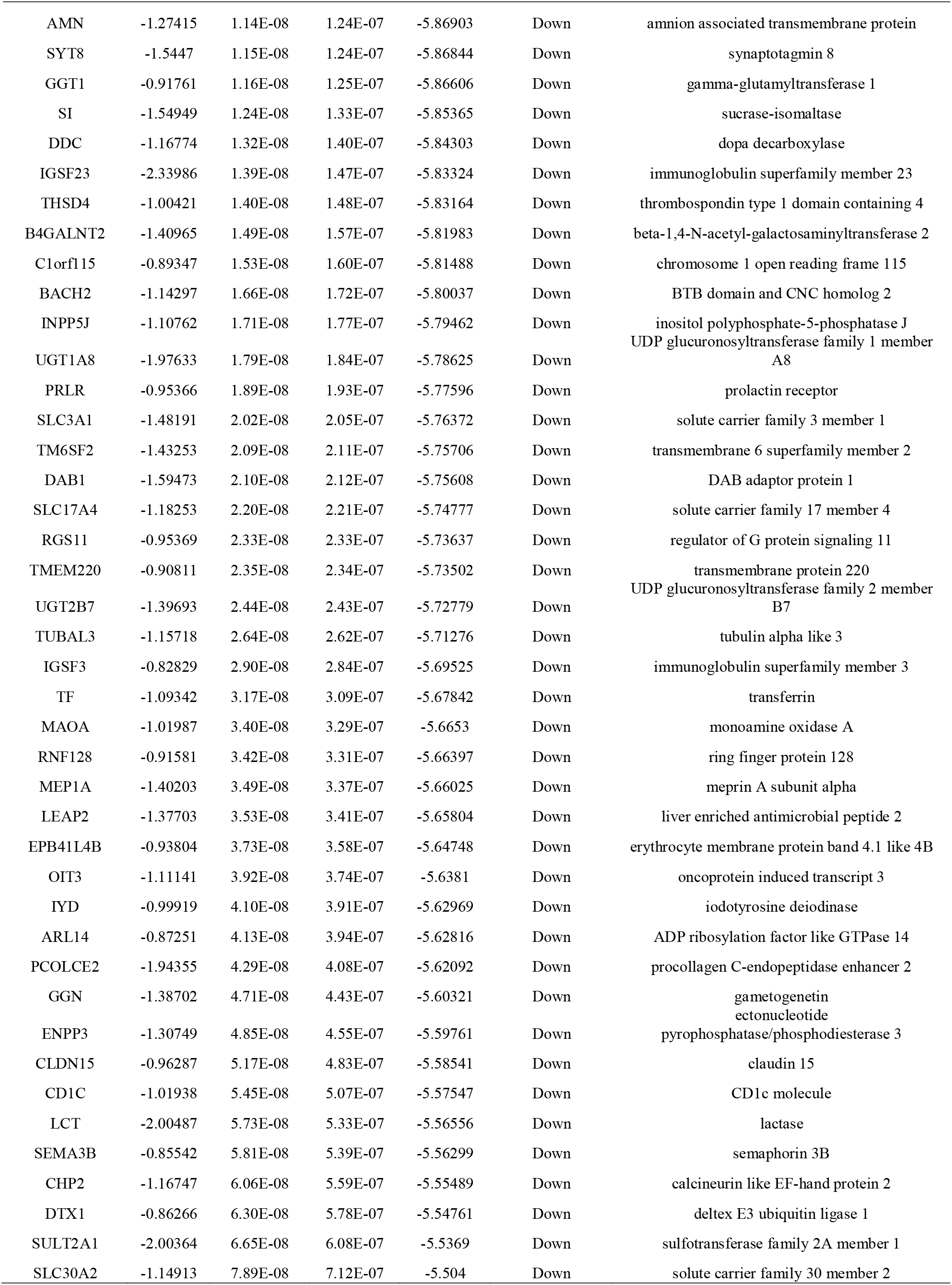
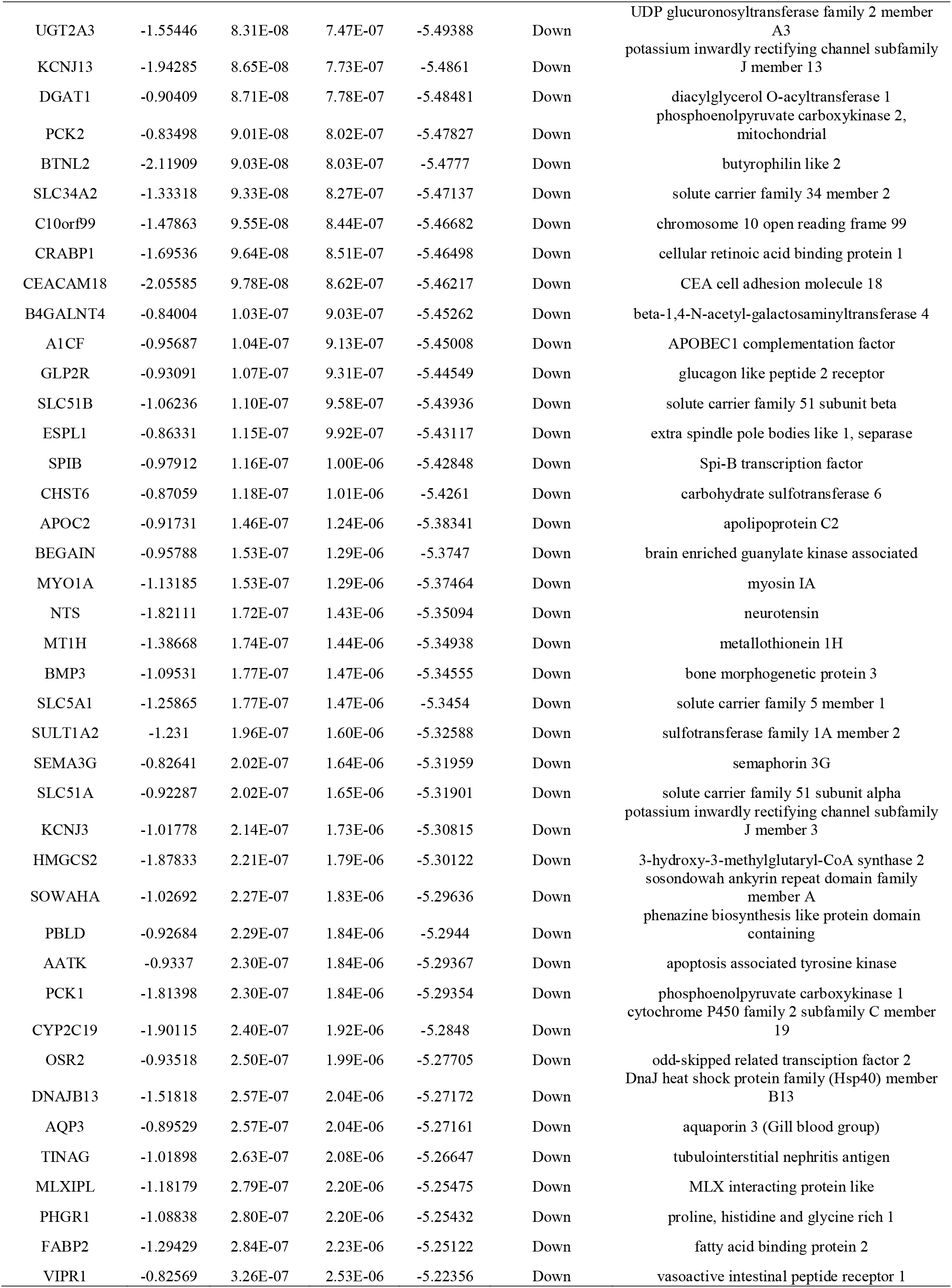
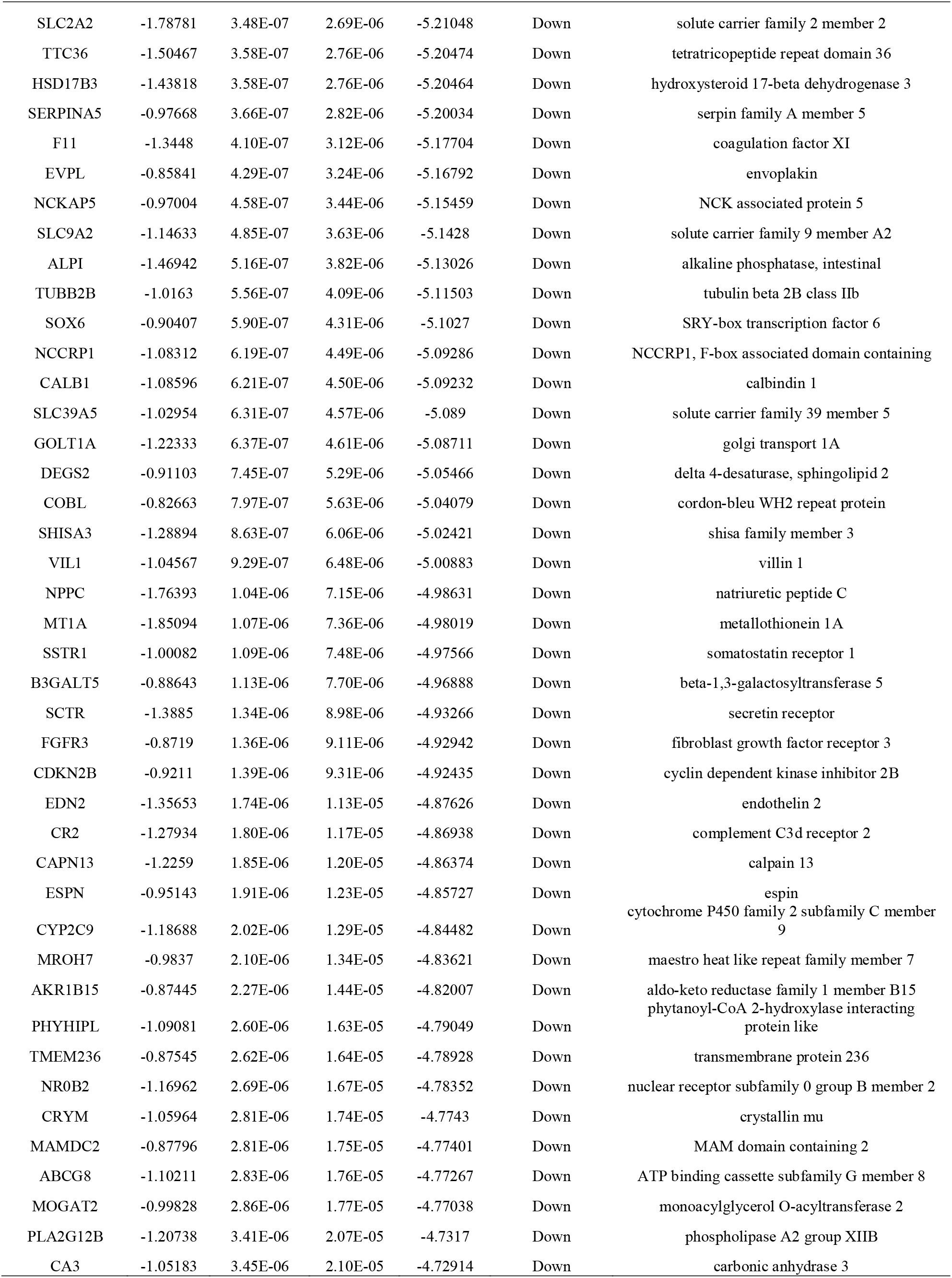
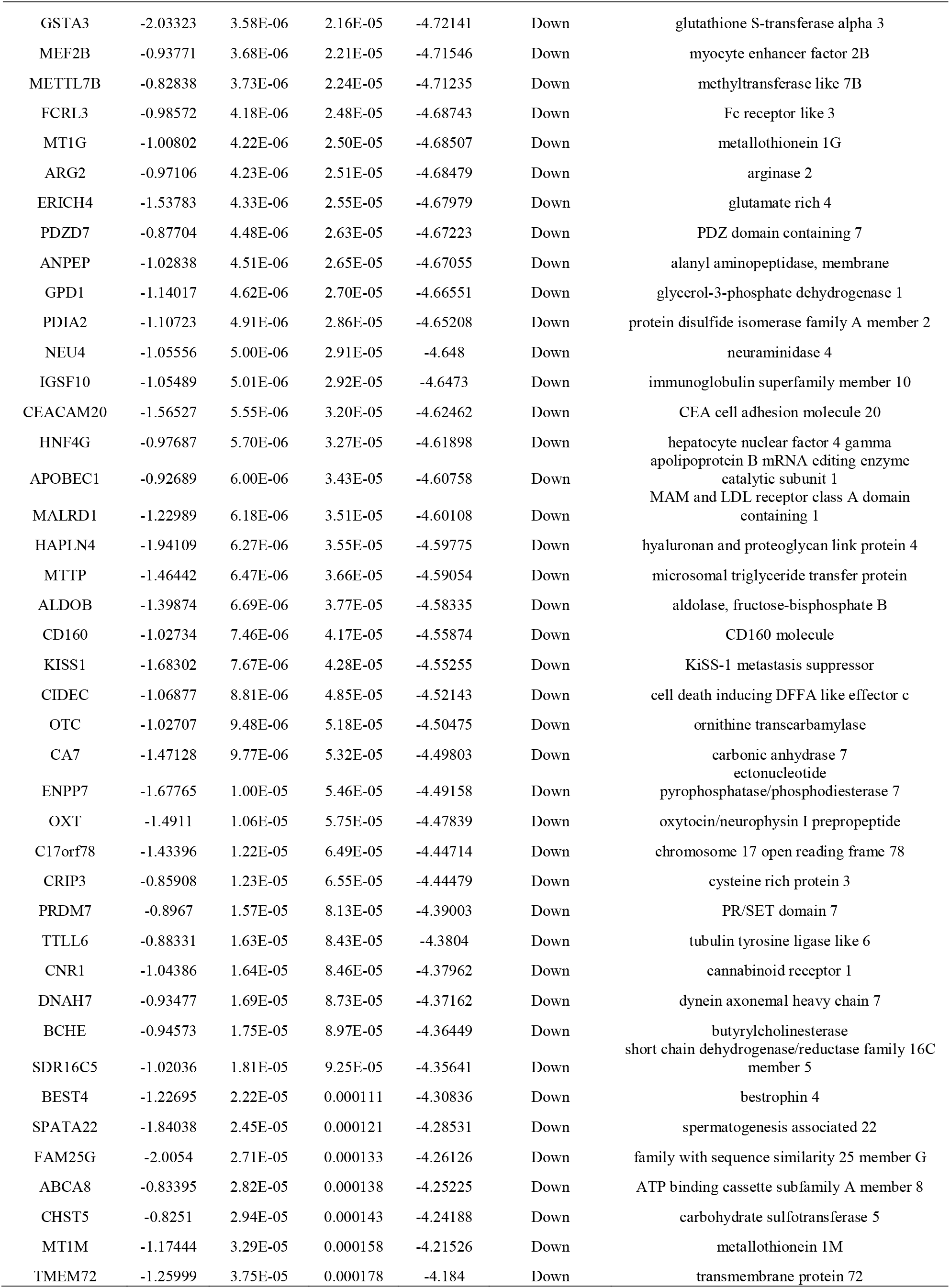
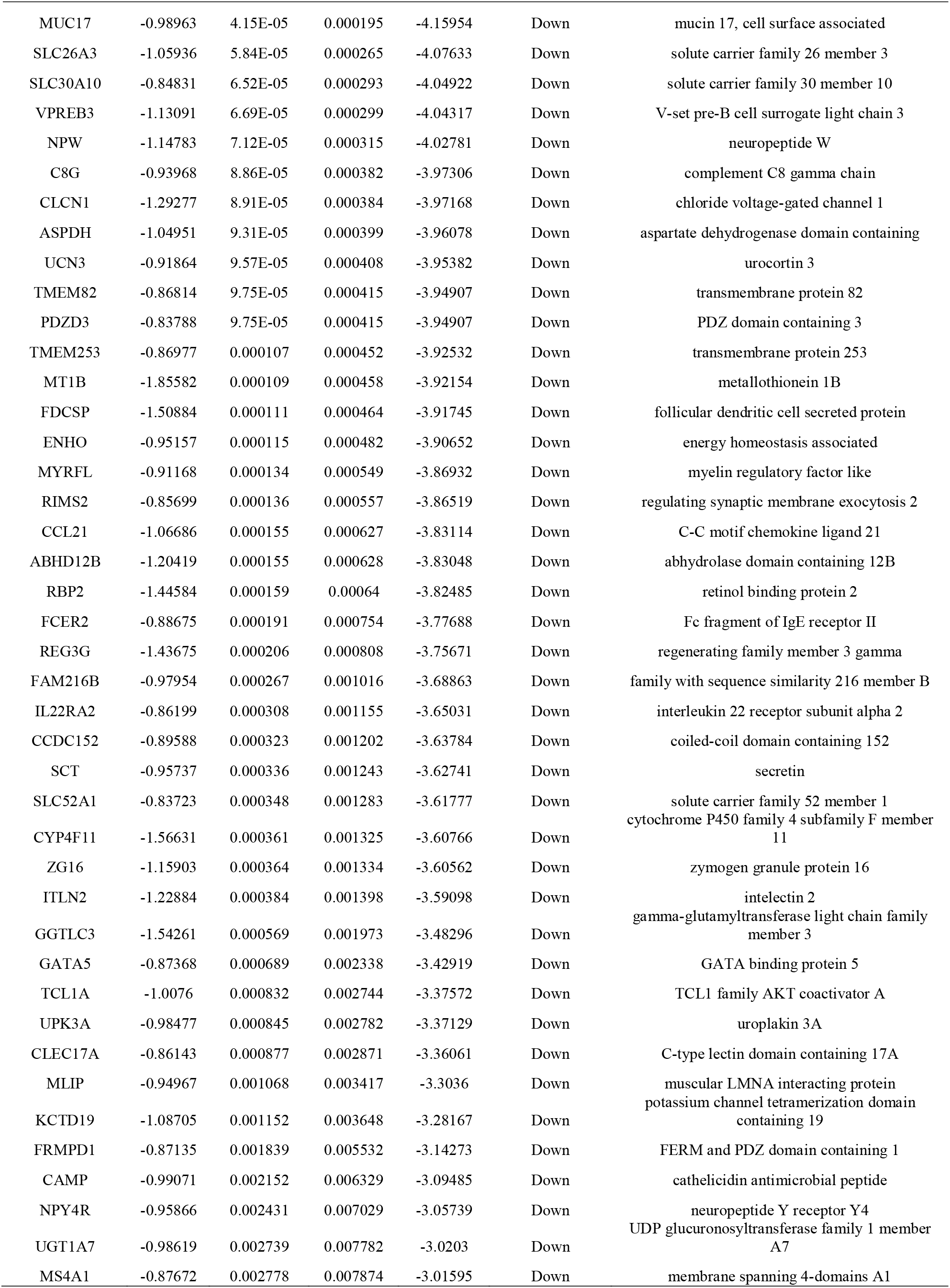
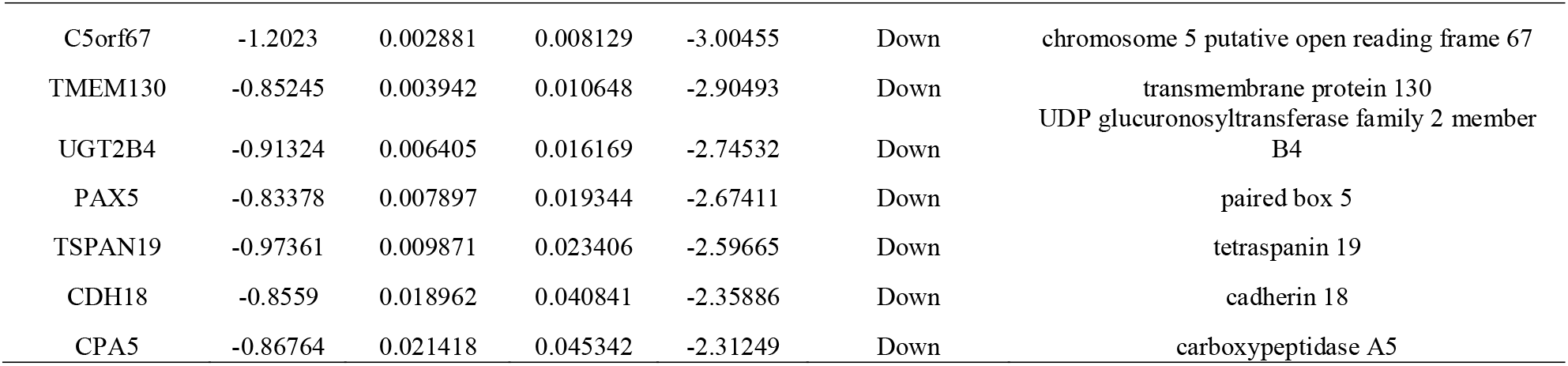
The statistical metrics for key differentially expressed genes (DEGs)

**Fig. 2.**
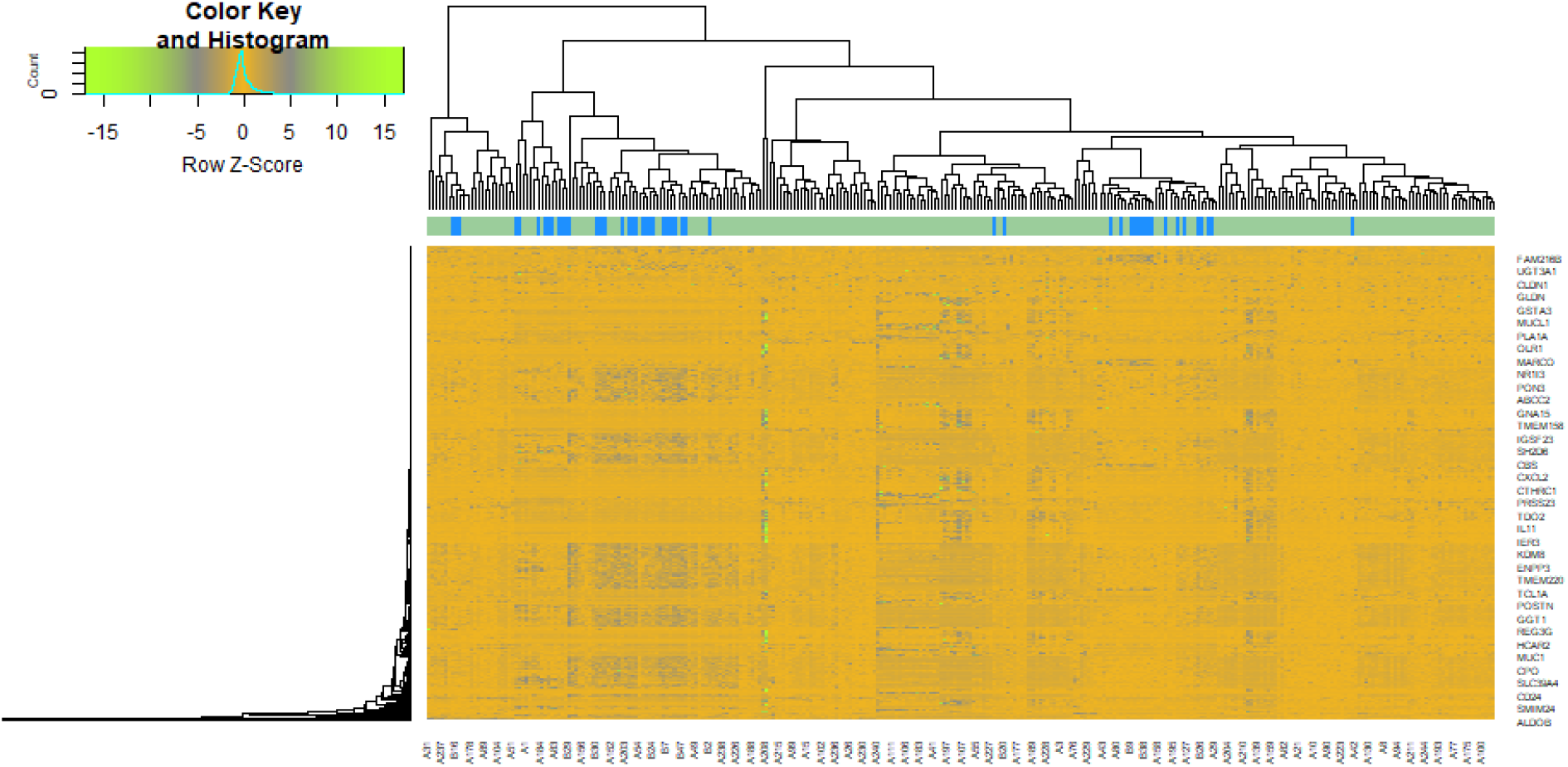
Heat map of differentially expressed genes. Legend on the top left indicate log fold change of genes. (A1 – A254 = CD samples; B1 – B50 = normal control samples)

### GO and pathway enrichment analyses of DEGs

A total of 957 DEGs were uploaded to g:Profiler for GO and REACTOME pathway enrichment analyses. The terms of each GO category are provided in Table 2. Most DEGs were enriched in the BP: response to stimulus, response to chemical, small molecule metabolic process and regulation of biological quality; CC: extracellular region, intrinsic component of membrane, membrane and cytoplasm; and MF: signaling receptor binding, molecular transducer activity, transporter activity and catalytic activity. The results of REACTOME pathway enrichment are shown in Table 3. The REACTOME pathway enrichment analysis confirmed that the DEGs were mainly associated with immune system, neutrophil degranulation, biological oxidations and metabolism.

**Table 2.**
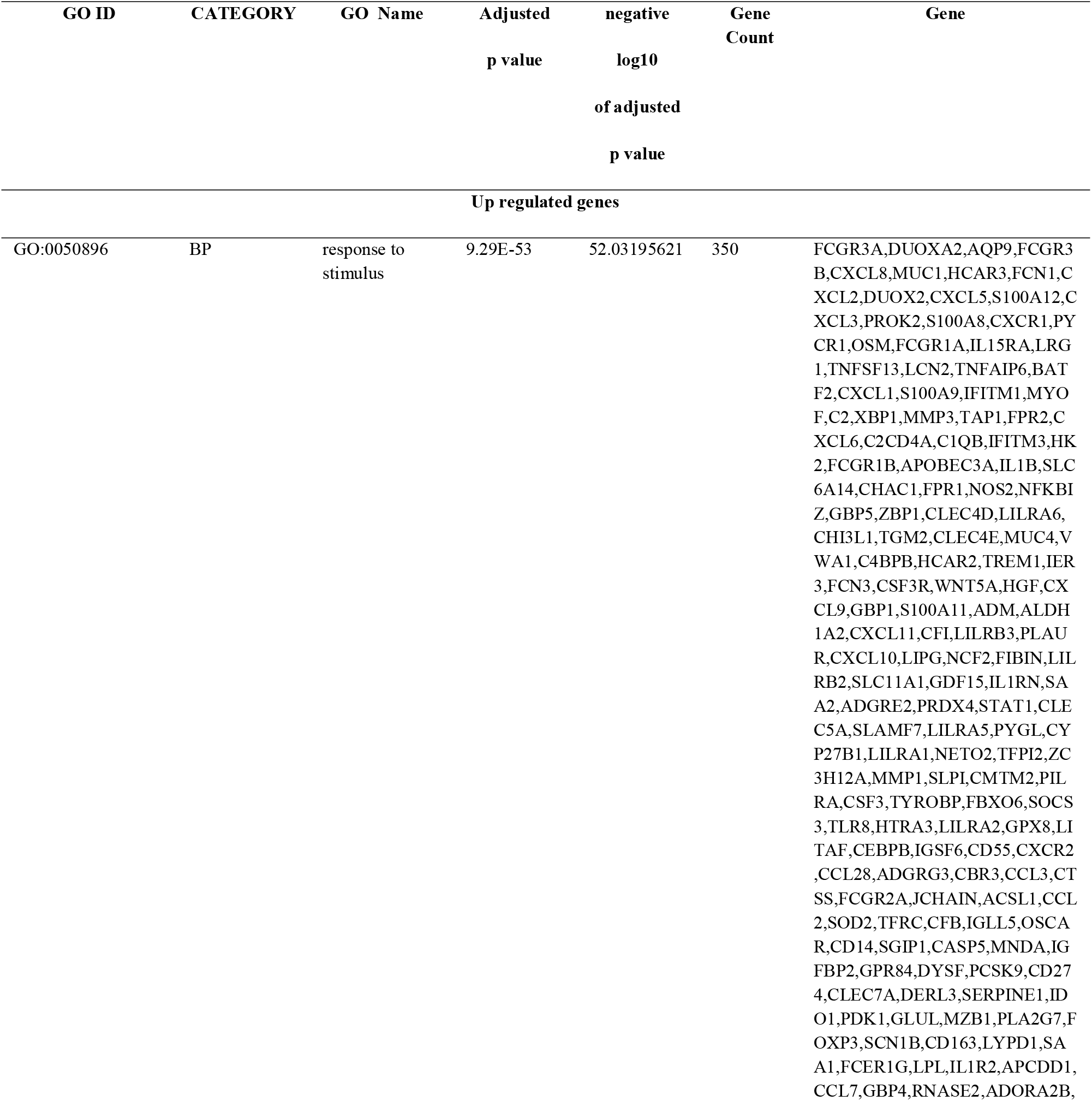
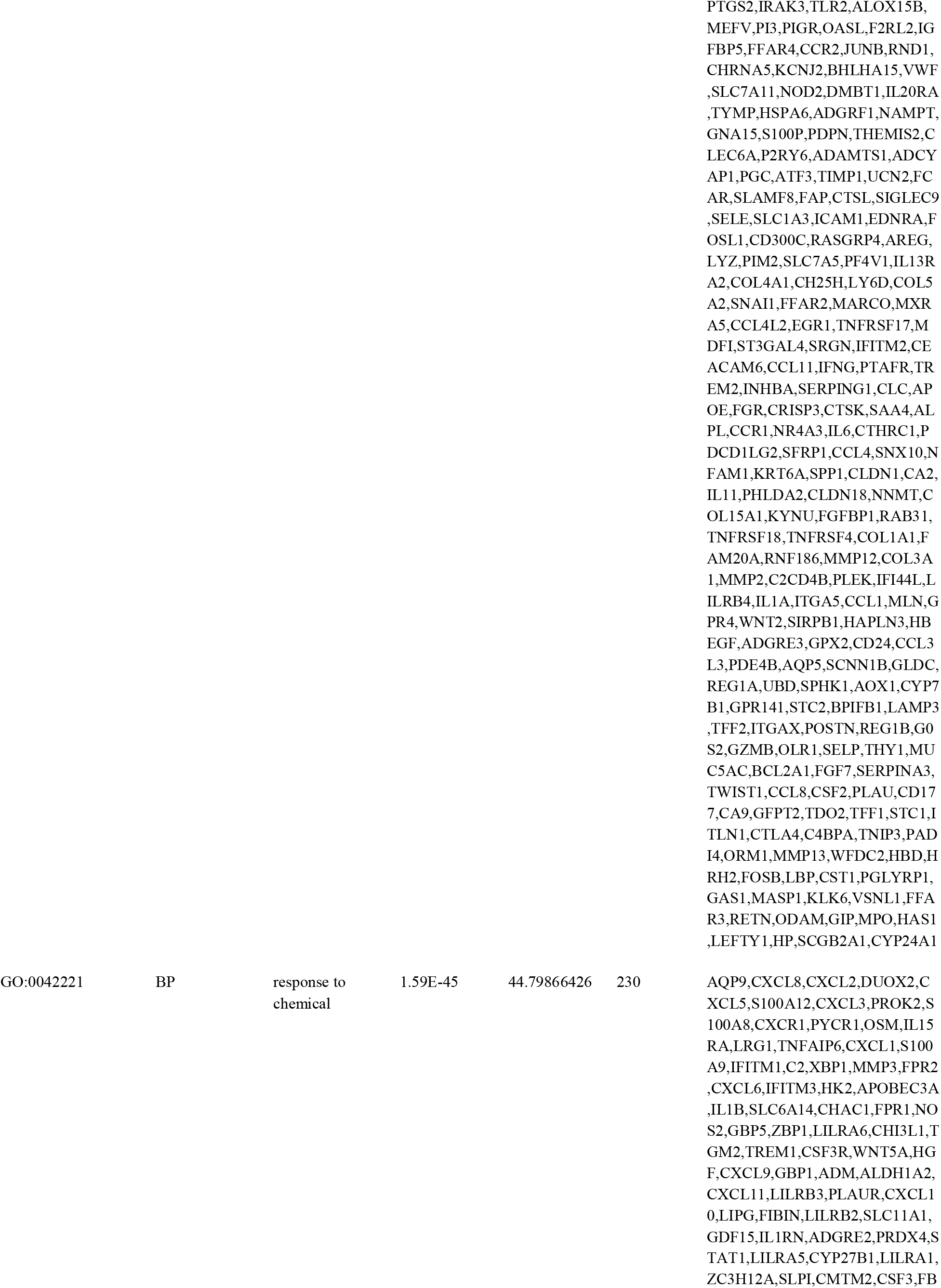
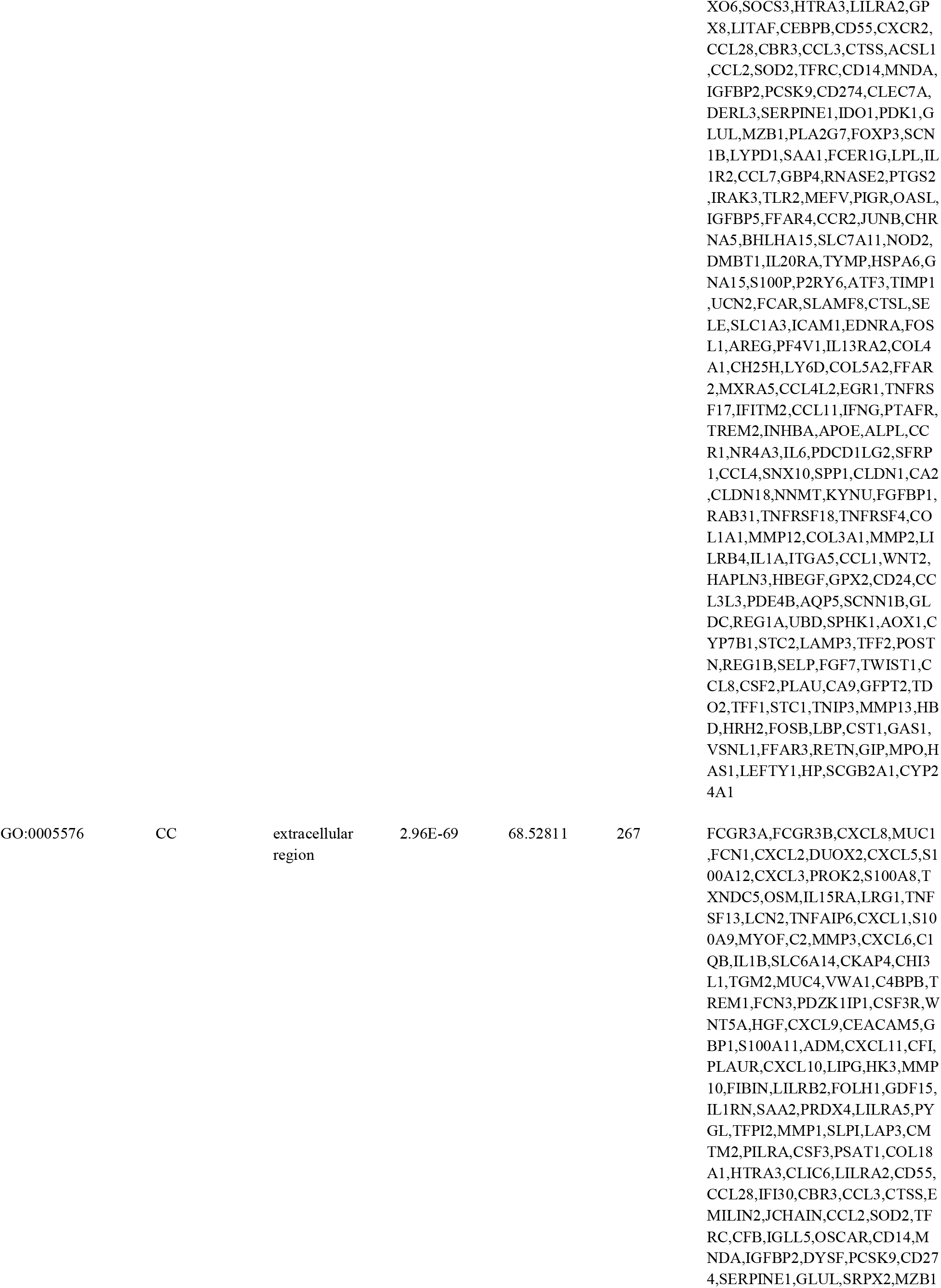
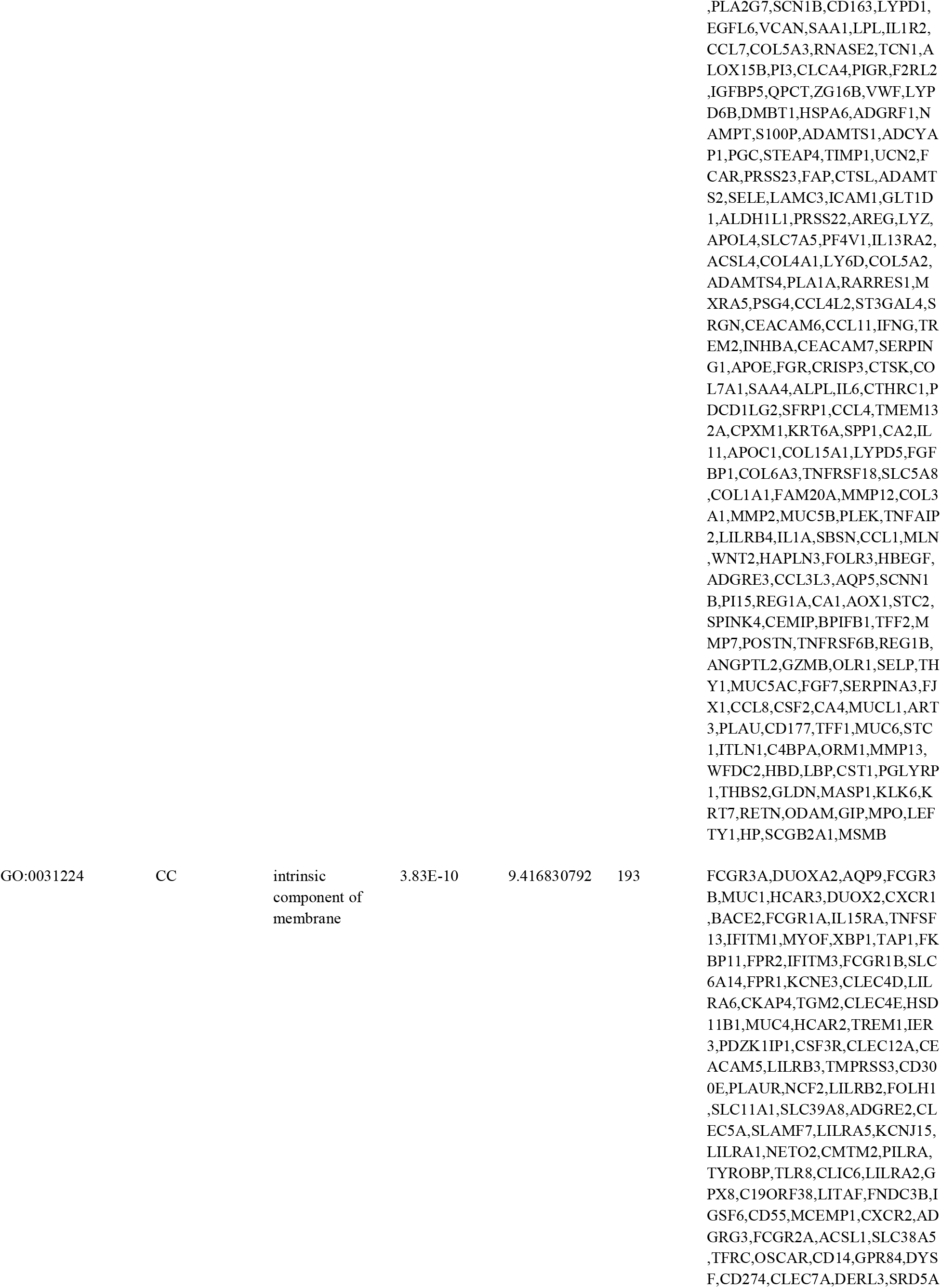
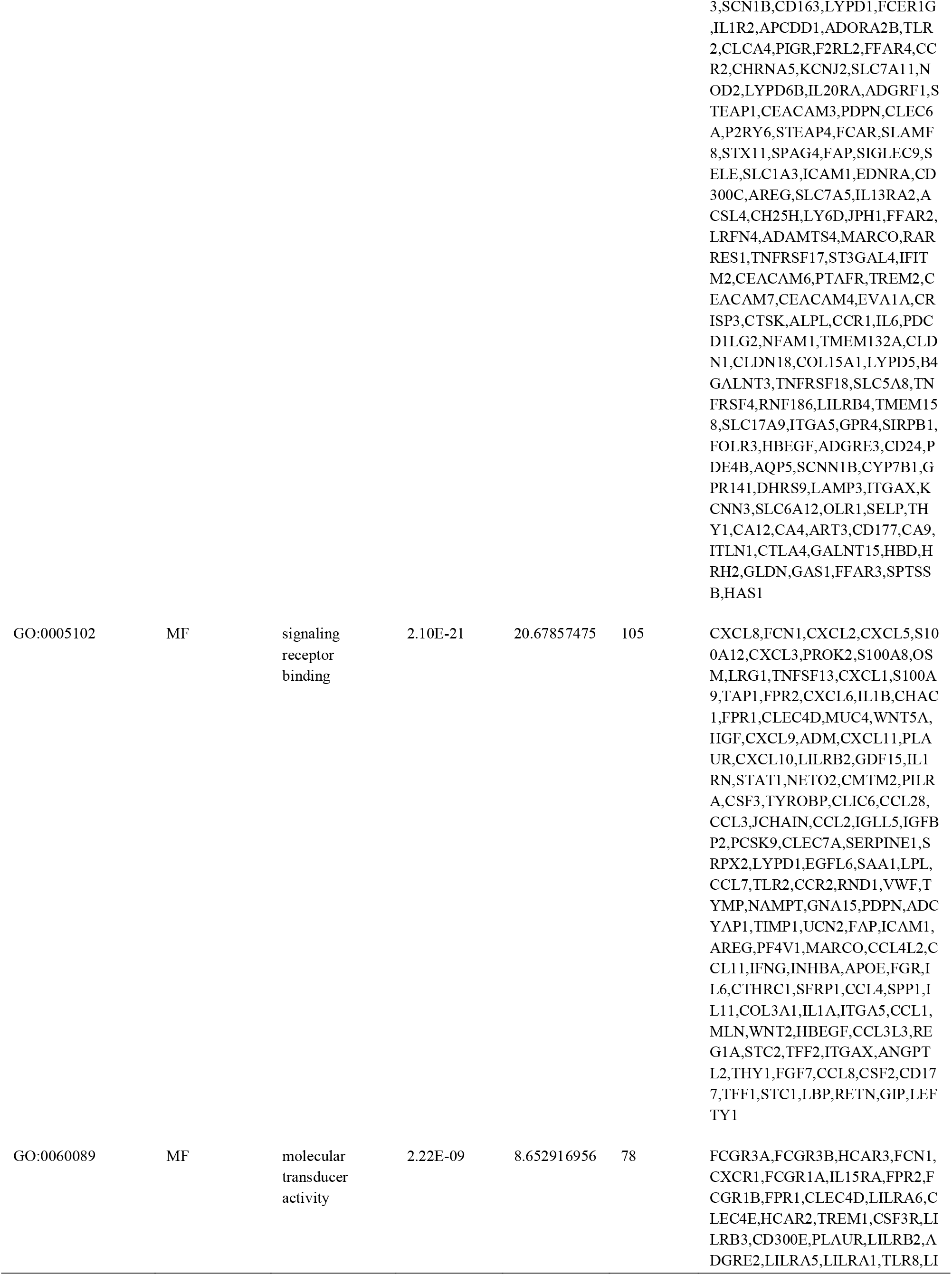
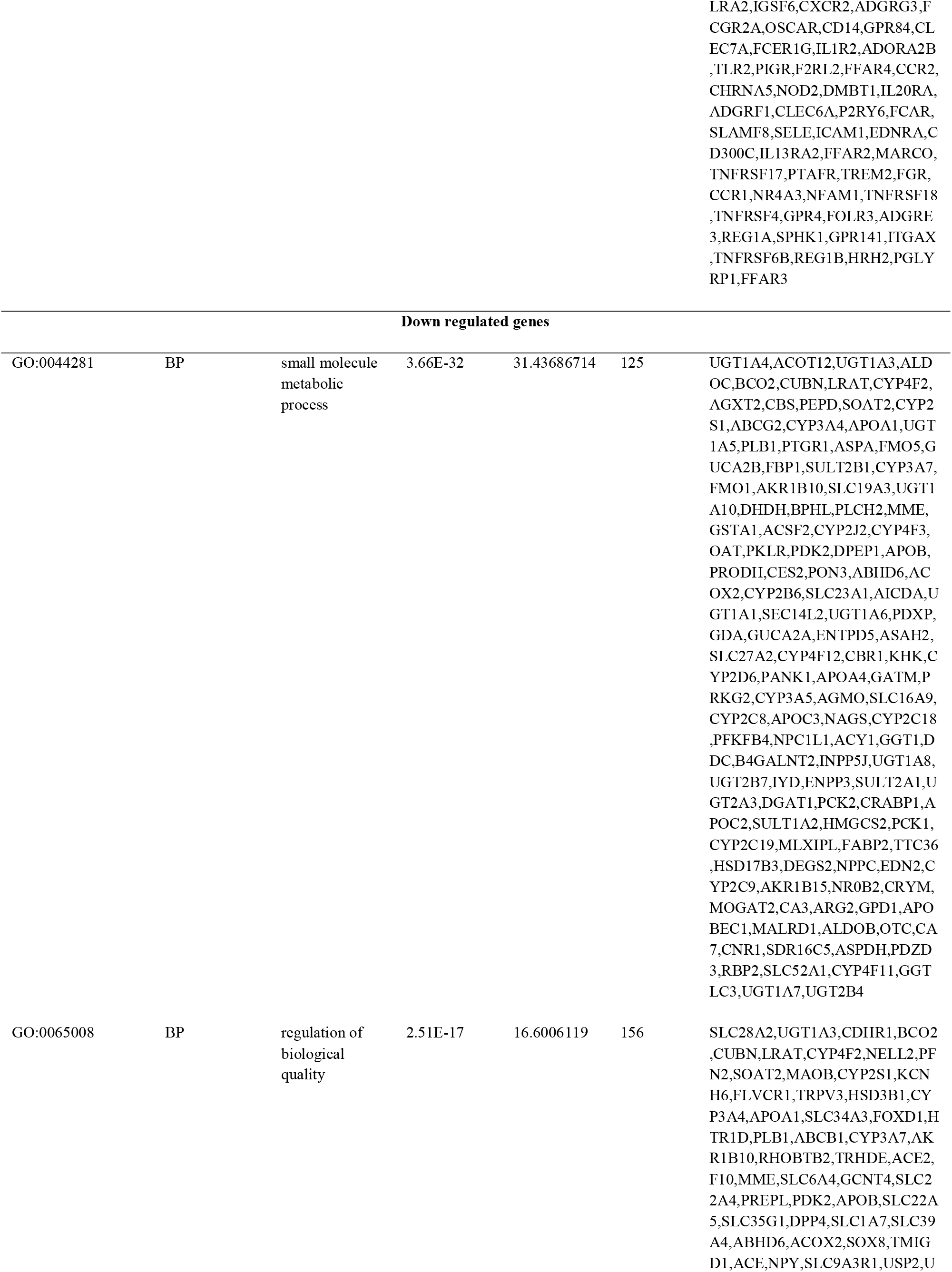
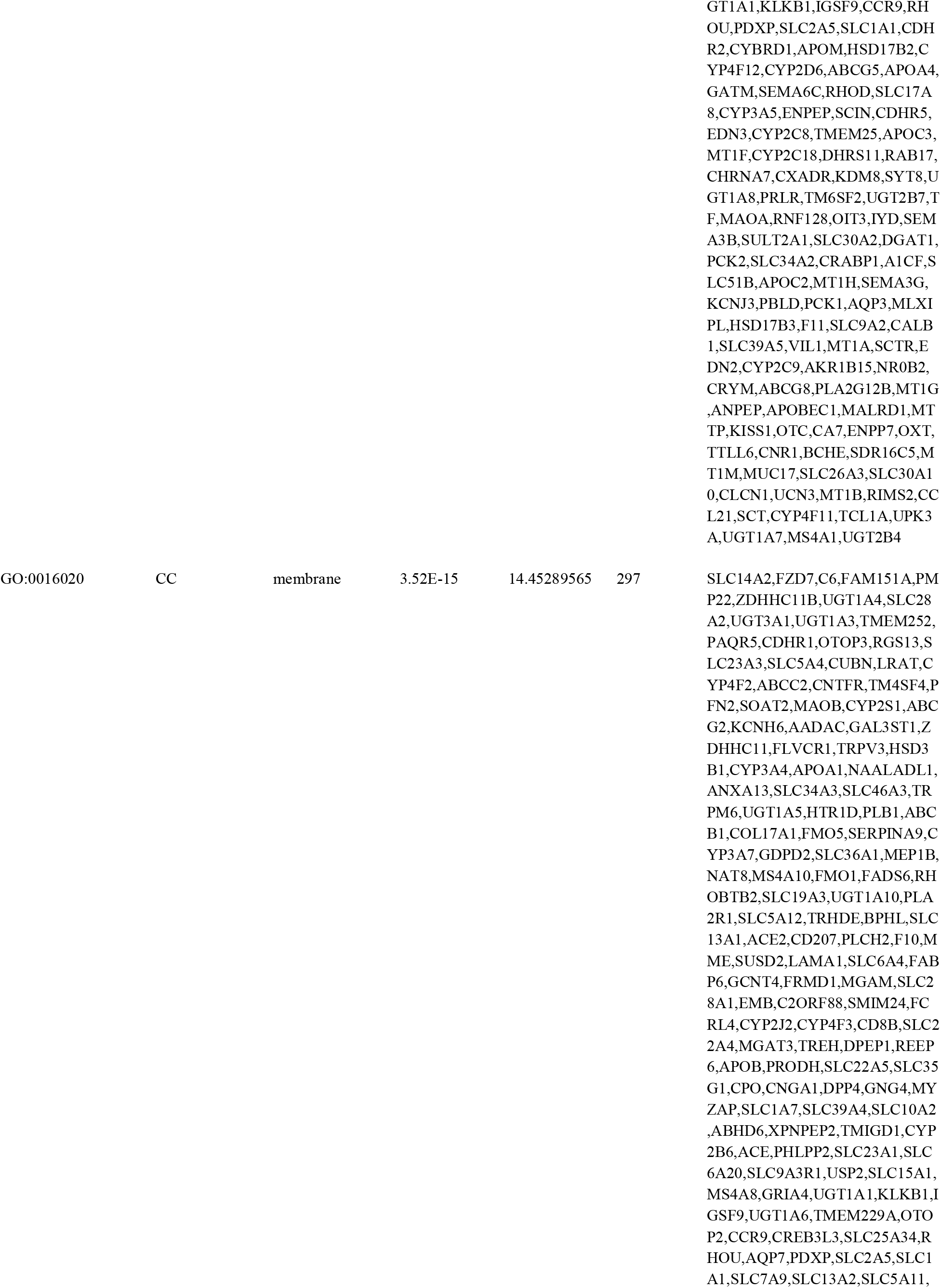
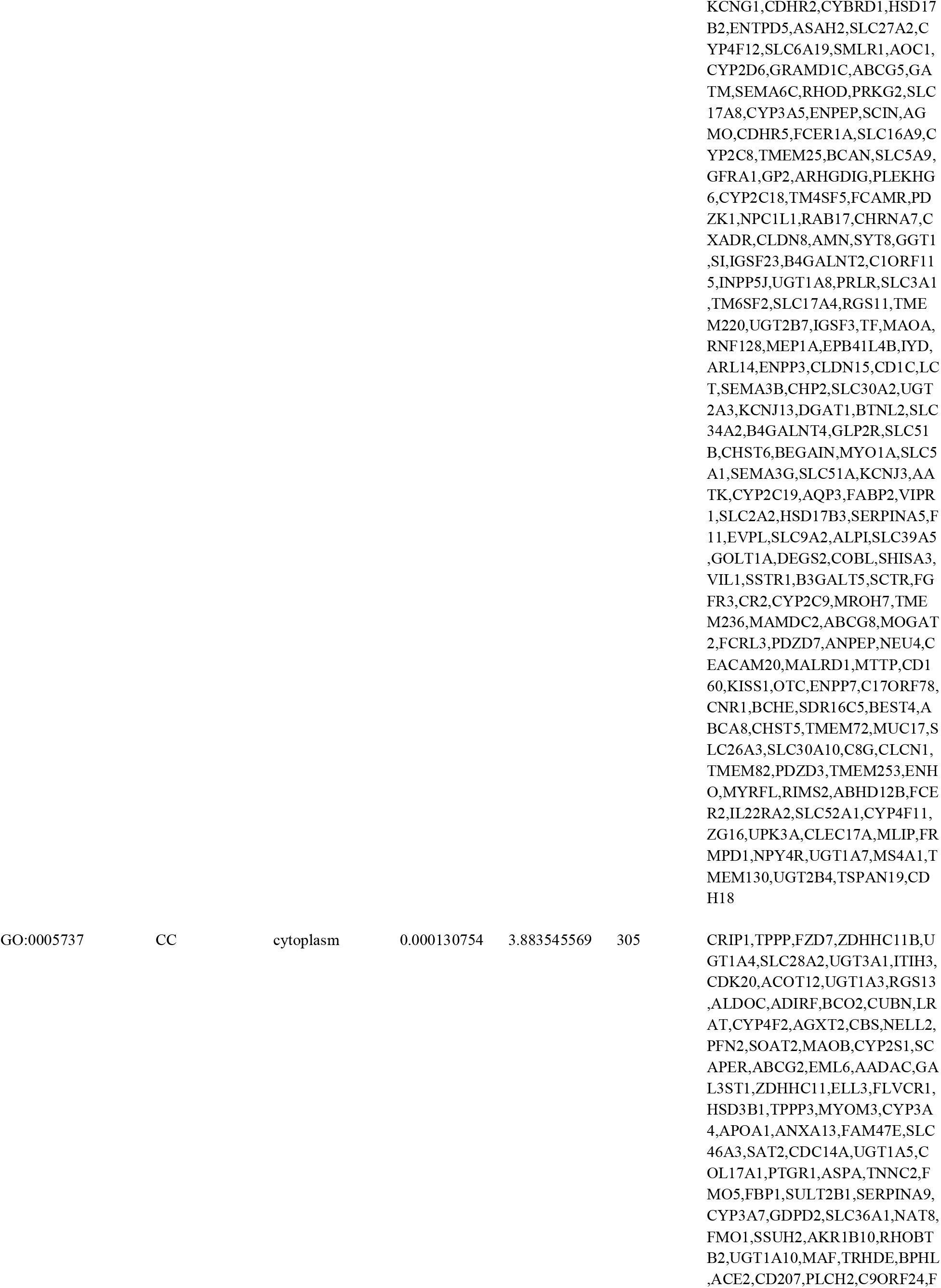
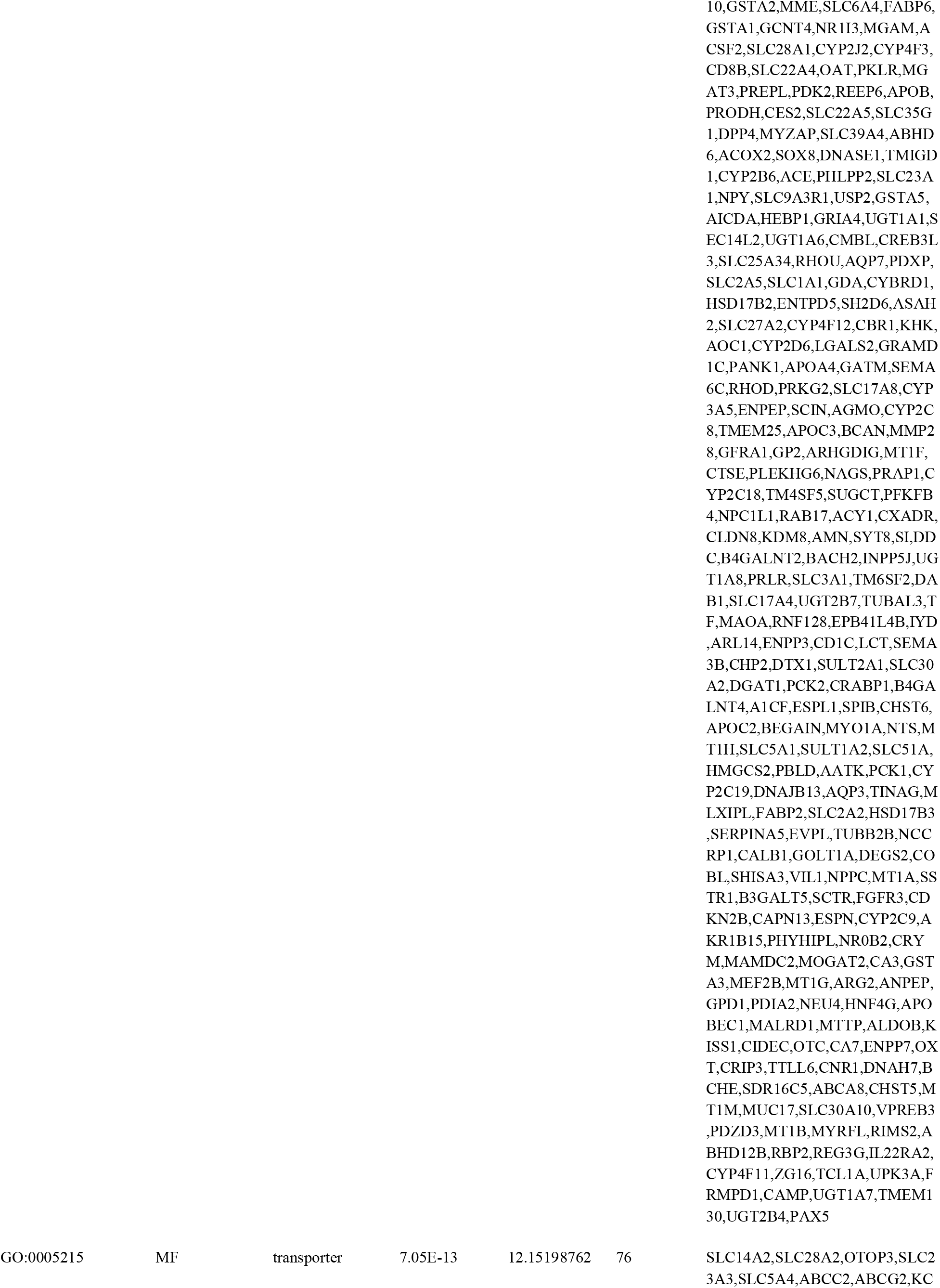
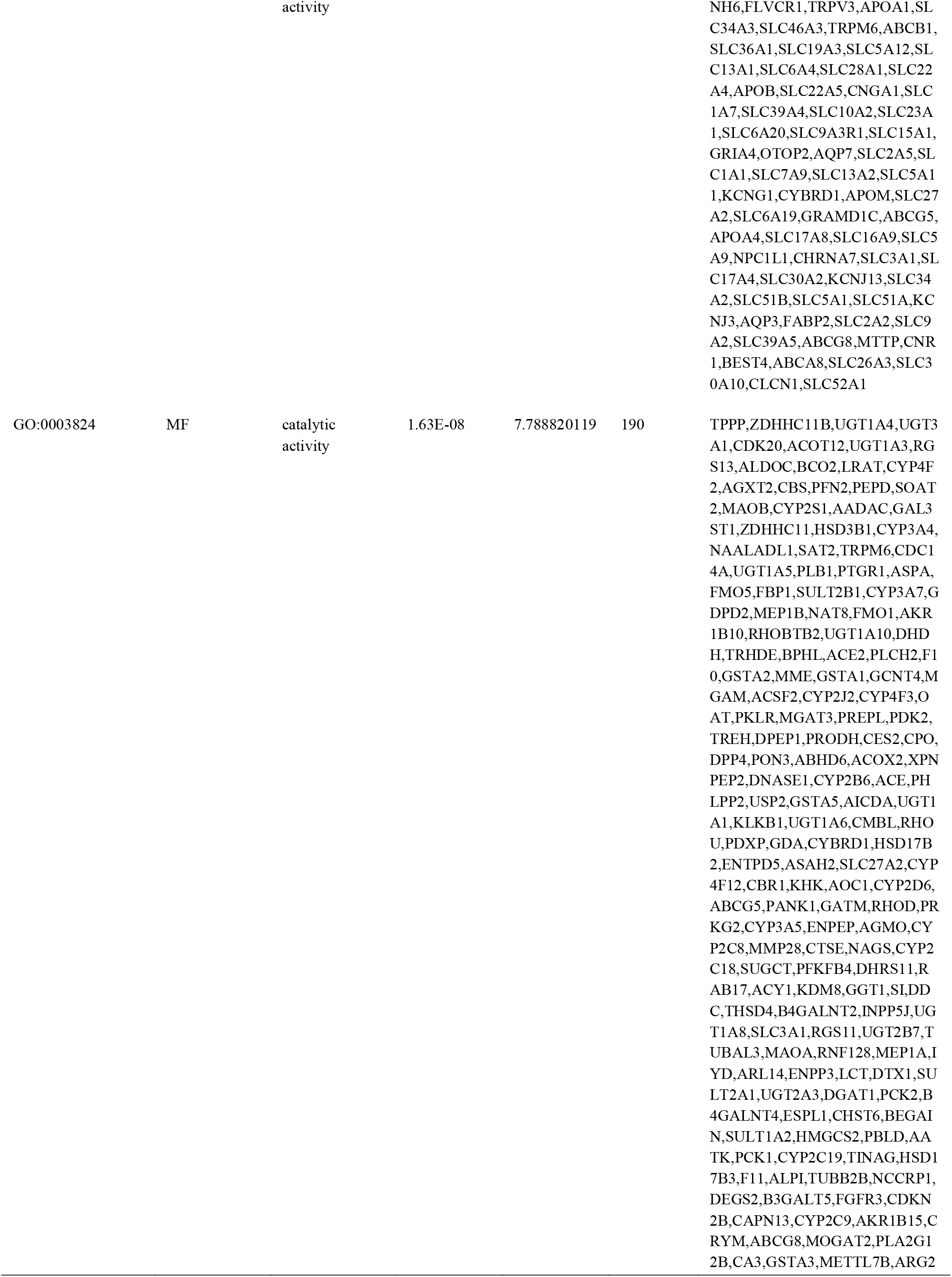
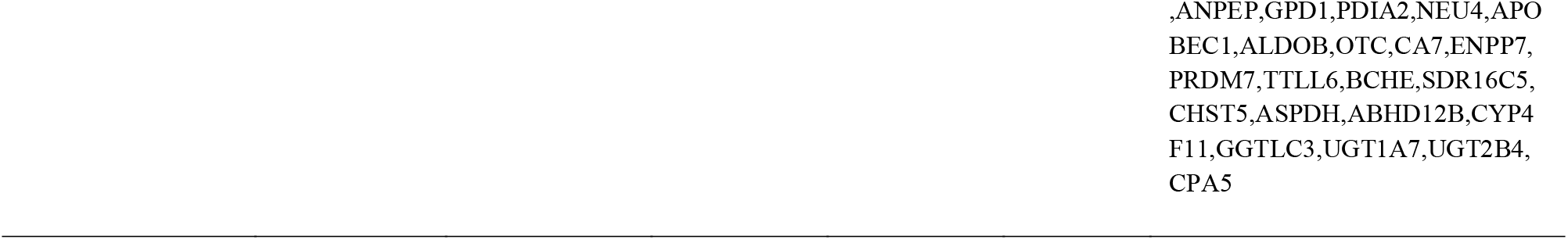
The enriched GO terms of the up and down regulated differentially expressed genes

**Table 3.**
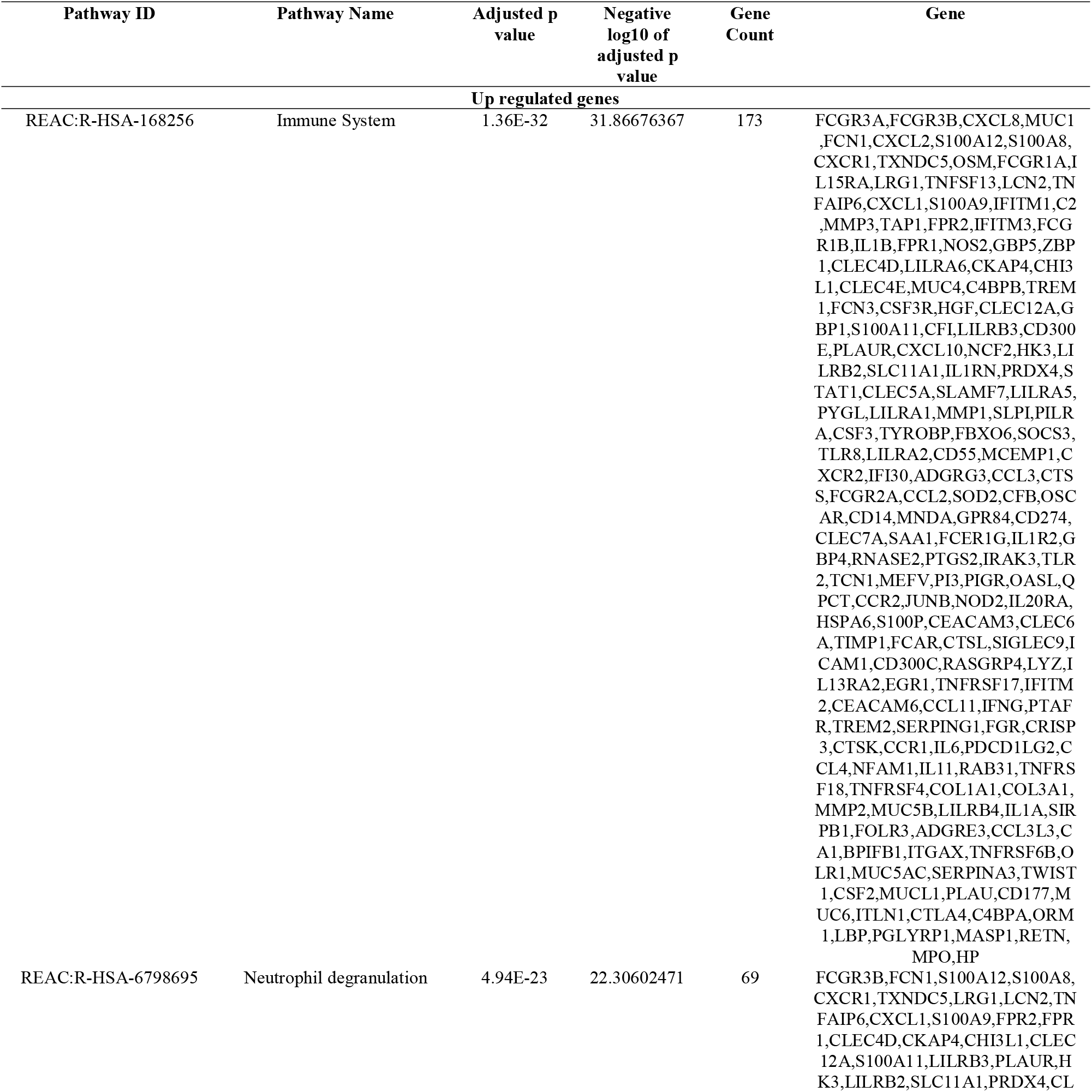
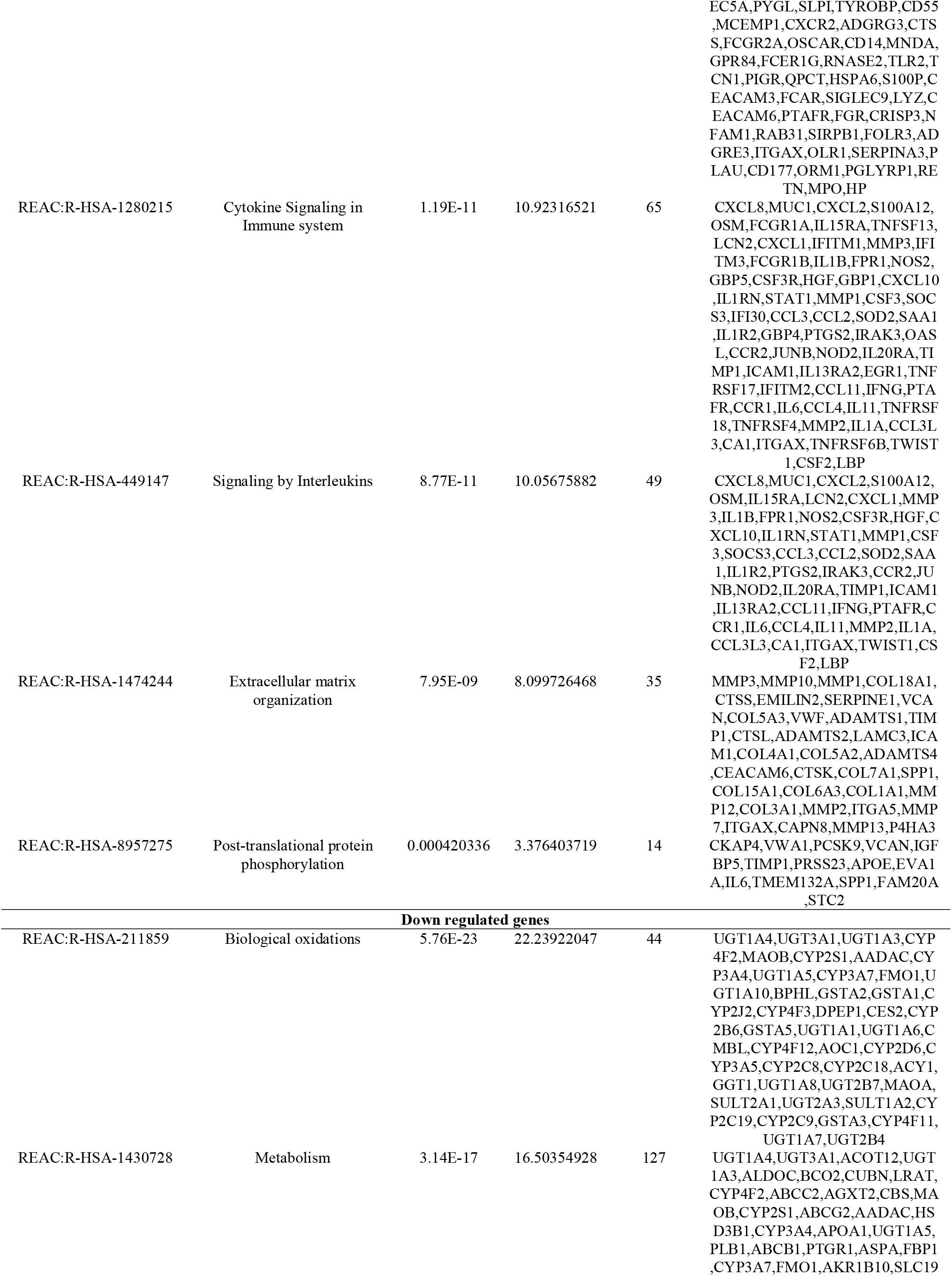
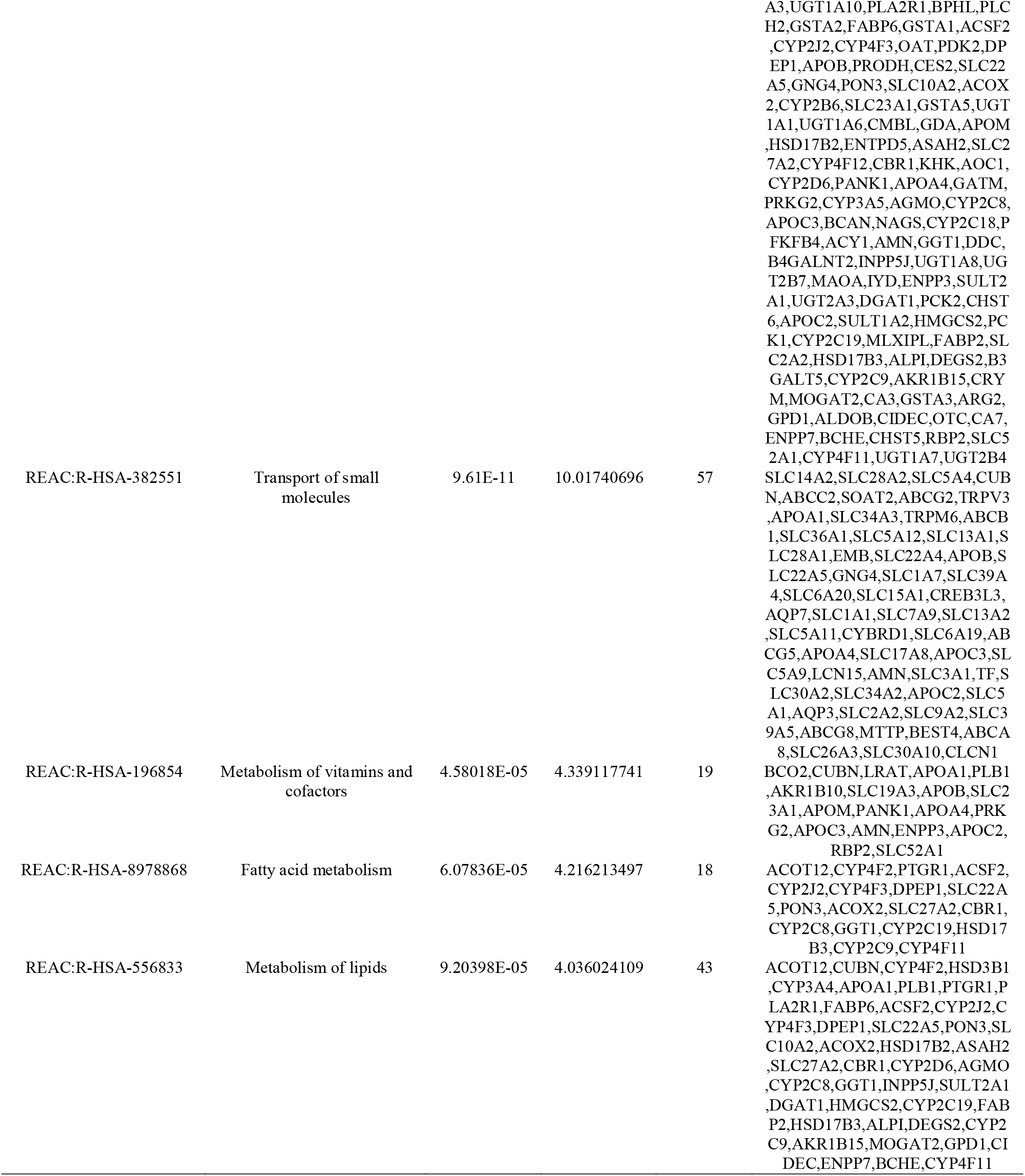
The enriched pathway terms of the up and down regulated differentially expressed genes

### Construction of the PPI network and module analysis

957 DEGs were imported into the HiPPIE database to explore the interrelationships between the various genes. 957 DEGs were used to establish the PPI network using the Cytoscape software. The PPI network consisted of 4030 nodes and 6196 edges (Fig. 3). The Network Analyzer in Cytoscape was used to screen the top genes with high node degree, betweenness, stress and closeness indicating the hub genes from the PPI network, including MDFI, MNDA, FBXO6, TFRC, STAT1, DPP4, MME, SLC39A4, APOA1 and TMEM25 and are listed in Table 4. Based on the degree of importance, two key modules were then screened from the PPI network using the PEWCC1 plug-in, and GO and pathway enrichment analysis was performed. Module 1 contained a total of 15 nodes and 49 edges (Fig. 4A), and module 2 contained a total of 15 nodes and 17 edges (Fig. 4B). The results showed that Module 1 was mainly related to immune system, response to stimulus, cytokine signaling in immune system, intrinsic component of membrane and response to chemical. Module 2 was primarily involved in the regulation of biological quality.

**Fig. 3.**
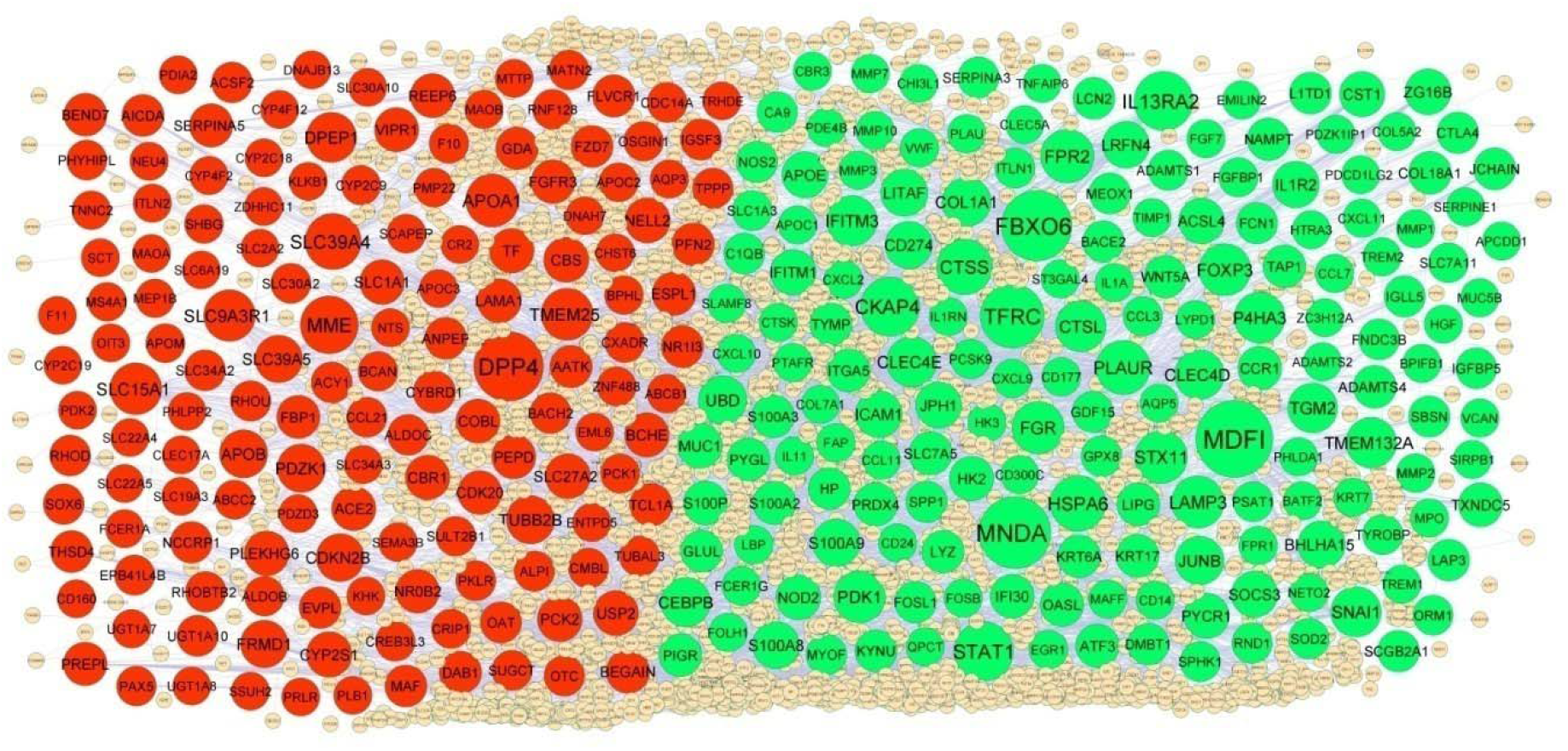
PPI network of DEGs. Up regulated genes are marked in green; down regulated genes are marked in red

**Table 4.**
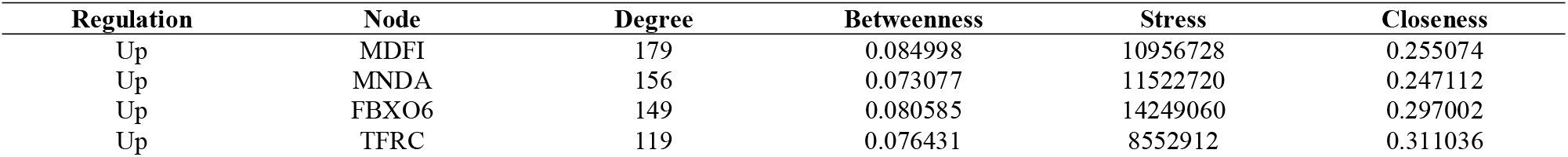
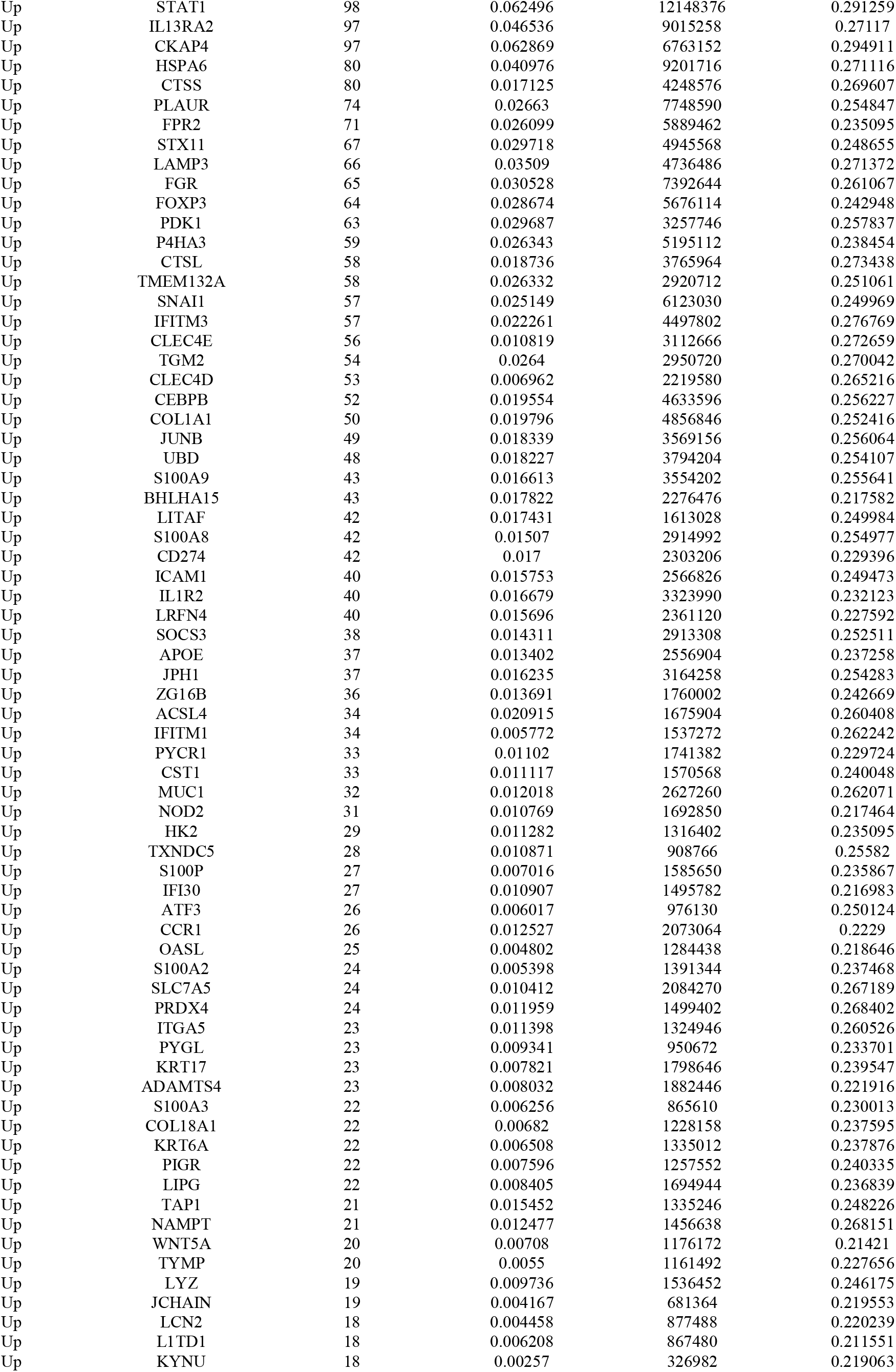
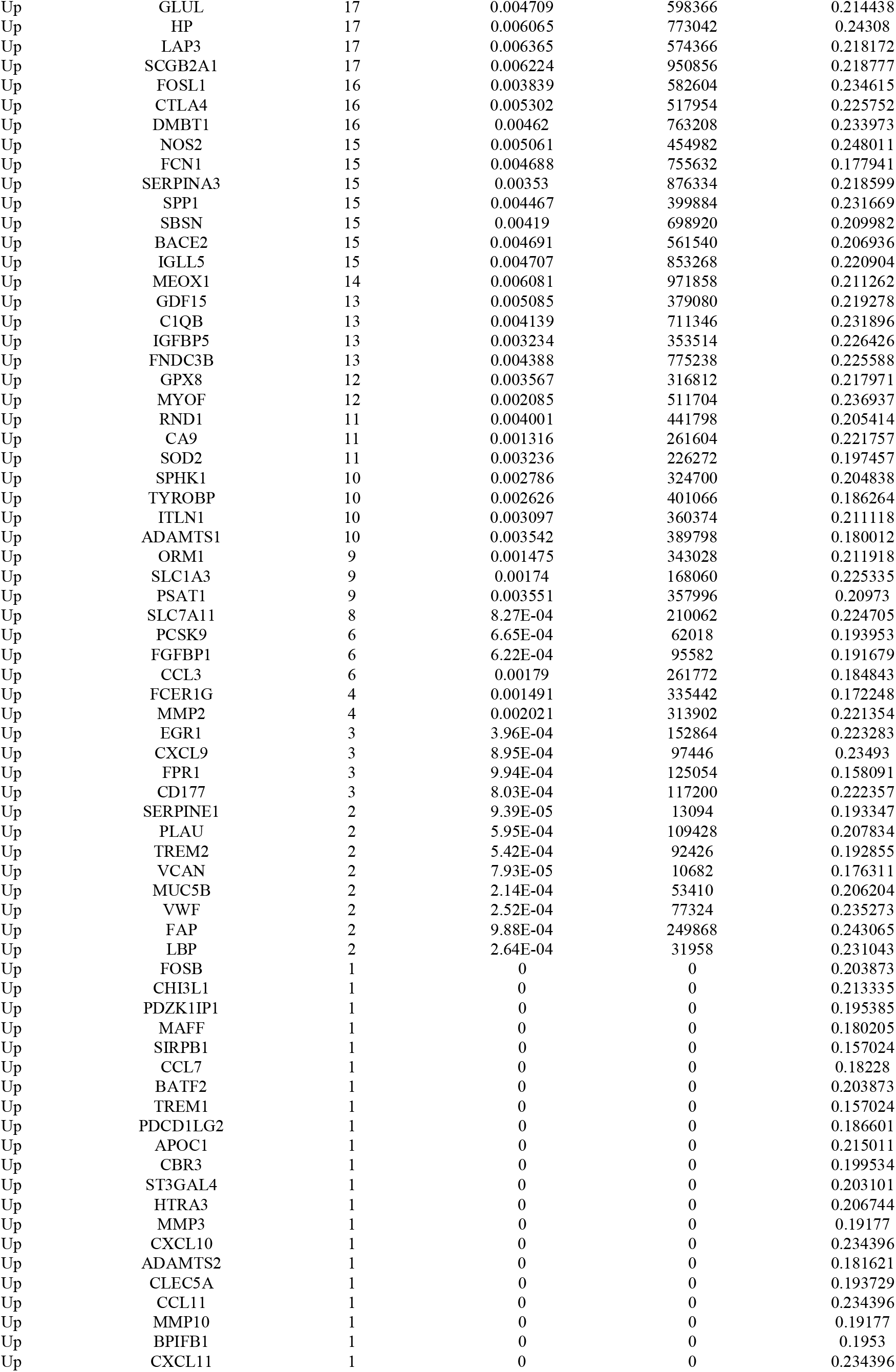
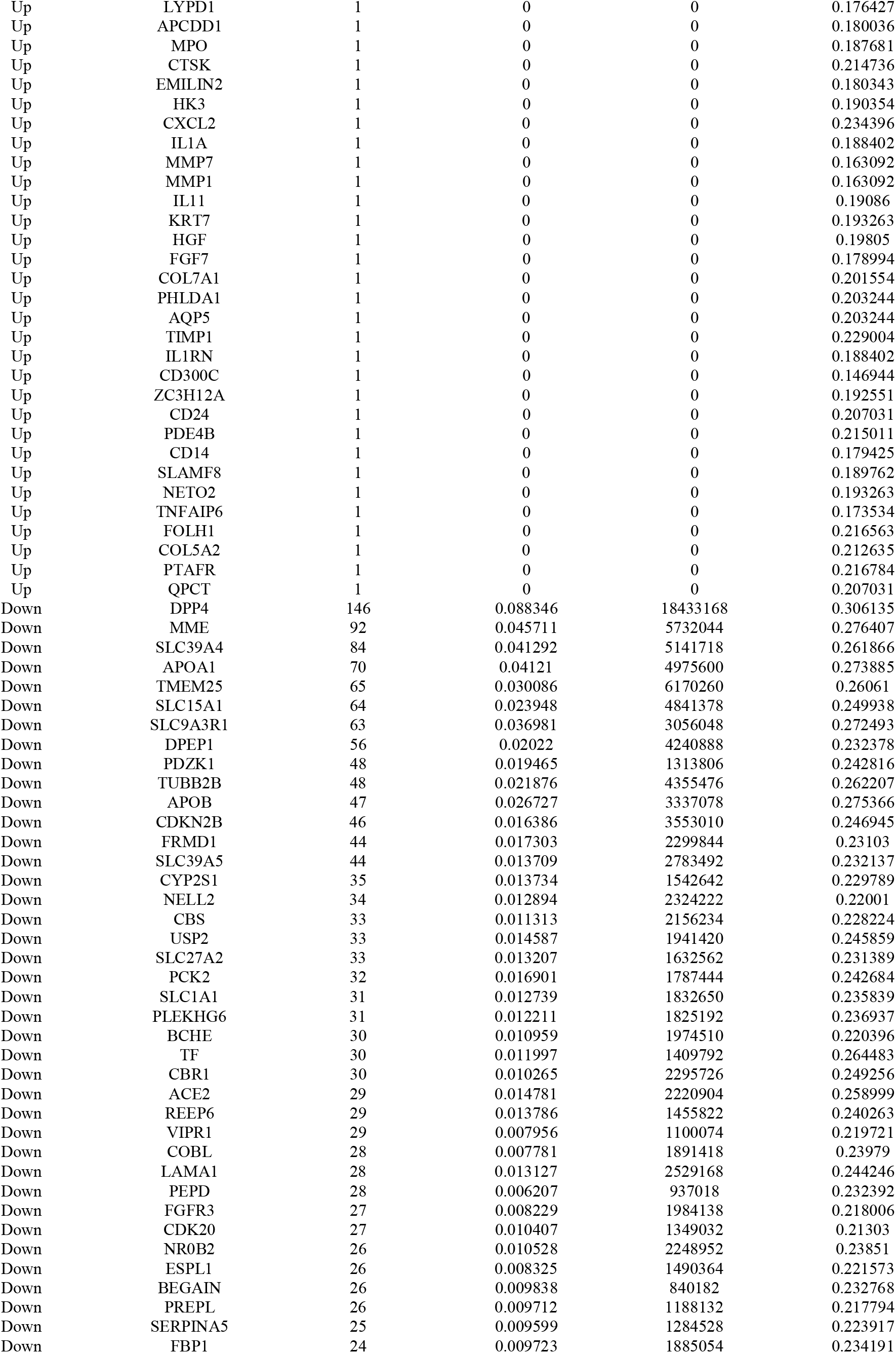
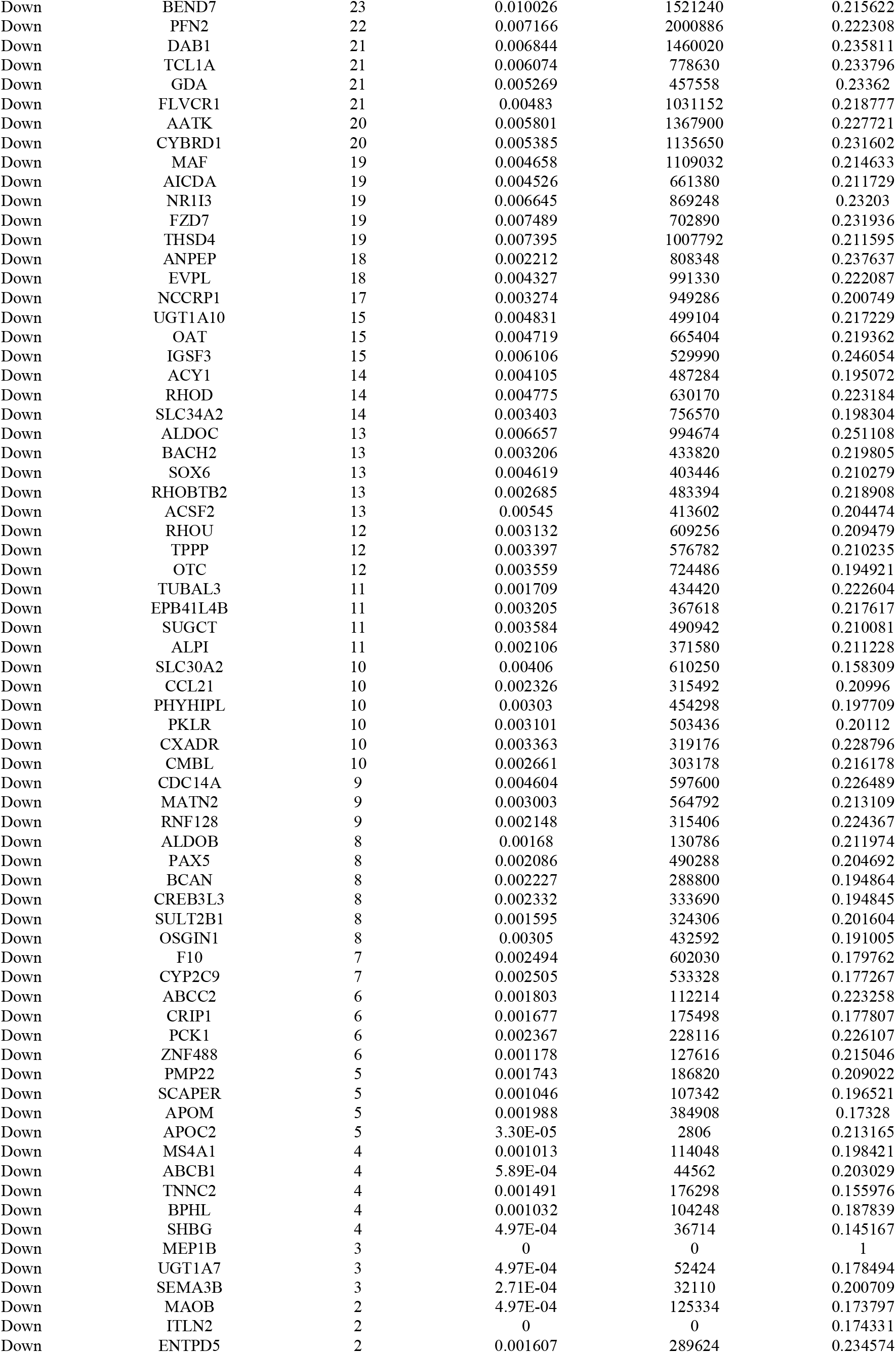
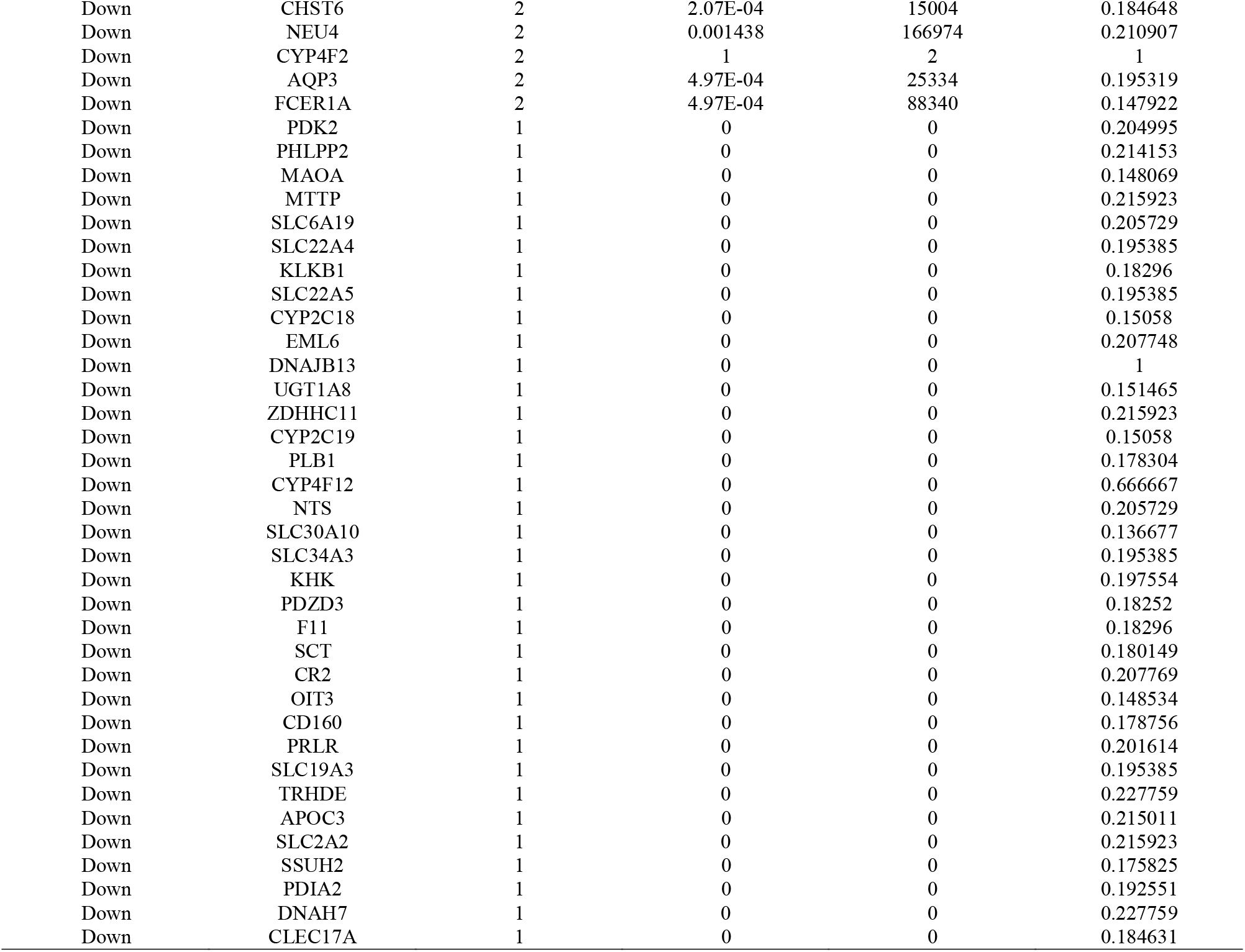
Topology table for up and down regulated genes

**Fig. 4.**
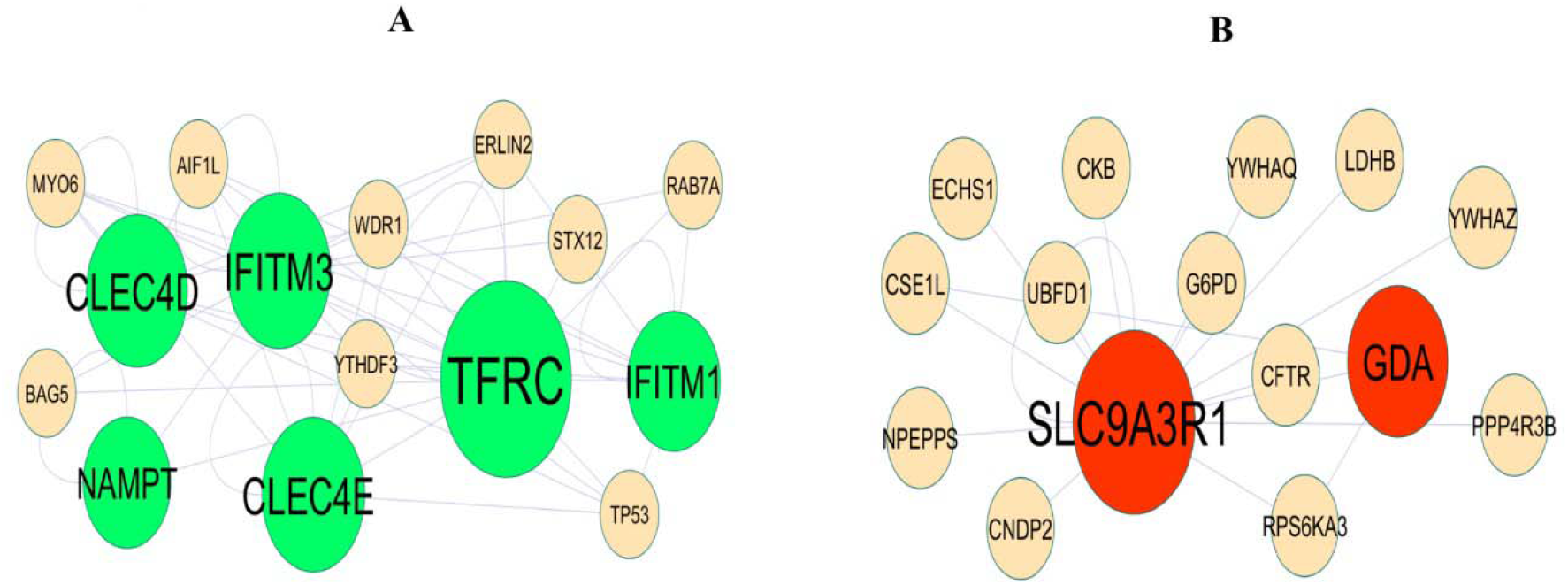
Modules selected from the DEG PPI between patients with FSGS and normal controls. (A) The most significant module was obtained from PPI network with 15 nodes and 49 edges for up regulated genes (B) The most significant module was obtained from PPI network with 15 nodes and 17 edges for down regulated genes. Up regulated genes are marked in green; down regulated genes are marked in red

### miRNA-hub gene regulatory network construction

To explore the key genes and miRNA involved in CD, miRNA-hub gene regulatory network of the hub genes was constructed. miRNA-hub gene regulatory network with 2377 nodes (miRNA: 2058; hub gene: 319) and 11302 edges (Fig. 5) and are listed in Table 5, which showed that 162 miRNAs (ex; hsa-mir-629-5p) could regulate TFRC expression; 102 miRNAs (ex; hsa-mir-25-5p) could regulate CKAP4 expression; 65 miRNAs (ex; hsa-mir-3714) could regulate HSPA6 expression; 63 miRNAs (ex; hsa-mir-146a) could regulate STAT1 expression; 43 miRNAs (ex; hsa-mir-4651) could regulate LAMP3 expression; 56 miRNAs (ex; hsa-mir-6130) could regulate SLC9A3R1 expression; 54 miRNAs (ex; hsa-mir-518f-5p) could regulate MME expression; 41 miRNAs (ex; hsa-mir-522-5p) could regulate TMEM25 expression; 35 miRNAs (ex; hsa-mir-641) could regulate CDKN2B expression; 27 miRNAs (ex; hsa-mir-148a-3p) could regulate TUBB2B expression.

**Fig. 5.**
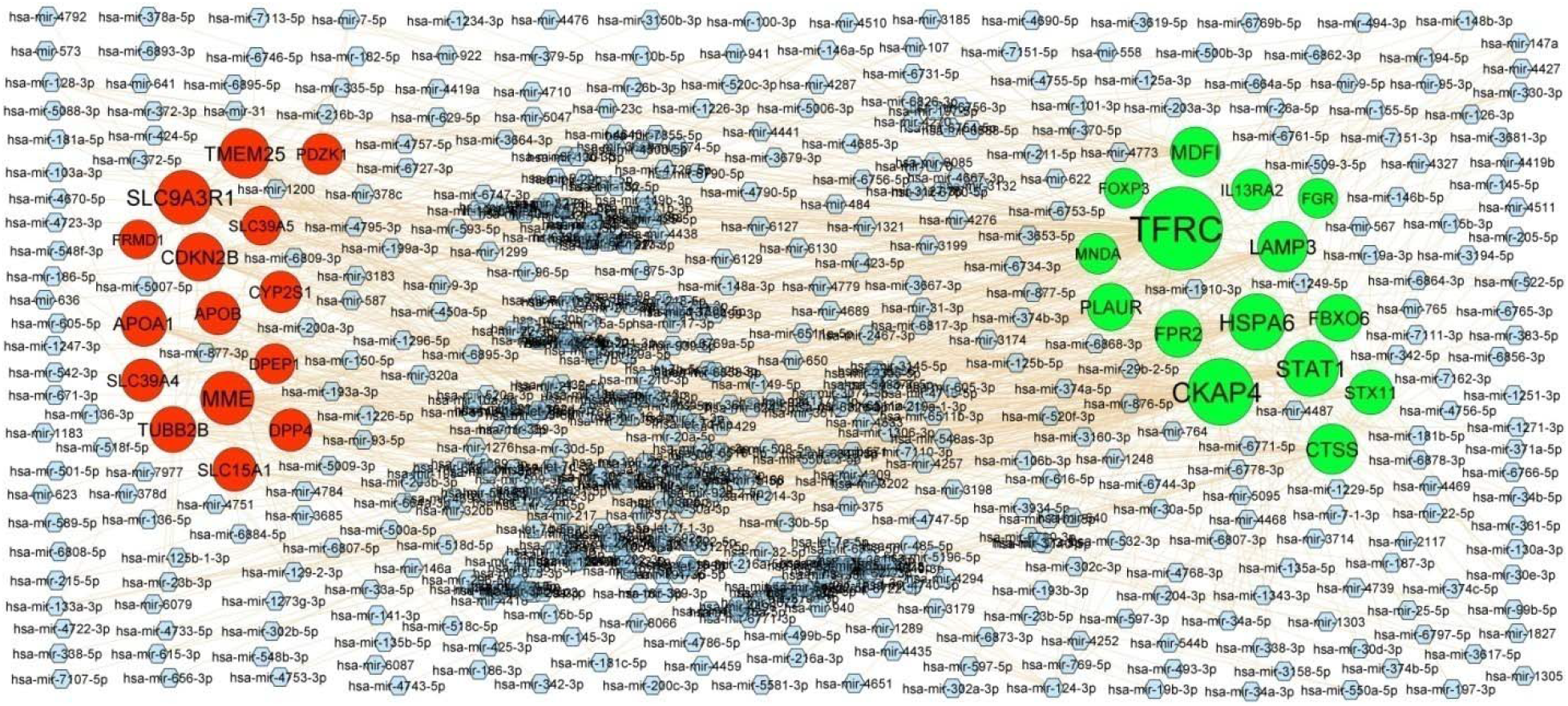
Target gene - miRNA regulatory network between target genes. The blue color diamond nodes represent the key miRNAs; up regulated genes are marked in green; down regulated genes are marked in red.

**Table 5.**
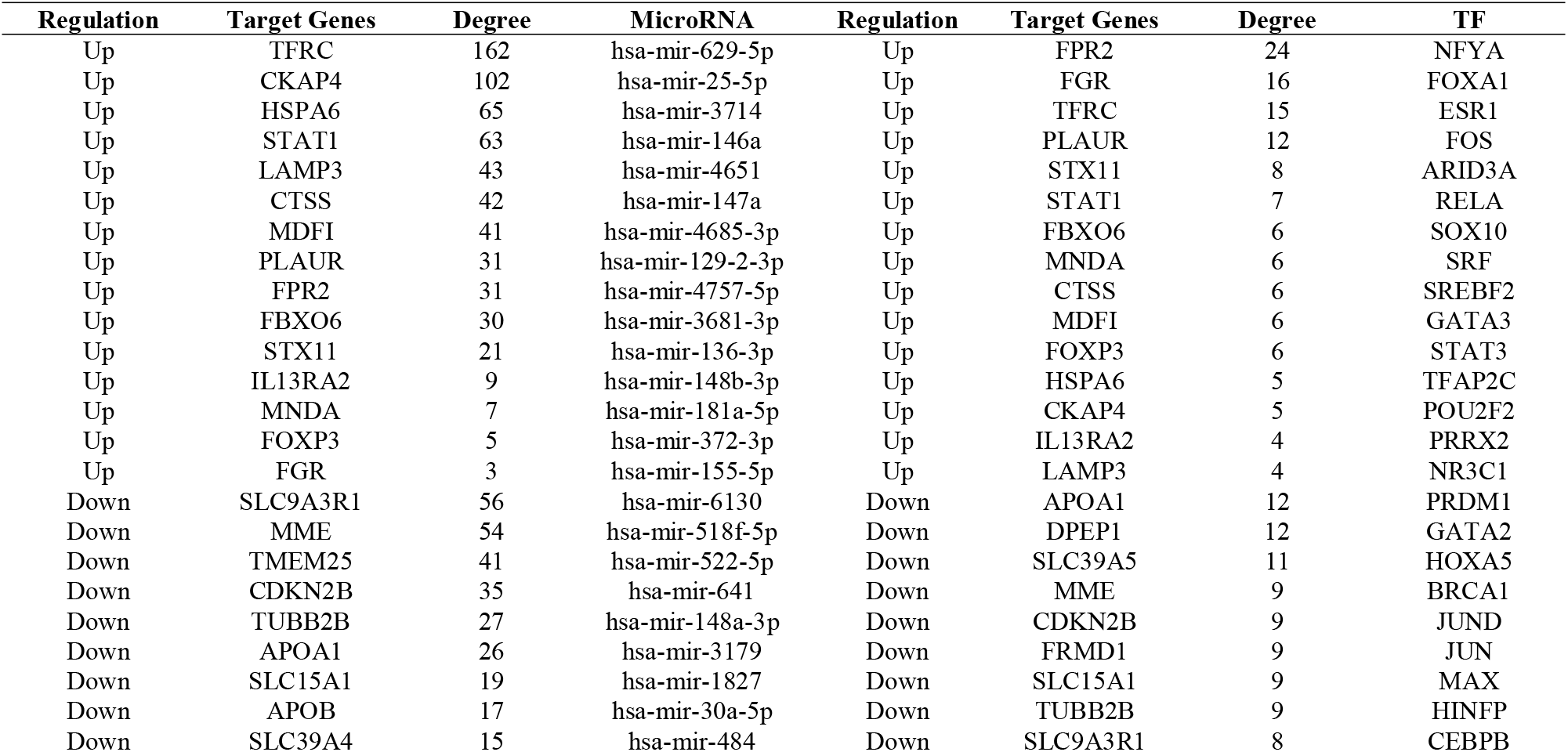
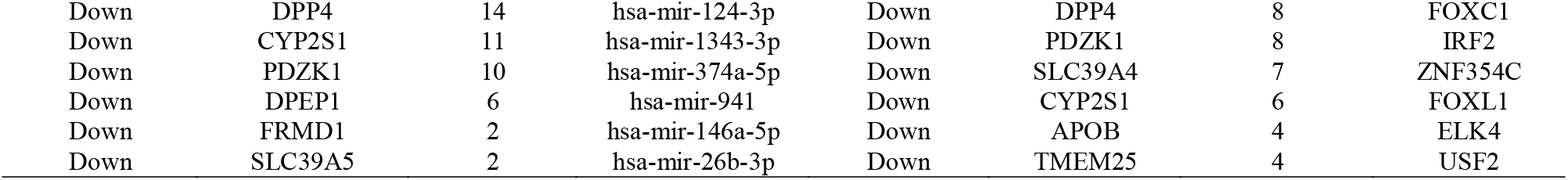
miRNA - target gene and TF - target gene interaction

### TF-hub gene regulatory network construction

To explore the key genes and TF involved in CD, TF-hub gene regulatory network of the hub genes was constructed. TF-hub gene regulatory network with 405 nodes (TF: 82; hub gene: 323) and 2514 edges (Fig. 6) and are listed in Table 5, which showed that 24 TFs (ex; NFYA) could regulate FPR2 expression; 16 TFs (ex; FOXA1) could regulate FGR expression; 15 TFs (ex; ESR1) could regulate TFRC expression; 12 TFs (ex; FOS) could regulate PLAUR expression; 8 TFs (ex; ARID3A) could regulate STX11 expression; 12 TFs (ex; PRDM1) could regulate APOA1 expression; 12 TFs (ex; GATA2) could regulate DPEP1 expression; 11 TFs (ex; HOXA5) could regulate SLC39A5 expression; 9 TFs (ex; BRCA1) could regulate MME expression; 9 TFs (ex; JUND) could regulate CDKN2B expression.

**Fig. 6.**
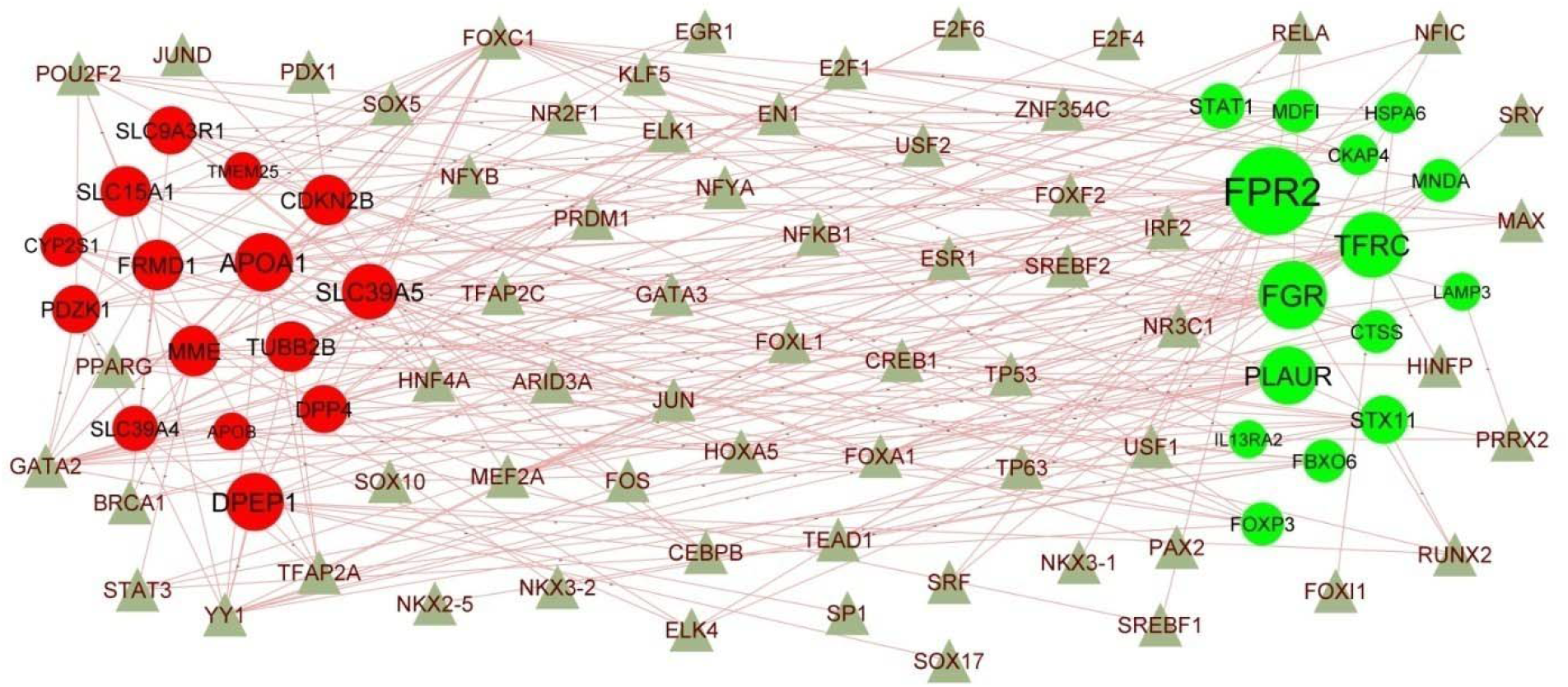
Target gene - TF regulatory network between target genes. The olive color triangle nodes represent the key TFs; up regulated genes are marked in green; down regulated genes are marked in red.

### Receiver operating characteristic curve (ROC) analysis

The ROC curves (Fig. 7) of these three genes showed that their AUC are as follows: MDFI (AUC = 0.933), MNDA (AUC = 0.915), FBXO6 (AUC = 0.922), TFRC (AUC = 0.922), STAT1 (AUC = 0.930), DPP4 (AUC = 0.943), MME (AUC = 0.900), SLC39A4 (AUC = 0.902), APOA1 (AUC = 0.939) and TMEM25 (AUC = 0.920). Since ROC curves had good specificity and sensitivity, MDFI, MNDA, FBXO6, TFRC, STAT1, DPP4, MME, SLC39A4, APOA1 and TMEM25 had excellent diagnostic efficiency for distinguishing CD and healthy control.

**Fig. 7.**
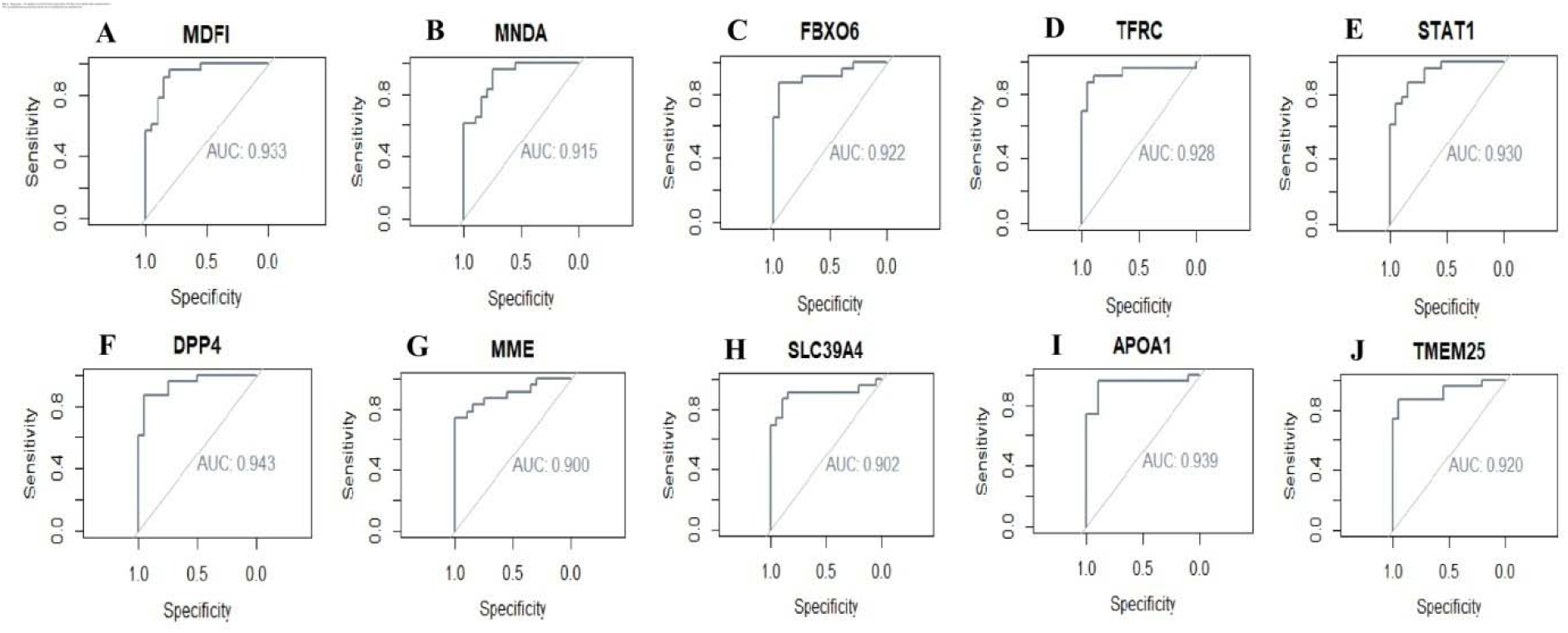
ROC curve analyses of hub genes. A) MDFI B) MNDA C) FBXO6 D) TFRC E) STAT1 F) DPP4 G) MME H) SLC39A4 I) APOA1 J) TMEM25

## Discussion

A NGS study is an ideal way to comprehensively investigate CD. In this investigation, NGS data was analyzed for identification potential biomarkers and explore molecular mechanisms of CD. Although CD is an inflammatory disease, it has recently been established that both inflammatory responses occur early during CD [40]. In the present investigation, NGS dataset (GSE101794) was downloaded from the GEO database. A total of 957 DEGs were screened: 478 up regulated genes and 479 down regulated genes. Lee et al [41], Gijsbers et al [42], Hashash et al [43], Wnorowski et al [44] and Kyodo et al [45] found that FCGR3A, CXCL8, MUC1, HCAR3 and DUOX2 might play an important role in the pathophysiology of CD. Nourse et al [46] and Nakashima et al [47] found that FCGR3A and MUC1 was altered expressed in coagulation and fibrinolysis. These coagulation and fibrinolysis responsible genes might positively linked with CD. Ying et al. [48], Huang et al. [49], Liu et al. [50], Yang et al [51], Bao et al [52], Zhang et al [53], Zhang et al [54], Ye et al [55] and Xi et al [56] showed that FCGR3A, AQP9, MUC1, HCAR3, CXCL2, DUOX2, CRIP1, TPPP (tubulin polymerization promoting protein) and FZD7 play an important role in the occurrence and development of colorectal cancer. These colorectal cancer responsible genes might be associated with CD. The altered expression of FCGR3A [57], FCGR3B [58], MUC1 [59] and CXCL2 [60] might be associated with autoimmune disease progression. These autoimmune disease responsible genes might found to be substantially related to CD. MacFie et al. [61], Walana et al. [62], Asano et al. [63] and Kvorjak et al. [64] studied the clinical and prognostic value of DUOXA2, CXCL8, FCGR3A, FCGR3B, MUC1 and DUOX2 in patients with ulcerative colitis. The altered expression of CXCL8 [65] and FCN1 [66] contributes to the type 1 diabetes mellitus progression. These type 1 diabetes mellitus responsible genes might play an important role in CD. Kerr et al. [67] and Ampuero et al [68] demonstrated that altered expression of C6 and SLC28A2 were found to be substantially related to anemia. These anemia responsible genes might be key genes in CD.

GO and REACTOME pathway enrichment analysis found that genes in patients with CD were enriched. Many pathways are associated with the pathogenesis of CD. Signaling pathway include immune system [69], neutrophil degranulation [70], cytokine signaling in immune system [71], extracellular matrix organization [72], post-translational protein phosphorylation [73], biological oxidations [74], metabolism [75] and metabolism of lipids [76] were responsible for progression of CD. CXCL5 [77], CXCL3 [78], PROK2 [79], CXCR1 [80], PYCR1 [81], OSM (oncostatin M) [82], IL15RA [83], LRG1 [84], LCN2 [85], BATF2 [86], CXCL1 [87], S100A9 [88], IFITM1 [89], MYOF (myoferlin) [90], XBP1 [91], MMP3 [92], TAP1 [93], FPR2 [94], CXCL6 [95], C2CD4A [96], IFITM3 [97], IL1B [98], SLC6A14 [99], FPR1 [100], NOS2 [101], CHI3L1 [102], TGM2 [103], MUC4 [104], TREM1 [105], WNT5A [106], HGF (hepatocyte growth factor) [107], CXCL9 [108], GBP1 [109], S100A11 [110], ADM (adrenomedullin) [111], CXCL11 [112], CXCL10 [113], LILRB2 [114], GDF15 [115], IL1RN [116] STAT1 [117], SLAMF7 [105], CYP27B1 [118], NETO2 [119], TFPI2 [120], ZC3H12A [121], MMP1 [122], CSF3 [123], SOCS3 [124], TLR8 [125], HTRA3 [126], CEBPB (CCAAT enhancer binding protein beta) [127], CD55 [128], CXCR2 [129], CCL28 [130], CBR3 [131], CCL3 [132], FCGR2A [48], ACSL1 [133], CCL2 [134], SOD2 [135], CD14 [136], IGFBP2 [137], CD274 [138], DERL3 [139], SERPINE1 [140], IDO1 [141], PDK1 [142], FOXP3 [143], CD163 [144], APCDD1 [145], CCL7 [146], PTGS2 [147], TLR2 [148], PIGR (polymeric immunoglobulin receptor) [149], IGFBP5 [150], CCR2 [151], VWF (von Willebrand factor) [152], SLC7A11 [153], NOD2 [154], DMBT1 [155], IL20RA [156], TYMP (thymidine phosphorylase) [144], S100P [157], PDPN (podoplanin) [158], ADAMTS1 [159], ATF3 [160], TIMP1 [161], UCN2 [162], SELE (selectin E) [163], ICAM1 [164], FOSL1 [165], AREG (amphiregulin) [166], PIM2 [167], SLC7A5 [168], CH25H [169], COL5A2 [170], SNAI1 [171], MXRA5 [172], EGR1 [173], TNFRSF17 [174], MDFI (MyoD family inhibitor) [175], SRGN (serglycin) [176], CEACAM6 [177], CCL11 [178], IFNG (interferon gamma) [179], TREM2 [180], INHBA (inhibin subunit beta A) [181], APOE (apolipoprotein E) [182], FGR (FGR proto-oncogene, Src family tyrosine kinase) [183], CTSK (cathepsin K) [184], CCR1 [185], IL6 [186], CTHRC1 [187], PDCD1LG2 [188], SFRP1 [189], CCL4 [190], SNX10 [191], SPP1 [192], CLDN1 [193], CA2 [194], IL11 [195], PHLDA2 [196], NNMT (nicotinamide N-methyltransferase) [197], FGFBP1 [198], RAB31 [199], COL1A1 [200], RNF186 [201], MMP12 [202], MMP2 [203], IL1A [204], ITGA5 [205], CCL1 [206], GPR4 [207], WNT2 [208], GPX2 [209], CD24 [210], PDE4B [211], AQP5 [212], REG1A [213], UBD (ubiquitin D) [214], SPHK1 [215], AOX1 [216], CYP7B1 [217], STC2 [218], TFF2 [219], POSTN (periostin) [220], GZMB (granzyme B) [221], MUC5AC [222], SERPINA3 [223], TWIST1 [224], CCL8 [225], CSF2 [226], PLAU (plasminogen activator, urokinase) [227], CD177 [228], CA9 [229], GFPT2 [230], TDO2 [231], TFF1 [232], STC1 [233], ITLN1 [234], CTLA4 [235], MMP13 [236], LBP (lipopolysaccharide binding protein) [237], CST1 [238], GAS1 [239], KLK6 [240], VSNL1 [241], RETN (resistin) [242], ODAM (odontogenic, ameloblast associated) [243], MPO (myeloperoxidase) [244], HP (haptoglobin) [245], SCGB2A1 [246], CYP24A1 [247], TXNDC5 [248], PDZK1IP1 [249], CEACAM5 [250], MMP10 [251], FOLH1 [252], LAP3 [253], PSAT1 [254], EMILIN2 [255], SRPX2 [256], EGFL6 [257], VCAN (versican) [258], TCN1 [259], CLCA4 [260], ZG16B [261], STEAP4 [262], ACSL4 [263], ADAMTS4 [264], RARRES1 [265], APOC1 [266], SLC5A8 [267], MUC5B [268], CA1 [194], SPINK4 [269], CEMIP (cell migration inducing hyaluronidase 1) [270], MMP7 [271], ANGPTL2 [114], FJX1 [272], MUCL1 [273], MUC6 [274], THBS2 [275], KRT7 [276], CEACAM3 [277], ALDOC (aldolase, fructose-bisphosphate C) [278], CUBN (cubilin) [279], LRAT (lecithin retinol acyltransferase) [280], CBS (cystathionine beta-synthase) [281], CYP2S1 [282], ABCG2 [283], CYP3A4 [284], APOA1 [285], FMO5 [286], GUCA2B [287], FBP1 [288], SULT2B1 [289], AKR1B10 [290], GSTA1 [291], CYP2J2 [292], PDK2 [293], DPEP1 [294], APOB (apolipoprotein B) [295], CES2 [296], SLC23A1 [297], UGT1A1 [298], UGT1A6 [299], GUCA2A [300], KHK (ketohexokinase) [301], CYP2D6 [302], CYP3A5 [303], SLC16A9 [304], CYP2C8 [305], NPC1L1 [306], ACY1 [307], DDC (dopa decarboxylase) [308],B4GALNT2 [309], UGT2B7 [310], CRABP1 [159], HMGCS2 [311], PCK1 [312], CYP2C19 [313], FABP2 [314], DEGS2 [315], EDN2 [316], CYP2C9 [313], NR0B2 [317], ALDOB (aldolase, fructose-bisphosphate B) [318], CNR1 [319], UGT1A7 [320], ABCC2 [321], TM4SF4 [322], PFN2 [323], MAOB (monoamine oxidase B) [324], ZDHHC11 [325], FLVCR1 [326], ANXA13 [327], HTR1D [328], ABCB1 [329], PLA2R1 [330], ACE2 [331], F10 [332], SLC6A4 [333], FABP6 [334], SLC22A5 [335], DPP4 [336], GNG4 [337], TMIGD1 [338], ACE (angiotensin I converting enzyme) [339], PHLPP2 [340], USP2 [341], GRIA4 [342], OTOP2 [343], SLC2A5 [344], SLC1A1 [304], HSD17B2 [345], AOC1 [346], ABCG5 [347], ENPEP (glutamyl aminopeptidase) [348], SCIN (scinderin) [349], GFRA1 [350], PLEKHG6 [351], TM4SF5 [352], CHRNA7 [353], CLDN8 [354], SI (sucrase-isomaltase) [355], TM6SF2 [356], TF (transferrin) [357], RNF128 [358], MEP1A [359], LCT (lactase) [360], SLC34A2 [361], AQP3 [362], ALPI (alkaline phosphatase, intestinal) [363], SHISA3 [364], SSTR1 [365], B3GALT5 [366], SCTR (secretin receptor) [367], FGFR3 [368], TMEM236 [369], NEU4 [370], KISS1 [371], BEST4 [372], MUC17 [373], SLC30A10 [374], ZG16 [375], MS4A1 [376], APOM (apolipoprotein M) [377] and GSTA2 [378] were previously reported to be critical for the development of colorectal cancer. These colorectal cancer responsible genes might be candidate genes for CD. Recently, increasing evidence demonstrated that CXCL5 [379], S100A8 [380], LCN2 [381], CXCL1 [382], S100A9 [383], CXCL9 [384], CXCL11 [385], CXCL10 [386], NCF2 [387], SLC11A1 [388], GDF15 [389], IL1RN [390], STAT1 [391], CYP27B1 [392], SOCS3 [393], TLR8 [394], CD55 [395], ADGRG3 [396], CCL3 [397], FCGR2A [398], CCL2 [399], CD14 [400], IGFBP2 [401], PCSK9 [402], IDO1 [403], FOXP3 [404], CD163 [405], CCL7 [406], TLR2 [407], CCR2 [408], IL20RA [409], S100P [410], ADAMTS1 [411], TIMP1 [412], ICAM1 [413], IFNG (interferon gamma) [414], TREM2 [415], APOE (apolipoprotein E) [416], CCR1 [417], IL6 [418], CTHRC1 [419], PDCD1LG2 [420], CCL4 [421], IL11 [422], COL1A1 [423], MMP2 [424], IL1A [425], CD24 [426], POSTN (periostin) [427], GZMB (granzyme B) [428], BCL2A1 [429], CSF2 [430], TDO2 [431], CTLA4 [432], PADI4 [433], MPO (myeloperoxidase) [434], HP (haptoglobin) [435], MUC5B [436], MMP7 [437], PON3 [438], ABHD6 [439], AICDA (activation induced cytidine deaminase) [440], UGT1A6 [441], CYP2D6 [442], CYP3A5 [443], DGAT1 [444], FCRL4 [445], SLC22A4 [446], DPP4 [447], ACE (angiotensin I converting enzyme) [448], SLC5A11 [449], VIPR1 [450], FCRL3 [451], CD160 [452] and IL22RA2 [453] were altered expression in autoimmune diseases. These autoimmune diseases responsible genes might be critical for the development of CD. Recently, study found that CXCL5 [454], S100A12 [455], OSM (oncostatin M) [456], LRG1 [457], LCN2 [458], CXCL1 [459], S100A9 [460], IFITM1 [461], XBP1 [462], MMP3 [457], IFITM3 [463], IL1B [464], GBP5 [465], HGF (hepatocyte growth factor) [466], CXCL9 [467], SLC11A1 [468], IL1RN [469], STAT1 [470], CYP27B1 [471], MMP1 [472], SOCS3 [473], TLR8 [474], CD55 [475], CCL28 [476], FCGR2A [477], CCL2 [478], CFB (complement factor B) [479], CD14 [480], GPR84 [481], PCSK9 [482], FOXP3 [483], LPL (lipoprotein lipase) [484], IL1R2 [485], TLR2 [486], MEFV (MEFV innate immuity regulator, pyrin) [487], VWF (von Willebrand factor) [488], NOD2 [489], DMBT1 [490], HSPA6 [491], TIMP1 [492], ICAM1 [493], EGR1 [494], CCL11 [495], IFNG (interferon gamma) [496], APOE (apolipoprotein E) [497] FGR (FGR proto-oncogene, Src family tyrosine kinase) [498], IL6 [499], SPP1 [192], IL11 [500], RNF186 [501], MMP2 [502], CD24 [503], SPHK1 [504], GZMB (granzyme B) [505], MUC5AC [506], SERPINA3 [507], TWIST1 [508], PLAU (plasminogen activator, urokinase) [509], CA2 [510], CA9 [510], CTLA4 [511], PADI4 [512], MMP13 [513], MPO (myeloperoxidase) [244], LEFTY1 [514], CA1 [515], MMP7 [513], ABCG2 [516], CYP2J2 [517], AICDA (activation induced cytidine deaminase) [518], CYP2D6 [519], CYP3A5 [520], CNR1 [521], TRPV3 [522], ABCB1 [523], SLC22A4 [524], SLC22A5[524], ACE (angiotensin I converting enzyme) [525], PHLPP2 [526], CCR9 [527], AOC1 [528], SI (sucrase-isomaltase) [529], BTNL2 [530] and SLC26A3 [531] are associated with the risk of ulcerative colitis. Many studies have indicated that S100A12 [532], S100A8 [533], OSM (oncostatin M) [534], LCN2 [535], TNFAIP6 [536], S100A9 [533], XBP1 [537], CXCL6 [42], IL1B [464], GBP5 [465], CHI3L1 [538], TREM1 [539], HGF (hepatocyte growth factor) [540], CXCL9 [541], GBP1 [542], ADM (adrenomedullin) [543], CXCL10 [544], SLC11A1 [468], IL1RN [545], STAT1 [546], CLEC5A [547], CYP27B1 [548], MMP1 [549], SOCS3 [550], TLR8 [474], CD55 [551], CXCR2 [552], FCGR2A [41], TFRC (transferrin receptor) [553], CFB (complement factor B) [554], CD14 [555], IGFBP2 [556], CLEC7A [557], IDO1 [558], FOXP3 [559], CD163 [560], ADORA2B [561], TLR2 [562], MEFV (MEFV innate immuity regulator, pyrin) [487], IGFBP5 [563], CCR2 [564], CHRNA5 [565], VWF (von Willebrand factor) [566], NOD2 [567], DMBT1 [568], IL20RA [569], ATF3 [570], TIMP1 [571], FAP (fibroblast activation protein alpha) [572], ICAM1 [493], IL13RA2 [573], EGR1 [494], TNFRSF17 [574], CEACAM6 [575], CCL11 [576], IFNG (interferon gamma) [577], APOE (apolipoprotein E) [578], IL6 [499], CCL4 [579], SPP1 [580], NNMT (nicotinamide N-methyltransferase) [44], COL1A1 [581], MMP2 [582], GPX2 [583], CD24 [584], REG1A [585], UBD (ubiquitin D) [586], AOX1 [587], POSTN (periostin) [588], GZMB (granzyme B) [589], SELP (selectin P) [590], MUC5AC [591], CCL8 [592], PLAU (plasminogen activator, urokinase) [593], CD177 [594], TFF1 [595], CTLA4 [596], MMP13 [597], LBP (lipopolysaccharide binding protein) [598], MPO (myeloperoxidase) [599], LEFTY1 [514], HP (haptoglobin) [600], CEACAM5 [601], FOLH1 [602], TNFAIP2 [603], MMP7 [597], CLEC12A [604], CBS (cystathionine beta-synthase) [605], PEPD (peptidase D) [606], ABCG2 [607], CYP3A4 [608], CYP2J2 [609], SLC23A1 [610], UGT1A1 [611], GUCA2A [612], CYP2D6 [613], APOC3 [614], CYP2C18 [615], DGAT1 [616], SULT1A2 [617], CYP2C19 [618], CNR1 [521], ABCB1 [619], ACE2 [620], SLC6A4 [621], SLC22A4 [622], MGAT3 [623], SLC22A5 [624], DPP4 [625], TMIGD1 [626], ACE (angiotensin I converting enzyme) [525], SLC15A1 [627], CCR9 [628], GP2 [629], CLDN8 [630], SI (sucrase-isomaltase) [631], TF (transferrin) [632], MEP1A [633], LCT (lactase) [634], BTNL2 [635], VIPR1 [636], F11 [637], ALPI (alkaline phosphatase, intestinal) [638], FCRL3 [639], BCHE (butyrylcholinesterase) [640] and SLC26A3 [641] plays a substantial role in CD. S100A8 [642], OSM (oncostatin M) [643], LCN2 [644], S100A9 [642], HGF (hepatocyte growth factor) [645], GDF15 [646], STAT1 [647], MMP1 [648], TLR8 [649], CD55 [650], SOD2 [651], DYSF (dysferlin) [652], FOXP3 [653], TLR2 [654], VWF (von Willebrand factor) [655], COL4A1 [656], IFNG (interferon gamma) [657], APOE (apolipoprotein E) [658], IL6 [659], CCL4 [421], IL11 [660], MMP2 [661], CSF2 [662], PLAU (plasminogen activator, urokinase) [663], CTLA4 [664], MPO (myeloperoxidase) [665], LEFTY1 [666], HP (haptoglobin) [667], CUBN (cubilin) [668], ABCG2 [669], CYP3A4 [670], APOA1 [671], UGT1A1 [672], CYP2D6 [673], CYP2C8 [674], PCK1 [675], ABCC2 [676], FLVCR1 [677] and ACE (angiotensin I converting enzyme) [678] were identified to be closely associated with anemia. These anemia responsible genes might be candidate biomarkers or therapeutic targets for CD. CXCR1 [679], CXCL1 [680], C2 [681], MMP3 [682], TAP1 [683], NOS2 [684], CXCL10 [685], SLC11A1 [686], GDF15 [687], IL1RN [688], PYGL (glycogen phosphorylase L) [689], CYP27B1 [690], SOCS3 [691], CD55 [692], CXCR2 [693], CCL3 [694], CCL2 [695], SOD2 [696], CD14 [697], IGFBP2 [698], PCSK9 [699], [700], IDO1 [701], FOXP3 [702], CD163 [703], LPL (lipoprotein lipase) [704], TLR2 [705], CCR2 [706], VWF (von Willebrand factor) [707], NOD2 [708], ATF3 [709], TIMP1 [682], ICAM1 [710], FFAR2 [711], IFNG (interferon gamma) [712], APOE (apolipoprotein E) [713], IL6 [714], CCL4 [715], CA2[716], MMP12 [717], MMP2 [718], CCL1 [719], CD24 [720], PLAU (plasminogen activator, urokinase) [721], CTLA4 [722], MASP1 [723], MPO (myeloperoxidase) [724], HP (haptoglobin) [725], MMP10 [726], CD300E [727], CUBN (cubilin) [728], SLC19A3 [729], APOB (apolipoprotein B) [730], APOC3 [731], HMGCS2 [732], OTC (ornithine transcarbamylase) [733], ACE2 [734], SLC22A4 [735], SLC22A5 [735], DPP4 [736], ACE (angiotensin I converting enzyme) [737], SLC6A19 [738], BTNL2 [739], FCRL3 [451], BCHE (butyrylcholinesterase) [740] and APOM (apolipoprotein M) [741] might be involved in type 1 diabetes mellitus. These type 1 diabetes mellitus responsible genes might be important participant in CD. IL1B [742], C4BPB [743], ADM (adrenomedullin) [744], GDF15 [745], IL1RN [746], SOD2 [747], TLR2 [748], VWF (von Willebrand factor) [749], APOE (apolipoprotein E) [750], IL6 [751], CCL4 [752], AQP5 [753], SELP (selectin P) [754], PLAU (plasminogen activator, urokinase) [755], MASP1 [756], MPO (myeloperoxidase) [757], MMP10 [758], EMILIN2 [759], ACE2 [760], F10 [761], ACE (angiotensin I converting enzyme) [762], F11 [763] and APOM (apolipoprotein M) [764] participates in the occurrence and development of coagulation and fibrinolysis. These coagulation and fibrinolysis responsible genes might be relevant to the pathological basis of CD susceptibility. Collectively, results of enriched GO and REACTOME pathway analysis were positively correlated with experimental findings. However, further studies are needed to explore and confirm the potentially significant GO terms and pathways for CD and to achieve a comprehensive understanding of this process.

Construction of PPI network a nd modules of DEGs might be helpful for understanding the relationship of developmental CD. TMEM25 is confirmed to be altered expressed in colorectal cancer [765]. This colorectal cancer gene might be linked with progression of CD. To the date, there are still no reports on the correlation between the hub genes of MNDA (myeloid cell nuclear differentiation antigen), FBXO6, CLEC4D, NAMPT (nicotinamide phosphoribosyltransferase), CLEC4E, MME (membrane metalloendopeptidase), SLC39A4, SLC9A3R1 and GDA (guanine deaminase) with CD and its associated complications. These hub genes might serve as potential biomarkers for CD.

We built a miRNA-hub gene regulatory network and TF-hub gene regulatory network of hub genes in CD based on the Cytoscape software. Finally, we got hub genes, miRNAs and TFs of CD. CDKN2B [766], hsa-mir-629-5p [767], hsa-mir-146a [768], ESR1 [769], PRDM1 [770] and GATA2 [771] have been discovered to be involved in the CD. CDKN2B [772], hsa-mir-146a [768] and ESR1 [773] have been thought of as a specific and exclusive biomarkers for ulcerative colitis. CDKN2B [774], hsa-mir-25-5p [775], hsa-mir-146a [768], hsa-mir-148a-3p [776], NFYA (Nuclear Transcription Factor Y Subunit Alpha) [777], FOXA1 [778], ESR1 [779], ARID3A [780], PRDM1 [781], GATA2 [782], HOXA5 [783] and BRCA1 [784] were significantly associated in colorectal cancer. Theses colorectal cancer biomarkers might plays important regulatory roles in CD. hsa-mir-146a [785], GATA2 [786] and BRCA1 [787] have been identified as a key biomarkers in anemia. These anemia biomarkers might closely related to the occurrence of CD. Recently, increasing evidence demonstrated that hsa-mir-146a [788] and hsa-mir-148a-3p [789] were altered expressed in type 1 diabetes mellitus. These type 1 diabetes mellitus responsible biomarkers might constitute a potential therapeutic target of CD. hsa-mir-4651 [790] levels are correlated with disease severity in patients with coagulation and fibrinolysis. This coagulation and fibrinolysis responsible biomarker might associated with CD. PRDM1 [791] and GATA2 [792] are associated with progression to autoimmune disease. These autoimmune disease related biomarkers might be associated with the development and progression of CD. CKAP4, LAMP3, PLAUR (plasminogen activator, urokinase receptor), STX11, TUBB2B, SLC39A5, hsa-mir-3714, hsa-mir-518f-5p, hsa-mir-522-5p, hsa-mir-641, hsa-mir-3179, FOS (Fos Proto-Oncogene, AP-1 Transcription Factor Subunit) and JUND (JunD Proto-Oncogene, AP-1 Transcription Factor Subunit) were defined as novel biomarkers that might provide new ideas for further studies on CD.

In conclusion, we identified several key genes that are potentially associated with the development of CD using bioinformatics analyses of DEGs between patients with CD and healthy controls. These genes and their pathways will further our understanding of CD etiology, and help improve diagnosis, prevention, and treatment. Our findings suggest that the MDFI, MNDA, FBXO6, TFRC, STAT1, DPP4, MME, SLC39A4, APOA1 and TMEM25 can be considered candidate biomarkers or therapeutic targets for CD.

## Acknowledgement

I thank Rebekah Karns, Cincinnati Children’s Hospital Medical Center, Gastroenterology, Hepatology, & Nutrition, Cincinnati, OH, USA, very much, the author who deposited their NGS dataset GSE101794, into the public GEO database.

## Conflict of interest

The authors declare that they have no conflict of interest.

## Ethical approval

This article does not contain any studies with human participants or animals performed by any of the authors.

## Informed consent

No informed consent because this study does not contain human or animals participants.

## Availability of data and materials

The datasets supporting the conclusions of this article are available in the GEO (Gene Expression Omnibus) (https://www.ncbi.nlm.nih.gov/geo/) repository. [(GSE101794) https://www.ncbi.nlm.nih.gov/geo/query/acc.cgi?acc=GSE101794]

## Consent for publication

Not applicable.

## Competing interests

The authors declare that they have no competing interests.

## Author Contributions

B. V. - Writing original draft, and review and editing

C. V. - Software and investigation

## Notes

### Competing Interest Statement

The authors have declared no competing interest.

### Summary of Updates

abstract only corrected

## References

1. Verburgt CM, Ghiboub M, Benninga MA, de Jonge WJ, Van Limbergen JE. Nutritional Therapy Strategies in Pediatric Crohn’s Disease. Nutrients. 2021;13(1):212. doi:10.3390/nu13010212

2. Turunen P, Ashorn M, Auvinen A, Iltanen S, Huhtala H, Kolho KL. Long-term health outcomes in pediatric inflammatory bowel disease: a population-based study. Inflamm Bowel Dis. 2009;15(1):56–62. doi:10.1002/ibd.20558

3. Benchimol EI, Bernstein CN, Bitton A, Carroll MW, Singh H, Otley AR, Vutcovici M, El-Matary W, Nguyen GC, Griffiths AM, et al. Trends in Epidemiology of Pediatric Inflammatory Bowel Disease in Canada: Distributed Network Analysis of Multiple Population-Based Provincial Health Administrative Databases. Am J Gastroenterol. 2017;112(7):1120–1134. doi:10.1038/ajg.2017.97

4. Rutgeerts P, Löfberg R, Malchow H, Lamers C, Olaison G, Jewell D, Danielsson A, Goebell H, Thomsen OO, Lorenz-Meyer H, et al. A comparison of budesonide with prednisolone for active Crohn’s disease. N Engl J Med. 1994;331(13):842–845. doi:10.1056/NEJM199409293311304

5. Valério de Azevedo S, Maltez C, Lopes AI. Pediatric Crohn’s disease, iron deficiency anemia and intravenous iron treatment: a follow-up study. Scand J Gastroenterol. 2017;52(1):29–33. doi:10.1080/00365521.2016.1224381

6. Nedelkopoulou N, Vadamalayan B, Vergani D, Mieli-Vergani G. Anti-TNFα Treatment in Children and Adolescents With Combined Inflammatory Bowel Disease and Autoimmune Liver Disease. J Pediatr Gastroenterol Nutr. 2018;66(1):100–105. doi:10.1097/MPG.0000000000001759

7. Lu S, Gong J, Tan Y, Liu D. Epidemiologic Association between Inflammatory Bowel Diseases and Type 1 Diabetes Mellitus: a Meta-Analysis. J Gastrointestin Liver Dis. 2020;29(3):407–413. doi:10.15403/jgld-798

8. Weber P, Husemann S, Vielhaber H, Zimmer KP, Nowak-Göttl U. Coagulation and fibrinolysis in children, adolescents, and young adults with inflammatory bowel disease. J Pediatr Gastroenterol Nutr. 1999;28(4):418–422. doi:10.1097/00005176-199904000-00013

9. Olén O, Erichsen R, Sachs MC, Pedersen L, Halfvarson J, Askling J, Ekbom A, Sørensen HT, Ludvigsson JF. Colorectal cancer in Crohn’s disease: a Scandinavian population-based cohort study. Lancet Gastroenterol Hepatol. 2020;5(5):475–484. doi:10.1016/S2468-1253(20)30005-4

10. Sim WH, Wagner J, Cameron DJ, Catto-Smith AG, Bishop RF, Kirkwood CD. Expression profile of genes involved in pathogenesis of pediatric Crohn’s disease. J Gastroenterol Hepatol. 2012;27(6):1083–1093. doi:10.1111/j.1440-1746.2011.06973.x

11. Wei J, Feng J. Signaling pathways associated with inflammatory bowel disease. Recent Pat Inflamm Allergy Drug Discov. 2010;4(2):105–117. doi:10.2174/187221310791163071

12. Bouzidi A, Mesbah-Amroun H, Boukercha A, Benhassine F, Belboueb R, Berkouk K, Messadi W, Touil-Boukoffa C.Association between MDR1 gene polymorphisms and the risk of Crohn’s disease in a cohort of Algerian pediatric patients. Pediatr Res. 2016;80(6):837–843. doi:10.1038/pr.2016.163

13. Potdar AA, Dube S, Naito T, Li K, Botwin G, Haritunians T, Li D, Casero D, Yang S, Bilsborough J, et al. Altered Intestinal ACE2 Levels Are Associated With Inflammation, Severe Disease, and Response to Anti-Cytokine Therapy in Inflammatory Bowel Disease. Gastroenterology. 2021;160(3):809–822.e7. doi:10.1053/j.gastro.2020.10.041

14. Lee YJ, Hwang EH, Park JH, Shin JH, Kang B, Kim SY. NUDT15 variant is the most common variant associated with thiopurine-induced early leukopenia and alopecia in Korean pediatric patients with Crohn’s disease. Eur J Gastroenterol Hepatol. 2016;28(4):475–478. doi:10.1097/MEG.0000000000000564

15. Marcil V, Sinnett D, Seidman E, Boudreau F, Gendron FP, Beaulieu JF, Menard D, Lambert M, Bitton A, Sanchez R, et al. Association between genetic variants in the HNF4A gene and childhood-onset Crohn’s disease. Genes Immun. 2012;13(7):556–565. doi:10.1038/gene.2012.37

16. Dubinsky MC, Wang D, Picornell Y, Wrobel I, Katzir L, Quiros A, Dutridge D, Wahbeh G, Silber G, Bahar R, et al. IL-23 receptor (IL-23R) gene protects against pediatric Crohn’s disease. Inflamm Bowel Dis. 2007;13(5):511–515. doi:10.1002/ibd.20126

17. Cordes F, Foell D, Ding JN, Varga G, Bettenworth D. Differential regulation of JAK/STAT-signaling in patients with ulcerative colitis and Crohn’s disease. World J Gastroenterol. 2020;26(28):4055–4075. doi:10.3748/wjg.v26.i28.4055

18. Zhao F, Zheng T, Gong W, Wu J, Xie H, Li W, Zhang R, Liu P, Liu J, Wu X, et al. Extracellular vesicles package dsDNA to aggravate Crohn’s disease by activating the STING pathway. Cell Death Dis. 2021;12(9):815. doi:10.1038/s41419-021-04101-z

19. Kopiasz Ł, Dziendzikowska K, Gajewska M, Oczkowski M, Majchrzak-Kuligowska K, Królikowski T, Gromadzka-Ostrowska J. Effects of Dietary Oat Beta-Glucans on Colon Apoptosis and Autophagy through TLRs and Dectin-1 Signaling Pathways-Crohn’s Disease Model Study. Nutrients. 2021;13(2):321. doi:10.3390/nu13020321

20. Zhang D, Zhu P, Liu Y, Shu Y, Zhou JY, Jiang F, Chen T, Yang BL, Chen YG. Total flavone of Abelmoschus manihot ameliorates Crohn’s disease by regulating the NF□κB and MAPK signaling pathways. Int J Mol Med. 2019;44(1):324–334. doi:10.3892/ijmm.2019.4180

21. Zhang J, Wang XJ, Wu LJ, Yang L, Yang YT, Zhang D, Hong J, Li XY, Dong XQ, Guo XC, et al. Herb-partitioned moxibustion alleviates colonic inflammation in Crohn’s disease rats by inhibiting hyperactivation of the NLRP3 inflammasome via regulation of the P2X7R-Pannexin-1 signaling pathway. PLoS One. 2021;16(5):e0252334. doi:10.1371/journal.pone.0252334

22. Lefterova MI, Suarez CJ, Banaei N, Pinsky BA. Next-Generation Sequencing for Infectious Disease Diagnosis and Management: A Report of the Association for Molecular Pathology. J Mol Diagn. 2015;17(6):623–634. doi:10.1016/j.jmoldx.2015.07.004

23. Charbit-Henrion F, Parlato M, Hanein S, Duclaux-Loras R, Nowak J, Begue B, Rakotobe S, Bruneau J, Fourrage C, Alibeu O, et al. Diagnostic Yield of Next-generation Sequencing in Very Early-onset Inflammatory Bowel Diseases: A Multicentre Study. J Crohns Colitis. 2018;12(9):1104–1112. doi:10.1093/ecco-jcc/jjy068

24. Haberman Y, Schirmer M, Dexheimer PJ, Karns R, Braun T, Kim MO, Walters TD, Baldassano RN, Noe JD, Rosh J, et al. Age-of-diagnosis dependent ileal immune intensification and reduced alpha-defensin in older versus younger pediatric Crohn Disease patients despite already established dysbiosis. Mucosal Immunol. 2019;12(2):491–502. doi:10.1038/s41385-018-0114-4

25. Clough E, Barrett T. The Gene Expression Omnibus Database. Methods Mol Biol. 2016;1418:93–110. doi:10.1007/978-1-4939-3578-9_5

26. Ritchie ME, Phipson B, Wu D, Hu Y, Law CW, Shi W, Smyth GK.limma powers differential expression analyses for RNA-sequencing and microarray studies. Nucleic Acids Res. 2015;43(7):e47. doi:10.1093/nar/gkv007

27. Reimand J, Kull M, Peterson H, Hansen J, Vilo J. g:Profiler--a web-based toolset for functional profiling of gene lists from large-scale experiments. Nucleic Acids Res. 2007;35(Web Server issue):W193–W200. doi:10.1093/nar/gkm226

28. Thomas PD. The Gene Ontology and the Meaning of Biological Function. Methods Mol Biol. 2017;1446:15□24. doi:10.1007/978-1-4939-3743-1_2

29. Fabregat A, Jupe S, Matthews L, Sidiropoulos K, Gillespie M, Garapati P, Haw R, Jassal B, Korninger F, May B et al The Reactome Pathway Knowledgebase. Nucleic Acids Res. 2018;46(D1):D649–D655. doi:10.1093/nar/gkx1132

30. Alanis-Lobato G, Andrade-Navarro MA, Schaefer MH. HIPPIE v2.0: enhancing meaningfulness and reliability of protein-protein interaction networks. Nucleic Acids Res. 2017;45(D1):D408–D414. doi:10.1093/nar/gkw985

31. Otasek D, Morris JH, Bouças J, Pico AR, Demchak B. Cytoscape Automation: empowering workflow-based network analysis. Genome Biol. 2019;20(1):185. doi:10.1186/s13059-019-1758-4

32. Luo X, Guo L, Dai XJ, Wang Q, Zhu W, Miao X, Gong H. Abnormal intrinsic functional hubs in alcohol dependence: evidence from a voxelwise degree centrality analysis. Neuropsychiatr Dis Treat. 2017;13:2011–2020. doi:10.2147/NDT.S142742

33. Li Y, Li W, Tan Y, Liu F, Cao Y, Lee KY. Hierarchical Decomposition for Betweenness Centrality Measure of Complex Networks. Sci Rep. 2017;7:46491.. doi:10.1038/srep46491

34. Gilbert M, Li Z, Wu XN, Rohr L, Gombos S, Harter K, Schulze WX. Comparison of path-based centrality measures in protein-protein interaction networks revealed proteins with phenotypic relevance during adaptation to changing nitrogen environments. J Proteomics. 2021;235:104114. doi:10.1016/j.jprot.2021.104114

35. Li G, Li M, Wang J, Li Y, Pan Y. United Neighborhood Closeness Centrality and Orthology for Predicting Essential Proteins. IEEE/ACM Trans Comput Biol Bioinform. 2020;17(4):1451–1458. doi:10.1109/TCBB.2018.2889978

36. Zaki N, Efimov D, Berengueres J. Protein complex detection using interaction reliability assessment and weighted clustering coefficient. BMC Bioinformatics. 2013;14:163. doi:10.1186/1471-2105-14

37. Fan Y, Xia J (2018) miRNet-Functional Analysis and Visual Exploration of miRNA-Target Interactions in a Network Context. Methods Mol Biol 1819:215–233. doi:10.1007/978-1-4939-8618-7_10

38. Zhou G, Soufan O, Ewald J, Hancock REW, Basu N, Xia J (2019) NetworkAnalyst 3.0: a visual analytics platform for comprehensive gene expression profiling and meta-analysis. Nucleic Acids Res 47:W234–W241. doi:10.1093/nar/gkz240

39. Robin X, Turck N, Hainard A, Tiberti N, Lisacek F, Sanchez JC, Müller M. pROC: an open-source package for R and S+ to analyze and compare ROC curves. BMC Bioinformatics 2011;12:77. doi:10.1186/1471-2105-12-77

40. Petagna L, Antonelli A, Ganini C, Bellato V, Campanelli M, Divizia A, Efrati C, Franceschilli M, Guida AM, Ingallinella S, et al. Pathophysiology of Crohn’s disease inflammation and recurrence. Biol Direct. 2020;15(1):23. doi:10.1186/s13062-020-00280-5

41. Lee YH, Choi SJ, Ji JD, Song GG. Associations between functional FCGR2A R131H and FCGR3A F158V polymorphisms and responsiveness to TNF blockers in spondyloarthropathy, psoriasis and Crohn’s disease: a meta-analysis. Pharmacogenomics. 2016;17(13):1465–1477. doi:10.2217/pgs.16.27

42. Gijsbers K, Van Assche G, Joossens S, Struyf S, Proost P, Rutgeerts P, Geboes K, Van Damme J. et al. CXCR1-binding chemokines in inflammatory bowel diseases: down-regulated IL-8/CXCL8 production by leukocytes in Crohn’s disease and selective GCP-2/CXCL6 expression in inflamed intestinal tissue. Eur J Immunol. 2004;34(7):1992–2000. doi:10.1002/eji.200324807

43. Hashash JG, Beatty PL, Critelli K, Hartman DJ, Regueiro M, Tamim H, Regueiro MD, Binion DG, Finn OJ. Altered Expression of the Epithelial Mucin MUC1 Accompanies Endoscopic Recurrence of Postoperative Crohn’s Disease. J Clin Gastroenterol. 2021;55(2):127–133. doi:10.1097/MCG.0000000000001340

44. Wnorowski A, Wnorowska S, Kurzepa J, Parada-Turska J. Alterations in Kynurenine and NAD+ Salvage Pathways during the Successful Treatment of Inflammatory Bowel Disease Suggest HCAR3 and NNMT as Potential Drug Targets. Int J Mol Sci. 2021;22(24):13497. doi:10.3390/ijms222413497

45. Kyodo R, Takeuchi I, Narumi S, Shimizu H, Hata K, Yoshioka T, Tanase-Nakao K, Shimizu T, Arai K. Novel biallelic mutations in the DUOX2 gene underlying very early-onset inflammatory bowel disease: A case report. Clin Immunol. 2022;238:109015. doi:10.1016/j.clim.2022.109015

46. Nourse JP, Lea R, Crooks P, Wright G, Tran H, Catalano J, Brighton T, Grigg A, Marlton P, Gandhi MK. The KIR2DS2/DL2 genotype is associated with adult persistent/chronic and relapsed immune thrombocytopenia independently of FCGR3a-158 polymorphisms. Blood Coagul Fibrinolysis. 2012;23(1):45–50. doi:10.1097/MBC.0b013e32834d7ce3

47. Nakashima T, Yokoyama A, Ohnishi H, Hamada H, Ishikawa N, Haruta Y, Hattori N, Tanigawa K, Kohno N. Circulating KL-6/MUC1 as an independent predictor for disseminated intravascular coagulation in acute respiratory distress syndrome. J Intern Med. 2008;263(4):432–439. doi:10.1111/j.1365-2796.2008.01929.x

48. Ying HQ, Wang F, Chen XL, He BS, Pan YQ, Jie C, Liu X, Cao WJ, Peng HX, Lin K, et al. FCGR2A, FCGR3A polymorphisms and therapeutic efficacy of anti-EGFR monoclonal antibody in metastatic colorectal cancer. Oncotarget. 2015;6(29):28071–28083. doi:10.18632/oncotarget.4872

49. Huang D, Feng X, Liu Y, Deng Y, Chen H, Chen D, Fang L, Cai Y, Liu H, Wang L, et al. AQP9-induced cell cycle arrest is associated with RAS activation and improves chemotherapy treatment efficacy in colorectal cancer. Cell Death Dis. 2017;8(6):e2894. doi:10.1038/cddis.2017.282

50. Liu B, Pan S, Xiao Y, Liu Q, Xu J, Jia L. LINC01296/miR-26a/GALNT3 axis contributes to colorectal cancer progression by regulating O-glycosylated MUC1 via PI3K/AKT pathway. J Exp Clin Cancer Res. 2018;37(1):316. doi:10.1186/s13046-018-0994-x

51. Yang X, Wei W, Tan S, Guo L, Qiao S, Yao B, Wang Z. Identification and verification of HCAR3 and INSL5 as new potential therapeutic targets of colorectal cancer. World J Surg Oncol. 2021;19(1):248. doi:10.1186/s12957-021-02335-x

52. Bao Z, Zeng W, Zhang D, Wang L, Deng X, Lai J, Li J, Gong J, Xiang G. SNAIL Induces EMT and Lung Metastasis of Tumours Secreting CXCL2 to Promote the Invasion of M2-Type Immunosuppressed Macrophages in Colorectal Cancer. Int J Biol Sci. 2022;18(7):2867–2881. doi:10.7150/ijbs.66854

53. Zhang X, Han J, Feng L, Zhi L, Jiang D, Yu B, Zhang Z, Gao B, Zhang C, Li M, et al. DUOX2 promotes the progression of colorectal cancer cells by regulating the AKT pathway and interacting with RPL3. Carcinogenesis. 2021;42(1):105–117. doi:10.1093/carcin/bgaa056

54. Zhang L, Zhou R, Zhang W, Yao X, Li W, Xu L, Sun X, Zhao L. Cysteine-rich intestinal protein 1 suppresses apoptosis and chemosensitivity to 5-fluorouracil in colorectal cancer through ubiquitin-mediated Fas degradation. J Exp Clin Cancer Res. 2019;38(1):120. doi:10.1186/s13046-019-1117-z

55. Ye K, Li Y, Zhao W, Wu N, Liu N, Li R, Chen L, He M, Lu B, Wang X, et al. Knockdown of Tubulin Polymerization Promoting Protein Family Member 3 inhibits cell proliferation and invasion in human colorectal cancer. J Cancer. 2017;8(10):1750–1758. doi:10.7150/jca.18943

56. Xi L, Liu Q, Zhang W, Luo L, Song J, Liu R, Wei S, Wang Y. Circular RNA circCSPP1 knockdown attenuates doxorubicin resistance and suppresses tumor progression of colorectal cancer via miR-944/FZD7 axis. Cancer Cell Int. 2021;21(1):153. doi:10.1186/s12935-021-01855-6

57. Robledo G, Márquez A, Dávila-Fajardo CL, Ortego-Centeno N, Rubio JL, Garrido Ede R, Sánchez-Román J, García-Hernández FJ, Ríos-Fernández R, González-Escribano MF, et al. Association of the FCGR3A-158F/V gene polymorphism with the response to rituximab treatment in Spanish systemic autoimmune disease patients. DNA Cell Biol. 2012;31(12):1671–1677. doi:10.1089/dna.2012.1799

58. Lee YH, Bae SC, Seo YH, Kim JH, Choi SJ, Ji JD, Song GG. Association between FCGR3B copy number variations and susceptibility to autoimmune diseases: a meta-analysis. Inflamm Res. 2015;64(12):983–991. doi:10.1007/s00011-015-0882-1

59. Yen JH, Xu S, Park YS, Ganea D, Kim KC. Higher susceptibility to experimental autoimmune encephalomyelitis in Muc1-deficient mice is associated with increased Th1/Th17 responses. Brain Behav Immun. 2013;29:70–81. doi:10.1016/j.bbi.2012.12.004

60. Zhang C, Chen J, Wang H, Chen J, Zheng MJ, Chen XG, Zhang L, Liang CZ, Zhan CS.IL-17 exacerbates experimental autoimmune prostatitis via CXCL1/CXCL2-mediated neutrophil infiltration. Andrologia. 2022;e14455. doi:10.1111/and.14455

61. MacFie TS, Poulsom R, Parker A, Warnes G, Boitsova T, Nijhuis A, Suraweera N, Poehlmann A, Szary J, Feakins R, et al. DUOX2 and DUOXA2 form the predominant enzyme system capable of producing the reactive oxygen species H2O2 in active ulcerative colitis and are modulated by 5-aminosalicylic acid. Inflamm Bowel Dis. 2014;20(3):514–524. doi:10.1097/01.MIB.0000442012.45038.0e

62. Walana W, Ye Y, Li M, Wang J, Wang B, Cheng JW, Gordon JR, Li F.IL-8 antagonist, CXCL8(3-72)K11R/G31P coupled with probiotic exhibit variably enhanced therapeutic potential in ameliorating ulcerative colitis. Biomed Pharmacother. 2018;103:253–261. doi:10.1016/j.biopha.2018.04.008

63. Asano K, Matsumoto T, Umeno J, Hirano A, Esaki M, Hosono N, Matsui T, Kiyohara Y, Nakamura Y, Kubo M, et al. Impact of allele copy number of polymorphisms in FCGR3A and FCGR3B genes on susceptibility to ulcerative colitis. Inflamm Bowel Dis. 2013;19(10):2061–2068. doi:10.1097/MIB.0b013e318298118e

64. Kvorjak M, Ahmed Y, Miller ML, Sriram R, Coronnello C, Hashash JG, Hartman DJ, Telmer CA, Miskov-Zivanov N, Finn OJ, et al. Cross-talk between Colon Cells and Macrophages Increases ST6GALNAC1 and MUC1-sTn Expression in Ulcerative Colitis and Colitis-Associated Colon Cancer. Cancer Immunol Res. 2020;8(2):167–178. doi:10.1158/2326-6066.CIR-19-0514

65. Abke S, Neumeier M, Weigert J, Wehrwein G, Eggenhofer E, Schäffler A, Maier K, Aslanidis C, Schölmerich J, Buechler C. et al. Adiponectin-induced secretion of interleukin-6 (IL-6), monocyte chemotactic protein-1 (MCP-1, CCL2) and interleukin-8 (IL-8, CXCL8) is impaired in monocytes from patients with type I diabetes. Cardiovasc Diabetol. 2006;5:17. doi:10.1186/1475-2840-5-17

66. Anjosa ZP, Santos MM, Rodrigues NJ, Lacerda GA, Araujo J, Silva JA, Tavares NA, Guimarães RL, Crovella S, Brandão LA. Polymorphism in ficolin-1 (FCN1) gene is associated with an earlier onset of type 1 diabetes mellitus in children and adolescents from northeast Brazil. J Genet. 2016;95(4):1031–1034. doi:10.1007/s12041-016-0719-x

67. Kerr RO, Dalmasso AP, Kaplan ME. Erythrocyte-bound C5 and C6 in autoimmune hemolytic anemia. J Immunol. 1971;107(4):1209–1210.

68. Ampuero J, Del Campo JA, Rojas L, Calleja JL, Cabezas J, Lens S, Crespo J, Forns X, Andrade RJ, Fernández I, et al. Role of ITPA and SLC28A2 genes in the prediction of anaemia associated with protease inhibitor plus ribavirin and peginterferon in hepatitis C treatment. J Clin Virol. 2015;68:56–60. doi:10.1016/j.jcv.2015.05.010

69. Miura S, Tsuzuki Y, Hokari R, Ishii H. Modulation of intestinal immune system by dietary fat intake: relevance to Crohn’s disease. J Gastroenterol Hepatol. 1998;13(12):1183–1190.

70. Denson LA, Jurickova I, Karns R, Shaw KA, Cutler DJ, Okou DT, Dodd A, Quinn K, Mondal K, Aronow BJ, et al. Clinical and Genomic Correlates of Neutrophil Reactive Oxygen Species Production in Pediatric Patients With Crohn’s Disease. Gastroenterology. 2018;154(8):2097–2110. doi:10.1053/j.gastro.2018.02.016

71. Carbonetto P, Stephens M. Integrated enrichment analysis of variants and pathways in genome-wide association studies indicates central role for IL-2 signaling genes in type 1 diabetes, and cytokine signaling genes in Crohn’s disease. PLoS Genet. 2013;9(10):e1003770. doi:10.1371/journal.pgen.1003770

72. Derkacz A, Olczyk P, Jura-Półtorak A, Olczyk K, Komosinska-Vassev K. The Diagnostic Usefulness of Circulating Profile of Extracellular Matrix Components: Sulfated Glycosaminoglycans (sGAG), Hyaluronan (HA) and Extracellular Part of Syndecan-1 (sCD138) in Patients with Crohn’s Disease and Ulcerative Colitis. J Clin Med. 2021;10(8):1722. doi:10.3390/jcm10081722

73. Hou Y, Sun X, Gheinani PT, Guan X, Sharma S, Zhou Y, Jin C, Yang Z, Naren AP, Yin J, et al. Epithelial SMYD5 Exaggerates IBD by Downregulating Mitochondrial Functions via Post-Translational Control of PGC-1α Stability. Cell Mol Gastroenterol Hepatol. 2022;S2352-345X(22)00088-1. doi:10.1016/j.jcmgh.2022.05.006

74. Moret-Tatay I, Iborra M, Cerrillo E, Tortosa L, Nos P, Beltrán B. Possible Biomarkers in Blood for Crohn’s Disease: Oxidative Stress and MicroRNAs-Current Evidences and Further Aspects to Unravel. Oxid Med Cell Longev. 2016;2016:2325162. doi:10.1155/2016/2325162

75. Scoville EA, Allaman MM, Brown CT, Motley AK, Horst SN, Williams CS, Koyama T, Zhao Z, Adams DW, Beaulieu DB, et al. Alterations in Lipid, Amino Acid, and Energy Metabolism Distinguish Crohn’s Disease from Ulcerative Colitis and Control Subjects by Serum Metabolomic Profiling. Metabolomics. 2018;14(1):17. doi:10.1007/s11306-017-1311-y

76. Steiner SJ, Pfefferkorn MD, Fitzgerald JF, Denne SC. Carbohydrate and lipid metabolism following infliximab therapy in pediatric Crohn’s disease. Pediatr Res. 2008;64(6):673–676. doi:10.1203/PDR.0b013e318186dde2

77. Luo M, Hu Z, Kong Y, Li L. MicroRNA-432-5p inhibits cell migration and invasion by targeting CXCL5 in colorectal cancer. Exp Ther Med. 2021;21(4):301. doi:10.3892/etm.2021.9732

78. Cui C, Zhang R, Gu F, Pei Y, Sun L, Huang Y, Niu G, Li J. Plasma CXCL3 Levels Are Associated with Tumor Progression and an Unfavorable Colorectal Cancer Prognosis. J Immunol Res. 2022;2022:1336509. doi:10.1155/2022/1336509

79. Kurebayashi H, Goi T, Shimada M, Tagai N, Naruse T, Nakazawa T, Kimura Y, Hirono Y, Yamaguchi A. Prokineticin 2 (PROK2) is an important factor for angiogenesis in colorectal cancer. Oncotarget. 2015;6(28):26242–26251. doi:10.18632/oncotarget.4385

80. Sahami-Fard MH, Yazd EF, Khazaei Z. The Relationship of CXCR1, IκBα and HIF-1α Expression Levels with Clinicopathological Parameters in Colorectal Cancer. Clin Lab. 2019;65(5):10.7754/Clin.Lab.2018.181004. doi:10.7754/Clin.Lab.2018.181004

81. Yan K, Xu X, Wu T, Li J, Cao G, Li Y, Ji Z.Knockdown of PYCR1 inhibits proliferation, drug resistance and EMT in colorectal cancer cells by regulating STAT3-Mediated p38 MAPK and NF-κB signalling pathway. Biochem Biophys Res Commun. 2019;520(2):486–491. doi:10.1016/j.bbrc.2019.10.059

82. Kim MS, Louwagie J, Carvalho B, Terhaar Sive Droste JS, Park HL, Chae YK, Yamashita K, Liu J, Ostrow KL, et al. Promoter DNA methylation of oncostatin m receptor-beta as a novel diagnostic and therapeutic marker in colon cancer. PLoS One. 2009;4(8):e6555. doi:10.1371/journal.pone.0006555

83. De Mattia E, Polesel J, Roncato R, Labriet A, Bignucolo A, Gagno S, Buonadonna A, D’Andrea M, Lévesque E, Jonker D, et al. IL15RA and SMAD3 Genetic Variants Predict Overall Survival in Metastatic Colorectal Cancer Patients Treated with FOLFIRI Therapy: A New Paradigm. Cancers (Basel). 2021;13(7):1705. doi:10.3390/cancers13071705

84. Zhong B, Cheng B, Huang X, Xiao Q, Niu Z, Chen YF, Yu Q, Wang W, Wu XJ. Colorectal cancer-associated fibroblasts promote metastasis by up-regulating LRG1 through stromal IL-6/STAT3 signaling. Cell Death Dis. 2021;13(1):16. doi:10.1038/s41419-021-04461-6

85. Zhang W, Pan R, Lu M, Zhang Q, Lin Z, Qin Y, Wang Z, Gong S, Lin H, Chong S, et al. Epigenetic induction of lipocalin 2 expression drives acquired resistance to 5-fluorouracil in colorectal cancer through integrin β3/SRC pathway. Oncogene. 2021;40(45):6369–6380. doi:10.1038/s41388-021-02029-4

86. Liu Z, Wei P, Yang Y, Cui W, Cao B, Tan C, Yu B, Bi R, Xia K, Chen W, et al. BATF2 Deficiency Promotes Progression in Human Colorectal Cancer via Activation of HGF/MET Signaling: A Potential Rationale for Combining MET Inhibitors with IFNs. Clin Cancer Res. 2015;21(7):1752–1763. doi:10.1158/1078-0432.CCR-14-1564

87. Park YL, Kim HP, Ock CY, Min DW, Kang JK, Lim YJ, Song SH, Han SW, Kim TY. EMT-mediated regulation of CXCL1/5 for resistance to anti-EGFR therapy in colorectal cancer. Oncogene. 2022;41(14):2026–2038. doi:10.1038/s41388-021-01920-4

88. Chen H, Huang J, Chen C, Jiang Y, Feng X, Liao Y, Yang Z. NGFR Increases the Chemosensitivity of Colorectal Cancer Cells by Enhancing the Apoptotic and Autophagic Effects of 5-fluorouracil via the Activation of S100A9. Front Oncol. 2021;11:652081.doi:10.3389/fonc.2021.652081

89. Kelemen A, Carmi I, Oszvald Á, Lőrincz P, Petővári G, Tölgyes T, Dede K, Bursics A, Buzás EI, Wiener Z. IFITM1 expression determines extracellular vesicle uptake in colorectal cancer. Cell Mol Life Sci. 2021;78(21-22):7009–7024. doi:10.1007/s00018-021-03949-w

90. He Y, Kan W, Li Y, Hao Y, Huang A, Gu H, Wang M, Wang Q, Chen J, Sun Z, et al. A potent and selective small molecule inhibitor of myoferlin attenuates colorectal cancer progression. Clin Transl Med. 2021;11(2):e289. doi:10.1002/ctm2.289

91. Zhao Y, Zhang W, Huo M, Wang P, Liu X, Wang Y, Li Y, Zhou Z, Xu N, Zhu H. XBP1 regulates the protumoral function of tumor-associated macrophages in human colorectal cancer. Signal Transduct Target Ther. 2021;6(1):357. doi:10.1038/s41392-021-00761-7

92. Ji Y, Li J, Li P, Wang L, Yang H, Jiang G. C/EBPβ Promotion of MMP3-Dependent Tumor Cell Invasion and Association with Metastasis in Colorectal Cancer. Genet Test Mol Biomarkers. 2018;22(1):5–10. doi:10.1089/gtmb.2017.0113

93. Ling A, Löfgren-Burström A, Larsson P, Li X, Wikberg ML, Öberg Å, Stenling R, Edin S, Palmqvist R. TAP1 down-regulation elicits immune escape and poor prognosis in colorectal cancer. Oncoimmunology. 2017;6(11):e1356143. doi:10.1080/2162402X.2017.1356143

94. Xiang Y, Yao X, Chen K, Wang X, Zhou J, Gong W, Yoshimura T, Huang J, Wang R, Wu Y, et al. The G-protein coupled chemoattractant receptor FPR2 promotes malignant phenotype of human colon cancer cells. Am J Cancer Res. 2016;6(11):2599–2610.

95. Xu L, Duda DG, di Tomaso E, Ancukiewicz M, Chung DC, Lauwers GY, Samuel R, Shellito P, Czito BG, Lin PC, et al. Direct evidence that bevacizumab, an anti-VEGF antibody, up-regulates SDF1alpha, CXCR4, CXCL6, and neuropilin 1 in tumors from patients with rectal cancer. Cancer Res. 2009;69(20):7905–7910. doi:10.1158/0008-5472.CAN-09-2099

96. Rong Z, Luo Z, Fu Z, Zhang P, Li T, Zhang J, Zhu Z, Yu Z, Li Q, Qiu Z, et al. The novel circSLC6A6/miR-1265/C2CD4A axis promotes colorectal cancer growth by suppressing p53 signaling pathway. J Exp Clin Cancer Res. 2021;40(1):324. doi:10.1186/s13046-021-02126-y

97. Li D, Peng Z, Tang H, Wei P, Kong X, Yan D, Huang F, Li Q, Le X, Li Q, et al. KLF4-mediated negative regulation of IFITM3 expression plays a critical role in colon cancer pathogenesis. Clin Cancer Res. 2011;17(11):3558–3568. doi:10.1158/1078-0432.CCR-10-2729

98. Qian H, Zhang D, Bao C. Two variants of Interleukin-1B gene are associated with the decreased risk, clinical features, and better overall survival of colorectal cancer: a two-center case-control study. Aging (Albany NY). 2018;10(12):4084–4092. doi:10.18632/aging.101695

99. Mao H, Sheng J, Jia J, Wang C, Zhang S, Li H, He F. Aberrant SLC6A14 Expression Promotes Proliferation and Metastasis of Colorectal Cancer via Enhancing the JAK2/STAT3 Pathway. Onco Targets Ther. 2021;14:379–392. doi:10.2147/OTT.S288709

100. Li SQ, Su N, Gong P, Zhang HB, Liu J, Wang D, Sun YP, Zhang Y, Qian F, Zhao B, et al. The Expression of Formyl Peptide Receptor 1 is Correlated with Tumor Invasion of Human Colorectal Cancer. Sci Rep. 2017;7(1):5918. doi:10.1038/s41598-017-06368-9

101. Schirripa M, Zhang W, Yang D, Cao S, Okazaki S, Loupakis F, Berger MD, Ning Y, Miyamoto Y, Suenaga M, et al. NOS2 polymorphisms in prediction of benefit from first-line chemotherapy in metastatic colorectal cancer patients. PLoS One. 2018;13(3):e0193640. doi:10.1371/journal.pone.0193640

102. Li H, Zhou X, Zhang H, Jiang J, Fu H, Wang F. Combined Efficacy of CXCL5, STC2, and CHI3L1 in the Diagnosis of Colorectal Cancer. J Oncol. 2022;2022:7271514. doi:10.1155/2022/7271514

103. Gu C, Cai J, Xu Z, Zhou S, Ye L, Yan Q, Zhang Y, Fang Y, Liu Y, Tu C, et al. MiR-532-3p suppresses colorectal cancer progression by disrupting the ETS1/TGM2 axis-mediated Wnt/β-catenin signaling. Cell Death Dis. 2019;10(10):739. doi:10.1038/s41419-019-1962-x

104. Lu S, Catalano C, Huhn S, Pardini B, Partu L, Vymetalkova V, Vodickova L, Levy M, Buchler T, Hemminki K, et al. Single nucleotide polymorphisms within MUC4 are associated with colorectal cancer survival. PLoS One. 2019;14(5):e0216666. doi:10.1371/journal.pone.0216666

105. Roh SA, Kwon YH, Lee JL, Kim SK, Kim JC. SLAMF7 and TREM1 Mediate Immunogenic Cell Death in Colorectal Cancer Cells: Focus on Microsatellite Stability. Anticancer Res. 2021;41(11):5431–5444. doi:10.21873/anticanres.15355

106. Sun G, Wu L, Sun G, Shi X, Cao H, Tang W. WNT5a in Colorectal Cancer: Research Progress and Challenges. Cancer Manag Res. 2021;13:2483–2498. doi:10.2147/CMAR.S289819

107. Joosten SPJ, Spaargaren M, Clevers H, Pals ST. Hepatocyte growth factor/MET and CD44 in colorectal cancer: partners in tumorigenesis and therapy resistance. Biochim Biophys Acta Rev Cancer. 2020;1874(2):188437. doi:10.1016/j.bbcan.2020.188437

108. Wu Z, Huang X, Han X, Li Z, Zhu Q, Yan J, Yu S, Jin Z, Wang Z, Zheng Q, et al. The chemokine CXCL9 expression is associated with better prognosis for colorectal carcinoma patients. Biomed Pharmacother. 2016;78:8–13. doi:10.1016/j.biopha.2015.12.021

109. Britzen-Laurent N, Lipnik K, Ocker M, Naschberger E, Schellerer VS, Croner RS, Vieth M, Waldner M, Steinberg P, Hohenadl C, et al. GBP-1 acts as a tumor suppressor in colorectal cancer cells. Carcinogenesis. 2013;34(1):153–162. doi:10.1093/carcin/bgs310

110. Guo AJ, Wang FJ, Ji Q, Geng HW, Yan X, Wang LQ, Tie WW, Liu XY, Thorne RF, Liu G, et al. Proteome Analyses Reveal S100A11, S100P, and RBM25 Are Tumor Biomarkers in Colorectal Cancer. Proteomics Clin Appl. 2021;15(1):e2000056. doi:10.1002/prca.202000056

111. Wang L, Gala M, Yamamoto M, Pino MS, Kikuchi H, Shue DS, Shirasawa S, Austin TR, Lynch MP, Rueda BR, et al. Adrenomedullin is a therapeutic target in colorectal cancer. Int J Cancer. 2014;134(9):2041–2050. doi:10.1002/ijc.28542

112. Gao YJ, Liu L, Li S, Yuan GF, Li L, Zhu HY, Cao GY. Down-regulation of CXCL11 inhibits colorectal cancer cell growth and epithelial-mesenchymal transition. Onco Targets Ther. 2018;11:7333–7343. doi:10.2147/OTT.S16787

113. Shang S, Yang YW, Chen F, Yu L, Shen SH, Li K, Cui B, Lv XX, Zhang C, Yang C, et al. TRIB3 reduces CD8+ T cell infiltration and induces immune evasion by repressing the STAT1-CXCL10 axis in colorectal cancer. Sci Transl Med. 2022;14(626):eabf0992. doi:10.1126/scitranslmed.abf0992

114. He J, Xu J, Yu X, Zhu H, Zeng Y, Fan D, Yi X. Overexpression of ANGPTL2 and LILRB2 as predictive and therapeutic biomarkers for metastasis and prognosis in colorectal cancer. Int J Clin Exp Pathol. 2018;11(5):2281–2294.

115. Zheng H, Yu S, Zhu C, Guo T, Liu F, Xu Y. HIF1α promotes tumor chemoresistance via recruiting GDF15-producing TAMs in colorectal cancer. Exp Cell Res. 2021;398(2):112394. doi:10.1016/j.yexcr.2020.112394

116. Ibrahimi M, Moossavi M, Mojarad EN, Musavi M, Mohammadoo-Khorasani M, Shahsavari Z. Positive correlation between interleukin-1 receptor antagonist gene 86bp VNTR polymorphism and colorectal cancer susceptibility: a case-control study. Immunol Res. 2019;67(1):151–156. doi:10.1007/s12026-018-9034-3

117. Tanaka A, Zhou Y, Ogawa M, Shia J, Klimstra DS, Wang JY, Roehrl MH. STAT1 as a potential prognosis marker for poor outcomes of early stage colorectal cancer with microsatellite instability. PLoS One. 2020;15(4):e0229252. doi:10.1371/journal.pone.0229252

118. Latacz M, Snarska J, Kostyra E, Wroński K, Fiedorowicz E, Savelkoul H, Jarmołowska B, Płomiński J, Grzybowski R, Cieślińska A. CYP27B1 Gene Polymorphism rs10877012 in Patients Diagnosed with Colorectal Cancer. Nutrients. 2020;12(4):998. doi:10.3390/nu12040998

119. Fedorova MS, Snezhkina AV, Lipatova AV, Pavlov VS, Kobelyatskaya AA, Guvatova ZG, Pudova EA, Savvateeva MV, Ishina IA, Demidova TB, et al. NETO2 Is Deregulated in Breast, Prostate, and Colorectal Cancer and Participates in Cellular Signaling. Front Genet. 2020;11:594933. doi:10.3389/fgene.2020.594933

120. Lei R, Zhao Y, Huang K, Wang Q, Wan K, Li T, Yang H, Lv X. The methylation of SDC2 and TFPI2 defined three methylator phenotypes of colorectal cancer. BMC Gastroenterol. 2022;22(1):88. doi:10.1186/s12876-022-02175-3

121. Chen T, Du D, Chen J, Zhou P, Weinstein JN, Yao L, Liu Y. ZC3H12A Expression in Different Stages of Colorectal Cancer. Oncoscience. 2019;6(3-4):301–311. doi:10.18632/oncoscience.480

122. Wang K, Zheng J, Yu J, Wu Y, Guo J, Xu Z, Sun X. Knockdown of MMP□1 inhibits the progression of colorectal cancer by suppressing the PI3K/Akt/c□myc signaling pathway and EMT. Oncol Rep. 2020;43(4):1103–1112. doi:10.3892/or.2020.7490

123. Saunders AS, Bender DE, Ray AL, Wu X, Morris KT. Colony-stimulating factor 3 signaling in colon and rectal cancers: Immune response and CMS classification in TCGA data. PLoS One. 2021;16(2):e0247233. doi:10.1371/journal.pone.0247233

124. Li L, Zhang J, Peng H, Jiang X, Liu Z, Tian H, Hou S, Xie X, Peng Q, Zhou T. Knockdown of miR-92a suppresses the stemness of colorectal cancer cells via mediating SOCS3. Bioengineered. 2022;13(3):5613–5624. doi:10.1080/21655979.2021.2022267

125. Grimm M, Kim M, Rosenwald A, Heemann U, Germer CT, Waaga-Gasser AM, Gasser M.Toll-like receptor (TLR) 7 and TLR8 expression on CD133+ cells in colorectal cancer points to a specific role for inflammation-induced TLRs in tumourigenesis and tumour progression. Eur J Cancer. 2010;46(15):2849–2857. doi:10.1016/j.ejca.2010.07.017

126. Forse CL, Rahimi M, Diamandis EP, Assarzadegan N, Dawson H, Grin A, Kennedy E, O’Connor B, Messenger DE, Riddell RH, et al. HtrA3 stromal expression is correlated with tumor budding in stage II colorectal cancer. Exp Mol Pathol. 2017;103(1):94–100. doi:10.1016/j.yexmp.2017.07.002

127. Shao K, Pu W, Zhang J, Guo S, Qian F, Glurich I, Jin Q, Ma Y, Ju S, Zhang Z, et al. DNA hypermethylation contributes to colorectal cancer metastasis by regulating the binding of CEBPB and TFCP2 to the CPEB1 promoter. Clin Epigenetics. 2021;13(1):89. doi:10.1186/s13148-021-01071-z

128. Xiao B, Zhang L, Liu H, Fang H, Wang C, Huang B, Liu X, Zhou X, Wang Y. Oncolytic Adenovirus CD55-Smad4 Suppresses Cell Proliferation, Metastasis, and Tumor Stemness in Colorectal Cancer by Regulating Wnt/β-Catenin Signaling Pathway. Biomedicines. 2020;8(12):593. doi:10.3390/biomedicines8120593

129. Wang H, Shao Q, Wang J, Zhao L, Wang L, Cheng Z, Yue C, Chen W, Wang H, Zhang Y. Decreased CXCR2 expression on circulating monocytes of colorectal cancer impairs recruitment and induces Re-education of tumor-associated macrophages. Cancer Lett. 2022;529:112–125. doi:10.1016/j.canlet.2022.01.004

130. Dimberg J, Hugander A, Wågsäter D. Protein expression of the chemokine, CCL28, in human colorectal cancer. Int J Oncol. 2006;28(2):315–319.

131. Yang M, Chen W, Liu H, Yu L, Tang M, Liu Y. Long Non-coding RNA CBR3 Antisense RNA 1 is Downregulated in Colorectal Cancer and Inhibits miR-29a-Mediated Cell Migration and Invasion. Mol Biotechnol. 2022;64(7):773–779. doi:10.1007/s12033-021-00444-2

132. Xie X, Jiang D, Zhou X, Ye X, Yang P, He Y. Recombinant Bacteroides fragilis enterotoxin-1 (rBFT-1) promotes proliferation of colorectal cancer via CCL3-related molecular pathways. Open Life Sci. 2021;16(1):408–418. doi:10.1515/biol-2021-0043

133. Vargas T, Moreno-Rubio J, Herranz J, Cejas P, Molina S, Mendiola M, Burgos E, Custodio AB, De Miguel M, Martín-Hernández R, et al. 3’UTR Polymorphism in ACSL1 Gene Correlates with Expression Levels and Poor Clinical Outcome in Colon Cancer Patients. PLoS One. 2016;11(12):e0168423. doi:10.1371/journal.pone.0168423

134. Nardelli C, Granata I, Nunziato M, Setaro M, Carbone F, Zulli C, Pilone V, Capoluongo ED, De Palma GD, Corcione F, et al. 16S rRNA of Mucosal Colon Microbiome and CCL2 Circulating Levels Are Potential Biomarkers in Colorectal Cancer. Int J Mol Sci. 2021;22(19):10747. doi:10.3390/ijms221910747

135. Paku M, Haraguchi N, Takeda M, Fujino S, Ogino T, Takahashi H, Miyoshi N, Uemura M, Mizushima T, Yamamoto H, et al. SIRT3-Mediated SOD2 and PGC-1α Contribute to Chemoresistance in Colorectal Cancer Cells. Ann Surg Oncol. 2021;28(8):4720–4732. doi:10.1245/s10434-020-09373-x

136. Chen D, Wang H. The clinical and immune features of CD14 in colorectal cancer identified via large-scale analysis. Int Immunopharmacol. 2020;88:106966. doi:10.1016/j.intimp.2020.106966

137. Walterskirchen N, Müller C, Ramos C, Zeindl S, Stang S, Herzog D, Sachet M, Schimek V, Unger L, Gerakopoulos V, et al. Metastatic colorectal carcinoma-associated fibroblasts have immunosuppressive properties related to increased IGFBP2 expression. Cancer Lett. 2022;540:215737. doi:10.1016/j.canlet.2022.215737

138. Alexander PG, McMillan DC, Park JH. A meta-analysis of CD274 (PD-L1) assessment and prognosis in colorectal cancer and its role in predicting response to anti-PD-1 therapy. Crit Rev Oncol Hematol. 2021;157:103147. doi:10.1016/j.critrevonc.2020.103147

139. Yu F, Mou B, Sheng J, Liu J, Qin X, Yu J. DERL3 suppresses colorectal cancer metastasis through negatively regulating MYCN level. Minerva Med. 2020;10.23736/S0026-4806.20.06657-4. doi:10.23736/S0026-4806.20.06657-4

140. Mazzoccoli G, Pazienza V, Panza A, Valvano MR, Benegiamo G, Vinciguerra M, Andriulli A, Piepoli A. ARNTL2 and SERPINE1: potential biomarkers for tumor aggressiveness in colorectal cancer. J Cancer Res Clin Oncol. 2012;138(3):501–511. doi:10.1007/s00432-011-1126-6

141. Li XM, Yuan DY, Liu YH, Zhu L, Qin HK, Yang YB, Li Y, Yan F, Wang YJ. Panax notoginseng saponins prevent colitis-associated colorectal cancer via inhibition IDO1 mediated immune regulation. Chin J Nat Med. 2022;20(4):258–269. doi:10.1016/S1875-5364(22)60179-1

142. Shi Y, Meng L, Zhang C, Zhang F, Fang Y. Extracellular vesicles of Lacticaseibacillus paracasei PC-H1 induce colorectal cancer cells apoptosis via PDK1/AKT/Bcl-2 signaling pathway. Microbiol Res. 2021;255:126921. doi:10.1016/j.micres.2021.126921

143. Liu S, Zhang C, Zhang K, Gao Y, Wang Z, Li X, Cheng G, Wang S, Xue X, Li W, et al. FOXP3 inhibits cancer stem cell self-renewal via transcriptional repression of COX2 in colorectal cancer cells. Oncotarget. 2017;8(27):44694–44704. doi:10.18632/oncotarget.17974

144. Kaidi D, Szeponik L, Yrlid U, Wettergren Y, Bexe Lindskog E. Impact of thymidine phosphorylase and CD163 expression on prognosis in stage II colorectal cancer. Clin Transl Oncol. 2022;10.1007/s12094-022-02839-2. doi:10.1007/s12094-022-02839-2

145. Skopelitou D, Miao B, Srivastava A, Kumar A, Kuswick M, Dymerska D, Paramasivam N, Schlesner M, Lubinski J, Hemminki K, et al. Whole Exome Sequencing Identifies APCDD1 and HDAC5 Genes as Potentially Cancer Predisposing in Familial Colorectal Cancer. Int J Mol Sci. 2021;22(4):1837. doi:10.3390/ijms22041837

146. Ren X, Xiao J, Zhang W, Wang F, Yan Y, Wu X, Zeng Z, He Y, Yang W, Liao W, et al. Inhibition of CCL7 derived from Mo-MDSCs prevents metastatic progression from latency in colorectal cancer. Cell Death Dis. 2021;12(5):484.doi:10.1038/s41419-021-03698-5

147. Venè R, Costa D, Augugliaro R, Carlone S, Scabini S, Casoni Pattacini G, Boggio M, Zupo S, Grillo F, Mastracci L, et al. Evaluation of Glycosylated PTGS2 in Colorectal Cancer for NSAIDS-Based Adjuvant Therapy. Cells. 2020;9(3):683. doi:10.3390/cells9030683

148. Fan L, Xu C, Ge Q, Lin Y, Wong CC, Qi Y, Ye B, Lian Q, Zhuo W, Si J, et al. A. Muciniphila Suppresses Colorectal Tumorigenesis by Inducing TLR2/NLRP3-Mediated M1-Like TAMs. Cancer Immunol Res. 2021;9(10):1111–1124. doi:10.1158/2326-6066.CIR-20-1019

149. Traicoff JL, De Marchis L, Ginsburg BL, Zamora RE, Khattar NH, Blanch VJ, Plummer S, Bargo SA, Templeton DJ, Casey G, et al. Characterization of the human polymeric immunoglobulin receptor (PIGR) 3’UTR and differential expression of PIGR mRNA during colon tumorigenesis. J Biomed Sci. 2003;10(6 Pt 2):792–804. doi:10.1159/000073967

150. Wu K, Zhou M, Wu QX, Yuan SX, Wang DX, Jin JL, Huang J, Yang JQ, Sun WJ, Wan LH, et al. The role of IGFBP-5 in mediating the anti-proliferation effect of tetrandrine in human colon cancer cells. Int J Oncol. 2015;46(3):1205–1213. doi:10.3892/ijo.2014.2800

151. Tu W, Gong J, Zhou Z, Tian D, Wang Z. TCF4 enhances hepatic metastasis of colorectal cancer by regulating tumor-associated macrophage via CCL2/CCR2 signaling. Cell Death Dis. 2021;12(10):882. doi:10.1038/s41419-021-04166-w

152. Garam N, Maláti É, Sinkovits G, Gombos T, Szederjesi A, Barabás L, Gráf L, Kocsis J, Prohászka Z. Platelet Count, ADAMTS13 Activity, von Willebrand Factor Level and Survival in Patients with Colorectal Cancer: 5-Year Follow-up Study. Thromb Haemost. 2018;118(1):123–131. doi:10.1160/TH17-07-0548

153. Zhang L, Liu W, Liu F, Wang Q, Song M, Yu Q, Tang K, Teng T, Wu D, Wang X, et al. IMCA Induces Ferroptosis Mediated by SLC7A11 through the AMPK/mTOR Pathway in Colorectal Cancer. Oxid Med Cell Longev. 2020;2020:1675613. doi:10.1155/2020/1675613

154. Branquinho D, Freire P, Sofia C. NOD2 mutations and colorectal cancer - Where do we stand?. World J Gastrointest Surg. 2016;8(4):284–293. doi:10.4240/wjgs.v8.i4.284

155. Robbe C, Paraskeva C, Mollenhauer J, Michalski JC, Sergi C, Corfield A. DMBT1 expression and glycosylation during the adenoma-carcinoma sequence in colorectal cancer. Biochem Soc Trans. 2005;33(Pt 4):730–732. doi:10.1042/BST0330730

156. Lamichhane S, Mo JS, Sharma G, Choi TY, Chae SC. MicroRNA 452 regulates IL20RA-mediated JAK1/STAT3 pathway in inflammatory colitis and colorectal cancer. Inflamm Res. 2021;70(8):903–914. doi:10.1007/s00011-021-01486-73

157. Schmid F, Dahlmann M, Röhrich H, Kobelt D, Hoffmann J, Burock S, Walther W, Stein U.Calcium-binding protein S100P is a new target gene of MACC1, drives colorectal cancer metastasis and serves as a prognostic biomarker. Br J Cancer. 2022;10.1038/s41416-022-01833-3. doi:10.1038/s41416-022-01833-3

158. Choi SY, Sung R, Lee SJ, Lee TG, Kim N, Yoon SM, Lee EJ, Chae HB, Youn SJ, Park SM. Podoplanin, α-smooth muscle actin or S100A4 expressing cancer-associated fibroblasts are associated with different prognosis in colorectal cancers. J Korean Med Sci. 2013;28(9):1293–1301. doi:10.3346/jkms.2013.28.9.1293

159. Lind GE, Kleivi K, Meling GI, Teixeira MR, Thiis-Evensen E, Rognum TO, Lothe RA. ADAMTS1, CRABP1, and NR3C1 identified as epigenetically deregulated genes in colorectal tumorigenesis. Cell Oncol. 2006;28(5-6):259–272. doi:10.1155/2006/949506

160. Inoue M, Uchida Y, Edagawa M, Hirata M, Mitamura J, Miyamoto D, Taketani K, Sekine S, Kawauchi J, Kitajima S. The stress response gene ATF3 is a direct target of the Wnt/β-catenin pathway and inhibits the invasion and migration of HCT116 human colorectal cancer cells. PLoS One. 2018;13(7):e0194160. doi:10.1371/journal.pone.0194160

161. Yang L, Jiang Q, Li DZ, Zhou X, Yu DS, Zhong J. TIMP1 mRNA in tumor-educated platelets is diagnostic biomarker for colorectal cancer. Aging (Albany NY). 2019;11(20):8998–9012. doi:10.18632/aging.102366

162. Pothoulakis C, Torre-Rojas M, Duran-Padilla MA, Gevorkian J, Zoras O, Chrysos E, Chalkiadakis G, Baritaki S. CRHR2/Ucn2 signaling is a novel regulator of miR-7/YY1/Fas circuitry contributing to reversal of colorectal cancer cell resistance to Fas-mediated apoptosis. Int J Cancer. 2018;142(2):334–346. doi:10.1002/ijc.31064

163. Li N, Xiao H, Shen J, Qiao X, Zhang F, Zhang W, Gao Y, Liu YD. SELE gene as a characteristic prognostic biomarker of colorectal cancer. J Int Med Res. 2021;49(4):3000605211004386. doi:10.1177/03000605211004386

164. Zhang Y, Zhang L, Zheng S, Li M, Xu C, Jia D, Qi Y, Hou T, Wang L, Wang B, et al. Fusobacterium nucleatum promotes colorectal cancer cells adhesion to endothelial cells and facilitates extravasation and metastasis by inducing ALPK1/NF-κB/ICAM1 axis. Gut Microbes. 2022;14(1):2038852. doi:10.1080/19490976.2022.2038852

165. Liu Y, Yue M, Li Z. FOSL1 promotes tumorigenesis in colorectal carcinoma by mediating the FBXL2/Wnt/β-catenin axis via Smurf1. Pharmacol Res. 2021;165:105405. doi:10.1016/j.phrs.2020.105405

166. Bormann F, Stinzing S, Tierling S, Morkel M, Markelova MR, Walter J, Weichert W, Roßner F, Kuhn N, Perner J, et al. Epigenetic regulation of Amphiregulin and Epiregulin in colorectal cancer. Int J Cancer. 2019;144(3):569–581. doi:10.1002/ijc.31892

167. Zhang XH, Yu HL, Wang FJ, Han YL, Yang WL. Pim-2 Modulates Aerobic Glycolysis and Energy Production during the Development of Colorectal Tumors. Int J Med Sci. 2015;12(6):487–493. doi:10.7150/ijms.10982

168. Najumudeen AK, Ceteci F, Fey SK, Hamm G, Steven RT, Hall H, Nikula CJ, Dexter A, Murta T, Race AM, et al. The amino acid transporter SLC7A5 is required for efficient growth of KRAS-mutant colorectal cancer. Nat Genet. 2021;53(1):16–26. doi:10.1038/s41588-020-00753-3

169. Zhou M, Wang S, Liu D, Zhou J. LINC01915 Facilitates the Conversion of Normal Fibroblasts into Cancer-Associated Fibroblasts Induced by Colorectal Cancer-Derived Extracellular Vesicles through the miR-92a-3p/KLF4/CH25H Axis. ACS Biomater Sci Eng. 2021;7(11):5255–5268. doi:10.1021/acsbiomaterials.1c00611

170. Wang J, Jiang YH, Yang PY, Liu F. Increased Collagen Type V α2 (COL5A2) in Colorectal Cancer is Associated with Poor Prognosis and Tumor Progression. Onco Targets Ther. 2021;14:2991–3002. doi:10.2147/OTT.S288422

171. Shen T, Yue C, Wang X, Wang Z, Wu Y, Zhao C, Chang P, Sun X, Wang W. NFATc1 promotes epithelial-mesenchymal transition and facilitates colorectal cancer metastasis by targeting SNAI1. Exp Cell Res. 2021;408(1):112854. doi:10.1016/j.yexcr.2021.112854

172. Wang GH, Yao L, Xu HW, Tang WT, Fu JH, Hu XF, Cui L, Xu XM. Identification of MXRA5 as a novel biomarker in colorectal cancer. Oncol Lett. 2013;5(2):544–548. doi:10.3892/ol.2012.1038

173. Gu H, Li Y, Cui X, Cao H, Hou Z, Ti Y, Liu D, Gao J, Wang Y, Wen P. MICAL1 inhibits colorectal cancer cell migration and proliferation by regulating the EGR1/β-catenin signaling pathway. Biochem Pharmacol. 2022;195:114870. doi:10.1016/j.bcp.2021.114870

174. Chae SC, Yu JI, Uhm TB, Lee SY, Kang DB, Lee JK, Park WC, Yun KJ. The haplotypes of TNFRSF17 polymorphisms are associated with colon cancer in a Korean population. Int J Colorectal Dis. 2012;27(6):701–707. doi:10.1007/s00384-011-1364-8

175. Sui Y, Li X, Oh S, Zhang B, Freeman WM, Shin S, Janknecht R. Opposite Roles of the JMJD1A Interaction Partners MDFI and MDFIC in Colorectal Cancer. Sci Rep. 2020;10(1):8710. doi:10.1038/s41598-020-65536-6

176. Xu Y, Xu J, Yang Y, Zhu L, Li X, Zhao W. SRGN Promotes Colorectal Cancer Metastasis as a Critical Downstream Target of HIF-1α. Cell Physiol Biochem. 2018;48(6):2429–2440. doi:10.1159/000492657

177. Yang T, Wang H, Li M, Yang L, Han Y, Liu C, Zhang B, Wu M, Wang G, Zhang Z, et al. CD151 promotes Colorectal Cancer progression by a crosstalk involving CEACAM6, LGR5 and Wnt signaling via TGFβ1. Int J Biol Sci. 2021;17(3):848–860. doi:10.7150/ijbs.536

178. Cho H, Lim SJ, Won KY, Bae GE, Kim GY, Min JW, Noh BJ. Eosinophils in Colorectal Neoplasms Associated with Expression of CCL11 and CCL24. J Pathol Transl Med. 2016;50(1):45–51. doi:10.4132/jptm.2015.10.16

179. Wang L, Wang Y, Song Z, Chu J, Qu X. Deficiency of interferon-gamma or its receptor promotes colorectal cancer development. J Interferon Cytokine Res. 2015;35(4):273–280. doi:10.1089/jir.2014.0132

180. Kim SM, Kim EM, Ji KY, Lee HY, Yee SM, Woo SM, Yi JW, Yun CH, Choi H, Kang HS. TREM2 Acts as a Tumor Suppressor in Colorectal Carcinoma through Wnt1/β-catenin and Erk Signaling. Cancers (Basel). 2019;11(9):1315. doi:10.3390/cancers11091315

181. He Z, Liang J, Wang B. Inhibin, beta A regulates the transforming growth factor-beta pathway to promote malignant biological behaviour in colorectal cancer. Cell Biochem Funct. 2021;39(2):258–266. doi:10.1002/cbf.3573

182. Zhao Z, Zou S, Guan X, Wang M, Jiang Z, Liu Z, Li C, Lin H, Liu X, Yang R, et al. Apolipoprotein E Overexpression Is Associated With Tumor Progression and Poor Survival in Colorectal Cancer. Front Genet. 2018;9:650. doi:10.3389/fgene.2018.00650

183. Lieu C, Kopetz S. The SRC family of protein tyrosine kinases: a new and promising target for colorectal cancer therapy. Clin Colorectal Cancer. 2010;9(2):89–94. doi:10.3816/CCC.2010.n.012

184. Li R, Zhou R, Wang H, Li W, Pan M, Yao X, Zhan W, Yang S, Xu L, Ding Y, et al. Gut microbiota-stimulated cathepsin K secretion mediates TLR4-dependent M2 macrophage polarization and promotes tumor metastasis in colorectal cancer. Cell Death Differ. 2019;26(11):2447–2463. doi:10.1038/s41418-019-0312-y

185. Yamamoto T, Kawada K, Itatani Y, Inamoto S, Okamura R, Iwamoto M, Miyamoto E, Chen-Yoshikawa TF, Hirai H, Hasegawa S, et al. Loss of SMAD4 Promotes Lung Metastasis of Colorectal Cancer by Accumulation of CCR1+ Tumor-Associated Neutrophils through CCL15-CCR1 Axis. Clin Cancer Res. 2017;23(3):833–844. doi:10.1158/1078-0432.CCR-16-0520

186. Waldner MJ, Foersch S, Neurath MF. Interleukin-6--a key regulator of colorectal cancer development. Int J Biol Sci. 2012;8(9):1248–1253. doi:10.7150/ijbs.4614

187. Zhang XL, Hu LP, Yang Q, Qin WT, Wang X, Xu CJ, Tian GA, Yang XM, Yao LL, Zhu L, et al. CTHRC1 promotes liver metastasis by reshaping infiltrated macrophages through physical interactions with TGF-β receptors in colorectal cancer. Oncogene. 2021;40(23):3959–3973. doi:10.1038/s41388-021-01827-0

188. Huang KC, Chiang SF, Chen TW, Chen WT, Yang PC, Ke TW, Chao KSC. Prognostic relevance of programmed cell death 1 ligand 2 (PDCD1LG2/PD-L2) in patients with advanced stage colon carcinoma treated with chemotherapy. Sci Rep. 2020;10(1):22330. doi:10.1038/s41598-020-79419-3

189. Liu X, Fu J, Bi H, Ge A, Xia T, Liu Y, Sun H, Li D, Zhao Y. DNA methylation of SFRP1, SFRP2, and WIF1 and prognosis of postoperative colorectal cancer patients. BMC Cancer. 2019;19(1):1212. doi:10.1186/s12885-019-6436-0

190. De la Fuente López M, Landskron G, Parada D, Dubois-Camacho K, Simian D, Martinez M, Romero D, Roa JC, Chahuán I, Gutiérrez R, et al. The relationship between chemokines CCL2, CCL3, and CCL4 with the tumor microenvironment and tumor-associated macrophage markers in colorectal cancer. Tumour Biol. 2018;40(11):1010428318810059. doi:10.1177/1010428318810059

191. Zhang S, Yang Z, Bao W, Liu L, You Y, Wang X, Shao L, Fu W, Kou X, Shen W, et al. SNX10 (sorting nexin 10) inhibits colorectal cancer initiation and progression by controlling autophagic degradation of SRC. Autophagy. 2020;16(4):735–749. doi:10.1080/15548627.2019.1632122

192. Giannos P, Triantafyllidis KK, Giannos G, Kechagias KS. SPP1 in infliximab resistant ulcerative colitis and associated colorectal cancer: an analysis of differentially expressed genes. Eur J Gastroenterol Hepatol. 2022;34(6):598–606. doi:10.1097/MEG.0000000000002349

193. Battagin AS, Bertuzzo CS, Carvalho PO, Ortega MM, Marson FAL. Single nucleotide variants c.-13G → C (rs17429833) and c.108C → T (rs72466472) in the CLDN1 gene and increased risk for familial colorectal cancer. Gene. 2021;768:145304. doi:10.1016/j.gene.2020.145304

194. Bekku S, Mochizuki H, Yamamoto T, Ueno H, Takayama E, Tadakuma T. Expression of carbonic anhydrase I or II and correlation to clinical aspects of colorectal cancer. Hepatogastroenterology. 2000;47(34):998–1001.

195. Nishina T, Deguchi Y, Ohshima D, Takeda W, Ohtsuka M, Shichino S, Ueha S, Yamazaki S, Kawauchi M, Nakamura E, et al. Interleukin-11-expressing fibroblasts have a unique gene signature correlated with poor prognosis of colorectal cancer. Nat Commun. 2021;12(1):2281. doi:10.1038/s41467-021-22450-3

196. Ma Z, Lou S, Jiang Z. PHLDA2 regulates EMT and autophagy in colorectal cancer via the PI3K/AKT signaling pathway. Aging (Albany NY). 2020;12(9):7985–8000. doi:10.18632/aging.103117

197. Ogawa M, Tanaka A, Namba K, Shia J, Wang JY, Roehrl MHA. Tumor stromal nicotinamide N-methyltransferase overexpression as a prognostic biomarker for poor clinical outcome in early-stage colorectal cancer. Sci Rep. 2022;12(1):2767. doi:10.1038/s41598-022-06772-w

198. Chen J, Liu QM, Du PC, Ning D, Mo J, Zhu HD, Wang C, Ge QY, Cheng Q, Zhang XW, et al. Type-2 11β-hydroxysteroid dehydrogenase promotes the metastasis of colorectal cancer via the Fgfbp1-AKT pathway. Am J Cancer Res. 2020;10(2):662–673.

199. Yang L, Tian X, Chen X, Lin X, Tang C, Gao Y, Chen S, Ge Z. Upregulation of Rab31 is associated with poor prognosis and promotes colorectal carcinoma proliferation via the mTOR/p70S6K/Cyclin D1 signalling pathway. Life Sci. 2020;257:118126. doi:10.1016/j.lfs.2020.118126

200. Sun J, Liu J, Zhu Q, Xu F, Kang L, Shi X. Hsa_circ_0001806 Acts as a ceRNA to Facilitate the Stemness of Colorectal Cancer Cells by Increasing COL1A1. Onco Targets Ther. 2020;13:6315–6327. doi:10.2147/OTT.S255485

201. Ji Y, Tu X, Hu X, Wang Z, Gao S, Zhang Q, Zhang W, Zhang H, Chen W. The role and mechanism of action of RNF186 in colorectal cancer through negative regulation of NF-κB. Cell Signal. 2020;75:109764. doi:10.1016/j.cellsig.2020.109764

202. VAN Nguyen S, Skarstedt M, Löfgren S, Zar N, Andersson RE, Lindh M, Matussek A, Dimberg J.Gene polymorphism of matrix metalloproteinase-12 and -13 and association with colorectal cancer in Swedish patients. Anticancer Res. 2013;33(8):3247–3250.

203. Sun J, Zhang Z, Chen J, Xue M, Pan X. ELTD1 promotes invasion and metastasis by activating MMP2 in colorectal cancer. Int J Biol Sci. 2021;17(12):3048–3058. doi:10.7150/ijbs.62293

204. Ji H, Lu L, Huang J, Liu Y, Zhang B, Tang H, Sun D, Zhang Y, Shang H, Li Y, et al. IL1A polymorphisms is a risk factor for colorectal cancer in Chinese Han population: a case control study. BMC Cancer. 2019;19(1):181. doi:10.1186/s12885-019-5395-9

205. Yu M, Chu S, Fei B, Fang X, Liu Z. O-GlcNAcylation of ITGA5 facilitates the occurrence and development of colorectal cancer. Exp Cell Res. 2019;382(2):111464. doi:10.1016/j.yexcr.2019.06.009

206. Li Z, Chan K, Qi Y, Lu L, Ning F, Wu M, Wang H, Wang Y, Cai S, Du J. Participation of CCL1 in Snail-Positive Fibroblasts in Colorectal Cancer Contribute to 5-Fluorouracil/Paclitaxel Chemoresistance. Cancer Res Treat. 2018;50(3):894–907. doi:10.4143/crt.2017.356

207. Yu M, Cui R, Huang Y, Luo Y, Qin S, Zhong M. Increased proton-sensing receptor GPR4 signalling promotes colorectal cancer progression by activating the hippo pathway. EBioMedicine. 2019;48:264–276. doi:10.1016/j.ebiom.2019.09.016

208. Kalhor H, Rahimi H, Akbari Eidgahi MR, Teimoori-Toolabi L. Novel Small Molecules against Two Binding Sites of Wnt2 Protein as potential Drug Candidates for Colorectal Cancer: A Structure Based Virtual Screening Approach. Iran J Pharm Res. 2020;19(2):160–174. doi:10.22037/ijpr.2019.15297.13037

209. Emmink BL, Laoukili J, Kipp AP, Koster J, Govaert KM, Fatrai S, Verheem A, Steller EJ, Brigelius-Flohé R, Jimenez CR, et al. GPx2 suppression of H2O2 stress links the formation of differentiated tumor mass to metastatic capacity in colorectal cancer. Cancer Res. 2014;74(22):6717–6730. doi:10.1158/0008-5472.CAN-14-1645

210. Wang TW, Chern E, Hsu CW, Tseng KC, Chao HM. SIRT1-Mediated Expression of CD24 and Epigenetic Suppression of Novel Tumor Suppressor miR-1185-1 Increases Colorectal Cancer Stemness. Cancer Res. 2020;80(23):5257–5269. doi:10.1158/0008-5472.CAN-19-3188

211. Pleiman JK, Irving AA, Wang Z, Toraason E, Clipson L, Dove WF, Deming DA, Newton MA. The conserved protective cyclic AMP-phosphodiesterase function PDE4B is expressed in the adenoma and adjacent normal colonic epithelium of mammals and silenced in colorectal cancer. PLoS Genet. 2018;14(9):e1007611.doi:10.1371/journal.pgen.1007611

212. Wang W, Li Q, Yang T, Li D, Ding F, Sun H, Bai G. Anti-cancer effect of Aquaporin 5 silencing in colorectal cancer cells in association with inhibition of Wnt/β-catenin pathway. Cytotechnology. 2018;70(2):615–624. doi:10.1007/s10616-017-0147-7

213. Astrosini C, Roeefzaad C, Dai YY, Dieckgraefe BK, Jöns T, Kemmner W. REG1A expression is a prognostic marker in colorectal cancer and associated with peritoneal carcinomatosis. Int J Cancer. 2008;123(2):409–413. doi:10.1002/ijc.23466

214. Su H, Qin M, Liu Q, Jin B, Shi X, Xiang Z. Ubiquitin-Like Protein UBD Promotes Cell Proliferation in Colorectal Cancer by Facilitating p53 Degradation. Front Oncol. 2021;11:691347. doi:10.3389/fonc.2021.691347

215. Shen Z, Feng X, Fang Y, Li Y, Li Z, Zhan Y, Lin M, Li G, Ding Y, Deng H. POTEE drives colorectal cancer development via regulating SPHK1/p65 signaling. Cell Death Dis. 2019;10(11):863. doi:10.1038/s41419-019-2046-7

216. Zhang W, Chai W, Zhu Z, Li X. Aldehyde oxidase 1 promoted the occurrence and development of colorectal cancer by up-regulation of expression of CD133. Int Immunopharmacol. 2020;85:106618. doi:10.1016/j.intimp.2020.106618

217. Xi XP, Zhuang J, Teng MJ, Xia LJ, Yang MY, Liu QG, Chen JB.MicroRNA-17 induces epithelial-mesenchymal transition consistent with the cancer stem cell phenotype by regulating CYP7B1 expression in colon cancer. Int J Mol Med. 2016;38(2):499–506. doi:10.3892/ijmm.2016.2624

218. Li H, Zhou X, Zhang H, Jiang J, Fu H, Wang F. Combined Efficacy of CXCL5, STC2, and CHI3L1 in the Diagnosis of Colorectal Cancer. J Oncol. 2022;2022:7271514. doi:10.1155/2022/7271514

219. Dubeykovskaya ZA, Duddempudi PK, Deng H, Valenti G, Cuti KL, Nagar K, Tailor Y, Guha C, Kitajewski J, Wang TC. Therapeutic potential of adenovirus-mediated TFF2-CTP-Flag peptide for treatment of colorectal cancer. Cancer Gene Ther. 2019;26(1-2):48–57. doi:10.1038/s41417-018-0036-z

220. Sueyama T, Kajiwara Y, Mochizuki S, Shimazaki H, Shinto E, Hase K, Ueno H. Periostin as a key molecule defining desmoplastic environment in colorectal cancer. Virchows Arch. 2021;478(5):865–874. doi:10.1007/s00428-020-02965-8

221. Daemen A, Udyavar AR, Sandmann T, Li C, Bosch LJW, O’Gorman W, Li Y, Au-Yeung A, Takahashi C, Kabbarah O, et al. Transcriptomic profiling of adjuvant colorectal cancer identifies three key prognostic biological processes and a disease specific role for granzyme B. PLoS One. 2021;16(12):e0262198. doi:10.1371/journal.pone.0262198

222. Pothuraju R, Rachagani S, Krishn SR, Chaudhary S, Nimmakayala RK, Siddiqui JA, Ganguly K, Lakshmanan I, Cox JL, Mallya K, et al. Molecular implications of MUC5AC-CD44 axis in colorectal cancer progression and chemoresistance. Mol Cancer. 2020;19(1):37. doi:10.1186/s12943-020-01156-y

223. Dimberg J, Ström K, Löfgren S, Zar N, Hugander A, Matussek A. Expression of the serine protease inhibitor serpinA3 in human colorectal adenocarcinomas. Oncol Lett. 2011;2(3):413–418. doi:10.3892/ol.2011.280

224. Shen X, Hu X, Mao J, Wu Y, Liu H, Shen J, Yu J, Chen W. The long noncoding RNA TUG1 is required for TGF-β/TWIST1/EMT-mediated metastasis in colorectal cancer cells. Cell Death Dis. 2020;11(1):65.doi:10.1038/s41419-020-2254-1

225. Klupp F, Schuler S, Kahlert C, Halama N, Franz C, Mayer P, Schmidt T, Ulrich A. Evaluation of the inflammatory markers CCL8, CXCL5, and LIF in patients with anastomotic leakage after colorectal cancer surgery. Int J Colorectal Dis. 2020;35(7):1221–1230. doi:10.1007/s00384-020-03582-2

226. Xu Z, Zhang Y, Xu M, Zheng X, Lin M, Pan J, Ye C, Deng Y, Jiang C, Lin Y, et al. Demethylation and Overexpression of CSF2 are Involved in Immune Response, Chemotherapy Resistance, and Poor Prognosis in Colorectal Cancer. Onco Targets Ther. 2019;12:11255–11269. doi:10.2147/OTT.S216829

227. Lin M, Zhang Z, Gao M, Yu H, Sheng H, Huang J. MicroRNA-193a-3p suppresses the colorectal cancer cell proliferation and progression through downregulating the PLAU expression. Cancer Manag Res. 2019;11:5353–5363. doi:10.2147/CMAR.S208233

228. Zhou G, Peng K, Song Y, Yang W, Shu W, Yu T, Yu L, Lin M, Wei Q, Chen C, et al. CD177+ neutrophils suppress epithelial cell tumourigenesis in colitis-associated cancer and predict good prognosis in colorectal cancer. Carcinogenesis. 2018;39(2):272–282. doi:10.1093/carcin/bgx142

229. Guan SS, Wu CT, Liao TZ, Lin KL, Peng CL, Shih YH, Weng MF, Chen CT, Yeh CH, Wang YC, et al. A novel 111indium-labeled dual carbonic anhydrase 9-targeted probe as a potential SPECT imaging radiotracer for detection of hypoxic colorectal cancer cells. Eur J Pharm Biopharm. 2021;168:38–52. doi:10.1016/j.ejpb.2021.08.004

230. Liu L, Pan Y, Ren X, Zeng Z, Sun J, Zhou K, Liang Y, Wang F, Yan Y, Liao W, et al. GFPT2 promotes metastasis and forms a positive feedback loop with p65 in colorectal cancer. Am J Cancer Res. 2020;10(8):2510–2522.

231. Lee R, Li J, Li J, Wu CJ, Jiang S, Hsu WH, Chakravarti D, Chen P, LaBella KA, Li J, et al. Synthetic Essentiality of Tryptophan 2,3-Dioxygenase 2 in APC-Mutated Colorectal Cancer. Cancer Discov. 2022;12(7):1702–1717. doi:10.1158/2159-8290.CD-21-0680

232. Yusufu A, Shayimu P, Tuerdi R, Fang C, Wang F, Wang H. TFF3 and TFF1 expression levels are elevated in colorectal cancer and promote the malignant behavior of colon cancer by activating the EMT process. Int J Oncol. 2019;55(4):789–804. doi:10.3892/ijo.2019.4854

233. Peña C, Céspedes MV, Lindh MB, Kiflemariam S, Mezheyeuski A, Edqvist PH, Hägglöf C, Birgisson H, Bojmar L, Jirström K, et al. STC1 expression by cancer-associated fibroblasts drives metastasis of colorectal cancer. Cancer Res. 2013;73(4):1287–1297. doi:10.1158/0008-5472.CAN-12-1875

234. Chen L, Jin XH, Luo J, Duan JL, Cai MY, Chen JW, Feng ZH, Guo AM, Wang FW, et al. ITLN1 inhibits tumor neovascularization and myeloid derived suppressor cells accumulation in colorectal carcinoma. Oncogene. 2021;40(40):5925–5937. doi:10.1038/s41388-021-01965-5

235. Cui K, Yao S, Zhang H, Zhou M, Liu B, Cao Y, Fei B, Huang S, Huang Z. Identification of an immune overdrive high-risk subpopulation with aberrant expression of FOXP3 and CTLA4 in colorectal cancer. Oncogene. 2021;40(11):2130–2145. doi:10.1038/s41388-021-01677-w

236. Ou B, Zhao J, Guan S, Feng H, Wangpu X, Zhu C, Zong Y, Ma J, Sun J, Shen X, et al. CCR4 promotes metastasis via ERK/NF-κB/MMP13 pathway and acts downstream of TNF-α in colorectal cancer. Oncotarget. 2016;7(30):47637–47649. doi:10.18632/oncotarget.10256

237. Turgunov Y, Ogizbayeva A, Akhmaltdinova L, Shakeyev K. Lipopolysaccharide-binding protein as a risk factor for development of infectious and inflammatory postsurgical complications in colorectal cancer paients. Contemp Oncol (Pozn). 2021;25(3):198–203. doi:10.5114/wo.2021.110051

238. Jiang J, Liu HL, Tao L, Lin XY, Yang YD, Tan SW, Wu B. Let□7d inhibits colorectal cancer cell proliferation through the CST1/p65 pathway. Int J Oncol. 2018;53(2):781–790. doi:10.3892/ijo.2018.4419

239. Li Q, Qin Y, Wei P, Lian P, Li Y, Xu Y, Li X, Li D, Cai S. Gas1 Inhibits Metastatic and Metabolic Phenotypes in Colorectal Carcinoma. Mol Cancer Res. 2016;14(9):830–840. doi:10.1158/1541-7786.MCR-16-0032

240. Tieng FYF, Abu N, Sukor S, Mohd Azman ZA, Mahamad Nadzir N, Lee LH, Mutalib NSA. L1CAM, CA9, KLK6, HPN, and ALDH1A1 as Potential Serum Markers in Primary and Metastatic Colorectal Cancer Screening. Diagnostics (Basel). 2020;10(7):444. doi:10.3390/diagnostics10070444

241. He C, Liu W, Xiong Y, Wang Y, Pan L, Luo L, Tu Y, Song R, Chen W. VSNL1 Promotes Cell Proliferation, Migration, and Invasion in Colorectal Cancer by Binding with COL10A1. Ann Clin Lab Sci. 2022;52(1):60–72.

242. Farahani H, Mahmoudi T, Asadi A, Nobakht H, Dabiri R, Hamta A. Insulin Resistance and Colorectal Cancer Risk: the Role of Elevated Plasma Resistin Levels. J Gastrointest Cancer. 2020;51(2):478–483. doi:10.1007/s12029-019-00260-7

243. Yu M, Mu Y, Qi Y, Qin S, Qiu Y, Cui R, Zhong M. Odontogenic ameloblast-associated protein (ODAM) inhibits human colorectal cancer growth by promoting PTEN elevation and inactivating PI3K/AKT signaling. Biomed Pharmacother. 2016;84:601–607. doi:10.1016/j.biopha.2016.09.076

244. Garrity-Park M, Loftus EV Jr, Sandborn WJ, Smyrk TC. Myeloperoxidase immunohistochemistry as a measure of disease activity in ulcerative colitis: association with ulcerative colitis-colorectal cancer, tumor necrosis factor polymorphism and RUNX3 methylation. Inflamm Bowel Dis. 2012;18(2):275–283. doi:10.1002/ibd.21681

245. Mariño-Crespo Ó, Cuevas-Álvarez E, Harding AL, Murdoch C, Fernández-Briera A, Gil-Martín E. Haptoglobin expression in human colorectal cancer. Histol Histopathol. 2019;34(8):953–963. doi:10.14670/HH-18-100

246. Munakata K, Uemura M, Takemasa I, Ozaki M, Konno M, Nishimura J, Hata T, Mizushima T, Haraguchi N, Noura S, et al. SCGB2A1 is a novel prognostic marker for colorectal cancer associated with chemoresistance and radioresistance. Int J Oncol. 2014;44(5):1521–1528. doi:10.3892/ijo.2014.2316

247. He L, Li H, Pan C, Hua Y, Peng J, Zhou Z, Zhao Y, Lin M. Squalene epoxidase promotes colorectal cancer cell proliferation through accumulating calcitriol and activating CYP24A1-mediated MAPK signaling. Cancer Commun (Lond). 2021;41(8):726–746. doi:10.1002/cac2.12187

248. Tan F, Zhu H, He X, Yu N, Zhang X, Xu H, Pei H. Role of TXNDC5 in tumorigenesis of colorectal cancer cells: In vivo and in vitro evidence. Int J Mol Med. 2018;42(2):935–945. doi:10.3892/ijmm.2018.3664

249. Rivero M, Peinado-Serrano J, Muñoz-Galvan S, Espinosa-Sánchez A, Suarez-Martinez E, Felipe-Abrio B, Fernández-Fernández MC, Ortiz MJ, Carnero A. MAP17 (PDZK1IP1) and pH2AX are potential predictive biomarkers for rectal cancer treatment efficacy. Oncotarget. 2018;9(68):32958–32971. doi:10.18632/oncotarget.26010

250. Huskey ALW, Merner ND. An investigation into the role of inherited CEACAM gene family variants and colorectal cancer risk. BMC Res Notes. 2022;15(1):26. doi:10.1186/s13104-022-05907-6

251. Klupp F, Neumann L, Kahlert C, Diers J, Halama N, Franz C, Schmidt T, Koch M, Weitz J, Schneider M, et al. Serum MMP7, MMP10 and MMP12 level as negative prognostic markers in colon cancer patients. BMC Cancer. 2016;16:494. doi:10.1186/s12885-016-2515-7

252. Eklöf V, Van Guelpen B, Hultdin J, Johansson I, Hallmans G, Palmqvist R. The reduced folate carrier (RFC1) 80G > A and folate hydrolase 1 (FOLH1) 1561C > T polymorphisms and the risk of colorectal cancer: a nested case-referent study. Scand J Clin Lab Invest. 2008;68(5):393–401. doi:10.1080/00365510701805431

253. Yang Q, Roehrl MH, Wang JY. Proteomic profiling of antibody-inducing immunogens in tumor tissue identifies PSMA1, LAP3, ANXA3, and maspin as colon cancer markers. Oncotarget. 2017;9(3):3996–4019. doi:10.18632/oncotarget.23583

254. Li S, Yang H, Li W, Liu JY, Ren LW, Yang YH, Ge BB, Zhang YZ, Fu WQ, Zheng XJ, et al. ADH1C inhibits progression of colorectal cancer through the ADH1C/PHGDH /PSAT1/serine metabolic pathway. Acta Pharmacol Sin. 2022;10.1038/s41401-022-00894-7. doi:10.1038/s41401-022-00894-7

255. Andreuzzi E, Fejza A, Polano M, Poletto E, Camicia L, Carobolante G, Tarticchio G, Todaro F, Di Carlo E, Scarpa M, et al. Colorectal cancer development is affected by the ECM molecule EMILIN-2 hinging on macrophage polarization via the TLR-4/MyD88 pathway. J Exp Clin Cancer Res. 2022;41(1):60. doi:10.1186/s13046-022-02271-y

256. Øster B, Linnet L, Christensen LL, Thorsen K, Ongen H, Dermitzakis ET, Sandoval J, Moran S, Esteller M, Hansen TF, et al. Non-CpG island promoter hypomethylation and miR-149 regulate the expression of SRPX2 in colorectal cancer. Int J Cancer. 2013;132(10):2303–2315. doi:10.1002/ijc.27921

257. Sung TY, Huang HL, Cheng CC, Chang FL, Wei PL, Cheng YW, Huang CC, Lee YC, HuangFu WC, Pan SL. Sung TY, Huang HL, Cheng CC, et al. EGFL6 promotes colorectal cancer cell growth and mobility and the anti-cancer property of anti-EGFL6 antibody. Cell Biosci. 2021;11(1):53. doi:10.1186/s13578-021-00561-0

258. Hope C, Emmerich PB, Papadas A, Pagenkopf A, Matkowskyj KA, Van De Hey DR, Payne SN, Clipson L, Callander NS, Hematti P, et al. Versican-Derived Matrikines Regulate Batf3-Dendritic Cell Differentiation and Promote T Cell Infiltration in Colorectal Cancer. J Immunol. 2017;199(5):1933–1941. doi:10.4049/jimmunol.1700529

259. Zhu X, Yi K, Hou D, Huang H, Jiang X, Shi X, Xing C. Clinicopathological Analysis and Prognostic Assessment of Transcobalamin I (TCN1) in Patients with Colorectal Tumors. Med Sci Monit. 2020;26:e923828. doi:10.12659/MSM.923828

260. Li H, Huang B. miR-19a targeting CLCA4 to regulate the proliferation, migration, and invasion of colorectal cancer cells. Eur J Histochem. 2022;66(1):3381. doi:10.4081/ejh.2022.3381

261. Escudero-Paniagua B, Bartolomé RA, Rodríguez S, De Los Ríos V, Pintado L, Jaén M, Lafarga M, Fernández-Aceñero MJ, Casal JI. PAUF/ZG16B promotes colorectal cancer progression through alterations of the mitotic functions and the Wnt/β-catenin pathway. Carcinogenesis. 2020;41(2):203–213. doi:10.1093/carcin/bgz093

262. Liao Y, Zhao J, Bulek K, Tang F, Chen X, Cai G, Jia S, Fox PL, Huang E, Pizarro TT, et al. Inflammation mobilizes copper metabolism to promote colon tumorigenesis via an IL-17-STEAP4-XIAP axis. Nat Commun. 2020;11(1):900. doi:10.1038/s41467-020-14698-y

263. Dai G, Wang D, Ma S, Hong S, Ding K, Tan X, Ju W. ACSL4 promotes colorectal cancer and is a potential therapeutic target of emodin. Phytomedicine. 2022;102:154149. doi:10.1016/j.phymed.2022.154149

264. Shang XQ, Liu KL, Li Q, Lao YQ, Li NS, Wu J. ADAMTS4 is upregulated in colorectal cancer and could be a useful prognostic indicator of colorectal cancer. Rev Assoc Med Bras (1992). 2020;66(1):42–47. doi:10.1590/1806-9282.66.1.42

265. Wu CC, Shyu RY, Chou JM, Jao SW, Chao PC, Kang JC, Wu ST, Huang SL, Jiang SY. RARRES1 expression is significantly related to tumour differentiation and staging in colorectal adenocarcinoma. Eur J Cancer. 2006;42(4):557–565. doi:10.1016/j.ejca.2005.11.015

266. Shen HY, Wei FZ, Liu Q. Differential analysis revealing APOC1 to be a diagnostic and prognostic marker for liver metastases of colorectal cancer. World J Clin Cases. 2021;9(16):3880–3894. doi:10.12998/wjcc.v9.i16.3880

267. Brim H, Kumar K, Nazarian J, Hathout Y, Jafarian A, Lee E, Green W, Smoot D, Park J, Nouraie M, et al. SLC5A8 gene, a transporter of butyrate: a gut flora metabolite, is frequently methylated in African American colon adenomas. PLoS One. 2011;6(6):e20216. doi:10.1371/journal.pone.0020216

268. Iranmanesh H, Majd A, Mojarad EN, Zali MR, Hashemi M. Investigating the Relationship Between the Expression Level of Mucin Gene Cluster (MUC2, MUC5A, and MUC5B) and Clinicopathological Characterization of Colorectal Cancer. Galen Med J. 2021;10:e2030. doi:10.31661/gmj.v10i0.2030

269. Wang X, Yu Q, Ghareeb WM, Zhang Y, Lu X, Huang Y, Huang S, Sun Y, Lin J, Liu J, et al. Downregulated SPINK4 is associated with poor survival in colorectal cancer. BMC Cancer. 2019;19(1):1258. doi:10.1186/s12885-019-6484-5

270. Wang XD, Lu J, Lin YS, Gao C, Qi F. Functional role of long non-coding RNA CASC19/miR-140-5p/CEMIP axis in colorectal cancer progression in vitro. World J Gastroenterol. 2019;25(14):1697–1714. doi:10.3748/wjg.v25.i14.1697

271. Gao Y, Nan X, Shi X, Mu X, Liu B, Zhu H, Yao B, Liu X, Yang T, Hu Y, et al. SREBP1 promotes the invasion of colorectal cancer accompanied upregulation of MMP7 expression and NF-κB pathway activation. BMC Cancer. 2019;19(1):685. doi:10.1186/s12885-019-5904-x

272. Liu L, Huang Y, Li Y, Wang Q, Hao Y, Liu L, Yao X, Yao X, Wei Y, Sun X, et al. FJX1 as a candidate diagnostic and prognostic serum biomarker for colorectal cancer. Clin Transl Oncol. 2022;10.1007/s12094-022-02852-5. doi:10.1007/s12094-022-02852-5

273. Abdulla M, Traiki TB, Vaali-Mohammed MA, El-Wetidy MS, Alhassan N, Al-Khayal K, Zubaidi A, Al-Obeed O, Ahmad R. Targeting MUCL1 protein inhibits cell proliferation and EMT by deregulating β□catenin and increases irinotecan sensitivity in colorectal cancer. Int J Oncol. 2022;60(3):22. doi:10.3892/ijo.2022.5312

274. Betge J, Schneider NI, Harbaum L, Pollheimer MJ, Lindtner RA, Kornprat P, Ebert MP, Langner C. MUC1, MUC2, MUC5AC, and MUC6 in colorectal cancer: expression profiles and clinical significance. Virchows Arch. 2016;469(3):255–265. doi:10.1007/s00428-016-1970-5

275. Qu HL, Hasen GW, Hou YY, Zhang CX. THBS2 promotes cell migration and invasion in colorectal cancer via modulating Wnt/β-catenin signaling pathway. Kaohsiung J Med Sci. 2022;38(5):469–478. doi:10.1002/kjm2.12528

276. Chen S, Su T, Zhang Y, Lee A, He J, Ge Q, Wang L, Si J, Zhuo W, Wang L.Fusobacterium nucleatum promotes colorectal cancer metastasis by modulating KRT7-AS/KRT7. Gut Microbes. 2020;11(3):511–525. doi:10.1080/19490976.2019.1695494

277. Taddei A, Castiglione F, Ringressi MN, Niccolai E, Tofani L, Boni L, Bechi P, Amedei A. The Trend of CEACAM3 Blood Expression as Number Index of the CTCs in the Colorectal Cancer Perioperative Course. Mediators Inflamm. 2015;2015:931784. doi:10.1155/2015/931784

278. Maruyama R, Nagaoka Y, Ishikawa A, Akabane S, Fujiki Y, Taniyama D, Sentani K, Oue N. Overexpression of aldolase, fructose-bisphosphate C and its association with spheroid formation in colorectal cancer. Pathol Int. 2022;72(3):176–186. doi:10.1111/pin.13200

279. Wu Y, Xu Y. Bioinformatics for The Prognostic Value and Function of Cubilin (CUBN) in Colorectal Cancer. Med Sci Monit. 2020;26:e922447. doi:10.12659/MSM.922447

280. Cheng YW, Pincas H, Huang J, Zachariah E, Zeng Z, Notterman DA, Paty P, Barany F.High incidence of LRAT promoter hypermethylation in colorectal cancer correlates with tumor stage. Med Oncol. 2014;31(11):254. doi:10.1007/s12032-014-0254-7

281. Módis K, Coletta C, Asimakopoulou A, Szczesny B, Chao C, Papapetropoulos A, Hellmich MR, Szabo C. Effect of S-adenosyl-L-methionine (SAM), an allosteric activator of cystathionine-β-synthase (CBS) on colorectal cancer cell proliferation and bioenergetics in vitro. Nitric Oxide. 2014;41:146–156. doi:10.1016/j.niox.2014.03.001

282. Yang C, Zhou Q, Li M, Tong X, Sun J, Qing Y, Sun L, Yang X, Hu X, Jiang J, et al. Upregulation of CYP2S1 by oxaliplatin is associated with p53 status in colorectal cancer cell lines. Sci Rep. 2016;6:33078. doi:10.1038/srep33078

283. Liu K, Li YC, Chen Y, Shi XB, Xing ZH, He ZJ, Wang ST, Liu WJ, Zhang PW, Yu ZZ, et al. AZ32 Reverses ABCG2-Mediated Multidrug Resistance in Colorectal Cancer. Front Oncol. 2021;11:680663. doi:10.3389/fonc.2021.680663

284. Gervasini G, García-Martín E, Ladero JM, Pizarro R, Sastre J, Martínez C, García M, Diaz-Rubio M, Agúndez JA. Genetic variability in CYP3A4 and CYP3A5 in primary liver, gastric and colorectal cancer patients. BMC Cancer. 2007;7:118. doi:10.1186/1471-2407-7-118

285. Sirniö P, Väyrynen JP, Klintrup K, Mäkelä J, Mäkinen MJ, Karttunen TJ, Tuomisto A. Decreased serum apolipoprotein A1 levels are associated with poor survival and systemic inflammatory response in colorectal cancer. Sci Rep. 2017;7(1):5374. doi:10.1038/s41598-017-05415-9

286. Zhang T, Yang P, Wei J, Li W, Zhong J, Chen H, Cao J. Overexpression of flavin-containing monooxygenase 5 predicts poor prognosis in patients with colorectal cancer. Oncol Lett. 2018;15(3):3923–3927. doi:10.3892/ol.2018.7724

287. Nomiri S, Hoshyar R, Chamani E, Rezaei Z, Salmani F, Larki P, Tavakoli T, Gholipour F, Tabrizi NJ, Derakhshani A, et al. Prediction and validation of GUCA2B as the hub-gene in colorectal cancer based on co-expression network analysis: In-silico and in-vivo study. Biomed Pharmacother. 2022;147:112691. doi:10.1016/j.biopha.2022.112691

288. Li Q, Wei P, Wu J, Zhang M, Li G, Li Y, Xu Y, Li X, Xie D, Cai S, et al. The FOXC1/FBP1 signaling axis promotes colorectal cancer proliferation by enhancing the Warburg effect. Oncogene. 2019;38(4):483–496. doi:10.1038/s41388-018-0469-8

289. Zhao T, Li Y, Shen K, Wang Q, Zhang J. Knockdown of OLR1 weakens glycolytic metabolism to repress colon cancer cell proliferation and chemoresistance by downregulating SULT2B1 via c-MYC. Cell Death Dis. 2021;13(1):4. doi:10.1038/s41419-021-04174-w

290. Yao Y, Wang X, Zhou D, Li H, Qian H, Zhang J, Jiang L, Wang B, Lin Q, Zhu X. Loss of AKR1B10 promotes colorectal cancer cells proliferation and migration via regulating FGF1-dependent pathway. Aging (Albany NY). 2020;12(13):13059–13075. doi:10.18632/aging.103393

291. Hezova R, Bienertova-Vasku J, Sachlova M, Brezkova V, Vasku A, Svoboda M, Radová L, Kiss I, Vyzula R, Slaby O.Common polymorphisms in GSTM1, GSTT1, GSTP1, GSTA1 and susceptibility to colorectal cancer in the Central European population. Eur J Med Res. 2012;17(1):17. doi:10.1186/2047-783X-17-17

292. Kong C, Yan X, Zhu Y, Zhu H, Luo Y, Liu P, Ferrandon S, Kalady MF, Gao R, He J, et al. Fusobacterium Nucleatum Promotes the Development of Colorectal Cancer by Activating a Cytochrome P450/Epoxyoctadecenoic Acid Axis via TLR4/Keap1/NRF2 Signaling. Cancer Res. 2021;81(17):4485–4498. doi:10.1158/0008-5472.CAN-21-0453

293. An J, Ha EM. Extracellular vesicles derived from Lactobacillus plantarum restore chemosensitivity through the PDK2-mediated glucose metabolic pathway in 5-FU-resistant colorectal cancer cells. J Microbiol. 2022;60(7):735–745. doi:10.1007/s12275-022-2201-1

294. Tachibana K, Saito M, Imai JI, Ito E, Yanagisawa Y, Honma R, Saito K, Ando J, Momma T, Ohki S, et al. Clinicopathological examination of dipeptidase 1 expression in colorectal cancer. Biomed Rep. 2017;6(4):423–428. doi:10.3892/br.2017.870

295. Yang DD, Chen ZH, Wang DS, Yu HE, Lu JH, Xu RH, Zeng ZL. Prognostic value of the serum apolipoprotein B to apolipoprotein A-I ratio in metastatic colorectal cancer patients. J Cancer. 2020;11(5):1063–1074. doi:10.7150/jca.35659

296. Ishimine M, Lee HC, Nakaoka H, Orita H, Kobayashi T, Mizuguchi K, Endo M, Inoue I, Sato K, Yokomizo T. The Relationship between TP53 Gene Status and Carboxylesterase 2 Expression in Human Colorectal Cancer. Dis Markers. 2018;2018:5280736. doi:10.1155/2018/5280736

297. Erichsen HC, Peters U, Eck P, Welch R, Schoen RE, Yeager M, Levine M, Hayes RB, Chanock S. Genetic variation in sodium-dependent vitamin C transporters SLC23A1 and SLC23A2 and risk of advanced colorectal adenoma. Nutr Cancer. 2008;60(5):652–659. doi:10.1080/01635580802033110

298. Liu X, Cheng D, Kuang Q, Liu G, Xu W. Association of UGT1A1*28 polymorphisms with irinotecan-induced toxicities in colorectal cancer: a meta-analysis in Caucasians. Pharmacogenomics J. 2014;14(2):120–129. doi:10.1038/tpj.2013.10

299. Osawa K, Nakarai C, Akiyama M, Hashimoto R, Tsutou A, Takahashi J, Takaoka Y, Kawamura S, Shimada E, Tanaka K, et al. Association between polymorphisms in UDP-glucuronosyltransferase 1A6 and 1A7 and colorectal cancer risk. Asian Pac J Cancer Prev. 2012;13(5):2311–2314. doi:10.7314/apjcp.2012.13.5.2311

300. Zhang H, Du Y, Wang Z, Lou R, Wu J, Feng J. Integrated Analysis of Oncogenic Networks in Colorectal Cancer Identifies GUCA2A as a Molecular Marker. Biochem Res Int. 2019;2019:6469420. doi:10.1155/2019/6469420

301. Shen Z, Li Z, Liu Y, Li Y, Feng X, Zhan Y, Lin M, Fang C, Fang Y, Deng H. GLUT5-KHK axis-mediated fructose metabolism drives proliferation and chemotherapy resistance of colorectal cancer. Cancer Lett. 2022;534:215617. doi:10.1016/j.canlet.2022.215617

302. Goetz MP, Suman VJ, Hoskin TL, Gnant M, Filipits M, Safgren SL, Kuffel M, Jakesz R, Rudas M, Greil R, et al. CYP2D6 metabolism and patient outcome in the Austrian Breast and Colorectal Cancer Study Group trial (ABCSG) 8. Clin Cancer Res. 2013;19(2):500–507. doi:10.1158/1078-0432.CCR-12-2153

303. Buck E, Sprick M, Gaida MM, Grüllich C, Weber TF, Herpel E, Bruckner T, Koschny R. Tumor response to irinotecan is associated with CYP3A5 expression in colorectal cancer. Oncol Lett. 2019;17(4):3890–3898. doi:10.3892/ol.2019.10043

304. Zhou J, Xie Z, Cui P, Su Q, Zhang Y, Luo L, Li Z, Ye L, Liang H, Huang J. SLC1A1, SLC16A9, and CNTN3 Are Potential Biomarkers for the Occurrence of Colorectal Cancer. Biomed Res Int. 2020;2020:1204605. doi:10.1155/2020/1204605

305. Ladero JM, Agúndez JA, Martínez C, Amo G, Ayuso P, García-Martín E. Analysis of the Functional Polymorphism in the Cytochrome P450 CYP2C8 Gene rs11572080 with Regard to Colorectal Cancer Risk. Front Genet. 2012;3:278. doi:10.3389/fgene.2012.00278

306. Kwon RJ, Park EJ, Lee SY, Lee Y, Hwang C, Kim C, Cho YH. Expression and prognostic significance of Niemann-Pick C1-Like 1 in colorectal cancer: a retrospective cohort study. Lipids Health Dis. 2021;20(1):104. doi:10.1186/s12944-021-01539-0

307. Yu B, Liu X, Cao X, Zhang M, Chang H. Study of the expression and function of ACY1 in patients with colorectal cancer. Oncol Lett. 2017;13(4):2459–2464. doi:10.3892/ol.2017.5702

308. Artemaki PI, Papatsirou M, Boti MA, Adamopoulos PG, Christodoulou S, Vassilacopoulou D, Scorilas A, Kontos CK. Revised Exon Structure of l-DOPA Decarboxylase (DDC) Reveals Novel Splice Variants Associated with Colorectal Cancer Progression. Int J Mol Sci. 2020;21(22):8568.. doi:10.3390/ijms21228568

309. Pucci M, Malagolini N, Dall’Olio F. Glycosyltransferase B4GALNT2 as a Predictor of Good Prognosis in Colon Cancer: Lessons from Databases. Int J Mol Sci. 2021;22(9):4331. doi:10.3390/ijms22094331

310. Yang ZZ, Li L, Xu MC, Ju HX, Hao M, Gu JK, Jim Wang ZJ, Jiang HD, Yu LS, Zeng S. Brain-derived neurotrophic factor involved epigenetic repression of UGT2B7 in colorectal carcinoma: A mechanism to alter morphine glucuronidation in tumor. Oncotarget. 2017;8(17):29138–29150. doi:10.18632/oncotarget.16251

311. Zou K, Hu Y, Li M, Wang H, Zhang Y, Huang L, Xie Y, Li S, Dai X, et al. Potential Role of HMGCS2 in Tumor Angiogenesis in Colorectal Cancer and Its Potential Use as a Diagnostic Marker. Can J Gastroenterol Hepatol. 2019;2019:8348967. doi:10.1155/2019/8348967

312. Yamaguchi N, Weinberg EM, Nguyen A, Liberti MV, Goodarzi H, Janjigian YY, Paty PB, Saltz LB, Kingham TP, Loo JM, et al. PCK1 and DHODH drive colorectal cancer liver metastatic colonization and hypoxic growth by promoting nucleotide synthesis. Elife. 2019;8:e52135. doi:10.7554/eLife.52135

313. Özhan G, Mutur M, Ercan G, Alpertunga B. Genetic variations in the xenobiotic-metabolizing enzymes CYP1A1, CYP1A2, CYP2C9, CYP2C19 and susceptibility to colorectal cancer among Turkish people. Genet Test Mol Biomarkers. 2014;18(4):223–228. doi:10.1089/gtmb.2013.0358

314. Hu X, Yuan P, Yan J, Feng F, Li X, Liu W, Yang Y. Gene Polymorphisms of ADIPOQ +45T>G, UCP2 -866G>A, and FABP2 Ala54Thr on the Risk of Colorectal Cancer: A Matched Case-Control Study. PLoS One. 2013;8(6):e67275. doi:10.1371/journal.pone.0067275

315. Guo W, Zhang C, Feng P, Li M, Wang X, Xia Y, Chen D, Li J. M6A methylation of DEGS2, a key ceramide-synthesizing enzyme, is involved in colorectal cancer progression through ceramide synthesis. Oncogene. 2021;40(40):5913–5924. doi:10.1038/s41388-021-01987-z

316. Wang R, Löhr CV, Fischer K, Dashwood WM, Greenwood JA, Ho E, Williams DE, Ashktorab H, Dashwood MR, Dashwood RH. Epigenetic inactivation of endothelin-2 and endothelin-3 in colon cancer. Int J Cancer. 2013;132(5):1004–1012. doi:10.1002/ijc.27762

317. Lam KK, Sethi R, Tan G, Tomar S, Lo M, Loi C, Tang CL, Tan E, Lai PS, Cheah PY. The orphan nuclear receptor NR0B2 could be a novel susceptibility locus associated with microsatellite-stable, APC mutation-negative early-onset colorectal carcinomas with metabolic manifestation. Genes Chromosomes Cancer. 2021;60(2):61–72. doi:10.1002/gcc.22904

318. Li Q, Li Y, Xu J, Wang S, Xu Y, Li X, Cai S. Aldolase B Overexpression is Associated with Poor Prognosis and Promotes Tumor Progression by Epithelial-Mesenchymal Transition in Colorectal Adenocarcinoma. Cell Physiol Biochem. 2017;42(1):397–406. doi:10.1159/000477484

319. Hasenoehrl C, Feuersinger D, Sturm EM, Bärnthaler T, Heitzer E, Graf R, Grill M, Pichler M, Beck S, Butcher L, et al. G protein-coupled receptor GPR55 promotes colorectal cancer and has opposing effects to cannabinoid receptor 1. Int J Cancer. 2018;142(1):121–132. doi:10.1002/ijc.31030

320. Carlini LE, Meropol NJ, Bever J, Andria ML, Hill T, Gold P, Rogatko A, Wang H, Blanchard RL. UGT1A7 and UGT1A9 polymorphisms predict response and toxicity in colorectal cancer patients treated with capecitabine/irinotecan. Clin Cancer Res. 2005;11(3):1226–1236.

321. Treenert A, Areepium N, Tanasanvimon S. Effects of ABCC2 and SLCO1B1 Polymorphisms on Treatment Responses in Thai Metastatic Colorectal Cancer Patients Treated with Irinotecan-Based Chemotherapy. Asian Pac J Cancer Prev. 2018;19(10):2757–2764. doi:10.22034/APJCP.2018.19.10.2757

322. Zhang F, Ye J, Guo W, Zhang F, Wang L, Han A. TYMS-TM4SF4 axis promotes the progression of colorectal cancer by EMT and upregulating stem cell marker. Am J Cancer Res. 2022;12(3):1009–1026.

323. Zhang H, Yang W, Yan J, Zhou K, Wan B, Shi P, Chen Y, He S, Li D. Loss of profilin 2 contributes to enhanced epithelial-mesenchymal transition and metastasis of colorectal cancer. Int J Oncol. 2018;53(3):1118–1128. doi:10.3892/ijo.2018.4475

324. Yang YC, Chien MH, Lai TC, Su CY, Jan YH, Hsiao M, Chen CL. Monoamine Oxidase B Expression Correlates with a Poor Prognosis in Colorectal Cancer Patients and Is Significantly Associated with Epithelial-to-Mesenchymal Transition-Related Gene Signatures. Int J Mol Sci. 2020;21(8):2813.doi:10.3390/ijms21082813

325. Murakami Y, Konishi H, Fujiya M, Takahashi K, Ando K, Ueno N, Kashima S, Moriichi K, Tanabe H, Okumura T. Testis-specific hnRNP is expressed in colorectal cancer cells and accelerates cell growth mediating ZDHHC11 mRNA stabilization. Cancer Med. 2022;10.1002/cam4.4738. doi:10.1002/cam4.4738

326. Han Y, Wang X, Mao E, Shen B, Huang L. lncRNA FLVCR1□AS1 drives colorectal cancer progression via modulation of the miR□381/RAP2A axis. Mol Med Rep. 2021;23(2):139. doi:10.3892/mmr.2020.11778

327. Jiang G, Wang P, Wang W, Li W, Dai L, Chen K. Annexin A13 promotes tumor cell invasion in vitro and is associated with metastasis in human colorectal cancer. Oncotarget. 2017;8(13):21663–21673. doi:10.18632/oncotarget.15523

328. Zeng C, Chen Y. HTR1D, TIMP1, SERPINE1, MMP3 and CNR2 affect the survival of patients with colon adenocarcinoma. Oncol Lett. 2019;18(3):2448–2454. doi:10.3892/ol.2019.10545

329. Yuan ZT, Shi XJ, Yuan YX, Qiu YY, Zou Y, Liu C, Yu H, He X, Xu K, Yin PH. Bufalin reverses ABCB1-mediated drug resistance in colorectal cancer. Oncotarget. 2017;8(29):48012–48026. doi:10.18632/oncotarget.18225

330. Lee JH, Jo YS, Kim MS, Yoo NJ, Lee SH. Inactivating frameshift mutation of putative tumor suppressor genes PLA2R1 and SRPK1 in gastric and colorectal cancers. Cancer Genet. 2017;210:34–35. doi:10.1016/j.cancergen.2016.11.005

331. Liu C, Wang K, Zhang M, Hu X, Hu T, Liu Y, Hu Q, Wu S, Yue J. High expression of ACE2 and TMPRSS2 and clinical characteristics of COVID-19 in colorectal cancer patients. NPJ Precis Oncol. 2021;5(1):1. doi:10.1038/s41698-020-00139-y

332. Sierko E, Wojtukiewicz MZ, Zimnoch L, Tokajuk P, Ostrowska-Cichocka K, Kisiel W. Co-localization of Protein Z, Protein Z-Dependent protease inhibitor and coagulation factor X in human colon cancer tissue: implications for coagulation regulation on tumor cells. Thromb Res. 2012;129(4):e112–e118. doi:10.1016/j.thromres.2011.10.027

333. Ouyang X, Zhang G, Pan H, Huang J. Susceptibility and severity of cancer-related fatigue in colorectal cancer patients is associated with SLC6A4 gene single nucleotide polymorphism rs25531 A>G genotype. Eur J Oncol Nurs. 2018;33:97–101. doi:10.1016/j.ejon.2018.02.003

334. Zhang Y, Zhao X, Deng L, Li X, Wang G, Li Y, Chen M. High expression of FABP4 and FABP6 in patients with colorectal cancer. World J Surg Oncol. 2019;17(1):171. doi:10.1186/s12957-019-1714-5

335. Zou D, Lou J, Ke J, Mei S, Li J, Gong Y, Yang Y, Zhu Y, Tian J, Chang J, et al. Integrative expression quantitative trait locus-based analysis of colorectal cancer identified a functional polymorphism regulating SLC22A5 expression. Eur J Cancer. 2018;93:1–9. doi:10.1016/j.ejca.2018.01.065

336. Ng L, Foo DC, Wong CK, Man AT, Lo OS, Law WL. Repurposing DPP-4 Inhibitors for Colorectal Cancer: A Retrospective and Single Center Study. Cancers (Basel). 2021;13(14):3588. doi:10.3390/cancers13143588

337. Zhao H, Sheng D, Qian Z, Ye S, Chen J, Tang Z. Identifying GNG4 might play an important role in colorectal cancer TMB. Cancer Biomark. 2021;32(4):435–450. doi:10.3233/CBM-203009

338. Mu L, Wang Y, Hu Y, Shi C, Alman BA, Zhang C, She J. The Role of TMIGD1 as a Tumor Suppressor in Colorectal Cancer. Genet Test Mol Biomarkers. 2022;26(4):174–183. doi:10.1089/gtmb.2021.0169

339. Cheung KS, Chan EW, Seto WK, Wong ICK, Leung WK. ACE (Angiotensin-Converting Enzyme) Inhibitors/Angiotensin Receptor Blockers Are Associated With Lower Colorectal Cancer Risk: A Territory-Wide Study With Propensity Score Analysis. Hypertension. 2020;76(3):968–975. doi:10.1161/HYPERTENSIONAHA.120.15317

340. Iyer DN, Foo DC, Lo OS, Wan TM, Li X, Sin RW, Pang RW, Law WL, Ng L. MiR-509-3p is oncogenic, targets the tumor suppressor PHLPP2, and functions as a novel tumor adjacent normal tissue based prognostic biomarker in colorectal cancer. BMC Cancer. 2022;22(1):351. doi:10.1186/s12885-021-09075-x

341. Davis MI, Pragani R, Fox JT, Shen M, Parmar K, Gaudiano EF, Liu L, Tanega C, McGee L, Hall MD, et al. Small Molecule Inhibition of the Ubiquitin-specific Protease USP2 Accelerates cyclin D1 Degradation and Leads to Cell Cycle Arrest in Colorectal Cancer and Mantle Cell Lymphoma Models. J Biol Chem. 2016;291(47):24628–24640. doi:10.1074/jbc.M116.738567

342. Hauptman N, Jevšinek Skok D, Spasovska E, Boštjančič E, Glavač D. Genes CEP55, FOXD3, FOXF2, GNAO1, GRIA4, and KCNA5 as potential diagnostic biomarkers in colorectal cancer. BMC Med Genomics. 2019;12(1):54. doi:10.1186/s12920-019-0501-z

343. Guo S, Sun Y. OTOP2, Inversely Modulated by miR-3148, Inhibits CRC Cell Migration, Proliferation and Epithelial-Mesenchymal Transition: Evidence from Bioinformatics Data Mining and Experimental Verification. Cancer Manag Res. 2022;14:1371–1384. doi:10.2147/CMAR.S345299

344. Lin M, Fang Y, Li Z, Li Y, Feng X, Zhan Y, Xie Y, Liu Y, Liu Z, Li G, et al. S100P contributes to promoter demethylation and transcriptional activation of SLC2A5 to promote metastasis in colorectal cancer. Br J Cancer. 2021;125(5):734–747. doi:10.1038/s41416-021-01306-z

345. Lee YE, He HL, Shiue YL, Lee SW, Lin LC, Wu TF, Chang IW, Lee HH, Li CF. The prognostic impact of lipid biosynthesis-associated markers, HSD17B2 and HMGCS2, in rectal cancer treated with neoadjuvant concurrent chemoradiotherapy. Tumour Biol. 2015;36(10):7675–7683. doi:10.1007/s13277-015-3503-2

346. Liu F, Ou W, Tang W, Huang Z, Zhu Z, Ding W, Fu J, Zhu Y, Liu C, Xu W, et al. Increased AOC1 Expression Promotes Cancer Progression in Colorectal Cancer. Front Oncol. 2021;11:657210. doi:10.3389/fonc.2021.657210

347. Hostettler L, Zlobec I, Terracciano L, Lugli A. ABCG5-positivity in tumor buds is an indicator of poor prognosis in node-negative colorectal cancer patients. World J Gastroenterol. 2010;16(6):732–739. doi:10.3748/wjg.v16.i6.732

348. Chuang HY, Jiang JK, Yang MH, Wang HW, Li MC, Tsai CY, Jhang YY, Huang JC. Aminopeptidase A initiates tumorigenesis and enhances tumor cell stemness via TWIST1 upregulation in colorectal cancer. Oncotarget. 2017;8(13):21266–21280. doi:10.18632/oncotarget.15072

349. Lin Q, Li J, Zhu D, Niu Z, Pan X, Xu P, Ji M, Wei Y, Xu J. Aberrant Scinderin Expression Correlates With Liver Metastasis and Poor Prognosis in Colorectal Cancer. Front Pharmacol. 2019;10:1183. doi:10.3389/fphar.2019.01183

350. Ma WR, Xu P, Liu ZJ, Zhou J, Gu LK, Zhang J, Deng DJ. Impact of GFRA1 gene reactivation by DNA demethylation on prognosis of patients with metastatic colon cancer. World J Gastroenterol. 2020;26(2):184–198. doi:10.3748/wjg.v26.i2.184

351. Ke J, Tian J, Mei S, Ying P, Yang N, Wang X, Zou D, Peng X, Yang Y, Zhu Y, et al. Genetic Predisposition to Colon and Rectal Adenocarcinoma Is Mediated by a Super-enhancer Polymorphism Coactivating CD9 and PLEKHG6. Cancer Epidemiol Biomarkers Prev. 2020;29(4):850–859. doi:10.1158/1055-9965.EPI-19-1116

352. Park BK, Park JY, Kim TH, Kim D, Wu G, Gautam A, Maharjan S, Lee SI, Lee Y, Kwon HJ, et al. Production of an anti-TM4SF5 monoclonal antibody and its application in the detection of TM4SF5 as a possible marker of a poor prognosis in colorectal cancer. Int J Oncol. 2018;53(1):275–285. doi:10.3892/ijo.2018.4385

353. Xiang T, Yu F, Fei R, Qian J, Chen W. CHRNA7 inhibits cell invasion and metastasis of LoVo human colorectal cancer cells through PI3K/Akt signaling. Oncol Rep. 2016;35(2):999–1005. doi:10.3892/or.2015.4462

354. Cheng B, Rong A, Zhou Q, Li W. CLDN8 promotes colorectal cancer cell proliferation, migration, and invasion by activating MAPK/ERK signaling. Cancer Manag Res. 2019;11:3741–3751. doi:10.2147/CMAR.S189558

355. Lise M, Loda M, Fiorentino M, Mercurio AM, Summerhayes IC, Lavin PT, Jessup JM. Association between sucrase-isomaltase and p53 expression in colorectal cancer. Ann Surg Oncol. 1997;4(2):176–183. doi:10.1007/BF02303802

356. Li Y, Liu S, Gao Y, Ma H, Zhan S, Yang Y, Xin Y, Xuan S. Association of TM6SF2 rs58542926 gene polymorphism with the risk of non-alcoholic fatty liver disease and colorectal adenoma in Chinese Han population. BMC Biochem. 2019;20(1):3. doi:10.1186/s12858-019-0106-3

357. Takashima Y, Shimada T, Yokozawa T. Clinical benefit of measuring both haemoglobin and transferrin concentrations in faeces: demonstration during a large-scale colorectal cancer screening trial in Japan. Diagnosis (Berl). 2015;2(1):53–59. doi:10.1515/dx-2014-0052

358. Zhuang Y, Liu PF, Zhan Y, Kong DL, Tian F, Zhao P. RING finger protein 128 (RNF128) regulates malignant biological behaviors of colorectal cancer cells via PI3K/AKT signaling pathway. Cell Biol Int. 2022;10.1002/cbin.11835. doi:10.1002/cbin.11835

359. Wang X, Chen J, Wang J, Yu F, Zhao S, Zhang Y, Tang H, Peng Z. Correction to: Metalloproteases meprin-□ (MEP1A) is a prognostic biomarker and promotes proliferation and invasion of colorectal cancer. BMC Cancer. 2018;18(1):70. doi:10.1186/s12885-017-3767-6

360. Gençdal G, Salman E, Özütemiz Ö, Akarca US. Association of LCT-13910 C/T Polymorphism and Colorectal Cancer. Ann Coloproctol. 2017;33(5):169–172. doi:10.3393/ac.2017.33.5.169

361. Yang Y, Wu J, Yu X, Wu Q, Cao H, Dai X, Chen H. SLC34A2 promotes cancer proliferation and cell cycle progression by targeting TMPRSS3 in colorectal cancer. Pathol Res Pract. 2022;229:153706. doi:10.1016/j.prp.2021.153706

362. Li A, Lu D, Zhang Y, Li J, Fang Y, Li F, Sun J.Critical role of aquaporin-3 in epidermal growth factor-induced migration of colorectal carcinoma cells and its clinical significance. Oncol Rep. 2013;29(2):535–540. doi:10.3892/or.2012.2144

363. Shin J, Carr A, Corner GA, Tögel L, Dávalos-Salas M, Tran H, Chueh AC, Al-Obaidi S, Chionh F, Ahmed N, et al. The intestinal epithelial cell differentiation marker intestinal alkaline phosphatase (ALPi) is selectively induced by histone deacetylase inhibitors (HDACi) in colon cancer cells in a Kruppel-like factor 5 (KLF5)-dependent manner. J Biol Chem. 2015;290(25):15392. doi:10.1074/jbc.A114.557546

364. Tang SH, Hsiao CW, Chen WL, Wu LW, Chang JB, Yang BH. Hypermethylation of SHISA3 DNA as a blood-based biomarker for colorectal cancer. Chin J Physiol. 2021;64(1):51–56. doi:10.4103/CJP.CJP_89_20

365. Modarai SR, Opdenaker LM, Viswanathan V, Fields JZ, Boman BM. Somatostatin signaling via SSTR1 contributes to the quiescence of colon cancer stem cells. BMC Cancer. 2016;16(1):941. doi:10.1186/s12885-016-2969-7

366. Ding Y, Feng W, Ge JK, Dai L, Liu TT, Hua XY, Lu X, Ju SQ, Yu J. Serum level of long noncoding RNA B3GALT5-AS1 as a diagnostic biomarker of colorectal cancer. Future Oncol. 2020;16(13):827–835. doi:10.2217/fon-2019-0820

367. Li D, Zhang L, Fu J, Huang H, Sun S, Zhang D, Zhao L, Ucheojor Onwuka J, Zhao Y, Cui B. SCTR hypermethylation is a diagnostic biomarker in colorectal cancer. Cancer Sci. 2020;111(12):4558–4566. doi:10.1111/cas.14661

368. Fromme JE, Schmitz K, Wachter A, Grzelinski M, Zielinski D, Koppel C, Conradi LC, Homayounfar K, Hugo T, Hugo S, et al. FGFR3 mRNA overexpression defines a subset of oligometastatic colorectal cancers with worse prognosis. Oncotarget. 2018;9(63):32204–32218. doi:10.18632/oncotarget.25941

369. Maurya NS, Kushwaha S, Chawade A, Mani A. Transcriptome profiling by combined machine learning and statistical R analysis identifies TMEM236 as a potential novel diagnostic biomarker for colorectal cancer. Sci Rep. 2021;11(1):14304. doi:10.1038/s41598-021-92692-0

370. Cai BH, Wu PH, Chou CK, Huang HC, Chao CC, Chung HY, Lee HY, Chen JY, Kannagi R. Synergistic activation of the NEU4 promoter by p73 and AP2 in colon cancer cells. Sci Rep. 2019;9(1):950. doi:10.1038/s41598-018-37521-7

371. Lin Y, Chen Z, Zheng Y, Liu Y, Gao J, Lin S, Chen S. MiR-506 Targets UHRF1 to Inhibit Colorectal Cancer Proliferation and Invasion via the KISS1/PI3K/NF-κB Signaling Axis. Front Cell Dev Biol. 2019;7:266. doi:10.3389/fcell.2019.00266

372. He XS, Ye WL, Zhang YJ, Yang XQ, Liu F, Wang JR, Ding XL, Yang Y, Zhang RN, Zhao YY, et al. Oncogenic potential of BEST4 in colorectal cancer via activation of PI3K/Akt signaling. Oncogene. 2022;41(8):1166–1177. doi:10.1038/s41388-021-02160-2

373. Senapati S, Ho SB, Sharma P, Das S, Chakraborty S, Kaur S, Niehans G, Batra SK. Expression of intestinal MUC17 membrane-bound mucin in inflammatory and neoplastic diseases of the colon. J Clin Pathol. 2010;63(8):702–707. doi:10.1136/jcp.2010.078717

374. Hou L, Liu P, Zhu T. Long noncoding RNA SLC30A10 promotes colorectal tumor proliferation and migration via miR-21c/APC axis. Eur Rev Med Pharmacol Sci. 2020;24(12):6682–6691. doi:10.26355/eurrev_202006_21655

375. Meng H, Li W, Boardman LA, Wang L. Loss of ZG16 is associated with molecular and clinicopathological phenotypes of colorectal cancer. BMC Cancer. 2018;18(1):433. doi:10.1186/s12885-018-4337-2

376. Mudd TW Jr, Lu C, Klement JD, Liu K. MS4A1 expression and function in T cells in the colorectal cancer tumor microenvironment. Cell Immunol. 2021;360:104260. doi:10.1016/j.cellimm.2020.104260

377. Mu Q, Luo G, Wei J, Zheng L, Wang H, Yu M, Xu N. Apolipoprotein M promotes growth and inhibits apoptosis of colorectal cancer cells through upregulation of ribosomal protein S27a. EXCLI J. 2021;20:145–159. doi:10.17179/excli2020-2867

378. Maekawa K, Hamaguchi T, Saito Y, Tatewaki N, Kurose K, Kaniwa N, Eguchi Nakajima T, Kato K, Yamada Y, Shimada Y, et al. Genetic variation and haplotype structures of the glutathione S-transferase genes GSTA1 and GSTA2 in Japanese colorectal cancer patients. Drug Metab Pharmacokinet. 2011;26(6):646–658. doi:10.2133/dmpk.DMPK-11-SC-050

379. Mostafa GA, Al-Ayadhi LY. The possible link between elevated serum levels of epithelial cell-derived neutrophil-activating peptide-78 (ENA-78/CXCL5) and autoimmunity in autistic children. Behav Brain Funct. 2015;11:11. doi:10.1186/s12993-015-0056-x

380. Zheng R, Chen S, Chen S. Correlation between myeloid-derived suppressor cells and S100A8/A9 in tumor and autoimmune diseases. Int Immunopharmacol. 2015;29(2):919–925. doi:10.1016/j.intimp.2015.10.01

381. Ding L, Hanawa H, Ota Y, Hasegawa G, Hao K, Asami F, Watanabe R, Yoshida T, Toba K, Yoshida K, et al. Lipocalin-2/neutrophil gelatinase-B associated lipocalin is strongly induced in hearts of rats with autoimmune myocarditis and in human myocarditis. Circ J. 2010;74(3):523–530. doi:10.1253/circj.cj-09-0485

382. Bachmaier K, Toya S, Malik AB. Therapeutic administration of the chemokine CXCL1/KC abrogates autoimmune inflammatory heart disease. PLoS One. 2014;9(2):e89647. doi:10.1371/journal.pone.0089647

383. Björk P, Björk A, Vogl T, Stenström M, Liberg D, Olsson A, Roth J, Ivars F, Leanderson T. Identification of human S100A9 as a novel target for treatment of autoimmune disease via binding to quinoline-3-carboxamides. PLoS Biol. 2009;7(4):e97. doi:10.1371/journal.pbio.1000097

384. Antonelli A, Ferrari SM, Frascerra S, Di Domenicantonio A, Nicolini A, Ferrari P, Ferrannini E, Fallahi P. Increase of circulating CXCL9 and CXCL11 associated with euthyroid or subclinically hypothyroid autoimmune thyroiditis. J Clin Endocrinol Metab. 2011;96(6):1859–1863. doi:10.1210/jc.2010-2905

385. Zohar Y, Wildbaum G, Novak R, Salzman AL, Thelen M, Alon R, Barsheshet Y, Karp CL, Karin N. CXCL11-dependent induction of FOXP3-negative regulatory T cells suppresses autoimmune encephalomyelitis. J Clin Invest. 2018;128(3):1200–1201. doi:10.1172/JCI120358

386. Antonelli A, Ferrari SM, Giuggioli D, Ferrannini E, Ferri C, Fallahi P. Chemokine (C-X-C motif) ligand (CXCL)10 in autoimmune diseases. Autoimmun Rev. 2014;13(3):272–280. doi:10.1016/j.autrev.2013.10.010

387. Chou J, Hsu JT, Bainter W, Al-Attiyah R, Al-Herz W, Geha RS. A novel mutation in NCF2 associated with autoimmune disease and a solitary late-onset infection. Clin Immunol. 2015;161(2):128–130. doi:10.1016/j.clim.2015.08.003

388. Archer NS, Nassif NT, O’Brien BA. Genetic variants of SLC11A1 are associated with both autoimmune and infectious diseases: systematic review and meta-analysis. Genes Immun. 2015;16(4):275–283. doi:10.1038/gene.2015.8

389. Lorenz G, Ribeiro A, von Rauchhaupt E, Würf V, Schmaderer C, Cohen CD, Vohra T, Anders HJ, Lindenmeyer M, Lech M. GDF15 Suppresses Lymphoproliferation and Humoral Autoimmunity in a Murine Model of Systemic Lupus Erythematosus. J Innate Immun. 2022;1–17. doi:10.1159/000523991

390. Reddy S, Jia S, Geoffrey R, Lorier R, Suchi M, Broeckel U, Hessner MJ, Verbsky J. An autoinflammatory disease due to homozygous deletion of the IL1RN locus. N Engl J Med. 2009;360(23):2438–2444. doi:10.1056/NEJMoa0809568

391. Wang X, Deckert M, Xuan NT, Nishanth G, Just S, Waisman A, Naumann M, Schlüter D. Astrocytic A20 ameliorates experimental autoimmune encephalomyelitis by inhibiting NF-κB- and STAT1-dependent chemokine production in astrocytes. Acta Neuropathol. 2013;126(5):711–724. doi:10.1007/s00401-013-1183-9

392. Ma X, Xie Z, Qin J, Luo S, Zhou Z. Association of Vitamin D Pathway Gene CYP27B1 and CYP2R1 Polymorphisms with Autoimmune Endocrine Disorders: A Meta-Analysis. J Clin Endocrinol Metab. 2020;105(11):dgaa525. doi:10.1210/clinem/dgaa525

393. Ham DW, Kim SG, Seo SH, Shin JH, Lee SH, Shin EH. Chronic Toxoplasma gondii Infection Alleviates Experimental Autoimmune Encephalomyelitis by the Immune Regulation Inducing Reduction in IL-17A/Th17 Via Upregulation of SOCS3. Neurotherapeutics. 2021;18(1):430–447. doi:10.1007/s13311-020-00957-9

394. Li B, Baylink DJ, Deb C, Zannetti C, Rajaallah F, Xing W, Walter MH, Lau KH, Qin X.1,25-Dihydroxyvitamin D3 suppresses TLR8 expression and TLR8-mediated inflammatory responses in monocytes in vitro and experimental autoimmune encephalomyelitis in vivo. PLoS One. 2013;8(3):e58808. doi:10.1371/journal.pone.0058808

395. Ruiz-Argüelles A, Llorente L. The role of complement regulatory proteins (CD55 and CD59) in the pathogenesis of autoimmune hemocytopenias. Autoimmun Rev. 2007;6(3):155–161. doi:10.1016/j.autrev.2006.09.008

396. Wang J, Wang X, Chen X, Lu S, Kuang Y, Fei J, Wang Z. Gpr97/Adgrg3 ameliorates experimental autoimmune encephalomyelitis by regulating cytokine expression. Acta Biochim Biophys Sin (Shanghai). 2018;50(7):666–675. doi:10.1093/abbs/gmy060

397. Quick ML, Mukherjee S, Rudick CN, Done JD, Schaeffer AJ, Thumbikat P. CCL2 and CCL3 are essential mediators of pelvic pain in experimental autoimmune prostatitis. Am J Physiol Regul Integr Comp Physiol. 2012;303(6):R580–R589. doi:10.1152/ajpregu.00240.2012

398. Dahlqvist J, Fulco CP, Ray JP, Liechti T, de Boer CG, Lieb DJ, Eisenhaure TM, Engreitz JM, Roederer M, Hacohen N. Systematic identification of genomic elements that regulate FCGR2A expression and harbor variants linked with autoimmune disease. Hum Mol Genet. 2022;31(12):1946–1961. doi:10.1093/hmg/ddab372

399. Kim RY, Hoffman AS, Itoh N, Ao Y, Spence R, Sofroniew MV, Voskuhl RR. Astrocyte CCL2 sustains immune cell infiltration in chronic experimental autoimmune encephalomyelitis. J Neuroimmunol. 2014;274(1-2):53–61. doi:10.1016/j.jneuroim.2014.06.009

400. Jia X, Wang B, Yao Q, Li Q, Zhang J. Variations in CD14 Gene Are Associated With Autoimmune Thyroid Diseases in the Chinese Population. Front Endocrinol (Lausanne). 2019;9:811. doi:10.3389/fendo.2018.00811

401. Liu X, Yao DL, Bondy CA, Brenner M, Hudson LD, Zhou J, Webster HD. Astrocytes express insulin-like growth factor-I (IGF-I) and its binding protein, IGFBP-2, during demyelination induced by experimental autoimmune encephalomyelitis. Mol Cell Neurosci. 1994;5(5):418–430. doi:10.1006/mcne.1994.1052

402. Ministrini S, Carbone F. PCSK9 and Inflammation: Their Role in Autoimmune Diseases, with a Focus on Rheumatoid Arthritis and Systemic Lupus Erythematosus. Curr Med Chem. 2022;29(6):970–979. doi:10.2174/0929867328666210810150940

403. Mondanelli G, Carvalho A, Puccetti P, Grohmann U, Volpi C. Reply to Han, et al.: On track for an IDO1-based personalized therapy in autoimmunity. Proc Natl Acad Sci U S A. 2020;117(39):24037–24038. doi:10.1073/pnas.2016277117

404. He Y, Na H, Li Y, Qiu Z, Li W. FoxP3 rs3761548 polymorphism predicts autoimmune disease susceptibility: a meta-analysis. Hum Immunol. 2013;74(12):1665–1671. doi:10.1016/j.humimm.2013.08.270

405. Zhou D, Wang Y, Chen LU, Zhang W, Luan J. Soluble CD163: A Novel Biomarker with Diagnostic and Therapeutic Implications in Autoimmune Diseases. J Rheumatol. 2016;43(4):830. doi:10.3899/jrheum.151317

406. Zhang Y, Han JJ, Liang XY, Zhao L, Zhang F, Rasouli J, Wang ZZ, Zhang GX, Li X. miR-23b Suppresses Leukocyte Migration and Pathogenesis of Experimental Autoimmune Encephalomyelitis by Targeting CCL7. Mol Ther. 2018;26(2):582–592. doi:10.1016/j.ymthe.2017.11.013

407. Shen Y, Yang R, Zhao J, Chen M, Chen S, Ji B, Chen H, Liu D, Li L, Du G. The histone deacetylase inhibitor belinostat ameliorates experimental autoimmune encephalomyelitis in mice by inhibiting TLR2/MyD88 and HDAC3/ NF-κB p65-mediated neuroinflammation. Pharmacol Res. 2022;176:105969. doi:10.1016/j.phrs.2021.105969

408. Chen YF, Zhou D, Metzger T, Gallup M, Jeanne M, Gould DB, Anderson MS, McNamara NA. Spontaneous development of autoimmune uveitis Is CCR2 dependent. Am J Pathol. 2014;184(6):1695–1705. doi:10.1016/j.ajpath.2014.02.024

409. McGovern A, Schoenfelder S, Martin P, Massey J, Duffus K, Plant D, Yarwood A, Pratt AG, Anderson AE, Isaacs JD, et al. Capture Hi-C identifies a novel causal gene, IL20RA, in the pan-autoimmune genetic susceptibility region 6q23. Genome Biol. 2016;17(1):212. doi:10.1186/s13059-016-1078-x

410. Hedegaard Jensen G, Mortensen MB, Klöppel G, Nielsen MFB, Nielsen O, Detlefsen S. Utility of pVHL, maspin, IMP3, S100P and Ki67 in the distinction of autoimmune pancreatitis from pancreatic ductal adenocarcinoma. Pathol Res Pract. 2020;216(5):152925. doi:10.1016/j.prp.2020.152925

411. Cross AK, Haddock G, Surr J, Plumb J, Bunning RA, Buttle DJ, Woodroofe MN. Differential expression of ADAMTS-1, -4, -5 and TIMP-3 in rat spinal cord at different stages of acute experimental autoimmune encephalomyelitis. J Autoimmun. 2006;26(1):16–23. doi:10.1016/j.jaut.2005.09.026

412. Jublanc C, Beaudeux JL, Aubart F, Raphael M, Chadarevian R, Chapman MJ, Bonnefont-Rousselot D, Bruckert E. Serum levels of adhesion molecules ICAM-1 and VCAM-1 and tissue inhibitor of metalloproteinases, TIMP-1, are elevated in patients with autoimmune thyroid disorders: relevance to vascular inflammation. Nutr Metab Cardiovasc Dis. 2011;21(10):817–822. doi:10.1016/j.numecd.2010.02.023

413. Bullard DC, Hu X, Schoeb TR, Collins RG, Beaudet AL, Barnum SR. Intercellular adhesion molecule-1 expression is required on multiple cell types for the development of experimental autoimmune encephalomyelitis. J Immunol. 2007;178(2):851–857. doi:10.4049/jimmunol.178.2.851

414. Lees JR. Interferon gamma in autoimmunity: A complicated player on a complex stage. Cytokine. 2015;74(1):18–26. doi:10.1016/j.cyto.2014.10.014

415. Piccio L, Buonsanti C, Mariani M, Cella M, Gilfillan S, Cross AH, Colonna M, Panina-Bordignon P. Blockade of TREM-2 exacerbates experimental autoimmune encephalomyelitis. Eur J Immunol. 2007;37(5):1290–1301. doi:10.1002/eji.200636837

416. Centa M, Prokopec KE, Garimella MG, Habir K, Hofste L, Stark JM, Dahdah A, Tibbitt CA, Polyzos KA, Gisterå A, et al. Acute Loss of Apolipoprotein E Triggers an Autoimmune Response That Accelerates Atherosclerosis. Arterioscler Thromb Vasc Biol. 2018;38(8):e145–e158. doi:10.1161/ATVBAHA.118.310802

417. Sunnemark D, Eltayeb S, Wallström E, Appelsved L, Malmberg A, Lassmann H, Ericsson-Dahlstrand A, Piehl F, Olsson T. Differential expression of the chemokine receptors CX3CR1 and CCR1 by microglia and macrophages in myelin-oligodendrocyte-glycoprotein-induced experimental autoimmune encephalomyelitis. Brain Pathol. 2003;13(4):617–629. doi:10.1111/j.1750-3639.2003.tb00490.x

418. Tanaka T, Narazaki M, Ogata A, Kishimoto T. A new era for the treatment of inflammatory autoimmune diseases by interleukin-6 blockade strategy. Semin Immunol. 2014;26(1):88–96. doi:10.1016/j.smim.2014.01.009

419. Bian Z, Miao Q, Zhong W, Zhang H, Wang Q, Peng Y, Chen X, Guo C, Shen L, Yang F, et al. Treatment of cholestatic fibrosis by altering gene expression of Cthrc1: Implications for autoimmune and non-autoimmune liver disease. J Autoimmun. 2015;63:76–87. doi:10.1016/j.jaut.2015.07.01

420. Wan S, Liu L, Ren B, Qu M, Wu H, Jiang W, Wang X, Shen H. DNA Methylation Patterns in the HLA-DPB1 and PDCD1LG2 Gene Regions in Patients with Autoimmune Thyroiditis from Different Water Iodine Areas. Thyroid. 2021;31(11):1741–1748. doi:10.1089/thy.2021.0221

421. Wu B, Wang W, Zhan Y, Li F, Zou S, Sun L, Cheng Y. CXCL13, CCL4, and sTNFR as circulating inflammatory cytokine markers in primary and SLE-related autoimmune hemolytic anemia. J Transl Med. 2015;13:112. doi:10.1186/s12967-015-0474-4

422. Zhang X, Kiapour N, Kapoor S, Khan T, Thamilarasan M, Tao Y, Cohen S, Miller R, Sobel RA, Markovic-Plese S. IL-11 Induces Encephalitogenic Th17 Cells in Multiple Sclerosis and Experimental Autoimmune Encephalomyelitis. J Immunol. 2019;203(5):1142–1150. doi:10.4049/jimmunol.1900311

423. Zhang S, Gong Y, Xiao J, Chai Y, Lei J, Huang H, Xiang T, Shen W. A COL1A1 Promoter-Controlled Expression of TGF-β Soluble Receptor Inhibits Hepatic Fibrosis Without Triggering Autoimmune Responses. Dig Dis Sci. 2018;63(10):2662–2672. doi:10.1007/s10620-018-5168-3

424. Choi EK, Kim MH, Jang SJ, Lee KH, Hwang CY, Moon SH, Lee TY, Koh CO, Park DH, Lee SS, et al. Differences in pancreatic immunohistochemical staining profiles of TGF-beta1, MMP-2, and TIMP-2 between autoimmune and alcoholic chronic pancreatitis. Pancreas. 2009;38(7):739–745. doi:10.1097/MPA.0b013e3181abab36

425. Su H, Rei N, Zhang L, Cheng J. Meta-analyses of IL1A polymorphisms and the risk of several autoimmune diseases published in databases. PLoS One. 2018;13(6):e0198693. doi:10.1371/journal.pone.0198693

426. Tan Y, Zhao M, Xiang B, Chang C, Lu Q. CD24: from a Hematopoietic Differentiation Antigen to a Genetic Risk Factor for Multiple Autoimmune Diseases. Clin Rev Allergy Immunol. 2016;50(1):70–83. doi:10.1007/s12016-015-8470-2

427. Allard DE, Wang Y, Li JJ, Conley B, Xu EW, Sailer D, Kimpston C, Notini R, Smith CJ, Koseoglu E, et al. Schwann cell-derived periostin promotes autoimmune peripheral polyneuropathy via macrophage recruitment. J Clin Invest. 2018;128(10):4727–4741. doi:10.1172/JCI99308

428. Russo V, Klein T, Lim DJ, Solis N, Machado Y, Hiroyasu S, Nabai L, Shen Y, Zeglinski MR, Zhao H, et al. Granzyme B is elevated in autoimmune blistering diseases and cleaves key anchoring proteins of the dermal-epidermal junction. Sci Rep. 2018;8(1):9690. doi:10.1038/s41598-018-28070-0

429. Iglesias M, Augustin JJ, Alvarez P, Santiuste I, Postigo J, Merino J, Merino R. Selective Impairment of TH17-Differentiation and Protection against Autoimmune Arthritis after Overexpression of BCL2A1 in T Lymphocytes. PLoS One. 2016;11(7):e0159714. doi:10.1371/journal.pone.0159714

430. Ghosh D, Curtis AD 2nd, Wilkinson DS, Mannie MD. Depletion of CD4+ CD25+ regulatory T cells confers susceptibility to experimental autoimmune encephalomyelitis (EAE) in GM-CSF-deficient Csf2-/- mice. J Leukoc Biol. 2016;100(4):747–760. doi:10.1189/jlb.3A0815-359R

431. Chang Y, Han P, Wang Y, Jia C, Zhang B, Zhao Y, Li S, Li S, Wang X, Yang X, et al. Tryptophan 2,3-dioxygenase 2 plays a key role in regulating the activation of fibroblast-like synoviocytes in autoimmune arthritis. Br J Pharmacol. 2022;179(12):3024–3042. doi:10.1111/bph.15787

432. Wang K, Zhu Q, Lu Y, Lu H, Zhang F, Wang X, Fan Y. CTLA-4 +49 G/A Polymorphism Confers Autoimmune Disease Risk: An Updated Meta-Analysis. Genet Test Mol Biomarkers. 2017;21(4):222–227. doi:10.1089/gtmb.2016.0335

433. Sawicka B, Borysewicz-Sańczyk H, Wawrusiewicz-Kurylonek N, Aversa T, Corica D, Gościk J, Krętowski A, Waśniewska M, Bossowski A. Analysis of Polymorphisms rs7093069-IL-2RA, rs7138803-FAIM2, and rs1748033-PADI4 in the Group of Adolescents With Autoimmune Thyroid Diseases. Front Endocrinol (Lausanne). 2020;11:544658. doi:10.3389/fendo.2020.544658

434. Ooi JD, Jiang JH, Eggenhuizen PJ, Chua LL, van Timmeren M, Loh KL, O’Sullivan KM, Gan PY, Zhong Y, Tsyganov K, et al. A plasmid-encoded peptide from Staphylococcus aureus induces anti-myeloperoxidase nephritogenic autoimmunity. Nat Commun. 2019;10(1):3392. doi:10.1038/s41467-019-11255-0

435. Galicia G, Maes W, Verbinnen B, Kasran A, Bullens D, Arredouani M, Ceuppens JL. Haptoglobin deficiency facilitates the development of autoimmune inflammation. Eur J Immunol. 2009;39(12):3404–3412. doi:10.1002/eji.200939291

436. Johnson C, Rosen P, Lloyd T, Horton M, Christopher-Stine L, Oddis CV, Mammen AL, Danoff SK.Exploration of the MUC5B promoter variant and ILD risk in patients with autoimmune myositis. Respir Med. 2017;130:52–54. doi:10.1016/j.rmed.2017.07.010

437. Kieseier BC, Clements JM, Pischel HB, Wells GM, Miller K, Gearing AJ, Hartung HP. Matrix metalloproteinases MMP-9 and MMP-7 are expressed in experimental autoimmune neuritis and the Guillain-Barré syndrome. Ann Neurol. 1998;43(4):427–434. doi:10.1002/ana.410430404

438. Marsillach J, Becker JO, Vaisar T, Hahn BH, Brunzell JD, Furlong CE, de Boer IH, McMahon MA, Hoofnagle AN. Paraoxonase-3 is depleted from the high-density lipoproteins of autoimmune disease patients with subclinical atherosclerosis. J Proteome Res. 2015;14(5):2046–2054. doi:10.1021/pr5011586

439. Manterola A, Bernal-Chico A, Cipriani R, Ruiz A, Pérez-Samartín A, Moreno-Rodríguez M, Hsu KL, Cravatt BF, Brown JM, Rodríguez-Puertas R, et al. Re-examining the potential of targeting ABHD6 in multiple sclerosis: Efficacy of systemic and peripherally restricted inhibitors in experimental autoimmune encephalomyelitis. Neuropharmacology. 2018;141:181–191. doi:10.1016/j.neuropharm.2018.08.038

440. Nagata K, Kumata K, Nakayama Y, Satoh Y, Sugihara H, Hara S, Matsushita M, Kuwamoto S, Kato M, Murakami I, et al. Epstein-Barr Virus Lytic Reactivation Activates B Cells Polyclonally and Induces Activation-Induced Cytidine Deaminase Expression: A Mechanism Underlying Autoimmunity and Its Contribution to Graves’ Disease. Viral Immunol. 2017;30(3):240–249. doi:10.1089/vim.2016.0179

441. Mori H, Shinoda M, Mizutani T. The N-terminal of human UGT1A6 is on the outside, as evidenced by ELISA with autoantibody in autoimmune hepatitis sera. Drug Metab Lett. 2007;1(4):261–266. doi:10.2174/187231207783221484

442. Lee YH, Bae SC. Association between Functional CYP2D6 Polymorphisms and Susceptibility to Autoimmune Diseases: A Meta-Analysis. Immunol Invest. 2017;46(2):109–122. doi:10.1080/08820139.2016.1226898

443. Muraki Y, Mizuno S, Nakatani K, Wakabayashi H, Ishikawa E, Araki T, Taniguchi A, Isaji S, Okuda M. Monitoring of peripheral blood cluster of differentiation 4+ adenosine triphosphate activity and CYP3A5 genotype to determine the pharmacokinetics, clinical effects and complications of tacrolimus in patients with autoimmune diseases. Exp Ther Med. 2018;15(1):532–538. doi:10.3892/etm.2017.5364

444. Graham KL, Werner BJ, Moyer KM, Patton AK, Krois CR, Yoo HS, Tverskoy M, LaJevic M, Napoli JL, Sobel RA, et al. DGAT1 inhibits retinol-dependent regulatory T cell formation and mediates autoimmune encephalomyelitis. Proc Natl Acad Sci U S A. 2019;116(8):3126–3135. doi:10.1073/pnas.1817669116

445. Amara K, Clay E, Yeo L, Ramsköld D, Spengler J, Sippl N, Cameron JA, Israelsson L, Titcombe PJ, Grönwall C, et al. B cells expressing the IgA receptor FcRL4 participate in the autoimmune response in patients with rheumatoid arthritis. J Autoimmun. 2017;81:34–43. doi:10.1016/j.jaut.2017.03.004

446. Hou X, Mao J, Li Y, Li J, Wang W, Fan C, Wang H, Zhang H, Shan Z, Teng W. Association of single nucleotide polymorphism rs3792876 in SLC22A4 gene with autoimmune thyroid disease in a Chinese Han population. BMC Med Genet. 2015;16:76. doi:10.1186/s12881-015-0222-x

447. Seong JM, Yee J, Gwak HS. Dipeptidyl peptidase-4 inhibitors lower the risk of autoimmune disease in patients with type 2 diabetes mellitus: A nationwide population-based cohort study. Br J Clin Pharmacol. 2019;85(8):1719–1727. doi:10.1111/bcp.13955

448. Platten M, Youssef S, Hur EM, Ho PP, Han MH, Lanz TV, Phillips LK, Goldstein MJ, Bhat R, Raine CS, et al. Blocking angiotensin-converting enzyme induces potent regulatory T cells and modulates TH1- and TH17-mediated autoimmunity. Proc Natl Acad Sci U S A. 2009;106(35):14948–14953. doi:10.1073/pnas.0903958106

449. Tsai LJ, Hsiao SH, Tsai LM, Lin CY, Tsai JJ, Liou DM, Lan JL. The sodium-dependent glucose cotransporter SLC5A11 as an autoimmune modifier gene in SLE. Tissue Antigens. 2008;71(2):114–126. doi:10.1111/j.1399-0039.2007.00975.x

450. Abad C, Jayaram B, Becquet L, Wang Y, O’Dorisio MS, Waschek JA, Tan YV. VPAC1 receptor (Vipr1)-deficient mice exhibit ameliorated experimental autoimmune encephalomyelitis, with specific deficits in the effector stage. J Neuroinflammation. 2016;13(1):169. doi:10.1186/s12974-016-0626-3

451. Duchatelet S, Caillat-Zucman S, Dubois-Laforgue D, Blanc H, Timsit J, Julier C. FCRL3 -169CT functional polymorphism in type 1 diabetes and autoimmunity traits. Biomed Pharmacother. 2008;62(3):153–157. doi:10.1016/j.biopha.2007.09.003

452. He W, Zhao J, Liu X, Li S, Mu K, Zhang J, Zhang JA. Associations between CD160 polymorphisms and autoimmune thyroid disease: a case-control study. BMC Endocr Disord. 2021;21(1):148. doi:10.1186/s12902-021-00810-w

453. Laaksonen H, Guerreiro-Cacais AO, Adzemovic MZ, Parsa R, Zeitelhofer M, Jagodic M, Olsson T. The multiple sclerosis risk gene IL22RA2 contributes to a more severe murine autoimmune neuroinflammation. Genes Immun. 2014;15(7):457–465. doi:10.1038/gene.2014.36

454. Cai M, Chen S, Hu W. MicroRNA-141 Is Involved in Ulcerative Colitis Pathogenesis via Aiming at CXCL5. J Interferon Cytokine Res. 2017;37(9):415–420. doi:10.1089/jir.2017.0019

455. Turner D, Leach ST, Mack D, Uusoue K, McLernon R, Hyams J, Leleiko N, Walters TD, Crandall W, Markowitz J, et al. Faecal calprotectin, lactoferrin, M2-pyruvate kinase and S100A12 in severe ulcerative colitis: a prospective multicentre comparison of predicting outcomes and monitoring response. Gut. 2010;59(9):1207–1212. doi:10.1136/gut.2010.211755

456. Li H, Feng C, Fan C, Yang Y, Yang X, Lu H, Lu Q, Zhu F, Xiang C, Zhang Z, et al. Intervention of oncostatin M-driven mucosal inflammation by berberine exerts therapeutic property in chronic ulcerative colitis. Cell Death Dis. 2020;11(4):271. doi:10.1038/s41419-020-2470-8

457. Kourkoulis P, Michalopoulos G, Katifelis H, Giannopoulou I, Lazaris AC, Papaconstantinou I, Karamanolis G, Gazouli M. Leucine-rich alpha-2 glycoprotein 1, high mobility group box 1, matrix metalloproteinase 3 and annexin A1 as biomarkers of ulcerative colitis endoscopic and histological activity. Eur J Gastroenterol Hepatol. 2020;32(9):1106–1115. doi:10.1097/MEG.0000000000001783

458. Li X, Gopinath SCB, Peng X, Lv J. A Zeolite Nanoparticle-Modified Anionic Surface for Aptasensing Lipocalin-2 in Ulcerative Colitis by Dual-Electrodes. J Biomed Nanotechnol. 2021;17(12):2495–2504. doi:10.1166/jbn.2021.3213

459. Egesten A, Eliasson M, Olin AI, et al. The proinflammatory CXC-chemokines GRO-alpha/CXCL1 and MIG/CXCL9 are concomitantly expressed in ulcerative colitis and decrease during treatment with topical corticosteroids. Int J Colorectal Dis. 2007;22(12):1421–1427. doi:10.1007/s00384-007-0370-3

460. Su S, Kong W, Zhang J, Wang X, Guo H. Integrated analysis of DNA methylation and gene expression profiles identified S100A9 as a potential biomarker in ulcerative colitis. Biosci Rep. 2020;40(12):BSR20202384. doi:10.1042/BSR20202384

461. Mo JS, Na KS, Yu JI, Chae SC. Identification of the polymorphisms in IFITM1 gene and their association in a Korean population with ulcerative colitis. Immunol Lett. 2013;156(1-2):118–122. doi:10.1016/j.imlet.2013.09.026

462. Chen Q, Fang X, Yao N, Wu F, Xu B, Chen Z. Suppression of miR-330-3p alleviates DSS-induced ulcerative colitis and apoptosis by upregulating the endoplasmic reticulum stress components XBP1. Hereditas. 2020;157(1):18. doi:10.1186/s41065-020-00135-z

463. Seo GS, Lee JK, Yu JI, Yun KJ, Chae SC, Choi SC. Identification of the polymorphisms in IFITM3 gene and their association in a Korean population with ulcerative colitis. Exp Mol Med. 2010;42(2):99–104. doi:10.3858/emm.2010.42.2.011

464. Lacruz-Guzmán D, Torres-Moreno D, Pedrero F, Romero-Cara P, García-Tercero I, Trujillo-Santos J, Conesa-Zamora P. Influence of polymorphisms and TNF and IL1β serum concentration on the infliximab response in Crohn’s disease and ulcerative colitis. Eur J Clin Pharmacol. 2013;69(3):431–438. doi:10.1007/s00228-012-1389-0

465. Gao S, Li Y, Wu D, Jiao N, Yang L, Zhao R, Xu Z, Chen W, Lin X, Cheng S, et al. IBD Subtype-Regulators IFNG and GBP5 Identified by Causal Inference Drive More Intense Innate Immunity and Inflammatory Responses in CD Than Those in UC. Front Pharmacol. 2022;13:869200. doi:10.3389/fphar.2022.869200

466. Li N, Zhang Y, Nepal N, Li G, Yang N, Chen H, Lin Q, Ji X, Zhang S, Jin S. Dental pulp stem cells overexpressing hepatocyte growth factor facilitate the repair of DSS-induced ulcerative colitis. Stem Cell Res Ther. 2021;12(1):30. doi:10.1186/s13287-020-02098-4

467. Elia G, Guglielmi G. CXCL9 chemokine in ulcerative colitis. Clin Ter. 2018;169(5):e235–e241. doi:10.7417/CT.2018.2085

468. Kotlowski R, Bernstein CN, Silverberg MS, Krause DO. Population-based case-control study of alpha 1-antitrypsin and SLC11A1 in Crohn’s disease and ulcerative colitis. Inflamm Bowel Dis. 2008;14(8):1112–1117. doi:10.1002/ibd.20425

469. Yamamoto-Furusho JK, Santiago-Hernández JJ, Pérez-Hernández N, Ramírez-Fuentes S, Fragoso JM, Vargas-Alarcón G. Interleukin 1 β (IL-1B) and IL-1 antagonist receptor (IL-1RN) gene polymorphisms are associated with the genetic susceptibility and steroid dependence in patients with ulcerative colitis. J Clin Gastroenterol. 2011;45(6):531–535. doi:10.1097/MCG.0b013e3181faec51

470. Xie F, Zhang H, Zheng C, Shen XF. Costunolide improved dextran sulfate sodium-induced acute ulcerative colitis in mice through NF-κB, STAT1/3, and Akt signaling pathways. Int Immunopharmacol. 2020;84:106567. doi:10.1016/j.intimp.2020.106567

471. Ding Q, Zhou H, Yun B, Zhou L, Zhang N, Yin G, Fan J. Interleukin-13 Inhibits Expression of cyp27b1 in Peripheral CD14+ Cells That Is Correlated With Vertebral Bone Mineral Density of Patients With Ulcerative Colitis. J Cell Biochem. 2017;118(2):376–381. doi:10.1002/jcb.25646

472. Wang YD, Tan XY, Zhang K. Correlation of plasma MMP-1 and TIMP-1 levels and the colonic mucosa expressions in patients with ulcerative colitis. Mediators Inflamm. 2009;2009:275072. doi:10.1155/2009/275072

473. Elhennawy MG, Abdelaleem EA, Zaki AA, Mohamed WR. Cinnamaldehyde and hesperetin attenuate TNBS-induced ulcerative colitis in rats through modulation of the JAk2/STAT3/SOCS3 pathway. J Biochem Mol Toxicol. 2021;35(5):e22730. doi:10.1002/jbt.22730

474. Saruta M, Targan SR, Mei L, Ippoliti AF, Taylor KD, Rotter JI. High-frequency haplotypes in the X chromosome locus TLR8 are associated with both CD and UC in females. Inflamm Bowel Dis. 2009;15(3):321–327. doi:10.1002/ibd.20754

475. Makidono C, Mizuno M, Nasu J, Hiraoka S, Okada H, Yamamoto K, Fujita T, Shiratori Y. Increased serum concentrations and surface expression on peripheral white blood cells of decay-accelerating factor (CD55) in patients with active ulcerative colitis. J Lab Clin Med. 2004;143(3):152–158. doi:10.1016/j.lab.2003.11.004

476. Lee DS, Lee KL, Jeong JB, Shin S, Kim SH, Kim JW. Expression of Chemokine CCL28 in Ulcerative Colitis Patients. Gut Liver. 2021;15(1):70–76. doi:10.5009/gnl19273

477. Xia SL, Lin DP, Lin QR, Sun L, Wang XQ, Hong WJ, Lin ZJ, Du CC, Jiang Y. A Case-Control Study on Association of Ulcerative Colitis with FCGR2A Gene Polymorphisms in Chinese Patients. Genet Test Mol Biomarkers. 2018;22(10):607–614. doi:10.1089/gtmb.2018.0042

478. Magnusson MK, Strid H, Isaksson S, Bajor A, Lasson A, Ung KA, Öhman L. Response to infliximab therapy in ulcerative colitis is associated with decreased monocyte activation, reduced CCL2 expression and downregulation of Tenascin C. J Crohns Colitis. 2015;9(1):56–65. doi:10.1093/ecco-jcc/jju008

479. Gupta A, Juyal G, Sood A, Midha V, Yamazaki K, Vich Vila A, Esaki M, Matsui T, Takahashi A, Kubo M, et al. A cross-ethnic survey of CFB and SLC44A4, Indian ulcerative colitis GWAS hits, underscores their potential role in disease susceptibility. Eur J Hum Genet. 2016;25(1):111–122. doi:10.1038/ejhg.2016.131

480. Sivaram G, Tiwari SK, Bardia A, Anjum F, Vishnupriya S, Habeeb A, Khan AA. Macrophage migration inhibitory factor, Toll-like receptor 4, and CD14 polymorphisms with altered expression levels in patients with ulcerative colitis. Hum Immunol. 2012;73(2):201–205. doi:10.1016/j.humimm.2011.12.006

481. Chen LH, Zhang Q, Xiao YF, Fang YC, Xie X, Nan FJ. Phosphodiesters as GPR84 Antagonists for the Treatment of Ulcerative Colitis. J Med Chem. 2022;65(5):3991–4006. doi:10.1021/acs.jmedchem.1c01813

482. Marinelli C, Zingone F, Lupo MG, Marin R, D’Incà R, Gubbiotti A, Massimi D, Casadei C, Barberio B, Ferri N, et al. Serum Levels of PCSK9 Are Increased in Patients With Active Ulcerative Colitis Representing a Potential Biomarker of Disease Activity: A Cross-sectional Study. J Clin Gastroenterol. 2021;10.1097/MCG.0000000000001607. doi:10.1097/MCG.0000000000001607

483. Xia SL, Ying SJ, Lin QR, Wang XQ, Hong WJ, Lin ZJ, Luo JK, Jiang Y. Association of Ulcerative Colitis with FOXP3 Gene Polymorphisms and Its Colonic Expression in Chinese Patients. Gastroenterol Res Pract. 2019;2019:4052168. doi:10.1155/2019/4052168

484. Kosaka T, Yoshino J, Inui K, Wakabayashi T, Okushima K, Kobayashi T, Miyoshi H, Nakamura Y, Hayashi S, Shiraishi T, et al. Impact of lipoprotein lipase gene polymorphisms on ulcerative colitis. World J Gastroenterol. 2006;12(39):6325–6330. doi:10.3748/wjg.v12.i39.6325

485. Mora-Buch R, Dotti I, Planell N, Calderón-Gómez E, Jung P, Masamunt MC, Llach J, Ricart E, Batlle E, Panés J, et al. Epithelial IL-1R2 acts as a homeostatic regulator during remission of ulcerative colitis. Mucosal Immunol. 2016;9(4):950–959. doi:10.1038/mi.2015.108

486. Zhang Y, Han D, Yu S, An C, Liu X, Zhong H, Xu Y, Jiang L, Wang Z. Protective Effect of Iridoid Glycosides of the Leaves of Syringa oblata Lindl. on Dextran Sulfate Sodium-Induced Ulcerative Colitis by Inhibition of the TLR2/4/MyD88/NF-κB Signaling Pathway. Biomed Res Int. 2020;2020:7650123. doi:10.1155/2020/7650123

487. Villani AC, Lemire M, Louis E, Silverberg MS, Collette C, Fortin G, Nimmo ER, Renaud Y, Brunet S, Libioulle C, et al. Genetic variation in the familial Mediterranean fever gene (MEFV) and risk for Crohn’s disease and ulcerative colitis. PLoS One. 2009;4(9):e7154. doi:10.1371/journal.pone.0007154

488. Zezos P, Papaioannou G, Nikolaidis N, Vasiliadis T, Giouleme O, Evgenidis N. Elevated plasma von Willebrand factor levels in patients with active ulcerative colitis reflect endothelial perturbation due to systemic inflammation. World J Gastroenterol. 2005;11(48):7639–7645. doi:10.3748/wjg.v11.i48.7639

489. Freire P, Cardoso R, Figueiredo P, Donato MM, Ferreira M, Mendes S, Ferreira AM, Vasconcelos H, Portela F, Sofia C. NOD2 gene mutations in ulcerative colitis: useless or misunderstood?. Int J Colorectal Dis. 2014;29(6):653–661. doi:10.1007/s00384-014-1850-x

490. Fukui H, Sekikawa A, Tanaka H, Fujimori Y, Katake Y, Fujii S, Ichikawa K, Tomita S, Imura J, Chiba T, et al. DMBT1 is a novel gene induced by IL-22 in ulcerative colitis. Inflamm Bowel Dis. 2011;17(5):1177–1188. doi:10.1002/ibd.21473

491. Regeling A, Imhann F, Volders HH, Blokzijl T, Bloks VW, Weersma RK, Dijkstra G, Faber KN. HSPA6 is an ulcerative colitis susceptibility factor that is induced by cigarette smoke and protects intestinal epithelial cells by stabilizing anti-apoptotic Bcl-XL. Biochim Biophys Acta. 2016;1862(4):788–796. doi:10.1016/j.bbadis.2016.01.020

492. Huang R, Wang K, Gao L, Gao W. TIMP1 Is A Potential Key Gene Associated With The Pathogenesis And Prognosis Of Ulcerative Colitis-Associated Colorectal Cancer. Onco Targets Ther. 2019;12:8895–8904. doi:10.2147/OTT.S222608

493. Vainer B, Nielsen OH, Horn T. Comparative studies of the colonic in situ expression of intercellular adhesion molecules (ICAM-1, -2, and -3), beta2 integrins (LFA-1, Mac-1, and p150,95), and PECAM-1 in ulcerative colitis and Crohn’s disease. Am J Surg Pathol. 2000;24(8):1115–1124. doi:10.1097/00000478-200008000-00009

494. Yu W, Lin Z, Hegarty JP, Chen X, Kelly AA, Wang Y, Poritz LS, Koltun WA. Genes differentially regulated by NKX2-3 in B cells between ulcerative colitis and Crohn’s disease patients and possible involvement of EGR1. Inflammation. 2012;35(3):889–899. doi:10.1007/s10753-011-9390-9

495. Ahrens R, Waddell A, Seidu L, Blanchard C, Carey R, Forbes E, Lampinen M, Wilson T, Cohen E, Stringer K, et al. Intestinal macrophage/epithelial cell-derived CCL11/eotaxin-1 mediates eosinophil recruitment and function in pediatric ulcerative colitis. J Immunol. 2008;181(10):7390–7399. doi:10.4049/jimmunol.181.10.7390

496. Padua D, Mahurkar-Joshi S, Law IK, Polytarchou C, Vu JP, Pisegna JR, Shih D, Iliopoulos D, Pothoulakis C. A long noncoding RNA signature for ulcerative colitis identifies IFNG-AS1 as an enhancer of inflammation. Am J Physiol Gastrointest Liver Physiol. 2016;311(3):G446–G457. doi:10.1152/ajpgi.00212.2016

497. Li K, Wang B, Sui H, Liu S, Yao S, Guo L, Mao D. Polymorphisms of the macrophage inflammatory protein 1 alpha and ApoE genes are associated with ulcerative colitis. Int J Colorectal Dis. 2009;24(1):13–17. doi:10.1007/s00384-008-0575-0

498. Cartwright CA, Coad CA, Egbert BM. Elevated c-Src tyrosine kinase activity in premalignant epithelia of ulcerative colitis. J Clin Invest. 1994;93(2):509–515. doi:10.1172/JCI117000

499. Parisinos CA, Serghiou S, Katsoulis M, George MJ, Patel RS, Hemingway H, Hingorani AD. Variation in Interleukin 6 Receptor Gene Associates With Risk of Crohn’s Disease and Ulcerative Colitis. Gastroenterology. 2018;155(2):303–306.e2. doi:10.1053/j.gastro.2018.05.022

500. Sabzevary-Ghahfarokhi M, Shohan M, Shirzad H, Rahimian G, Bagheri N, Soltani A, Deris F, Ghatreh-Samani M, Razmara E. The expression analysis of Fra-1 gene and IL-11 protein in Iranian patients with ulcerative colitis. BMC Immunol. 2018;19(1):17. doi:10.1186/s12865-018-0257-9

501. Fujimoto K, Kinoshita M, Tanaka H, Okuzaki D, Shimada Y, Kayama H, Okumura R, Furuta Y, Narazaki M, Tamura A, et al. Regulation of intestinal homeostasis by the ulcerative colitis-associated gene RNF186. Mucosal Immunol. 2017;10(2):446–459. doi:10.1038/mi.2016.58

502. Bai X, Bai G, Tang L, Liu L, Li Y, Jiang W. Changes in MMP-2, MMP-9, inflammation, blood coagulation and intestinal mucosal permeability in patients with active ulcerative colitis. Exp Ther Med. 2020;20(1):269–274. doi:10.3892/etm.2020.8710

503. Diaz-Gallo LM, Medrano LM, Gómez-García M, Cardeña C, Rodrigo L, Mendoza JL, Taxonera C, Nieto A, Alcain G, Cueto I, et al. Analysis of the influence of two CD24 genetic variants in Crohn’s disease and ulcerative colitis. Hum Immunol. 2011;72(10):969–972. doi:10.1016/j.humimm.2011.05.028

504. Liu J, Jiang B. Sphk1 promotes ulcerative colitis via activating JAK2/STAT3 signaling pathway. Hum Cell. 2020;33(1):57–66. doi:10.1007/s13577-019-00283-z

505. Jenkins D, Seth R, Kummer JA, Scott BB, Hawkey CJ, Robins RA. Differential levels of granzyme B, regulatory cytokines, and apoptosis in Crohn’s disease and ulcerative colitis at first presentation. J Pathol. 2000;190(2):184–189. doi:10.1002/(SICI)1096-9896(200002)190:2<184::AID-PATH531>3.0.CO;2-E

506. Mizoshita T, Tanida S, Tsukamoto H, Ozeki K, Katano T, Ebi M, Mori Y, Kataoka H, Kamiya T, Joh T. Colon Mucosa Exhibits Loss of Ectopic MUC5AC Expression in Patients with Ulcerative Colitis Treated with Oral Tacrolimus. ISRN Gastroenterol. 2013;2013:304894. doi:10.1155/2013/304894

507. Zhang J, Wang W, Zhu S, Chen Y. Increased SERPINA3 Level Is Associated with Ulcerative Colitis. Diagnostics (Basel). 2021;11(12):2371. doi:10.3390/diagnostics11122371

508. Liu C, Mo LH, Feng BS, Jin QR, Li Y, Lin J, Shu Q, Liu ZG, Liu Z, Sun X, et al. Twist1 contributes to developing and sustaining corticosteroid resistance in ulcerative colitis. Theranostics. 2021;11(16):7797–7812. doi:10.7150/thno.62256

509. Kurose I, Miura S, Suematsu M, Serizawa H, Fukumura D, Asako H, Hibi T, Tsuchiya M. Tissue-type plasminogen activator of colonic mucosa in ulcerative colitis. Evidence of endothelium-derived fibrinolytic activation. Dig Dis Sci. 1992;37(2):307–311. doi:10.1007/BF0130818

510. Nakada N, Mikami T, Horie K, Nagashio R, Sakurai Y, Sanoyama I, Yoshida T, Sada M, Kobayashi K, Sato Y, et al. Expression of CA2 and CA9 carbonic anhydrases in ulcerative colitis and ulcerative colitis-associated colorectal cancer. Pathol Int. 2020;70(8):523–532. doi:10.1111/pin.12949

511. Magnusson MK, Vidal A, Maasfeh L, Isaksson S, Malhotra R, Olsson HK, Öhman L. Impaired Butyrate Induced Regulation of T Cell Surface Expression of CTLA-4 in Patients with Ulcerative Colitis. Int J Mol Sci. 2021;22(6):3084. doi:10.3390/ijms22063084

512. Chen CC, Isomoto H, Narumi Y, Sato K, Oishi Y, Kobayashi T, Yanagihara K, Mizuta Y, Kohno S, Tsukamoto K. Haplotypes of PADI4 susceptible to rheumatoid arthritis are also associated with ulcerative colitis in the Japanese population. Clin Immunol. 2008;126(2):165–171. doi:10.1016/j.clim.2007.09.001

513. Rath T, Roderfeld M, Halwe JM, Tschuschner A, Roeb E, Graf J. Cellular sources of MMP-7, MMP-13 and MMP-28 in ulcerative colitis. Scand J Gastroenterol. 2010;45(10):1186–1196. doi:10.3109/00365521.2010.499961

514. Skok DJ, Hauptman N, Jerala M, Zidar N. Expression of Cytokine-Coding Genes BMP8B, LEFTY1 and INSL5 Could Distinguish between Ulcerative Colitis and Crohn’s Disease. Genes (Basel). 2021;12(10):1477. doi:10.3390/genes12101477

515. Fonti R, Latella G, Caprilli R, Frieri G, Marcheggiano A, Sambuy Y. Carbonic anhydrase I reduction in colonic mucosa of patients with active ulcerative colitis. Dig Dis Sci. 1998;43(9):2086–2092. doi:10.1023/a:1018819600645

516. Englund G, Jacobson A, Rorsman F, Artursson P, Kindmark A, Rönnblom A. Efflux transporters in ulcerative colitis: decreased expression of BCRP (ABCG2) and Pgp (ABCB1). Inflamm Bowel Dis. 2007;13(3):291–297. doi:10.1002/ibd.20030

517. Qiu YE, Qin J, Luo Y, Qin SL, Mu YF, Cun R, Jiang HL, Chen JJ, Yu MH, Zhong M. Increased epoxyeicosatrienoic acids may be part of a protective mechanism in human ulcerative colitis, with increased CYP2J2 and reduced soluble epoxide hydrolase expression. Prostaglandins Other Lipid Mediat. 2018;136:9–14. doi:10.1016/j.prostaglandins.2018.03.004

518. Hale LP. Deficiency of activation-induced cytidine deaminase in a murine model of ulcerative colitis. PLoS One. 2020;15(9):e0239295. doi:10.1371/journal.pone.0239295

519. Ramirez Garcia SA, Flores Alvarado LJ, Baltazar Rodriguez LM, Garcia Cruz D. Are the CYP2D6*G and MDR1, 3435T Alleles Associated with the Risk of Ulcerative Colitis in Iranian Population?. Arch Iran Med. 2019;22(11):680–681.

520. Yamamoto Y, Nakase H, Matsuura M, Maruyama S, Masuda S. CYP3A5 Genotype as a Potential Pharmacodynamic Biomarker for Tacrolimus Therapy in Ulcerative Colitis in Japanese Patients. Int J Mol Sci. 2020;21(12):4347. doi:10.3390/ijms21124347

521. Storr M, Emmerdinger D, Diegelmann J, Pfennig S, Ochsenkühn T, Göke B, Lohse P, Brand S. The cannabinoid 1 receptor (CNR1) 1359 G/A polymorphism modulates susceptibility to ulcerative colitis and the phenotype in Crohn’s disease. PLoS One. 2010;5(2):e9453. doi:10.1371/journal.pone.0009453

522. Toledo Mauriño JJ, Fonseca-Camarillo G, Furuzawa-Carballeda J, Barreto-Zuñiga R, Martínez Benítez B, Granados J, Yamamoto-Furusho JK. TRPV Subfamily (TRPV2, TRPV3, TRPV4, TRPV5, and TRPV6) Gene and Protein Expression in Patients with Ulcerative Colitis. J Immunol Res. 2020;2020:2906845. doi:10.1155/2020/2906845

523. Cao Y, Qu C, Chen Y, Li L, Wang X. Association of ABCB1 polymorphisms and ulcerative colitis susceptibility. Int J Clin Exp Pathol. 2015;8(1):943–947.

524. Magyari L, Bene J, Komlósi K, Talián G, Faragó B, Csöngei V, Járomi L, Sáfrány E, Sipeky C, Lakner L, et al. Prevalence of SLC22A4 1672T and SLC22A5 -207C combination defined TC haplotype in Hungarian ulcerative colitis patients. Pathol Oncol Res. 2007;13(1):53–56. doi:10.1007/BF02893441

525. Matsuda T, Suzuki J, Furuya K, Masutani M, Kawakami Y. Serum angiotensin I-converting enzyme is reduced in Crohn’s disease and ulcerative colitis irrespective of genotype. Am J Gastroenterol. 2001;96(9):2705–2710. doi:10.1111/j.1572-0241.2001.03945.x

526. Li T, Liu W, Hui W, Shi T, Liu H, Feng Y, Gao F. Integrated Analysis of Ulcerative Colitis Revealed an Association between PHLPP2 and Immune Infiltration. Dis Markers. 2022;2022:4983471. doi:10.1155/2022/4983471

527. Eberhardson M, Karlén P, Linton L, Jones P, Lindberg A, Kostalla MJ, Lindh E, Odén A, Glise H, Winqvist O. Randomised, Double-blind, Placebo-controlled Trial of CCR9-targeted Leukapheresis Treatment of Ulcerative Colitis Patients. J Crohns Colitis. 2017;11(5):534–542. doi:10.1093/ecco-jcc/jjw196

528. Rong Y, Hong G, Zhu N, Liu Y, Jiang Y, Liu T. Photodynamic Therapy of Novel Photosensitizer Ameliorates TNBS-Induced Ulcerative Colitis via Inhibition of AOC1. Front Pharmacol. 2021;12:746725. doi:10.3389/fphar.2021.746725

529. Andrews CW Jr, O’Hara CJ, Goldman H, Mercurio AM, Silverman ML, Steele GD Jr. Sucrase-isomaltase expression in chronic ulcerative colitis and dysplasia. Hum Pathol. 1992;23(7):774–779. doi:10.1016/0046-8177(92)90347-6

530. Pathan S, Gowdy RE, Cooney R, Beckly JB, Hancock L, Guo C, Barrett JC, Morris A, Jewell DP. Confirmation of the novel association at the BTNL2 locus with ulcerative colitis. Tissue Antigens. 2009;74(4):322–329. doi:10.1111/j.1399-0039.2009.01314.x

531. Lohi H, Mäkelä S, Pulkkinen K, Höglund P, Karjalainen-Lindsberg ML, Puolakkainen P, Kere J. Upregulation of CFTR expression but not SLC26A3 and SLC9A3 in ulcerative colitis. Am J Physiol Gastrointest Liver Physiol. 2002;283(3):G567–G575. doi:10.1152/ajpgi.00356.2001

532. Sipponen T, Haapamäki J, Savilahti E, Alfthan H, Hämäläinen E, Rautiainen H, Koskenpato J, Nuutinen H, Färkkilä M. Fecal calprotectin and S100A12 have low utility in prediction of small bowel Crohn’s disease detected by wireless capsule endoscopy. Scand J Gastroenterol. 2012;47(7):778–784. doi:10.3109/00365521.2012.677953

533. Azramezani Kopi T, Amini Kadijani A, Parsian H, Shahrokh S, Asadzadeh Aghdaei H, Mirzaei A, Balaii H, Zali MR. The value of mRNA expression of S100A8 and S100A9 as blood-based biomarkers of inflammatory bowel disease. Arab J Gastroenterol. 2019;20(3):135–140. doi:10.1016/j.ajg.2019.07.002

534. Guo A, Ross C, Chande N, Gregor J, Ponich T, Khanna R, Sey M, Beaton M, Yan B, Kim RB, et al. High oncostatin M predicts lack of clinical remission for patients with inflammatory bowel disease on tumor necrosis factor α antagonists. Sci Rep. 2022;12(1):1185. doi:10.1038/s41598-022-05208-9

535. Csillag C, Nielsen OH, Vainer B, Olsen J, Dieckgraefe BK, Hendel J, Vind I, Dupuy C, Nielsen FC, Borup R. Expression of the genes dual oxidase 2, lipocalin 2 and regenerating islet-derived 1 alpha in Crohn’s disease. Scand J Gastroenterol. 2007;42(4):454–463. doi:10.1080/00365520600976266

536. Yu Q, Zhang S, Wang H, Zhang Y, Feng T, Chen B, He Y, Zeng Z, Chen M. TNFAIP6 is a potential biomarker of disease activity in inflammatory bowel disease. Biomark Med. 2016;10(5):473–483. doi:10.2217/bmm.16.9

537. Kaser A, Lee AH, Franke A, Glickman JN, Zeissig S, Tilg H, Nieuwenhuis EE, Higgins DE, Schreiber S, Glimcher LH, et al. XBP1 links ER stress to intestinal inflammation and confers genetic risk for human inflammatory bowel disease. Cell. 2008;134(5):743–756. doi:10.1016/j.cell.2008.07.021

538. Deutschmann C, Roggenbuck D, Schierack P. The loss of tolerance to CHI3L1 - A putative role in inflammatory bowel disease?. Clin Immunol. 2019;199:12–17. doi:10.1016/j.clim.2018.12.005

539. Verstockt B, Verstockt S, Dehairs J, Ballet V, Blevi H, Wollants WJ, Breynaert C, Van Assche G, Vermeire S, Ferrante M. Low TREM1 expression in whole blood predicts anti-TNF response in inflammatory bowel disease. EBioMedicine. 2019;40:733–742. doi:10.1016/j.ebiom.2019.01.027

540. Srivastava M, Zurakowski D, Cheifetz P, Leichtner A, Bousvaros A. Elevated serum hepatocyte growth factor in children and young adults with inflammatory bowel disease. J Pediatr Gastroenterol Nutr. 2001;33(5):548–553. doi:10.1097/00005176-200111000-00007

541. Lacher M, Kappler R, Berkholz S, Baurecht H, von Schweinitz D, Koletzko S. Association of a CXCL9 polymorphism with pediatric Crohn’s disease. Biochem Biophys Res Commun. 2007;363(3):701–707. doi:10.1016/j.bbrc.2007.09.020

542. de Buhr MF, Mähler M, Geffers R, Hansen W, Westendorf AM, Lauber J, Buer J, Schlegelberger B, Hedrich HJ, Bleich A. Cd14, Gbp1, and Pla2g2a: three major candidate genes for experimental IBD identified by combining QTL and microarray analyses. Physiol Genomics. 2006;25(3):426–434. doi:10.1152/physiolgenomics.00022.2005

543. Kominato K, Yamasaki H, Mitsuyama K, Takedatsu H, Yoshioka S, Kuwaki K, Kobayashi T, Yamauchi R, Fukunaga S, Tsuruta O, et al. Increased levels of circulating adrenomedullin following treatment with TU□100 in patients with Crohn’s disease. Mol Med Rep. 2016;14(3):2264–2268. doi:10.3892/mmr.2016.5488

544. Grip O, Janciauskiene S. Atorvastatin reduces plasma levels of chemokine (CXCL10) in patients with Crohn’s disease. PLoS One. 2009;4(5):e5263. doi:10.1371/journal.pone.0005263

545. Tang J, Zhang CB, Lyu KS, Jin ZM, Guan SX, You N, Huang M, Wang XD, Gao X. Association of polymorphisms in C1orf106, IL1RN, and IL10 with post-induction infliximab trough level in Crohn’s disease patients. Gastroenterol Rep (Oxf). 2019;8(5):367–373. doi:10.1093/gastro/goz056

546. Günther C, Ruder B, Stolzer I, Dorner H, He GW, Chiriac MT, Aden K, Strigli A, Bittel M, Zeissig S, et al. Interferon Lambda Promotes Paneth Cell Death Via STAT1 Signaling in Mice and Is Increased in Inflamed Ileal Tissues of Patients With Crohn’s Disease. Gastroenterology. 2019;157(5):1310–1322.e13. doi:10.1053/j.gastro.2019.07.031

547. Elleisy N, Rohde S, Huth A, Gittel N, Glass Ä, Möller S, Lamprecht G, Schäffler H, Jaster R. Genetic association analysis of CLEC5A and CLEC7A gene single-nucleotide polymorphisms and Crohn’s disease. World J Gastroenterol. 2020;26(18):2194–2202. doi:10.3748/wjg.v26.i18.2194

548. Huang J, Chen T, Liu Y, Lyu L, Li X, Yue W. How would serum 25(OH)D level change in patients with inflammatory bowel disease depending on intestinal mucosa vitamin D receptor (VDR) and vitamin D1-α hydroxylase (CYP27B1)?. Turk J Gastroenterol. 2019;30(2):132–138. doi:10.5152/tjg.2018.17828

549. Arihiro S, Ohtani H, Hiwatashi N, Torii A, Sorsa T, Nagura H. Vascular smooth muscle cells and pericytes express MMP-1, MMP-9, TIMP-1 and type I procollagen in inflammatory bowel disease. Histopathology. 2001;39(1):50–59. doi:10.1046/j.1365-2559.2001.01142.x

550. Cheng X, Zhang X, Su J, Zhang Y, Zhou W, Zhou J, Wang C, Liang H, Chen X, Shi R, et al. miR-19b downregulates intestinal SOCS3 to reduce intestinal inflammation in Crohn’s disease. Sci Rep. 2015;5:10397. doi:10.1038/srep10397

551. Chongsrisawat V, Suratannon N, Chatchatee P, Ittiwut R, Ittiwut C, Weerapakorn W, Theamboonlers A, Rohlfs M, Klein C, Kotlarz D, et al. Novel CD55 Mutation Associated With Severe Small Bowel Ulceration Mimicking Inflammatory Bowel Disease in a Pair of Siblings. Inflamm Bowel Dis. 2022;izac001. doi:10.1093/ibd/izac001

552. Li Q, Lian Y, Deng Y, Chen J, Wu T, Lai X, Zheng B, Qiu C, Peng Y, Li W, et al. mRNA-engineered mesenchymal stromal cells expressing CXCR2 enhances cell migration and improves recovery in IBD. Mol Ther Nucleic Acids. 2021;26:222–236. doi:10.1016/j.omtn.2021.07.009

553. Fagundes RR, Bourgonje AR, Hu S, Barbieri R, Jansen BH, Sinnema N, Blokzijl T, Taylor CT, Weersma RK, Faber KN, et al. HIF1α-Dependent Induction of TFRC by a Combination of Intestinal Inflammation and Systemic Iron Deficiency in Inflammatory Bowel Disease. Front Physiol. 2022;13:889091. doi:10.3389/fphys.2022.889091

554. Ostvik AE, Granlund Av, Gustafsson BI, Torp SH, Espevik T, Mollnes TE, Damås JK, Sandvik AK. Mucosal toll-like receptor 3-dependent synthesis of complement factor B and systemic complement activation in inflammatory bowel disease. Inflamm Bowel Dis. 2014;20(6):995–1003. doi:10.1097/MIB.0000000000000035

555. Azzam N, Nounou H, Alharbi O, Aljebreen A, Shalaby M. CARD15/NOD2, CD14 and toll-like 4 receptor gene polymorphisms in Saudi patients with Crohn’s Disease. Int J Mol Sci. 2012;13(4):4268–4280. doi:10.3390/ijms13044268

556. Street ME, de’Angelis G, Camacho-Hübner C, Giovannelli G, Ziveri MA, Bacchini PL, Bernasconi S, Sansebastiano G, Savage MO. Relationships between serum IGF-1, IGFBP-2, interleukin-1beta and interleukin-6 in inflammatory bowel disease. Horm Res. 2004;61(4):159–164. doi:10.1159/000075699

557. Elleisy N, Rohde S, Huth A, Gittel N, Glass Ä, Möller S, Lamprecht G, Schäffler H, Jaster R. Genetic association analysis of CLEC5A and CLEC7A gene single-nucleotide polymorphisms and Crohn’s disease. World J Gastroenterol. 2020;26(18):2194–2202. doi:10.3748/wjg.v26.i18.2194

558. Lee A, Kanuri N, Zhang Y, Sayuk GS, Li E, Ciorba MA. IDO1 and IDO2 non-synonymous gene variants: correlation with crohn’s disease risk and clinical phenotype. PLoS One. 2014;9(12):e115848. doi:10.1371/journal.pone.0115848

559. Xia S, Zhang D, Zheng S, Wu C, Lin Q, Ying S, Shao X, Jiang Y. Association of Crohn’s disease with Foxp3 gene polymorphisms and its colonic expression in Chinese patients. J Clin Lab Anal. 2019;33(4):e22835. doi:10.1002/jcla.22835

560. Demetter P, De Vos M, Van Huysse JA, Baeten D, Ferdinande L, Peeters H, Mielants H, Veys EM, De Keyser F, Cuvelier CA. Colon mucosa of patients both with spondyloarthritis and Crohn’s disease is enriched with macrophages expressing the scavenger receptor CD163. Ann Rheum Dis. 2005;64(2):321–324. doi:10.1136/ard.2003.018382

561. Dammen R, Haugen M, Svejda B, Alaimo D, Brenna O, Pfragner R, Gustafsson BI, Kidd M. The stimulatory adenosine receptor ADORA2B regulates serotonin (5-HT) synthesis and release in oxygen-depleted EC cells in inflammatory bowel disease. PLoS One. 2013;8(4):e62607. doi:10.1371/journal.pone.0062607

562. Qasem A, Naser AE, Naser SA. Enteropathogenic infections modulate intestinal serotonin transporter (SERT) function by activating Toll-like receptor 2 (TLR-2) in Crohn’s disease. Sci Rep. 2021;11(1):22624. doi:10.1038/s41598-021-02050-3

563. Adali G, Yorulmaz E, Ozkanli S, Ulasoglu C, Bayraktar B, Orhun A, Colak Y, Tuncer I. Serum concentrations of insulin-like growth factor-binding protein 5 in Crohn’s disease. World J Gastroenterol. 2013;19(47):9049–9056. doi:10.3748/wjg.v19.i47.9049

564. Connor SJ, Paraskevopoulos N, Newman R, Cuan N, Hampartzoumian T, Lloyd AR, Grimm MC. CCR2 expressing CD4+ T lymphocytes are preferentially recruited to the ileum in Crohn’s disease. Gut. 2004;53(9):1287–1294. doi:10.1136/gut.2003.028225

565. Cushing KC, Chiplunker A, Li A, Sung YJ, Geisman T, Chen LS, Cresci S, Gutierrez AM. Smoking Interacts With CHRNA5, a Nicotinic Acetylcholine Receptor Subunit Gene, to Influence the Risk of IBD-Related Surgery. Inflamm Bowel Dis. 2018;24(5):1057–1064. doi:10.1093/ibd/izx094

566. Cibor D, Owczarek D, Butenas S, Salapa K, Mach T, Undas A. Levels and activities of von Willebrand factor and metalloproteinase with thrombospondin type-1 motif, number 13 in inflammatory bowel diseases. World J Gastroenterol. 2017;23(26):4796–4805. doi:10.3748/wjg.v23.i26.4796

567. Yamamoto S, Ma X. Role of Nod2 in the development of Crohn’s disease. Microbes Infect. 2009;11(12):912–918. doi:10.1016/j.micinf.2009.06.005

568. Polley S, Prescott N, Nimmo E, Veal C, Vind I, Munkholm P, Fode P, Mansfield J, Skyt Andersen P, Satsangi J, et al. Copy number variation of scavenger-receptor cysteine-rich domains within DMBT1 and Crohn’s disease. Eur J Hum Genet. 2016;24(9):1294–1300. doi:10.1038/ejhg.2015.280

569. Ungaro F, Garlatti V, Massimino L, Spinelli A, Carvello M, Sacchi M, Spanò S, Colasante G, Valassina N, Vetrano S, et al. mTOR-Dependent Stimulation of IL20RA Orchestrates Immune Cell Trafficking through Lymphatic Endothelium in Patients with Crohn’s Disease. Cells. 2019;8(8):924. doi:10.3390/cells8080924

570. Gu L, Ge Z, Wang Y, Shen M, Zhao P. Activating transcription factor 3 promotes intestinal epithelial cell apoptosis in Crohn’s disease. Pathol Res Pract. 2018;214(6):862–870. doi:10.1016/j.prp.2018.04.013

571. Jakubowska K, Pryczynicz A, Iwanowicz P, Niewiński A, Maciorkowska E, Hapanowicz J, Jagodzińska D, Kemona A, Guzińska-Ustymowicz K. Expressions of Matrix Metalloproteinases (MMP-2, MMP-7, and MMP-9) and Their Inhibitors (TIMP-1, TIMP-2) in Inflammatory Bowel Diseases. Gastroenterol Res Pract. 2016;2016:2456179. doi:10.1155/2016/2456179

572. Truffi M, Sorrentino L, Monieri M, Fociani P, Mazzucchelli S, Bonzini M, Zerbi P, Sampietro GM, Di Sabatino A, Corsi F. Inhibition of Fibroblast Activation Protein Restores a Balanced Extracellular Matrix and Reduces Fibrosis in Crohn’s Disease Strictures Ex Vivo. Inflamm Bowel Dis. 2018;24(2):332–345. doi:10.1093/ibd/izx008

573. Verstockt B, Verstockt S, Creyns B, Tops S, Van Assche G, Gils A, Ceuppens JL, Vermeire S, Ferrante M, Breynaert C. Mucosal IL13RA2 expression predicts nonresponse to anti-TNF therapy in Crohn’s disease. Aliment Pharmacol Ther. 2019;49(5):572–581. doi:10.1111/apt.15126

574. Chae SC, Yu JI, Oh GJ, Choi CS, Choi SC, Yang YS, Yun KJ. Identification of single nucleotide polymorphisms in the TNFRSF17 gene and their association with gastrointestinal disorders. Mol Cells. 2010;29(1):21–28. doi:10.1007/s10059-010-0002-6

575. Keita ÅV, Alkaissi LY, Holm EB, Heil SDS, Chassaing B, Darfeuille-Michaud A, McKay DM, Söderholm JD. Enhanced E. coli LF82 Translocation through the Follicle-associated Epithelium in Crohn’s Disease is Dependent on Long Polar Fimbriae and CEACAM6 expression, and Increases Paracellular Permeability. J Crohns Colitis. 2020;14(2):216–229. doi:10.1093/ecco-jcc/jjz144

576. Roy S, Ghosh S, Banerjee M, Laha S, Bhattacharjee D, Sarkar R, Ray S, Banerjee A, Ghosh R, Halder A, et al. A combination of circulating microRNA-375-3p and chemokines CCL11, CXCL12, and G-CSF differentiate Crohn’s disease and intestinal tuberculosis. Sci Rep. 2021;11(1):23303. doi:10.1038/s41598-021-02383-z

577. Cui D, Huang G, Yang D, Huang B, An B. Efficacy and safety of interferon-gamma-targeted therapy in Crohn’s disease: a systematic review and meta-analysis of randomized controlled trials. Clin Res Hepatol Gastroenterol. 2013;37(5):507–513. doi:10.1016/j.clinre.2012.12.004

578. Glapa-Nowak A, Szczepanik M, Iwańczak B, Kwiecień J, Szaflarska-Popławska AB, Grzybowska-Chlebowczyk U, Osiecki M, Dziekiewicz M, Stawarski A, Kierkuś J, et al. Apolipoprotein E variants correlate with the clinical presentation of paediatric inflammatory bowel disease: A cross-sectional study. World J Gastroenterol. 2021;27(14):1483–1496. doi:10.3748/wjg.v27.i14.1483

579. Gong W, Yu J, Zheng T, Liu P, Zhao F, Liu J, Hong Z, Ren H, Gu G, Wang G, et al. CCL4-mediated targeting of spleen tyrosine kinase (Syk) inhibitor using nanoparticles alleviates inflammatory bowel disease. Clin Transl Med. 2021;11(2):e339. doi:10.1002/ctm2.339

580. Glas J, Seiderer J, Bayrle C, Wetzke M, Fries C, Tillack C, Olszak T, Beigel F, Steib C, Friedrich M, et al. The role of osteopontin (OPN/SPP1) haplotypes in the susceptibility to Crohn’s disease. PLoS One. 2011;6(12):e29309. doi:10.1371/journal.pone.0029309

581. Todhunter CE, Sutherland-Craggs A, Bartram SA, Donaldson PT, Daly AK, Francis RM, Mansfield JC, Thompson NP. nfluence of IL-6, COL1A1, and VDR gene polymorphisms on bone mineral density in Crohn’s disease. Gut. 2005;54(11):1579–1584. doi:10.1136/gut.2005.064212

582. Romagnoli C, Marcucci T, Picariello L, Tonelli F, Vincenzini MT, Iantomasi T. Role of N-acetylcysteine and GSH redox system on total and active MMP-2 in intestinal myofibroblasts of Crohn’s disease patients. Int J Colorectal Dis. 2013;28(7):915–924. doi:10.1007/s00384-012-1632-2

583. Te Velde AA, Pronk I, de Kort F, Stokkers PC. Glutathione peroxidase 2 and aquaporin 8 as new markers for colonic inflammation in experimental colitis and inflammatory bowel diseases: an important role for H2O2?. Eur J Gastroenterol Hepatol. 2008;20(6):555–560. doi:10.1097/MEG.0b013e3282f45751

584. Zheng Y, Ge W, Ma Y, Xie G, Wang W, Han L, Bian B, Li L, Shen L. miR-155 Regulates IL-10-Producing CD24hiCD27+ B Cells and Impairs Their Function in Patients with Crohn’s Disease. Front Immunol. 2017;8:914. doi:10.3389/fimmu.2017.00914

585. Mao H, Jia J, Sheng J, Zhang S, Huang K, Li H, He F. Protective and anti-inflammatory role of REG1A in inflammatory bowel disease induced by JAK/STAT3 signaling axis. Int Immunopharmacol. 2021;92:107304. doi:10.1016/j.intimp.2020.107304

586. Kawamoto A, Nagata S, Anzai S, Takahashi J, Kawai M, Hama M, Nogawa D, Yamamoto K, Kuno R, Suzuki K, et al. Ubiquitin D is Upregulated by Synergy of Notch Signalling and TNF-α in the Inflamed Intestinal Epithelia of IBD Patients. J Crohns Colitis. 2019;13(4):495–509. doi:10.1093/ecco-jcc/jjy180

587. Mahasneh S, Sharab A, Al Shhab M, Rashid M, Zihlif M. AOX1 and XDH Enzymes Genotyping and its Effect on Clinical Response to Azathioprine in Inflammatory Bowel Disease Patients Among Jordanian Population. Curr Drug Metab. 2020;21(2):140–144. doi:10.2174/1389200221666200413125011

588. Keskin M, Topkaç A. The Predictive Value of Periostin to Diagnose Crohn’s Disease. Turk J Gastroenterol. 2022;33(2):127–135. doi:10.5152/tjg.2022.21875

589. Lefferts AR, Regner EH, Stahly A, O’Rourke B, Gerich ME, Fennimore BP, Scott FI, Freeman AE, Jones K, Kuhn KA. Circulating mature granzyme B+ T cells distinguish Crohn’s disease-associated axial spondyloarthritis from axial spondyloarthritis and Crohn’s disease. Arthritis Res Ther. 2021;23(1):147. doi:10.1186/s13075-021-02531-w

590. Schürmann GM, Bishop AE, Facer P, Vecchio M, Lee JC, Rampton DS, Polak JM. Increased expression of cell adhesion molecule P-selectin in active inflammatory bowel disease. Gut. 1995;36(3):411–418. doi:10.1136/gut.36.3.411

591. Mizoshita T, Tanida S, Tsukamoto H, Ozeki K, Katano T, Nishiwaki H, Ebi M, Mori Y, Kubota E, Kataoka H, et al. Adalimumab Treatment in Biologically Naïve Crohn’s Disease: Relationship with Ectopic MUC5AC Expression and Endoscopic Improvement. Gastroenterol Res Pract. 2014;2014:687257. doi:10.1155/2014/687257

592. Irak K, Bayram M, Cifci S, Sener G. Serum levels of NLRC4 and MCP-2/CCL8 in patients with active Crohn’s disease. PLoS One. 2021;16(11):e0260034. doi:10.1371/journal.pone.0260034

593. Miseljic S, Galandiuk S, Myers SD, Wittliff JL. Expression of urokinase-type plasminogen activator and plasminogen activator inhibitor in colon disease. J Clin Lab Anal. 1995;9(6):413–417. doi:10.1002/jcla.1860090613

594. Zhou G, Yu L, Fang L, Yang W, Yu T, Miao Y, Chen M, Wu K, Chen F, Cong Y, et al. CD177+ neutrophils as functionally activated neutrophils negatively regulate IBD. Gut. 2018;67(6):1052–1063. doi:10.1136/gutjnl-2016-313535

595. Hemmati S, Sanati G, Sadeghi MA, Ebrahimi Daryani N, Rezaei N. Association between Promotor hypomethylation of TFF1 and Crohn’s Disease. Acta Biomed. 2022;93(1):e2022176. doi:10.23750/abm.v93i1.12073

596. Zeissig S, Petersen BS, Tomczak M, Melum E, Huc-Claustre E, Dougan SK, Laerdahl JK, Stade B, Forster M, Schreiber S, et al. Early-onset Crohn’s disease and autoimmunity associated with a variant in CTLA-4. Gut. 2015;64(12):1889–1897. doi:10.1136/gutjnl-2014-308541

597. Rath T, Roderfeld M, Graf J, Wagner S, Vehr AK, Dietrich C, Geier A, Roeb E. Enhanced expression of MMP-7 and MMP-13 in inflammatory bowel disease: a precancerous potential?. Inflamm Bowel Dis. 2006;12(11):1025–1035. doi:10.1097/01.mib.0000234133.97594.04

598. Lakatos PL, Kiss LS, Palatka K, Altorjay I, Antal-Szalmas P, Palyu E, Udvardy M, Molnar T, Farkas K, Veres G, et al. Serum lipopolysaccharide-binding protein and soluble CD14 are markers of disease activity in patients with Crohn’s disease. Inflamm Bowel Dis. 2011;17(3):767–777. doi:10.1002/ibd.21402

599. Rodriguez-Palacios A, Harding A, Menghini P, Himmelman C, Retuerto M, Nickerson KP, Lam M, Croniger CM, McLean MH, Durum SK, et al. The Artificial Sweetener Splenda Promotes Gut Proteobacteria, Dysbiosis, and Myeloperoxidase Reactivity in Crohn’s Disease-Like Ileitis. Inflamm Bowel Dis. 2018;24(5):1005–1020. doi:10.1093/ibd/izy060

600. Maza I, Miller-Lotan R, Levy AP, Nesher S, Karban A, Eliakim R. The association of Haptoglobin polymorphism with Crohn’s disease in Israel. J Crohns Colitis. 2008;2(3):214–218. doi:10.1016/j.crohns.2008.03.005

601. Dunkin D, Merlino F, Correale C; CEACAM5 working group, Yeretssian G, Marinelli L, Roda G. A small CEACAM5 peptide restores the protective function of CD8+ regulatory T cells in Crohn’s disease. Gastroenterology. 2022;S0016-5085(22)00630-8. doi:10.1053/j.gastro.2022.06.025

602. Date AA, Rais R, Babu T, Ortiz J, Kanvinde P, Thomas AG, Zimmermann SC, Gadiano AJ, Halpert G, Slusher BS, et al. Local enema treatment to inhibit FOLH1/GCPII as a novel therapy for inflammatory bowel disease. J Control Release. 2017;263:132–138. doi:10.1016/j.jconrel.2017.01.036

603. Yu YL, Chen M, Zhu H, Zhuo MX, Chen P, Mao YJ, Li LY, Zhao Q, Wu M, Ye M. STAT1 epigenetically regulates LCP2 and TNFAIP2 by recruiting EP300 to contribute to the pathogenesis of inflammatory bowel disease. Clin Epigenetics. 2021;13(1):127. doi:10.1186/s13148-021-01101-w

604. Begun J, Lassen KG, Jijon HB, Baxt LA, Goel G, Heath RJ, Ng A, Tam JM, Kuo SY, Villablanca EJ, et al. Integrated Genomics of Crohn’s Disease Risk Variant Identifies a Role for CLEC12A in Antibacterial Autophagy. Cell Rep. 2015;11(12):1905–1918. doi:10.1016/j.celrep.2015.05.045

605. Alsahli S, Al Anazi A, Al Hatlani MM, Kashgari A, Al Sufiani F, Alfadhel M, Al Mutairi F. Severe Crohn’s Disease Manifestations in a Child with Cystathionine β-Synthase Deficiency. ACG Case Rep J. 2018;5:e93. doi:10.14309/crj.2018.93

606. Bergemalm D, Kruse R, Sapnara M, Halfvarson J, Hörnquist EH. Elevated fecal peptidase D at onset of colitis in Galphai2-/- mice, a mouse model of IBD. PLoS One. 2017;12(3):e0174275. doi:10.1371/journal.pone.0174275

607. Deuring JJ, de Haar C, Koelewijn CL, Kuipers EJ, Peppelenbosch MP, van der Woude CJ. Absence of ABCG2-mediated mucosal detoxification in patients with active inflammatory bowel disease is due to impeded protein folding. Biochem J. 2012;441(1):87–93. doi:10.1042/BJ20111281

608. Vyhlidal CA, Chapron BD, Ahmed A, Singh V, Casini R, Shakhnovich V. Effect of Crohn’s Disease on Villous Length and CYP3A4 Expression in the Pediatric Small Intestine. Clin Transl Sci. 2021;14(2):729–736. doi:10.1111/cts.12938

609. Bystrom J, Thomson SJ, Johansson J, Edin ML, Zeldin DC, Gilroy DW, Smith AM, Bishop-Bailey D. Inducible CYP2J2 and its product 11,12-EET promotes bacterial phagocytosis: a role for CYP2J2 deficiency in the pathogenesis of Crohn’s disease?. PLoS One. 2013;8(9):e75107. doi:10.1371/journal.pone.0075107

610. Amir Shaghaghi M, Bernstein CN, Serrano León A, El-Gabalawy H, Eck P. Polymorphisms in the sodium-dependent ascorbate transporter gene SLC23A1 are associated with susceptibility to Crohn disease. Am J Clin Nutr. 2014;99(2):378–383. doi:10.3945/ajcn.113.068015

611. de Vries HS, Te Morsche RH, Jenniskens K, Peters WH, de Jong DJ. A functional polymorphism in UGT1A1 related to hyperbilirubinemia is associated with a decreased risk for Crohn’s disease. J Crohns Colitis. 2012;6(5):597–602. doi:10.1016/j.crohns.2011.11.010

612. Tronstad RR, Polushina T, Brattbakk HR, Stansberg C, von Volkmann HL, Hanevik K, Ellinghaus E, Jørgensen SF, Ersland KM, Pham KD, et al. Genetic and transcriptional analysis of inflammatory bowel disease-associated pathways in patients with GUCY2C-linked familial diarrhea. Scand J Gastroenterol. 2018;53(10-11):1264–1273. doi:10.1080/00365521.2018.1521867

613. Dudarewicz M, Rychlik-Sych M, Barańska M, Wojtczak A, Trzciński R, Dziki A, Skrętkowicz J. Significance of the genetic polymorphism of CYP2D6 and NAT2 in patients with inflammatory bowel diseases. Pharmacol Rep. 2014;66(4):686–690. doi:10.1016/j.pharep.2014.04.002

614. Hernández-Camba A, Carrillo-Palau M, Ramos L, de Armas-Rillo L, Vela M, Arranz L, González-Gay MÁ, Ferraz-Amaro I. Apolipoprotein C3 Is Downregulated in Patients With Inflammatory Bowel Disease. Clin Transl Gastroenterol. 2022;13(6):e00500. doi:10.14309/ctg.0000000000000500

615. Du J, Yin J, Du H, Zhang J. Revisiting an Expression Dataset of Discordant Inflammatory Bowel Disease Twin Pairs Using a Mutation Burden Test Reveals CYP2C18 as a Novel Marker. Front Genet. 2021;12:680125. doi:10.3389/fgene.2021.680125

616. Deolet E, Callewaert B, Geldof J, Van Biervliet S, Vande Velde S, Van Dorpe J, Van Winckel M, Hoorens A. Apoptotic enteropathy, gluten intolerance, and IBD-like inflammation associated with lipotoxicity in DGAT1 deficiency-related diarrhea: a case report of a 17-year-old patient and literature review. Virchows Arch. 2022;10.1007/s00428-022-03365-w. doi:10.1007/s00428-022-03365-w

617. Zhou T, Patel K, Harris RA, Seghers V, Walsh SM, Rodriguez R, Kellermayer R, Wu H. SULT1A1 and SULT1A2 Associated Extensive Prolapse-Type Inflammatory Polyposis in Crohn’s Colitis. Ann Clin Lab Sci. 2021;51(6):868–874.

618. Feng R, Xu PP, Chen BL, Mao R, Zhang SH, Qiu Y, Zeng ZR, Chen MH, He Y. CYP2C19 polymorphism has no correlation with the efficacy and safety of thalidomide in the treatment of immune-related bowel disease. J Dig Dis. 2020;21(2):98–103. doi:10.1111/1751-2980.12842

619. Krupoves A, Mack D, Seidman E, Deslandres C, Amre D. Associations between variants in the ABCB1 (MDR1) gene and corticosteroid dependence in children with Crohn’s disease. Inflamm Bowel Dis. 2011;17(11):2308–2317. doi:10.1002/ibd.21608

620. Toyonaga T, Araba KC, Kennedy MM, Keith BP, Wolber EA, Beasley C, Steinbach EC, Schaner MR, Jain A, Long MD, et al. Increased colonic expression of ACE2 associates with poor prognosis in Crohn’s disease. Sci Rep. 2021;11(1):13533. doi:10.1038/s41598-021-92979-2

621. Kumar S, Ranjan P, Mittal B, Ghoshal UC. Serotonin transporter gene (SLC6A4) polymorphism in patients with irritable bowel syndrome and healthy controls. J Gastrointestin Liver Dis. 2012;21(1):31–38.

622. Pochini L, Scalise M, Galluccio M, Pani G, Siminovitch KA, Indiveri C. The human OCTN1 (SLC22A4) reconstituted in liposomes catalyzes acetylcholine transport which is defective in the mutant L503F associated to the Crohn’s disease. Biochim Biophys Acta. 2012;1818(3):559–565. doi:10.1016/j.bbamem.2011.12.014

623. Klasić M, Markulin D, Vojta A, Samaržija I, Biruš I, Dobrinić P, Ventham NT, Trbojević-Akmačić I, Šimurina M, Štambuk J, et al. Promoter methylation of the MGAT3 and BACH2 genes correlates with the composition of the immunoglobulin G glycome in inflammatory bowel disease. Clin Epigenetics. 2018;10:75. doi:10.1186/s13148-018-0507-y

624. Chua KH, Lian LH, Kee BP, Thum CM, Lee WS, Hilmi I, Goh KL. Identification of DLG5 and SLC22A5 gene polymorphisms in Malaysian patients with Crohn’s disease. J Dig Dis. 2011;12(6):459–466. doi:10.1111/j.1751-2980.2011.00533.x

625. Rose M, Walter OB, Fliege H, Hildebrandt M, Mönnikes H, Klapp BF. DPP IV and mental depression in Crohn’s disease. Adv Exp Med Biol. 2003;524:321–331. doi:10.1007/0-306-47920-6_38

626. Zabana Y, Lorén V, Domènech E, Aterido A, Garcia-Jaraquemada A, Julià A, Vicario M, Pedrosa E, Ferreiro M, Troya J, Lozano JJ, et al. Transcriptomic identification of TMIGD1 and its relationship with the ileal epithelial cell differentiation in Crohn’s disease. Am J Physiol Gastrointest Liver Physiol. 2020;319(2):G109–G120. doi:10.1152/ajpgi.00027.2020

627. Zucchelli M, Torkvist L, Bresso F, Halfvarson J, Hellquist A, Anedda F, Assadi G, Lindgren GB, Svanfeldt M, Janson M, et al. PepT1 oligopeptide transporter (SLC15A1) gene polymorphism in inflammatory bowel disease. Inflamm Bowel Dis. 2009;15(10):1562–1569. doi:10.1002/ibd.20963

628. Hassan-Zahraee M, Banerjee A, Cheng JB, Zhang W, Ahmad A, Page K, von Schack D, Zhang B, Martin SW, Nayak S, et al. Anti-MAdCAM Antibody Increases ß7+ T Cells and CCR9 Gene Expression in the Peripheral Blood of Patients With Crohn’s Disease. J Crohns Colitis. 2018;12(1):77–86. doi:10.1093/ecco-jcc/jjx121

629. Bogdanos DP, Roggenbuck D, Reinhold D, Wex T, Pavlidis P, von Arnim U, Malfertheiner P, Forbes A, Conrad K, Laass MW. Pancreatic-specific autoantibodies to glycoprotein 2 mirror disease location and behaviour in younger patients with Crohn’s disease. BMC Gastroenterol. 2012;12:102. doi:10.1186/1471-230X-12-102

630. Zhuang X, Chen B, Huang S, Han J, Zhou G, Xu S, Chen M, Zeng Z, Zhang S. Hypermethylation of miR-145 promoter-mediated SOX9-CLDN8 pathway regulates intestinal mucosal barrier in Crohn’s disease. EBioMedicine. 2022;76:103846. doi:10.1016/j.ebiom.2022.103846

631. Foley A, Halmos EP, Husein DM, Fehily SR, Löscher BS, Franke A, Naim HY, Gibson PR, D’Amato M. Adult sucrase-isomaltase deficiency masquerading as IBS. Gut. 2022;71(6):1237–1238. doi:10.1136/gutjnl-2021-326153

632. Repnik K, Koder S, Skok P, Ferkolj I, Potočnik U. Transferrin Level Before Treatment and Genetic Polymorphism in HFE Gene as Predictive Markers for Response to Adalimumab in Crohn’s Disease Patients. Biochem Genet. 2016;54(4):476–486. doi:10.1007/s10528-016-9734-0

633. Banerjee S, Oneda B, Yap LM, Jewell DP, Matters GL, Fitzpatrick LR, Seibold F, Sterchi EE, Ahmad T, Lottaz D, et al. MEP1A allele for meprin A metalloprotease is a susceptibility gene for inflammatory bowel disease. Mucosal Immunol. 2009;2(3):220–231. doi:10.1038/mi.2009.3

634. Nolan DJ, Han DY, Lam WJ, Morgan AR, Fraser AG, Tapsell LC, Ferguson LR. Genetic adult lactase persistence is associated with risk of Crohn’s Disease in a New Zealand population. BMC Res Notes. 2010;3:339. doi:10.1186/1756-0500-3-339

635. Johnson CM, Traherne JA, Jamieson SE, Tremelling M, Bingham S, Parkes M, Blackwell JM, Trowsdale J. Analysis of the BTNL2 truncating splice site mutation in tuberculosis, leprosy and Crohn’s disease. Tissue Antigens. 2007;69(3):236–241. doi:10.1111/j.1399-0039.2006.00795.x

636. Yukawa T, Oshitani N, Yamagami H, Watanabe K, Higuchi K, Arakawa T. Differential expression of vasoactive intestinal peptide receptor 1 expression in inflammatory bowel disease. Int J Mol Med. 2007;20(2):161–167.

637. Undas A, Owczarek D, Gissel M, Salapa K, Mann KG, Butenas S. Activated factor XI and tissue factor in inflammatory bowel disease. Inflamm Bowel Dis. 2010;16(9):1447–1448. doi:10.1002/ibd.21206

638. Parlato M, Charbit-Henrion F, Pan J, Romano C, Duclaux-Loras R, Le Du MH, Warner N, Francalanci P, Bruneau J, Bras M, et al. Human ALPI deficiency causes inflammatory bowel disease and highlights a key mechanism of gut homeostasis. EMBO Mol Med. 2018;10(4):e8483. doi:10.15252/emmm.201708483

639. Mendoza JL, Lana R, Martin MC, de la Concha EG, Urcelay E, Diaz-Rubio M, Abreu MT, Mitchell AA. FcRL3 gene promoter variant is associated with peripheral arthritis in Crohn’s disease. Inflamm Bowel Dis. 2009;15(9):1351–1357. doi:10.1002/ibd.20895

640. Futagami M, Sakamoto T, Sakamoto A, Shigetou T, Taniguchi R, Sato S, Tutaya S, Kojima K, Yasujima M, Mizunuma H. A pregnant woman with genetic variants of butyrylcholinesterase and inflammatory bowel disease. J Obstet Gynaecol. 2006;26(6):562–563. doi:10.1080/01443610600821440

641. Kim ES, Song JS, Ki CS, Choe YH, Kang B. Development of Crohn’s Disease in a Child With SLC26A3-related Congenital Chloride Diarrhea: Report of the First Case in East Asia and a Novel Missense Variant. Ann Lab Med. 2021;41(2):255–257. doi:10.3343/alm.2021.41.2.255

642. Giudice V, Wu Z, Kajigaya S, Fernandez Ibanez MDP, Rios O, Cheung F, Ito S, Young NS. Circulating S100A8 and S100A9 protein levels in plasma of patients with acquired aplastic anemia and myelodysplastic syndromes. Cytokine. 2019;113:462–465. doi:10.1016/j.cyto.2018.06.025

643. López-Karpovitch X, Barrales-Benítez O, Flores M, Piedras J. Effect of azacytidine in the release of leukemia inhibitory factor, oncostatin m, interleukin (IL)-6, and IL-11 by mononuclear cells of patients with refractory anemia. Cytokine. 2002;20(4):154–162. doi:10.1006/cyto.2002.1998

644. Jiang W, Constante M, Santos MM. Anemia upregulates lipocalin 2 in the liver and serum. Blood Cells Mol Dis. 2008;41(2):169–174. doi:10.1016/j.bcmd.2008.04.006

645. Klahr S, Morrissey J. Obstructive nephropathy and renal fibrosis: The role of bone morphogenic protein-7 and hepatocyte growth factor. Kidney Int Suppl. 2003;(87):S105–S112. doi:10.1046/j.1523-1755.64.s87.16.x

646. Hong JH, Choi YK, Min BK, Park KS, Seong K, Lee IK, Kim JG. Relationship between hepcidin and GDF15 in anemic patients with type 2 diabetes without overt renal impairment. Diabetes Res Clin Pract. 2015;109(1):64–70. doi:10.1016/j.diabres.2015.05.001

647. Rosenberg JM, Peters JM, Hughes T, Lareau CA, Ludwig LS, Massoth LR, Austin-Tse C, Rehm HL, Bryson B, Chen YB, et al. AK inhibition in a patient with a STAT1 gain-of-function variant reveals STAT1 dysregulation as a common feature of aplastic anemia. Med (N Y). 2022;3(1):42–57.e5. doi:10.1016/j.medj.2021.12.003

648. Büyüklü M, Kürüm AT, Tatlý E, Set T. Effects of levosimendan on TNF-alpha, BNP and MMP-1 in patients with heart failure with anemia. Arq Bras Cardiol. 2012;99(1):659–664. doi:10.1590/s0066-782x2012005000055

649. Fejtkova M, Sukova M, Hlozkova K, Skvarova Kramarzova K, Rackova M, Jakubec D, Bakardjieva M, Bloomfield M, Klocperk A, Parackova Z, et al. TLR8/TLR7 dysregulation due to a novel TLR8 mutation causes severe autoimmune hemolytic anemia and autoinflammation in identical twins. Am J Hematol. 2022;97(3):338–351. doi:10.1002/ajh.26452

650. Al-Faris L, Al-Humood S, Behbehani F, Sallam H. Altered Expression Pattern of CD55 and CD59 on Red Blood Cells in Anemia of Chronic Kidney Disease. Med Princ Pract. 2017;26(6):516–522. doi:10.1159/000481823

651. Martin FM, Bydlon G, Friedman JS. SOD2-deficiency sideroblastic anemia and red blood cell oxidative stress. Antioxid Redox Signal. 2006;8(7-8):1217–1225. doi:10.1089/ars.2006.8.1217

652. Pesciotta EN, Sriswasdi S, Tang HY, Speicher DW, Mason PJ, Bessler M. Dysferlin and other non-red cell proteins accumulate in the red cell membrane of Diamond-Blackfan Anemia patients. PLoS One. 2014;9(1):e85504. doi:10.1371/journal.pone.0085504

653. In JW, Lee N, Roh EY, Shin S, Park KU, Song EY. Association of aplastic anemia and FoxP3 gene polymorphisms in Koreans. Hematology. 2017;22(3):149–154. doi:10.1080/10245332.2016.1238645

654. David S, Aguiar P, Antunes L, Dias A, Morais A, Sakuntabhai A, Lavinha J. Variants in the non-coding region of the TLR2 gene associated with infectious subphenotypes in pediatric sickle cell anemia. Immunogenetics. 2018;70(1):37–51. doi:10.1007/s00251-017-1013-7

655. Chen YC, Chao TY, Cheng SN, Hu SH, Liu JY. Prevalence of von Willebrand disease in women with iron deficiency anaemia and menorrhagia in Taiwan. Haemophilia. 2008;14(4):768–774. doi:10.1111/j.1365-2516.2008.01777.x

656. Ogura H, Ohga S, Aoki T, Utsugisawa T, Takahashi H, Iwai A, Watanabe K, Okuno Y, Yoshida K, Ogawa S, et al. Novel COL4A1 mutations identified in infants with congenital hemolytic anemia in association with brain malformations. Hum Genome Var. 2020;7(1):42. doi:10.1038/s41439-020-00130-w

657. Welsh JP, Rutherford TR, Flynn J, Foukaneli T, Gordon-Smith EC, Gibson FM. In vitro effects of interferon-gamma and tumor necrosis factor-alpha on CD34+ bone marrow progenitor cells from aplastic anemia patients and normal donors. Hematol J. 2004;5(1):39–46. doi:10.1038/sj.thj.6200340

658. Paul A, Calleja L, Vilella E, Martínez R, Osada J, Joven J. Reduced progression of atherosclerosis in apolipoprotein E-deficient mice with phenylhydrazine-induced anemia. Atherosclerosis. 1999;147(1):61–68. doi:10.1016/s0021-9150(99)00164-1

659. Hohaus S, Massini G, Giachelia M, Vannata B, Bozzoli V, Cuccaro A, D’Alo’ F, Larocca LM, Raymakers RA, Swinkels DW, et al. Anemia in Hodgkin’s lymphoma: the role of interleukin-6 and hepcidin. J Clin Oncol. 2010;28(15):2538–2543. doi:10.1200/JCO.2009.27.6873

660. Wang Y, Niu ZY, Guo YJ, Wang LH, Lin FR, Zhang JY. IL-11 promotes the treatment efficacy of hematopoietic stem cell transplant therapy in aplastic anemia model mice through a NF-κB/microRNA-204/thrombopoietin regulatory axis. Exp Mol Med. 2017;49(12):e410. doi:10.1038/emm.2017.217

661. Roomi MW, Kalinovsky T, Rath M, Niedzwiecki A. Cytokines, inducers and inhibitors modulate MMP-2 and MMP□9 secretion by human Fanconi anemia immortalized fibroblasts. Oncol Rep. 2017;37(3):1842–1848. doi:10.3892/or.2017.5368

662. Kisia LE, Cheng Q, Raballah E, Munde EO, McMahon BH, Hengartner NW, Ong’echa JM, Chelimo K, Lambert CG, Ouma C, et al. Genetic variation in CSF2 (5q31.1) is associated with longitudinal susceptibility to pediatric malaria, severe malarial anemia, and all-cause mortality in a high-burden malaria and HIV region of Kenya. Trop Med Health. 2022;50(1):41. doi:10.1186/s41182-022-00432-5

663. Phillips G, Hartman J, Keller VA, Santiago MA, Pizzo S. Regulation of tissue plasminogen activator in sickle cell anemia. Am J Hematol. 1990;35(3):167–170. doi:10.1002/ajh.2830350305

664. Liu B, Shao Y, Liang X, Lu D, Yan L, Churov A, Fu R. CTLA-4 and HLA-DQ are key molecules in the regulation of mDC-mediated cellular immunity by Tregs in severe aplastic anemia. J Clin Lab Anal. 2020;34(10):e23443. doi:10.1002/jcla.23443

665. Ustabas Kahraman F, Çakir FB, Buhur Pirimoglu M, Torun E, Ergen HA, Doğan Demir A. Association of Myeloperoxidase Gene Polymorphism With Iron Deficiency Anemia in Turkish Children. J Pediatr Hematol Oncol. 2021;43(7):e941–e945. doi:10.1097/MPH.0000000000002125

666. Ear J, Huang H, Wilson T, Tehrani Z, Lindgren A, Sung V, Laadem A, Daniel TO, Chopra R, Lin S. RAP-011 improves erythropoiesis in zebrafish model of Diamond-Blackfan anemia through antagonizing lefty1. Blood. 2015;126(7):880–890. doi:10.1182/blood-2015-01-622522

667. Kengne Fotsing CB, Pieme CA, Biapa Nya PC, Chedjou JP, Dabou S, Nguemeni C, Teto G, Mbacham WF, Gatsing D. Relation between haptoglobin polymorphism and oxidative stress status, lipid profile, and cardiovascular risk in sickle cell anemia patients. Health Sci Rep. 2022;5(1):e465. doi:10.1002/hsr2.465

668. Kristiansen M, Aminoff M, Jacobsen C, de La Chapelle A, Krahe R, Verroust PJ, Moestrup SK. Cubilin P1297L mutation associated with hereditary megaloblastic anemia 1 causes impaired recognition of intrinsic factor-vitamin B(12) by cubilin. Blood. 2000;96(2):405–409.

669. Fujita S, Kashiwagi H, Tomimatsu T, Ito S, Mimura K, Kanagawa T, Endo M, Miyoshi T, Okamura Y, Tani Y, et al. Expression levels of ABCG2 on cord red blood cells and study of fetal anemia associated with anti-Jr(a). Transfusion. 2016;56(5):1171–1181. doi:10.1111/trf.13515

670. Dufour C, Svahn J, Bacigalupo A, Longoni D, Varotto S, Iori AP, Bagnasco F, Locasciulli A, Menna G, Ramenghi U, et al. Genetic polymorphisms of CYP3A4, GSTT1, GSTM1, GSTP1 and NQO1 and the risk of acquired idiopathic aplastic anemia in Caucasian patients. Haematologica. 2005;90(8):1027–1031.

671. Simpson DC, Kabyemela E, Muehlenbachs A, Ogata Y, Mutabingwa TK, Duffy PE, Fried M. Plasma levels of apolipoprotein A1 in malaria-exposed primigravidae are associated with severe anemia. PLoS One. 2010;5(1):e8822. doi:10.1371/journal.pone.0008822

672. Batista JVGF, Arcanjo GS, Batista THC, Sobreira MJ, Santana RM, Domingos IF, Hatzlhofer BL, Falcão DA, Pereira-Martins DA, Oliveira JM, et al. Influence of UGT1A1 promoter polymorphism, α-thalassemia and βs haplotype in bilirubin levels and cholelithiasis in a large sickle cell anemia cohort. Ann Hematol. 2021;100(4):903–911. doi:10.1007/s00277-021-04422-1

673. Yahouédéhou SCMA, Neres JSDS, da Guarda CC, Carvalho SP, Santiago RP, Figueiredo CVB, Fiuza LM, Ndidi US, de Oliveira RM, Fonseca CA, et al. Sickle Cell Anemia: Variants in the CYP2D6, CAT, and SLC14A1 Genes Are Associated With Improved Hydroxyurea Response. Front Pharmacol. 2020;11:553064. doi:10.3389/fphar.2020.553064

674. Bouamar R, Elens L, Shuker N, van Schaik RH, Weimar W, Hesselink DA, van Gelder T. Mycophenolic acid-related anemia and leucopenia in renal transplant recipients are related to genetic polymorphisms in CYP2C8. Transplantation. 2012;93(10):e39–e42. doi:10.1097/TP.0b013e3182488bb4

675. Culpepper C, Wesolowski SR, Benjamin J, Bruce JL, Brown LD, Jonker SS, Wilkening RB, Hay WW Jr, Rozance PJ. Chronic anemic hypoxemia increases plasma glucagon and hepatic PCK1 mRNA in late-gestation fetal sheep. Am J Physiol Regul Integr Comp Physiol. 2016;311(1):R200–R208. doi:10.1152/ajpregu.00037.2016

676. Braga CCB, Benites BD, de Albuquerque DM, Alvarez MC, Seva-Pereira T, Duarte BKL, Costa FF, Gilli SCO, Saad STO. Deferasirox associated with liver failure and death in a sickle cell anemia patient homozygous for the -1774delG polymorphism in the Abcc2 gene. Clin Case Rep. 2017;5(8):1218–1221. doi:10.1002/ccr3.1040

677. Rey MA, Duffy SP, Brown JK, Kennedy JA, Dick JE, Dror Y, Tailor CS. Enhanced alternative splicing of the FLVCR1 gene in Diamond Blackfan anemia disrupts FLVCR1 expression and function that are critical for erythropoiesis. Haematologica. 2008;93(11):1617–1626. doi:10.3324/haematol.13359

678. Yaren A, Oztop I, Turgut S, Turgut G, Degirmencioglu S, Demirpence M. Angiotensin-converting enzyme gene polymorphism is associated with anemia in non small-cell lung cancer. Exp Biol Med (Maywood). 2008;233(1):32–37. doi:10.3181/0705-RM-141

679. Piemonti L, Keymeulen B, Gillard P, Linn T, Bosi E, Rose L, Pozzilli P, Giorgino F, Cossu E, Daffonchio L, et al. Ladarixin, an inhibitor of IL-8 receptors CXCR1 and CXCR2, in new-onset type 1 diabetes: a multicenter, randomized, double-blind, placebo-controlled trial. Diabetes Obes Metab. 2022;10.1111/dom.14770. doi:10.1111/dom.14770

680. Takahashi K, Ohara M, Sasai T, Homma H, Nagasawa K, Takahashi T, Yamashina M, Ishii M, Fujiwara F, Kajiwara T, et al. Serum CXCL1 concentrations are elevated in type 1 diabetes mellitus, possibly reflecting activity of anti-islet autoimmune activity. Diabetes Metab Res Rev. 2011;27(8):830–833. doi:10.1002/dmrr.1257

681. Hägglöf B, Holmgren G, Holmlund G, Lindblom B, Olaisen B, Teisberg P. Studies of HLA, factor B (Bf), complement C2 and C4 haplotypes in type 1 diabetic and control families from northern Sweden. Hum Hered. 1986;36(4):201–212. doi:10.1159/000153627

682. Peeters SA, Engelen L, Buijs J, Chaturvedi N, Fuller JH, Schalkwijk CG, Stehouwer CD; EURODIAB Prospective Complications Study Group.Plasma levels of matrix metalloproteinase-2, -3, -10, and tissue inhibitor of metalloproteinase-1 are associated with vascular complications in patients with type 1 diabetes: the EURODIAB Prospective Complications Study. Cardiovasc Diabetol. 2015;14:31. doi:10.1186/s12933-015-0195-2

683. Li YY, Gao W, Pang SS, Min XY, Yang ZJ, Wang H, Lu XZ, Wang LS, Wang XM, Qian Y, et al. TAP1 I333V gene polymorphism and type 1 diabetes mellitus: a meta-analysis of 2248 cases. J Cell Mol Med. 2014;18(5):929–937. doi:10.1111/jcmm.12244

684. Johannesen J, Pociot F, Kristiansen OP, Karlsen AE, Nerup J; DIEGG and DSGD. Danish IDDM Epidemiology and Genetics Group. Danish Study Group of Diabetis in Childhood. No evidence for linkage in the promoter region of the inducible nitric oxide synthase gene (NOS2) in a Danish type 1 diabetes population. Genes Immun. 2000;1(6):362–366. doi:10.1038/sj.gene.6363686

685. Nigi L, Brusco N, Grieco GE, Licata G, Krogvold L, Marselli L, Gysemans C, Overbergh L, Marchetti P, Mathieu C, et al. Pancreatic Alpha-Cells Contribute Together With Beta-Cells to CXCL10 Expression in Type 1 Diabetes. Front Endocrinol (Lausanne). 2020;11:630. doi:10.3389/fendo.2020.00630

686. Yang JH, Downes K, Howson JM, Nutland S, Stevens HE, Walker NM, Todd JA. Evidence of association with type 1 diabetes in the SLC11A1 gene region. BMC Med Genet. 2011;12:59. doi:10.1186/1471-2350-12-59

687. Sarkar S, Melchior JT, Henry HR, Syed F, Mirmira RG, Nakayasu ES, Metz TO. GDF15: a potential therapeutic target for type 1 diabetes. Expert Opin Ther Targets. 2022;26(1):57–67. doi:10.1080/14728222.2022.2029410

688. Abed NT, Ramadan IA, Mohammed SA, El-Shanawany EM. Genetic polymorphism of interleukin-1 receptor antagonist in Type 1 diabetic children. Pediatr Res. 2022;91(6):1536–1541. doi:10.1038/s41390-021-01569-5

689. Tomihira M, Kawasaki E, Nakajima H, Imamura Y, Sato Y, Sata M, Kage M, Sugie H, Nunoi K.Intermittent and recurrent hepatomegaly due to glycogen storage in a patient with type 1 diabetes: genetic analysis of the liver glycogen phosphorylase gene (PYGL). Diabetes Res Clin Pract. 2004;65(2):175–182. doi:10.1016/j.diabres.2003.12.004

690. Hussein AG, Mohamed RH, Alghobashy AA. Synergism of CYP2R1 and CYP27B1 polymorphisms and susceptibility to type 1 diabetes in Egyptian children. Cell Immunol. 2012;279(1):42–45. doi:10.1016/j.cellimm.2012.08.006

691. Mori H, Shichita T, Yu Q, Yoshida R, Hashimoto M, Okamoto F, Torisu T, Nakaya M, Kobayashi T, Takaesu G, et al. Suppression of SOCS3 expression in the pancreatic beta-cell leads to resistance to type 1 diabetes. Biochem Biophys Res Commun. 2007;359(4):952–958. doi:10.1016/j.bbrc.2007.05.198

692. Irie J, Reck B, Wu Y, Wicker LS, Howlett S, Rainbow D, Feingold E, Ridgway WM. Genome-wide microarray expression analysis of CD4+ T Cells from nonobese diabetic congenic mice identifies Cd55 (Daf1) and Acadl as candidate genes for type 1 diabetes. J Immunol. 2008;180(2):1071–1079. doi:10.4049/jimmunol.180.2.1071

693. Zhong X, Zhao X, Zhang L, Liu N, Shi S, Wang Y. Sodium hydrosulfide inhibiting endothelial cells injury and neutrophils activation via IL-8/CXCR2/ROS/NF-κB axis in type 1 diabetes mellitus rat. Biochem Biophys Res Commun. 2022;606:1–9. doi:10.1016/j.bbrc.2022.03.072

694. Huang G, Mo X, Li M, Xiang Y, Li X, Luo S, Zhou Z. Autoantibodies to CCL3 are of low sensitivity and specificity for the diagnosis of type 1 diabetes. Acta Diabetol. 2012;49(5):395–399. doi:10.1007/s00592-012-0380-7

695. Guan R, Purohit S, Wang H, Bode B, Reed JC, Steed RD, Anderson SW, Steed L, Hopkins D, Xia C, et al. Chemokine (C-C motif) ligand 2 (CCL2) in sera of patients with type 1 diabetes and diabetic complications. PLoS One. 2011;6(4):e17822. doi:10.1371/journal.pone.0017822

696. Wegner M, Mostowska A, Araszkiewicz A, Choudhury M, Piorunska-Stolzmann M, Zozulinska-Ziolkiewicz D, Wierusz-Wysocka B, Jagodzinski PP. Association investigation of BACH2 rs3757247 and SOD2 rs4880 polymorphisms with the type 1 diabetes and diabetes long-term complications risk in the Polish population. Biomed Rep. 2015;3(3):327–332. doi:10.3892/br.2015.424

697. Dezsofi A, Szebeni B, Hermann CS, Kapitány A, Veres G, Sipka S, Körner A, Madácsy L, Korponay-Szabó I, Rajczy K, et al. Frequencies of genetic polymorphisms of TLR4 and CD14 and of HLA-DQ genotypes in children with celiac disease, type 1 diabetes mellitus, or both. J Pediatr Gastroenterol Nutr. 2008;47(3):283–287. doi:10.1097/MPG.0b013e31816de885

698. Bereket A, Lang CH, Blethen SL, Wilson TA. Insulin-like growth factor-binding protein-2 and insulin: studies in children with type 1 diabetes mellitus and maturity-onset diabetes of the young. J Clin Endocrinol Metab. 1995;80(12):3647–3652. doi:10.1210/jcem.80.12.8530614

699. Bojanin D, Vekic J, Milenkovic T, Vukovic R, Zeljkovic A, Stefanovic A, Janac J, Ivanisevic J, Mitrovic K, Miljkovic M, et al. Association between proprotein convertase subtilisin/kexin 9 (PCSK9) and lipoprotein subclasses in children with type 1 diabetes mellitus: Effects of glycemic control. Atherosclerosis. 2019;280:14–20. doi:10.1016/j.atherosclerosis.2018.11.020

700. Fang C, Huang Y, Pei Y, Zhang HH, Chen X, Guo H, Li S, Ji X, Hu J. Genome-wide gene expression profiling reveals that CD274 is up-regulated new-onset type 1 diabetes mellitus. Acta Diabetol. 2017;54(8):757–767. doi:10.1007/s00592-017-1005-y

701. Anquetil F, Mondanelli G, Gonzalez N, Rodriguez Calvo T, Zapardiel Gonzalo J, Krogvold L, Dahl-Jørgensen K, Van den Eynde B, Orabona C, Grohmann U, et al. Loss of IDO1 Expression From Human Pancreatic β-Cells Precedes Their Destruction During the Development of Type 1 Diabetes. Diabetes. 2018;67(9):1858–1866. doi:10.2337/db17-1281

702. Wang Z, Zheng Y, Hou C, Yang L, Li X, Lin J, Huang G, Lu Q, Wang CY, Zhou Z. DNA methylation impairs TLR9 induced Foxp3 expression by attenuating IRF-7 binding activity in fulminant type 1 diabetes. J Autoimmun. 2013;41:50–59. doi:10.1016/j.jaut.2013.01.009

703. Melin EO, Dereke J, Hillman M. Female sex, high soluble CD163, and low HDL-cholesterol were associated with high galectin-3 binding protein in type 1 diabetes. Biol Sex Differ. 2019;10(1):51. doi:10.1186/s13293-019-0268-0

704. Calderon RM, Diaz S, Szeto A, Llinas JA, Hughes TA, Mendez AJ, Goldberg RB.Elevated Lipoprotein Lipase Activity Does Not Account for the Association Between Adiponectin and HDL in Type 1 Diabetes. J Clin Endocrinol Metab. 2015;100(7):2581–2588. doi:10.1210/jc.2015-1357

705. Demirci M, Bahar Tokman H, Taner Z, Keskin FE, Çağatay P, Ozturk Bakar Y, Özyazar M, Kiraz N, Kocazeybek BS. Bacteroidetes and Firmicutes levels in gut microbiota and effects of hosts TLR2/TLR4 gene expression levels in adult type 1 diabetes patients in Istanbul, Turkey. J Diabetes Complications. 2020;34(2):107449. doi:10.1016/j.jdiacomp.2019.107449

706. Gambelunghe G, Ghaderi M, Brozzetti A, Del Sindaco P, Gharizadeh B, Nyren P, Hjelmström P, Nikitina-Zake L, Sanjeevi CB, Falorni A. Lack of association of CCR2-64I and CCR5-Delta 32 with type 1 diabetes and latent autoimmune diabetes in adults. Hum Immunol. 2003;64(6):629–632. doi:10.1016/s0198-8859(03)00064-8

707. Chan NN, Fuller JH, Rubens M, Colhoun HM. Von Willebrand factor in type 1 diabetes: its production and coronary artery calcification. Med Sci Monit. 2003;9(7):CR297–CR303.

708. Li YY, Pearson JA, Chao C, Peng J, Zhang X, Zhou Z, Liu Y, Wong FS, Wen L. Nucleotide-binding oligomerization domain-containing protein 2 (Nod2) modulates T1DM susceptibility by gut microbiota. J Autoimmun. 2017;82:85–95. doi:10.1016/j.jaut.2017.05.00

709. Badr G, Sayed LH, Omar HEM, Abd El-Rahim AM, Ahmed EA, Mahmoud MH. Camel Whey Protein Protects B and T Cells from Apoptosis by Suppressing Activating Transcription Factor-3 (ATF-3)-Mediated Oxidative Stress and Enhancing Phosphorylation of AKT and IκB-α in Type I Diabetic Mice. Cell Physiol Biochem. 2017;41(1):41–54. doi:10.1159/000455935

710. Heilman K, Zilmer M, Zilmer K, Tillmann V. Lower bone mineral density in children with type 1 diabetes is associated with poor glycemic control and higher serum ICAM-1 and urinary isoprostane levels. J Bone Miner Metab. 2009;27(5):598–604. doi:10.1007/s00774-009-0076-4

711. Traisaeng S, Batsukh A, Chuang TH, Herr DR, Huang YF, Chimeddorj B, Huang CM.Leuconostoc mesenteroides fermentation produces butyric acid and mediates Ffar2 to regulate blood glucose and insulin in type 1 diabetic mice. Sci Rep. 2020;10(1):7928. doi:10.1038/s41598-020-64916-2

712. Jahromi M, Millward A, Demaine A. A CA repeat polymorphism of the IFN-gamma gene is associated with susceptibility to type 1 diabetes. J Interferon Cytokine Res. 2000;20(2):187–190. doi:10.1089/107999000312595

713. Atabek ME, Özkul Y, Eklioğlu BS, Kurtoğlu S, Baykara M. Association between apolipoprotein E polymorphism and subclinic atherosclerosis in patients with type 1 diabetes mellitus. J Clin Res Pediatr Endocrinol. 2012;4(1):8–13. doi:10.4274/jcrpe.521

714. Purohit S, Sharma A, Zhi W, Bai S, Hopkins D, Steed L, Bode B, Anderson SW, Reed JC, Steed RD, et al. Proteins of TNF-α and IL6 Pathways Are Elevated in Serum of Type-1 Diabetes Patients with Microalbuminuria. Front Immunol. 2018;9:154. doi:10.3389/fimmu.2018.00154

715. Meagher C, Beilke J, Arreaza G, Mi QS, Chen W, Salojin K, Horst N, Cruikshank WW, Delovitch TL. Neutralization of interleukin-16 protects nonobese diabetic mice from autoimmune type 1 diabetes by a CCL4-dependent mechanism. Diabetes. 2010;59(11):2862–2871. doi:10.2337/db09-0131

716. Taniguchi T, Okazaki K, Okamoto M, Seko S, Tanaka J, Uchida K, Nagashima K, Kurose T, Yamada Y, Chiba T, et al. High prevalence of autoantibodies against carbonic anhydrase II and lactoferrin in type 1 diabetes: concept of autoimmune exocrinopathy and endocrinopathy of the pancreas. Pancreas. 2003;27(1):26–30. doi:10.1097/00006676-200307000-00004

717. Bister V, Kolho KL, Karikoski R, Westerholm-Ormio M, Savilahti E, Saarialho-Kere U. Metalloelastase (MMP-12) is upregulated in the gut of pediatric patients with potential celiac disease and in type 1 diabetes. Scand J Gastroenterol. 2005;40(12):1413–1422. doi:10.1080/00365520510023918

718. Symeonidis C, Papakonstantinou E, Galli A, Tsinopoulos I, Mataftsi A, Batzios S, Dimitrakos SA. Matrix metalloproteinase (MMP-2, -9) and tissue inhibitor (TIMP-1, -2) activity in tear samples of pediatric type 1 diabetic patients: MMPs in tear samples from type 1 diabetes. Graefes Arch Clin Exp Ophthalmol. 2013;251(3):741–749. doi:10.1007/s00417-012-2221-3

719. Cantor J, Haskins K. Recruitment and activation of macrophages by pathogenic CD4 T cells in type 1 diabetes: evidence for involvement of CCR8 and CCL1. J Immunol. 2007;179(9):5760–5767. doi:10.4049/jimmunol.179.9.5760

720. Wang Y, Qin Y, Wang X, Zhang L, Wang J, Xu X, Chen H, Hsu HT, Zhang M. Decrease in the proportion of CD24hi CD38hi B cells and impairment of their regulatory capacity in type 1 diabetes patients. Clin Exp Immunol. 2020;200(1):22–32. doi:10.1111/cei.13408

721. Rotbain Curovic V, Theilade S, Winther SA, Tofte N, Eugen-Olsen J, Persson F, Hansen TW, Jeppesen J, Rossing P. Soluble Urokinase Plasminogen Activator Receptor Predicts Cardiovascular Events, Kidney Function Decline, and Mortality in Patients With Type 1 Diabetes. Diabetes Care. 2019;42(6):1112–1119. doi:10.2337/dc18-1427

722. Mosaad YM, Elsharkawy AA, El-Deek BS. Association of CTLA-4 (+49A/G) gene polymorphism with type 1 diabetes mellitus in Egyptian children. Immunol Invest. 2012;41(1):28–37. doi:10.3109/08820139.2011.579215

723. Jenny L, Ajjan R, King R, Thiel S, Schroeder V. Plasma levels of mannan-binding lectin-associated serine proteases MASP-1 and MASP-2 are elevated in type 1 diabetes and correlate with glycaemic control. Clin Exp Immunol. 2015;180(2):227–232. doi:10.1111/cei.12574

724. Heilman K, Zilmer M, Zilmer K, Lintrop M, Kampus P, Kals J, Tillmann V. Arterial stiffness, carotid artery intima-media thickness and plasma myeloperoxidase level in children with type 1 diabetes. Diabetes Res Clin Pract. 2009;84(2):168–173. doi:10.1016/j.diabres.2009.01.014

725. Costacou T, Orchard TJ. The Haptoglobin genotype predicts cardio-renal mortality in type 1 diabetes. J Diabetes Complications. 2016;30(2):221–226. doi:10.1016/j.jdiacomp.2015.11.011

726. Toni M, Hermida J, Goñi MJ, Fernández P, Parks WC, Toledo E, Montes R, Díez N. Matrix metalloproteinase-10 plays an active role in microvascular complications in type 1 diabetic patients. Diabetologia. 2013;56(12):2743–2752. doi:10.1007/s00125-013-3052-4

727. Haseda F, Imagawa A, Nishikawa H, Mitsui S, Tsutsumi C, Fujisawa R, Sano H, Murase-Mishiba Y, Terasaki J, Sakaguchi S, et al. Antibody to CMRF35-Like Molecule 2, CD300e A Novel Biomarker Detected in Patients with Fulminant Type 1 Diabetes. PLoS One. 2016;11(8):e0160576. doi:10.1371/journal.pone.0160576

728. Thrailkill KM, Nimmo T, Bunn RC, Cockrell GE, Moreau CS, Mackintosh S, Edmondson RD, Fowlkes JL. Microalbuminuria in type 1 diabetes is associated with enhanced excretion of the endocytic multiligand receptors megalin and cubilin. Diabetes Care. 2009;32(7):1266–1268. doi:10.2337/dc09-0112

729. Porta M, Toppila I, Sandholm N, Hosseini SM, Forsblom C, Hietala K, Borio L, Harjutsalo V, Klein BE, Klein R, et al. Variation in SLC19A3 and Protection From Microvascular Damage in Type 1 Diabetes. Diabetes. 2016;65(4):1022–1030. doi:10.2337/db15-1247

730. Bjornstad P, Eckel RH, Pyle L, Rewers M, Maahs DM, Snell-Bergeon JK. Relation of Combined Non-High-Density Lipoprotein Cholesterol and Apolipoprotein B With Atherosclerosis in Adults With Type 1 Diabetes Mellitus. Am J Cardiol. 2015;116(7):1057–1062. doi:10.1016/j.amjcard.2015.07.020

731. Valladolid-Acebes I, Berggren PO, Juntti-Berggren L. Apolipoprotein CIII Is an Important Piece in the Type-1 Diabetes Jigsaw Puzzle. Int J Mol Sci. 2021;22(2):932. doi:10.3390/ijms22020932

732. Shukla SK, Liu W, Sikder K, Addya S, Sarkar A, Wei Y, Rafiq K.HMGCS2 is a key ketogenic enzyme potentially involved in type 1 diabetes with high cardiovascular risk. Sci Rep. 2017;7(1):4590. doi:10.1038/s41598-017-04469-z

733. Grünert SC, Villavicencio-Lorini P, Wermuth B, Lehnert W, Sass JO, Schwab KO. Ornithine transcarbamylase deficiency combined with type 1 diabetes mellitus - a challenge in clinical and dietary management. J Diabetes Metab Disord. 2013;12(1):37. doi:10.1186/2251-6581-12-37

734. Cherney DZ, Xiao F, Zimpelmann J, Har RL, Lai V, Scholey JW, Reich HN, Burns KD. Urinary ACE2 in healthy adults and patients with uncomplicated type 1 diabetes. Can J Physiol Pharmacol. 2014;92(8):703–706. doi:10.1139/cjpp-2014-0065

735. Santiago JL, Martínez A, de la Calle H, Fernández-Arquero M, Figueredo MA, de la Concha EG, Urcelay E. Evidence for the association of the SLC22A4 and SLC22A5 genes with type 1 diabetes: a case control study. BMC Med Genet. 2006;7:54. doi:10.1186/1471-2350-7-54

736. Davis H, Jones Briscoe V, Dumbadze S, Davis SN. Using DPP-4 inhibitors to modulate beta cell function in type 1 diabetes and in the treatment of diabetic kidney disease. Expert Opin Investig Drugs. 2019;28(4):377–388. doi:10.1080/13543784.2019.1592156

737. Pedersen-Bjergaard U, Agerholm-Larsen B, Pramming S, Hougaard P, Thorsteinsson B. Prediction of severe hypoglycaemia by angiotensin-converting enzyme activity and genotype in type 1 diabetes. Diabetologia. 2003;46(1):89–96. doi:10.1007/s00125-002-0969-4

738. Waters MF, Delghingaro-Augusto V, Javed K, Dahlstrom JE, Burgio G, Bröer S, Nolan CJ. Knockout of the Amino Acid Transporter SLC6A19 and Autoimmune Diabetes Incidence in Female Non-Obese Diabetic (NOD) Mice. Metabolites. 2021;11(10):665. doi:10.3390/metabo11100665

739. Orozco G, Eerligh P, Sánchez E, Zhernakova S, Roep BO, González-Gay MA, López-Nevot MA, Callejas JL, Hidalgo C, Pascual-Salcedo D, et al. Analysis of a functional BTNL2 polymorphism in type 1 diabetes, rheumatoid arthritis, and systemic lupus erythematosus. Hum Immunol. 2005;66(12):1235–1241. doi:10.1016/j.humimm.2006.02.003

740. Ma SG, Yang LX, Qiu XQ. Assessment of the platelet parameters and serum butyrylcholinesterase activity in type 1 diabetes patients with ketoacidosis. Platelets. 2013;24(7):544–548. doi:10.3109/09537104.2012.735720

741. Safi M, Borup A, Stevns Hansen C, Rossing P, Thorsten Jensen M, Christoffersen C. Association between plasma apolipoprotein M and cardiac autonomic neuropathy in type 1 diabetes. Diabetes Res Clin Pract. 2022;189:109943. doi:10.1016/j.diabres.2022.109943

742. Fernández-Cadenas I, Del Río-Espínola A, Giralt D, Domingues-Montanari S, Quiroga A, Mendióroz M, Ruíz A, Ribó M, Serena J, Obach V, et al. IL1B and VWF variants are associated with fibrinolytic early recanalization in patients with ischemic stroke. Stroke. 2012;43(10):2659–2665. doi:10.1161/STROKEAHA.112.657007

743. Buil A, Trégouët DA, Souto JC, Saut N, Germain M, Rotival M, Tiret L, Cambien F, Lathrop M, Zeller T, et al. C4BPB/C4BPA is a new susceptibility locus for venous thrombosis with unknown protein S-independent mechanism: results from genome-wide association and gene expression analyses followed by case-control studies. Blood. 2010;115(23):4644–4650. doi:10.1182/blood-2010-01-263038

744. Sugano T, Tsuji H, Masuda H, Nishimura H, Yoshizumi M, Kawano H, Kimura S, Ukimura N, Yano S, Kunieda Y, et al. Adrenomedullin inhibits angiotensin II-induced expression of tissue factor and plasminogen activator inhibitor-1 in cultured rat aortic endothelial cells. Arterioscler Thromb Vasc Biol. 2001;21(6):1078–1083. doi:10.1161/01.atv.21.6.1078

745. Matusik PT, Małecka B, Lelakowski J, Undas A. Association of NT-proBNP and GDF-15 with markers of a prothrombotic state in patients with atrial fibrillation off anticoagulation. Clin Res Cardiol. 2020;109(4):426–434. doi:10.1007/s00392-019-01522-x

746. van Minkelen R, de Visser MC, Houwing-Duistermaat JJ, Vos HL, Bertina RM, Rosendaal FR. Haplotypes of IL1B, IL1RN, IL1R1, and IL1R2 and the risk of venous thrombosis. Arterioscler Thromb Vasc Biol. 2007;27(6):1486–1491. doi:10.1161/ATVBAHA.107.140384

747. Jerotic D, Ranin J, Bukumiric Z, Djukic T, Coric V, Savic-Radojevic A, Todorovic N, Asanin M, Ercegovac M, Milosevic I, et al. SOD2 rs4880 and GPX1 rs1050450 polymorphisms do not confer risk of COVID-19, but influence inflammation or coagulation parameters in Serbian cohort. Redox Rep. 2022;27(1):85–91. doi:10.1080/13510002.2022.2057707

748. Shin HS, Xu F, Bagchi A, Herrup E, Prakash A, Valentine C, Kulkarni H, Wilhelmsen K, Warren S, Hellman J. Bacterial lipoprotein TLR2 agonists broadly modulate endothelial function and coagulation pathways in vitro and in vivo. J Immunol. 2011;186(2):1119–1130. doi:10.4049/jimmunol.1001647

749. Bombeli T, Jutzi M, De Conno E, Seifert B, Fehr J. In patients with deep-vein thrombosis elevated levels of factor VIII correlate only with von Willebrand factor but not other endothelial cell-derived coagulation and fibrinolysis proteins. Blood Coagul Fibrinolysis. 2002;13(7):577–581. doi:10.1097/00001721-200210000-00001

750. Beppu T, Gil-Bernabe P, Boveda-Ruiz D, D’Alessandro-Gabazza C, Matsuda Y, Toda M, Miyake Y, Shiraki K, Murata M, Murata T, et al. High incidence of tumors in diabetic thrombin activatable fibrinolysis inhibitor and apolipoprotein E double-deficient mice. J Thromb Haemost. 2010;8(11):2514–2522. doi:10.1111/j.1538-7836.2010.04023.x

751. Stouthard JM, Levi M, Hack CE, Veenhof CH, Romijn HA, Sauerwein HP, van der Poll T. Interleukin-6 stimulates coagulation, not fibrinolysis, in humans. Thromb Haemost. 1996;76(5):738–742.

752. Arıcı OF, Cetin N. Protective role of ghrelin against carbon tetrachloride (CCl□)-induced coagulation disturbances in rats. Regul Pept. 2011;166(1-3):139–142. doi:10.1016/j.regpep.2010.10.009

753. Zhang Y, Chen M, Zhang Y, Peng P, Li J, Xin X. miR-96 and miR-330 overexpressed and targeted AQP5 in lipopolysaccharide-induced rat lung damage of disseminated intravascular coagulation. Blood Coagul Fibrinolysis. 2014;25(7):731–737. doi:10.1097/MBC.0000000000000133

754. Garabet L, Ghanima W, Monceyron Jonassen C, Skov V, Holst R, Mowinckel MC, C Hasselbalch H, A Kruse T, Thomassen M, Liebman H, et al. Effect of thrombopoietin receptor agonists on markers of coagulation and P-selectin in patients with immune thrombocytopenia. Platelets. 2019;30(2):206–212. doi:10.1080/09537104.2017.1394451

755. Shenkman B, Livnat T, Budnik I, Tamarin I, Einav Y, Martinowitz U. Plasma tissue-type plasminogen activator increases fibrinolytic activity of exogenous urokinase-type plasminogen activator. Blood Coagul Fibrinolysis. 2012;23(8):729–733. doi:10.1097/MBC.0b013e32835897d5

756. Dobó J, Schroeder V, Jenny L, Cervenak L, Závodszky P, Gál P. Multiple roles of complement MASP-1 at the interface of innate immune response and coagulation. Mol Immunol. 2014;61(2):69–78. doi:10.1016/j.molimm.2014.05.013

757. Misztal T, Golaszewska A, Tomasiak-Lozowska MM, Iwanicka M, Marcinczyk N, Leszczynska A, Chabielska E, Rusak T.The myeloperoxidase product, hypochlorous acid, reduces thrombus formation under flow and attenuates clot retraction and fibrinolysis in human blood. Free Radic Biol Med. 2019;141:426–437. doi:10.1016/j.freeradbiomed.2019.07.003

758. Navarro-Oviedo M, Roncal C, Salicio A, Belzunce M, Rabal O, Toledo E, Zandio B, Rodríguez JA, Páramo JA, Muñoz R, et al. MMP10 Promotes Efficient Thrombolysis After Ischemic Stroke in Mice with Induced Diabetes. Transl Stroke Res. 2019;10(4):389–401. doi:10.1007/s12975-018-0652-9

759. Sa Q, Hoover-Plow JL. EMILIN2 (Elastin microfibril interface located protein), potential modifier of thrombosis. Thromb J. 2011;9:9. doi:10.1186/1477-9560-9-9

760. Gemmati D, Bramanti B, Serino ML, Secchiero P, Zauli G, Tisato V. COVID-19 and Individual Genetic Susceptibility/Receptivity: Role of ACE1/ACE2 Genes, Immunity, Inflammation and Coagulation. Might the Double X-chromosome in Females Be Protective against SARS-CoV-2 Compared to the Single X-Chromosome in Males?. Int J Mol Sci. 2020;21(10):3474. doi:10.3390/ijms21103474

761. Erem C. Blood coagulation, fibrinolytic activity and lipid profile in subclinical thyroid disease: subclinical hyperthyroidism increases plasma factor X activity. Clin Endocrinol (Oxf). 2006;64(3):323–329. doi:10.1111/j.1365-2265.2006.02464.x

762. Ay C, Bencur P, Vormittag R, et al. The angiotensin-converting enzyme insertion/deletion polymorphism and serum levels of angiotensin-converting enzyme in venous thromboembolism. Data from a case control study. Thromb Haemost. 2007;98(4):777–782.

763. Meijers JC, Tekelenburg WL, Bouma BN, Bertina RM, Rosendaal FR. High levels of coagulation factor XI as a risk factor for venous thrombosis. N Engl J Med. 2000;342(10):696–701. doi:10.1056/NEJM200003093421004

764. Ahmad A, Sundquist K, Zöller B, Dahlbäck B, Elf J, Svensson PJ, Strandberg K, Sundquist J, Memon AA. Evaluation of Expression Level of Apolipoprotein M as a Diagnostic Marker for Primary Venous Thromboembolism. Clin Appl Thromb Hemost. 2018;24(3):416–422. doi:10.1177/1076029617730639

765. Hrašovec S, Hauptman N, Glavač D, Jelenc F, Ravnik-Glavač M. TMEM25 is a candidate biomarker methylated and down-regulated in colorectal cancer. Dis Markers. 2013;34(2):93–104. doi:10.3233/DMA-120948

766. Rankin CR, Lokhandwala ZA, Huang R, Pekow J, Pothoulakis C, Padua D. Linear and circular CDKN2B-AS1 expression is associated with Inflammatory Bowel Disease and participates in intestinal barrier formation. Life Sci. 2019;231:116571. doi:10.1016/j.lfs.2019.116571

767. Verstockt S, De Hertogh G, Van der Goten J, Verstockt B, Vancamelbeke M, Machiels K, Van Lommel L, Schuit F, Van Assche G, Rutgeerts P, et al. Gene and Mirna Regulatory Networks During Different Stages of Crohn’s Disease. J Crohns Colitis. 2019;13(7):916–930. doi:10.1093/ecco-jcc/jjz007

768. Bai J, Li Y, Shao T, Zhao Z, Wang Y, Wu A, Chen H, Li S, Jiang C, Xu J, Li X. Integrating analysis reveals microRNA-mediated pathway crosstalk among Crohn’s disease, ulcerative colitis and colorectal cancer. Mol Biosyst. 2014;10(9):2317–2328. doi:10.1039/c4mb00169a

769. Krela-Kaźmierczak I, Skrzypczak-Zielińska M, Kaczmarek-Ryś M, Michalak M, Szymczak-Tomczak A, Hryhorowicz ST, Szalata M, Łykowska-Szuber L, Eder P, Stawczyk-Eder K, et al. ESR1 Gene Variants Are Predictive of Osteoporosis in Female Patients with Crohn’s Disease. J Clin Med. 2019;8(9):1306. doi:10.3390/jcm8091306

770. Ellinghaus D, Zhang H, Zeissig S, Lipinski S, Till A, Jiang T, Stade B, Bromberg Y, Ellinghaus E, Keller A, et al. Association between variants of PRDM1 and NDP52 and Crohn’s disease, based on exome sequencing and functional studies. Gastroenterology. 2013;145(2):339–347. doi:10.1053/j.gastro.2013.04.040

771. Li P, Zhang HY, Gao JZ, Du WQ, Tang D, Wang W, Wang LH. Mesenchymal stem cells-derived extracellular vesicles containing miR-378a-3p inhibit the occurrence of inflammatory bowel disease by targeting GATA2. J Cell Mol Med. 2022;26(11):3133–3146. doi:10.1111/jcmm.17176

772. Tian Y, Cui L, Lin C, Wang Y, Liu Z, Miao X. LncRNA CDKN2B-AS1 relieved inflammation of ulcerative colitis via sponging miR-16 and miR-195. Int Immunopharmacol. 2020;88:106970. doi:10.1016/j.intimp.2020.106970

773. Arasaradnam RP, Khoo K, Bradburn M, Mathers JC, Kelly SB. DNA methylation of ESR-1 and N-33 in colorectal mucosa of patients with ulcerative colitis (UC). Epigenetics. 2010;5(5):422–426. doi:10.4161/epi.5.5.11959

774. Yang L, Ma DW, Cao YP, Li DZ, Zhou X, Feng JF, Bao J. PRMT5 functionally associates with EZH2 to promote colorectal cancer progression through epigenetically repressing CDKN2B expression. Theranostics. 2021;11(8):3742–3759. doi:10.7150/thno.53023

775. Jung JH, Shin EA, Kim JH, Sim DY, Lee H, Park JE, Lee HJ, Kim SH.NEDD9 Inhibition by miR-25-5p Activation Is Critically Involved in Co-Treatment of Melatonin- and Pterostilbene-Induced Apoptosis in Colorectal Cancer Cells. Cancers (Basel). 2019;11(11):1684. doi:10.3390/cancers11111684

776. Kara M, Yumrutas O, Ozcan O, Celik OI, Bozgeyik E, Bozgeyik I, Tasdemir S. Differential expressions of cancer-associated genes and their regulatory miRNAs in colorectal carcinoma. Gene. 2015;567(1):81–86. doi:10.1016/j.gene.2015.04.065

777. Han F, Zhang L, Liao S, Zhang Y, Qian L, Hou F, Gong J, Lai M, Zhang H. The interaction between S100A2 and KPNA2 mediates NFYA nuclear import and is a novel therapeutic target for colorectal cancer metastasis. Oncogene. 2022;41(5):657–670. doi:10.1038/s41388-021-02116-6

778. Lazar SB, Pongor L, Li XL, Grammatikakis I, Muys BR, Dangelmaier EA, Redon CE, Jang SM, Walker RL, Tang W, et al. Genome-Wide Analysis of the FOXA1 Transcriptional Network Identifies Novel Protein-Coding and Long Noncoding RNA Targets in Colorectal Cancer Cells. Mol Cell Biol. 2020;40(21):e00224–20. doi:10.1128/MCB.00224-20

779. Ye SB, Cheng YK, Deng R, Deng Y, Li P, Zhang L, Lan P. The Predictive Value of Estrogen Receptor 1 on Adjuvant Chemotherapy in Locally Advanced Colorectal Cancer: A Retrospective Analysis With Independent Validation and Its Potential Mechanism. Front Oncol. 2020;10:214. doi:10.3389/fonc.2020.00214

780. Tang J, Yang L, Li Y, Ning X, Chaulagain A, Wang T, Wang D. ARID3A promotes the development of colorectal cancer by upregulating AURKA. Carcinogenesis. 2021;42(4):578–586. doi:10.1093/carcin/bgaa118

781. Chai B, Guo Y, Cui X, Liu J, Suo Y, Dou Z, Li N. MiR-223-3p promotes the proliferation, invasion and migration of colon cancer cells by negative regulating PRDM1. Am J Transl Res. 2019;11(7):4516–4523.

782. Pan Y, Zhu Y, Zhang J, Jin L, Cao P. A feedback loop between GATA2-AS1 and GATA2 promotes colorectal cancer cell proliferation, invasion, epithelial-mesenchymal transition and stemness via recruiting DDX3X. J Transl Med. 2022;20(1):287. doi:10.1186/s12967-022-03483-8

783. Yaiche H, Tounsi-Kettiti H, Ben Jemii N, Jaballah Gabteni A, Mezghanni N, Ardhaoui M, Fehri E, Maaloul A, Abdelhak S, Boubaker S. New insights in the clinical implication of HOXA5 as prognostic biomarker in patients with colorectal cancer. Cancer Biomark. 2021;30(2):213–221. doi:10.3233/CBM-201758

784. Leaf S, Carlsen L, El-Deiry WS. Opposing effects of BRCA1 mRNA expression on patient survival in breast and colorectal cancer and variations among African American, Asian, and younger patients. Oncotarget. 2021;12(20):1992–2005. doi:10.18632/oncotarget.28082

785. Cagnan I, Keles M, Keskus AG, Tombaz M, Sahan OB, Aerts-Kaya F, Uckan-Cetinkaya D, Konu O, Gunel-Ozcan A. Global miRNA expression of bone marrow mesenchymal stem/stromal cells derived from Fanconi anemia patients. Hum Cell. 2022;35(1):111–124. doi:10.1007/s13577-021-00626-9

786. Ganapathi KA, Townsley DM, Hsu AP, Arthur DC, Zerbe CS, Cuellar-Rodriguez J, Hickstein DD, Rosenzweig SD, Braylan RC, Young NS, et al. GATA2 deficiency-associated bone marrow disorder differs from idiopathic aplastic anemia. Blood. 2015;125(1):56–70. doi:10.1182/blood-2014-06-580340

787. Freire BL, Homma TK, Funari MFA, Lerario AM, Leal AM, Velloso EDRP, Malaquias AC, Jorge AAL. Homozygous loss of function BRCA1 variant causing a Fanconi-anemia-like phenotype, a clinical report and review of previous patients. Eur J Med Genet. 2018;61(3):130–133. doi:10.1016/j.ejmg.2017.11.003

788. Samandari N, Mirza AH, Nielsen LB, Kaur S, Hougaard P, Fredheim S, Mortensen HB, Pociot F. Circulating microRNA levels predict residual beta cell function and glycaemic control in children with type 1 diabetes mellitus. Diabetologia. 2017;60(2):354–363. doi:10.1007/s00125-016-4156-4

789. Mirza AH, Kaur S, Nielsen LB, Størling J, Yarani R, Roursgaard M, Mathiesen ER, Damm P, Svare J, Mortensen HB, et al. Breast Milk-Derived Extracellular Vesicles Enriched in Exosomes From Mothers With Type 1 Diabetes Contain Aberrant Levels of microRNAs. Front Immunol. 2019;10:2543. doi:10.3389/fimmu.2019.02543

790. Gabler J, Basílio J, Steinbrecher O, Kollars M, Kyrle PA, Eichinger S. MicroRNA Signatures in Plasma of Patients With Venous Thrombosis: A Preliminary Report. Am J Med Sci. 2021;361(4):509–516. doi:10.1016/j.amjms.2020.12.002

791. Roberts NA, Adams BD, McCarthy NI, Tooze RM, Parnell SM, Anderson G, Kaech SM, Horsley V. Prdm1 Regulates Thymic Epithelial Function To Prevent Autoimmunity. J Immunol. 2017;199(4):1250–1260. doi:10.4049/jimmunol.1600941

792. Liu Z, Qing P, Zhao Y, Liu Y, Marion TN. Combined Mutation of the GATA2 Gene and STAT5B Gene in a Patient with Hypogammaglobulinemia and Autoimmunity. Tohoku J Exp Med. 2021;255(2):143–146. doi:10.1620/tjem.255.143

